# Refining physico-chemical rules for herbicides using an antimalarial library

**DOI:** 10.1101/2020.10.27.356576

**Authors:** Kirill V. Sukhoverkov, Maxime G. Corral, Julie Leroux, Joel Haywood, Philipp Johnen, Trevor Newton, Keith A. Stubbs, Joshua S. Mylne

## Abstract

Successful herbicides, like drugs, have physico-chemical properties that usually fall within certain limits. A recent analysis of 334 herbicides showed similar properties to the ‘rule of five’ for human orally-delivered drugs, but herbicides diverged from this for proton donors, partition coefficients and molecular weight. To refine rules for herbicides, we exploited the close evolutionary relationship between *P. falciparum* and plants by screening the Malaria Box, a 400-compound library composed of novel chemical scaffolds with activity against blood stage malaria parasite *Plasmodium falciparum*. A high proportion (52 of 400) were herbicidal to *Arabidopsis thaliana* on agar plates. Thirty-nine of these 52 herbicidal compounds were tested on soil and 16 compounds were herbicidal. These data were used to predict whether a herbicidal hit found on agar will work on soil-grown plants. The physico-chemical parameters were weighted to logP and formal charge and used to generate weighted scores to a large chemical library of liver-stage effective antimalarial leads. Of the six top-scoring compounds, one had a potency comparable to commercial herbicides. This novel compound MMV1206386 had no close structural matches among commercial herbicides. Physiological profiling suggested that MMV1206386 has a new mode of action and overall demonstrates how weighted rules can help during herbicide discovery programs.

## Introduction

The implementation of herbicides in agriculture in the 1940s improved crop productivity, but the emergence of herbicide resistance in the last few decades threatens those gains in yield. The first case of herbicide resistance, triazine-resistant *Senecio vulgaris*, was documented in 1968 ^1^ and since then, the number of weed species said to be resistant to one or more herbicides is now 262 ^2^. Although practices such as herbicide rotation helps avoid the evolution of resistance and extends the life of herbicides, it relies on switching between modes of action ^3^. From the 1950s to the 1970s a new mode of action was introduced every two to three years, but this slowed in the 1980s. During the next 30 years no new herbicide mode of action was commercialised until tetflupyrolimet (inhibitor of plant dihydroorotate dehydrogenase) was announced by FMC Agricultural Solutions in 2019 ^4, 5^. Two old herbicides, cinmethylin and aclonifen, were rediscovered as having new modes of action ^6, 7^, while new compounds with new modes of action have emerged, such as cyclopyrimorate (an inhibitor of homogentisate solanesyltransferase, recently commercialised by Mitsui Chemical Agro Inc) ^8^ and several novel cyclic methylphosphonates that target the pyruvate dehydrogenase complex ^5^. The traditional approach to discover herbicides is through mass chemical screening and only later determining the mode of action; as a result this process gives diminishing returns as highly active molecules are increasingly found to match known herbicides or a known mode of action.

The close relationship between the malaria parasite *Plasmodium falciparum* and plants like *Arabidopsis thaliana* ^9, 10^ means many antimalarial compounds are herbicidal and vice versa e.g. herbicides such as glyphosate, trifluralin, endothall and prodiamine are lethal to *Plasmodium* species ^11–14^ as antimalarials cycloguanil, pyrimethamine, sulfadoxine, dihydroartemisinin and artesunate are herbicidal ^15^. The chemical tools for antimalarial drug discovery have burgeoned in the last decade with chemical libraries composed of chemically novel antimalarials made publicly available by large, international consortia ^16, 17^. Exploiting this, two herbicidal compounds were discovered by screening a small subset of compounds from the Malaria Box chemical library against *A. thaliana* ^18, 19^. Overall, the Malaria Box is a 400-compound set of structurally diverse and chemically simple drug-like molecules that are toxic to blood stage *P. falciparum* and have physico-chemical properties suitable for orally delivered drugs ^17^. The screening of this subset against *A. thaliana* grown on Murashige-Skoog agar medium revealed twenty highly herbicidal compounds ^18, 19^. Twelve of these were tested against soil-grown *A. thaliana*, and only two remained highly herbicidal, despite all twelve molecules having physico-chemical properties within the broad range observed for commercial herbicides ^18, 19^. There are lessons to be learned from looking at compounds that succeed as herbicides on plate-based assays, but fail in soil-based assays. Instead of a bias towards successes, considering the physico-chemical properties of compounds that fail to translate to soil-based assays might provide more sophisticated rules for predicting which herbicidal hits will continue to work when sprayed on the leaves of soil-grown plants.

Previous analyses ^20–22^, including a recent study of 334 commercial herbicides ^23^, demonstrated that although the physico-chemical properties of herbicides were similar to orally delivered drugs, herbicides possess fewer proton donors, a lower partition coefficient and molecular weight ^23, 24^. To better understand the physico-chemical properties favoured by herbicides, we tested the entire Malaria Box for herbicidal activity of plate-based hits and determined which remained against soil-grown plants. Our analysis found correlations between some molecular properties and success in soil tests, which improved predictive ability. We used this newly refined set of rules to rank 631 liver-stage effective antimalarials *in silico*. Of the six highest-scoring compounds we found MMV1206386, a highly herbicidal tetrahydroquinoline derivative and chemically novel herbicidal compound, for which we obtained structure-activity information and determined that the compound potentially possesses a novel mode of action.

## Results

### Comparison of physico-chemical parameters of antimalarials and herbicides

To evaluate the overall potential *in planta* bioavailability of compounds from the Malaria Box library (MMV400) we compared their solubility in water (logS), partition (logP) and distribution (logD) coefficients, molar mass, proportion of aromatic atoms and polar surface area with the corresponding parameters of 360 commercial herbicides (**Supporting Dataset 1**). To visualise the distribution of values they were plotted as: molar mass – logS (**Figure 1A**), molar mass – logP (**Figure 1B**), proportion of aromatic atoms – logP (**Figure 1C**) and logD – polar surface area (**Figure 1D**). Overall, the physico-chemical properties of commercial herbicides and the MMV400 compounds were similar, however the range for each specific parameter was narrower for the MMV400 compounds. Compared to the herbicides, in general the MMV400 possessed lower water solubility and higher molar weight (**Figure 1A**): 80% of the MMV400 had values of logS between −3.6 and −5.8, and molecular masses between 270 and 470 Da, whereas for 80% of the herbicides logS values varied between −2.2 and −5.5 and 430 and 200 Da respectively. The overall lower solubility of the MMV400 correlated with higher logP values: only 65% of MMV400 compounds had a logP below 4.6, whereas the logP of 90% of commercial herbicides did not exceed this value (**Figure 1B**). Despite the higher logP of the MMV400, their proportion of aromatic atoms (20-50%) was below that of the herbicides (20-60%) (**Figure 1C**). The MMV400 were also less polar: 90% of the MMV400 had a polar surface area less than 80 Å, and logD between 1.5 and 6, in contrast for herbicides 90^th^ percentile of polar surface area was 130 Å, and logD fell within a range of −1.0 to 4.7 (**Figure 1D**). Thus, in comparison with herbicides the MMV400 are in general more hydrophobic, less polar and possess lower water solubility, however their physico-chemical properties are still within the limits characteristic for commercial herbicides.

**Figure 1.**
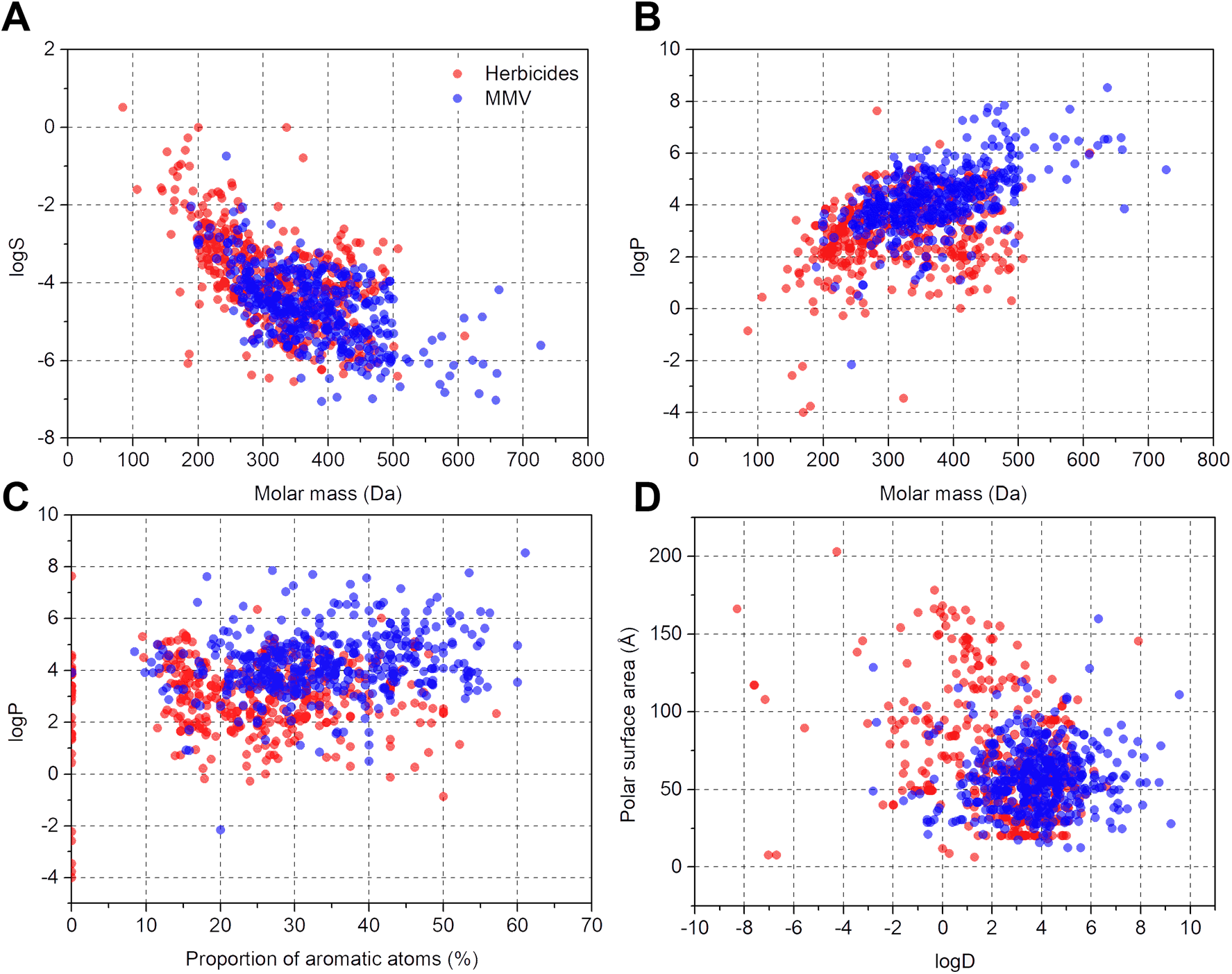
Cluster analysis of physico-chemical parameters of herbicides and the Malaria Box compounds. Red circles represent the 360 commercial herbicides and blue circles represent the MMV400 compounds. In general, the MMV400 have a narrower range for all physico-chemical properties assessed.

### Screening for herbicidal activity within the MMV400

MMV Plates B and C (160 compounds) were screened previously ^18, 19^ so to create a wider dataset of antimalarials to help correlate physico-chemical properties with herbicidal activity, we screened the remaining 240 compounds (plates A, D and E) of the MMV400 against *Arabidopsis thaliana* grown aseptically on an agar medium. Seeds were sown on solid agar medium containing 80 μM of the MMV400 compound (**Supporting Dataset 2**) or 80 μM control compound (typically a herbicide) left to germinate and grow for 16 days (**Supporting Figure 1**). In total, 52 of the MMV400 demonstrated herbicidal activity with phenotypes from stunted growth to bleaching and inhibition of germination.

To evaluate the herbicidal activity of the active antimalarials under more relevant conditions, we chose 39 compounds based on commercial availability and structural diversity. These antimalarials were tested against *A. thaliana* pre- and post-emergence on soil with a concentration range from 0 to 400 mg/L. Oryzalin and glyphosate were used as controls, being typical pre- and post-emergence herbicides respectively. Only 16 of the 39 compounds were active against plants grown on soil (**Figure 2**), including those two previously known ^18, 19^. The remaining 23 antimalarials did not inhibit the growth of the plants even at 400 mg/L (**Supporting Dataset 2**).

**Figure 2.**
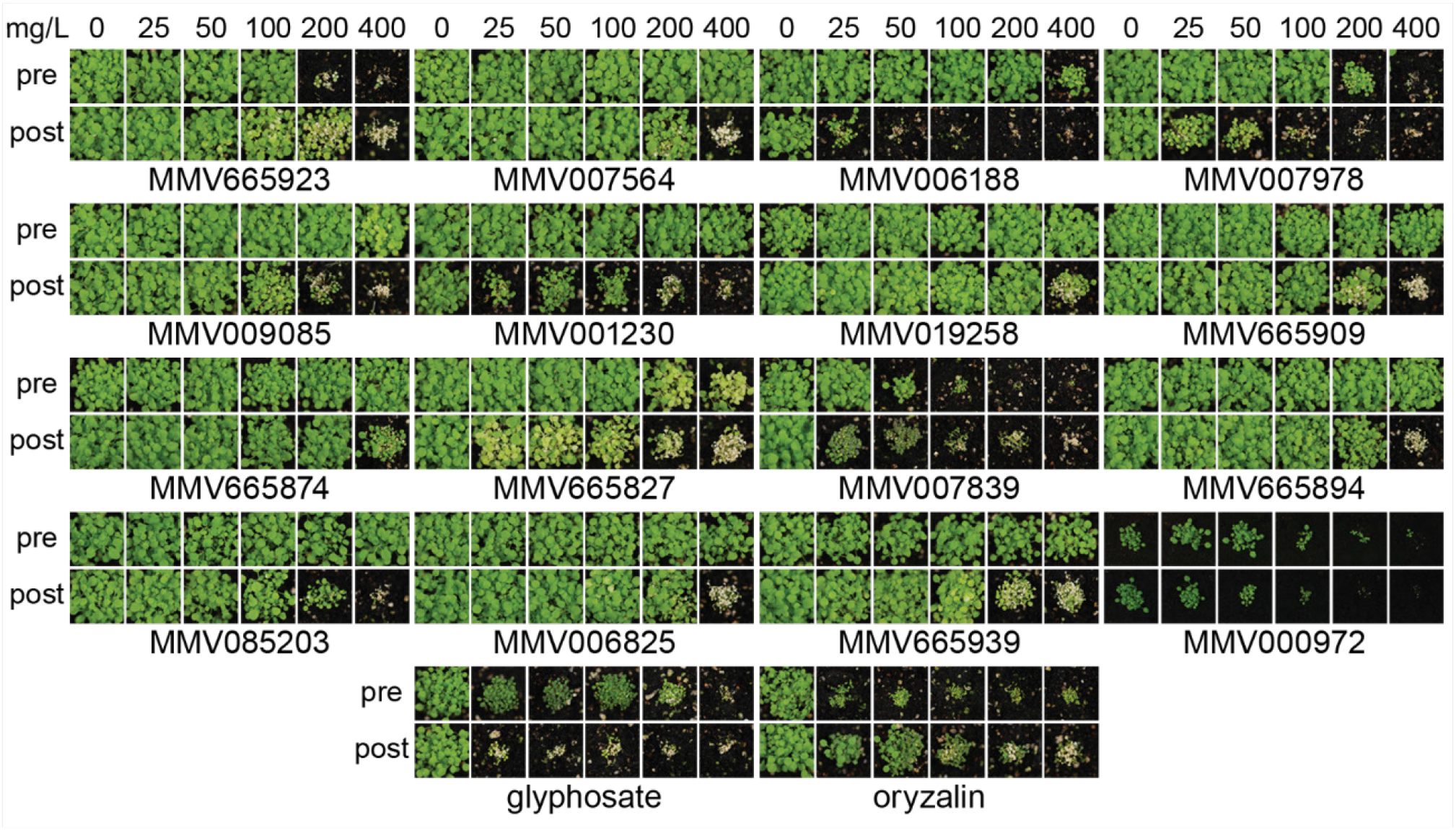
Herbicidal activity of selected MMV400 compounds against soil-grown *A. thaliana*. Each compound was applied on *A. thaliana* seeds on soil (pre) or on seedlings 3 and 6 days after germination (post). The antimalarials MMV007839 and MMV000972 were the most herbicidal. Commercial herbicides glyphosate and oryzalin were used as positive controls.

Overall, the subset of 16 herbicidal MMV400 was mainly effective post-emergence. Only two compounds (MMV000972 and MMV007839) were effective both pre- and post-emergence, with strong growth inhibition at an application rate of 50 mg/L (**Figure 2**). As shown previously ^18, 19^, two compounds MMV007978 and MMV006188 were effective post-emergence and weakly active pre-emergence. MMV001230 was only active post-emergence (**Figure 2**). These data show that, although the molecular properties of antimalarials are in general close to those of herbicides, just over half the antimalarials that were herbicidal on agar plate assays were inactive on soil. As a result, we thought it was beneficial to investigate how the physico-chemical properties of both the soil-active and inactive compounds correlated with activity.

### Correlating physico-chemical properties and herbicidal activity against soil-grown plants

To examine which properties might correlate with herbicidal activity for agar plates and soil, we examined the physico-chemical properties of the MMV400 compounds that were active on soil-grown plants versus compounds that were only herbicidal on agar. To better describe their molecular properties we used the set of descriptors that included logP, logD, logS, molecular mass, proportion of aromatic atoms, number of hydrogen bond donors and acceptors, number of rotatable bonds, polar surface area and formal charge. For each parameter we compared an average value for compounds active both on soil and agar plates (n = 16) and compounds which were not active on soil, but were active on agar plates (n = 23) and we used a two-sample t-test to determine statistical significance of any differences (**Table 1**).

**Table 1.**
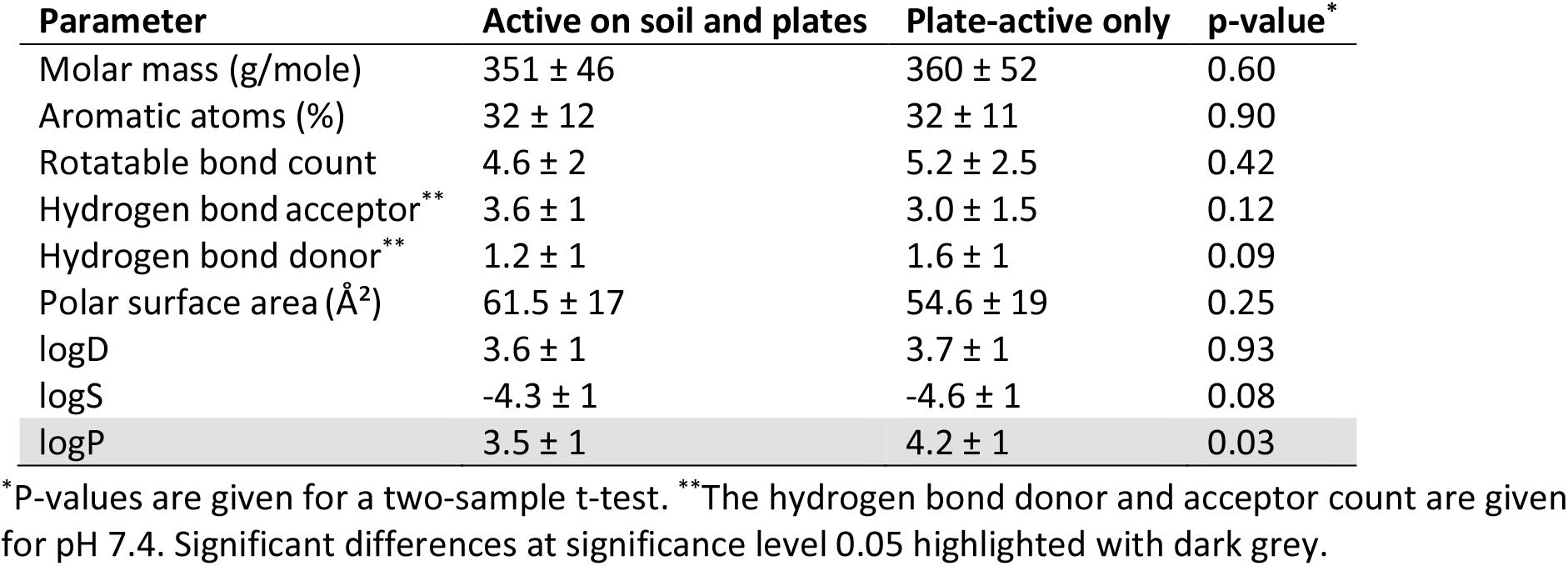
Average values of physico-chemical parameters for soil and plate-active antimalarials (± standard deviation) vs plant-only active antimalarials (± standard deviation).

The only significant (p <0.05) difference in the average values was for logP. The mean logP for soil-active compounds (3.5 ± 1) was lower than for compounds that were inactive on soil (4.2 ± 1) and closer to the mean logP for commercial herbicides (2.9 ± 1.5). This indicated that high logP reduced activity against soil-grown plants for antimalarials that were herbicidal on agar. Herbicide-like values for logS (p = 0.08) and number of hydrogen bond donors (p = 0.09) were important, but just showed a trend (p < 0.1).

To examine whether formal charge might correlate with activity against soil-grown plants, we calculated cumulative distribution functions of charge for soil-active antimalarials, commercial herbicides and antimalarials inactive on soil (**Figure 3A**). It is worth noting that only three of the 360 commercial herbicides (**Figure 3A**, dark grey bars) had a positive charge, with all having either neutral or slightly negative charge. By contrast, the soil inactive antimalarials had significantly higher proportion of positively charged compounds (Fisher’s exact test p < 0.05) (**Figure 3A**, dark grey bars). The soil-active antimalarials (**Figure 3A**, black bars) more closely resembled commercial herbicides with most being neutral or having a single negative charge, however one of the 16 compounds had a single positive charge.

**Figure 3.**
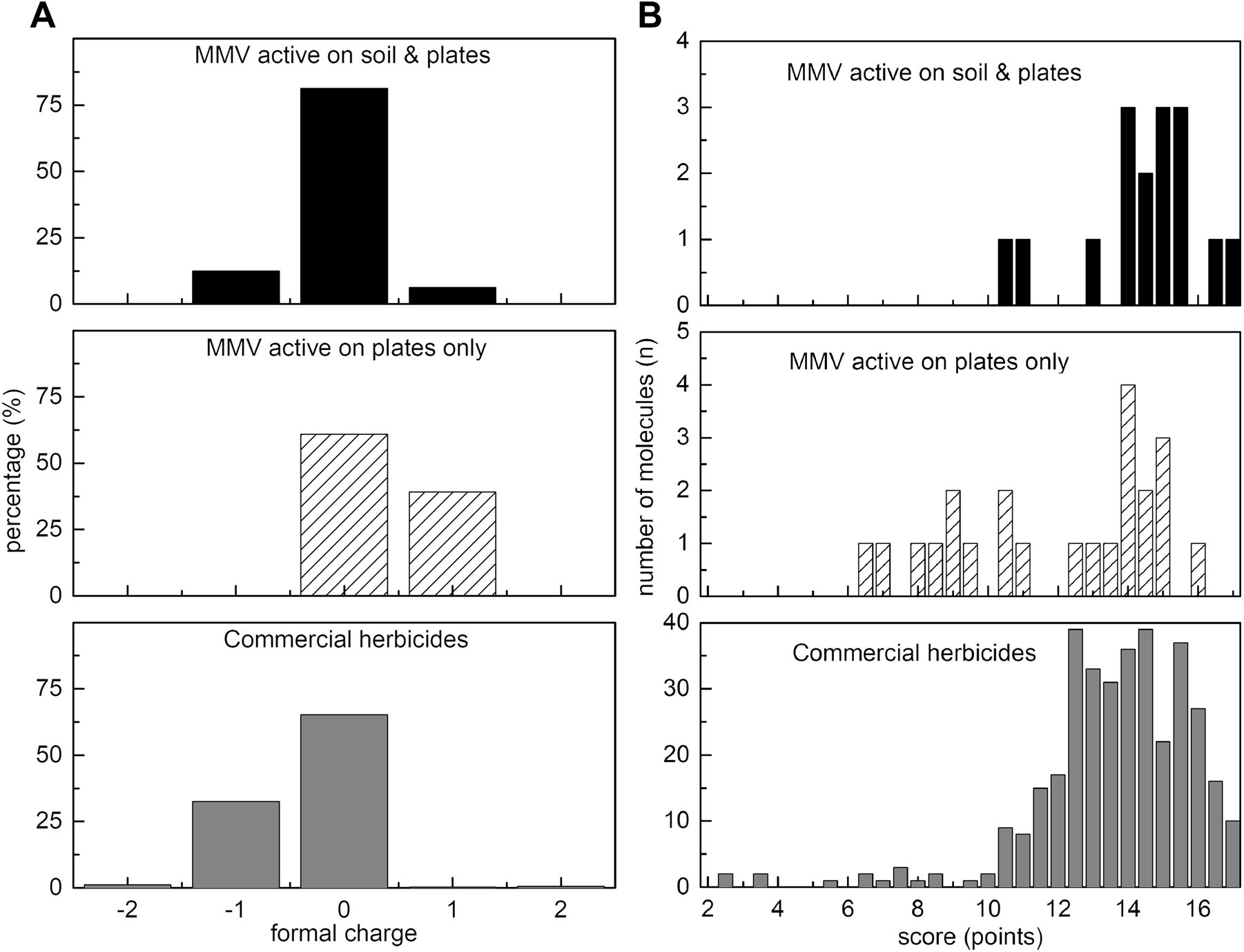
Charge and scores for herbicidal MMV400 compounds vs commercial herbicides. (A) Herbicidal MMV400 and commercial herbicides tend to be neutral or negatively charged. The formal charge was calculated at pH 7.4: positive formal charge means that a molecule exists in solution in cationic form, negative charge – anionic form, and zero charge corresponds to neutral form; (B) Distributions of scores for ‘herbicide-likeness’ for herbicidal MMV400 compounds and commercial herbicides. The MMV400 compounds that were only herbicidal on plates demonstrate a much wider distribution of scores, whereas the soil-active MMV400 closely matched the pattern of commercial herbicides.

To quantify how similar a particular compound was to the herbicides, a weighted scoring system was developed based on the closeness to the average of each herbicide physico-chemical parameter. Scoring rules were split into continuous and discrete physicochemical parameters. Physicochemical properties that are continuous variables include molar mass, proportion of aromatic atoms, polar surface area, logD, logS and logP. For these continuous parameters, we developed a scoring system based on the number of standard deviations away from the herbicidal mean that a particular parameter for an MMV compound fell (**Table 2**). The standard score for a parameter within one standard deviation was 1.5, two standard deviations was 1.0, three standard deviations was 0.5 and more than three standard deviations the score given was −1.0. Discrete physicochemical parameters (that is parameters possessing whole number values only) included rotatable bond count, number of H-bond acceptors/donors and formal charge. For these discrete parameters, we used scoring rules based on how far a parameter value is from the corresponding mode value for the 360 commercial herbicide set (**Table 3**).

**Table 2.** Weighted scoring for continuous parameters.

| Parameter | Average value ±SD* | 1 SD | 2 SD | 3 SD | > 3 SD |
| --- | --- | --- | --- | --- | --- |
| Molar mass (g/mole) | 317 ± 88 | 1.5 | 1.0 | 0.5 | -1.0 |
| Aromatic atoms (%) | 25 ± 12 | 1.5 | 1.0 | 0.5 | -1.0 |
| PSA ($\text{\AA}^2$) | 72 ± 39 | 1.5 | 1.0 | 0.5 | -1.0 |
| logD | 2.2 ± 2.4 | 1.5 | 1.0 | 0.5 | -1.0 |
| logS | -3.5 ± 1 | 1.5 | 1.0 | 0.5 | -1.0 |
| logP | 2.9 ± 1.5 | 3.0 | 2.0 | 1.0 | -1.0 |

\*The score given is based on being within 1, 2, 3 or >3 standard deviations (SD) from the average value (for 360 commercial herbicides). Note that the logP score is weighted differently.

**Table 3.**
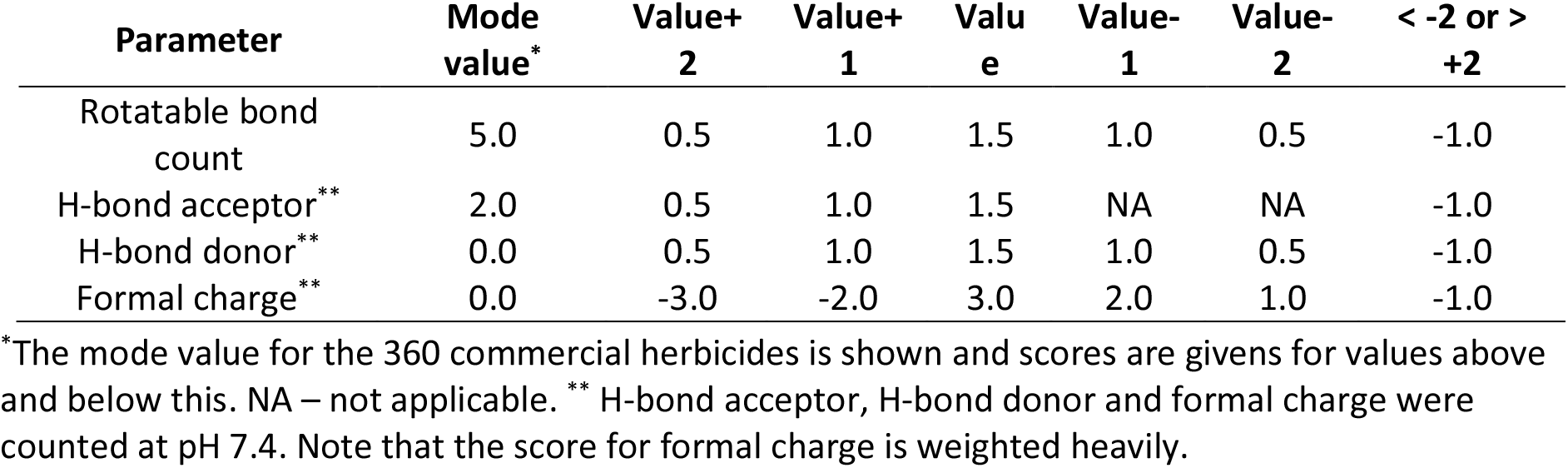
Weighted scoring for discrete parameters.

To weight the scoring system to account for the aforementioned importance of logP and formal charge (**Table 1**, **Figure 3A**) we doubled the standard score for logP values falling within one to three standard deviations, but no additional penalty if logP fell outside three standard deviations (**Table 2**). For the discrete parameter of formal charge, positive values were heavily penalised and neutral or negative values had a higher weighting (**Table 3**). The total score for herbicide-likeness was the sum of scores for all parameters with the maximum possible score being 18 points (**Table 2** and **Table 3**).

Using this scoring system, we plotted distribution curves for antimalarials active only on plates (**Figure 3B**, white, diagonally hashed), the antimalarials active on plates and soil-grown plants (**Figure 3B**, black) and commercial herbicides (**Figure 3B**, dark grey). For the 39 MMV-antimalarials that were tested against soil-grown *A. thaliana* (**Supporting Dataset 2**) the majority of soil-active antimalarials had scores of 14 or above, with the average score of 14.4 points (n = 16), that was close to 13.6 points for commercial herbicides (n = 360). Scores for plate active-only antimalarials for herbicide likeness varied from 6.5 to 17, with the average value 11.9 points (n = 23) which is significantly different (p < 0.05) from the average value for herbicides and soil-active antimalarials. This scoring system appears to give values consistent for compounds showing herbicidal activity in more natural, soil-grown conditions.

To test this weighted scoring system on a different dataset, we performed an *in silico* pre-screening of 631 molecules that were active against liver-stage malaria parasites, which possess low hepatotoxicity and potentially good oral bioavailability ^25^. Consisting of antimalarial compounds a high proportion should be herbicidal, but this liver-active set were structurally different to the blood stage MMV400 antimalarials. We hoped to demonstrate whether the aforementioned rules could focus attention only on compounds with appropriate physico-chemical properties of soil-active herbicidal compounds. We scored all 631 (whose names also have an MMV prefix) for herbicide-likeness and tested the top six scoring compounds that were commercially available.

Four of these six top-scoring liver-stage active antimalarial compounds were herbicidal, but only MMV1206386 had activity commensurate with commercial herbicides. MMV1206386 completely inhibited the growth of *A. thaliana* at 200 mg/L application rate in both pre- and post-emergence application (**Figure 4A**). Two compounds: MMV1085491 and MMV1266305 were completely inactive, despite a high score for herbicide-likeness (**Figure 4A**), probably due to the absence of a plant target. Notably, both compounds share a similar structural core (**Supporting Dataset 3**).

**Figure 4.**
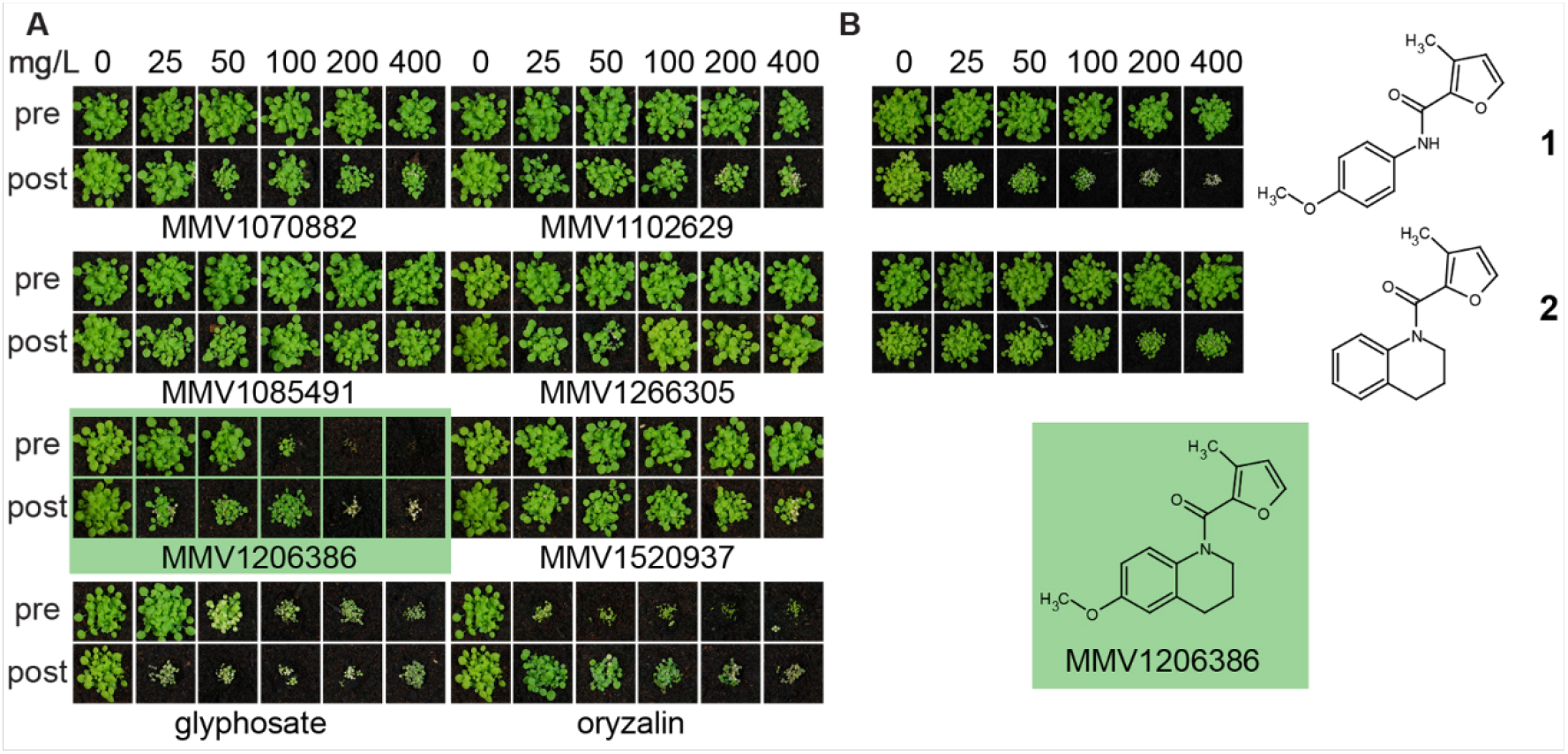
Herbicidal activity of liver-stage active MMV compounds on soil. (A) Herbicidal activity of liver-stage active MMV compounds against soil-grown *A. thaliana*. Comparison of herbicidal activity of selected antimalarials against *A. thaliana* treated pre- and post-emergently show that MMV1260386 (highlighted in green) was the most herbicidal and comparable with commercial herbicides; (B) Analogs of MMV1260386 showed herbicidal activity did not depend on quinolone motif. Compound **1** which lacks the second ring was decreased in pre-, but nor post-emergence activity. Elimination of p-methoxy group (**2**), virtually eliminated herbicidal activity.

To assess the opportunities for improving herbicidal activity of MMV1206386 we screened 25 structural analogues and found that none of structural changes increased herbicidal activity; some had no effect, while many of them decreased or abolished herbicidal activity (**Figure 4B**, **Figure 5**).

**Figure 5.**
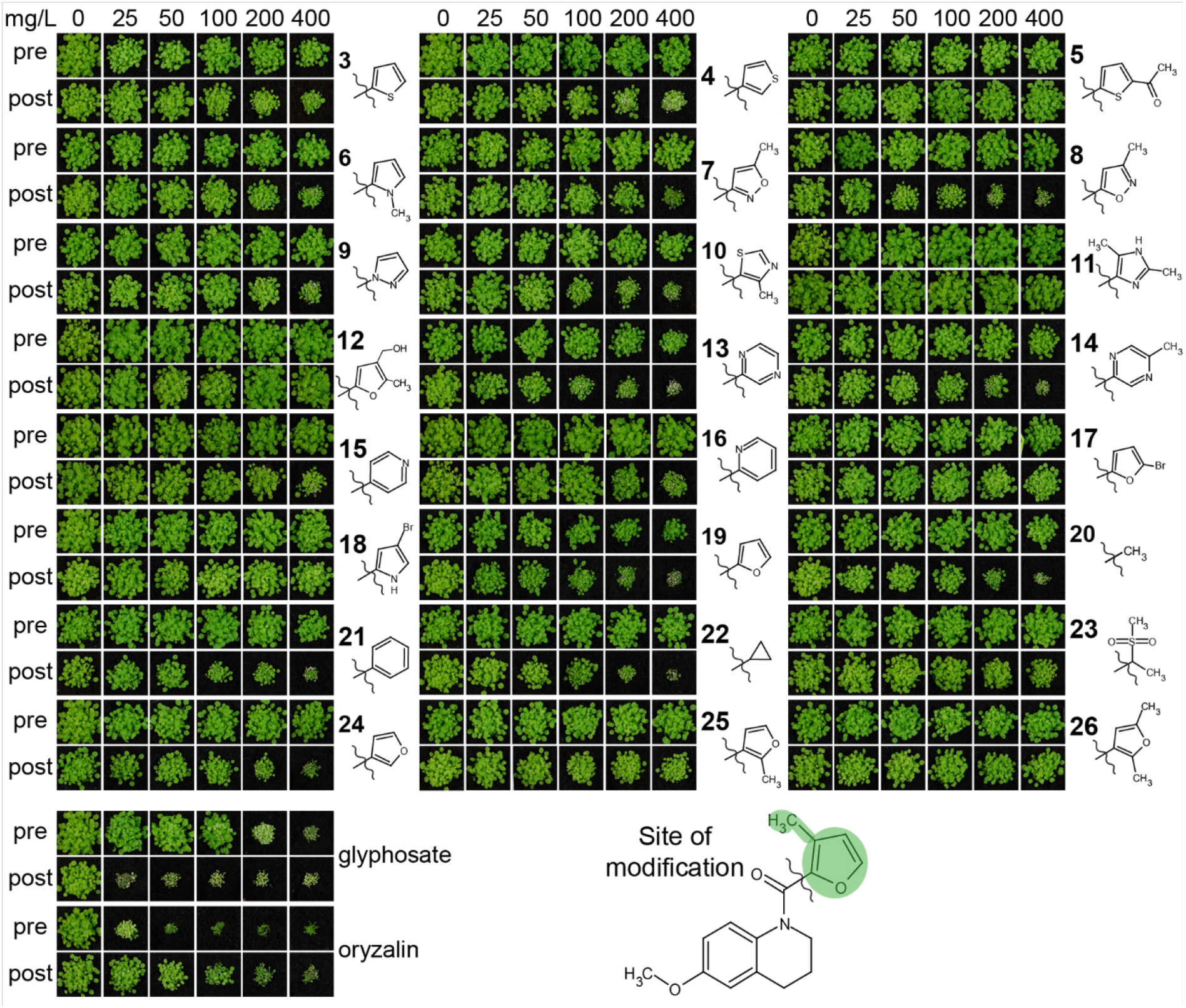
Effect of the 2-methyl furanyl group in MMV1260386 on herbicidal activity. Pre-emergence activity was eliminated in all molecules except **19**. Compounds **5, 11, 12, 17, 18, 23, 25,** and **26** completely lost herbicidal activity.

First of all, we checked if the tetrahydroquinoline motif was essential for herbicidal activity. We found that replacement of the motif with the appropriate methoxyphenyl group as in **1**, did not change post-emergence herbicidal activity, but eliminated pre-emergence. In contrast, the methoxy group in the position (6) of quinolone ring was crucial for herbicidal activity: removal of this group in **2** completely eliminated pre-emergence activity and significantly reduced post-emergence activity.

To assess the importance of the furanyl motif (marked, **Figure 5**) we replaced it with other heterocycles as in **3**-**19**, **24**-**26,** aromatic **21** and aliphatic groups **20, 22** and **23**. In most cases the replacement of this group greatly reduced the activity both pre- and post-emergence. Replacing the furan motif with other five-membered heterocycles such as those in **3**-**10** eliminated pre-emergence activity and drastically decreased post-emergence activity. Replacement of the furanyl motif with a pyrimidinyl motif as in **13** and **14** also decreased herbicidal activity, but to a smaller extent than for replacement with a pyridinyl moiety, as in **15** and **16**.

Increasing the heterocyclic nature of the five-membered ring as in **11** and **12** or increasing the functionality of the five membered ring as in **5**, **17**, **18**, **26** also lost activity. Removal of the 3-methyl group as in **19** clearly reduced pre- and post-emergence herbicidal activity. Replacement of furan motif with groups which did not contain heterocyclic core **20**-**22** similarly decreased herbicidal activity or virtually eliminated as in **23**. Changing the position of the link between the furan motif and the carbonyl group greatly reduced herbicidal activity in **24**, and completely eliminated it in **25**. With these results available, we examined the potential of MMV1206386 to act as a herbicide against more relevant weed species. Eight crop specific weed species in pre- and post-emergence application were assayed (**Table 4**).

**Table 4.**
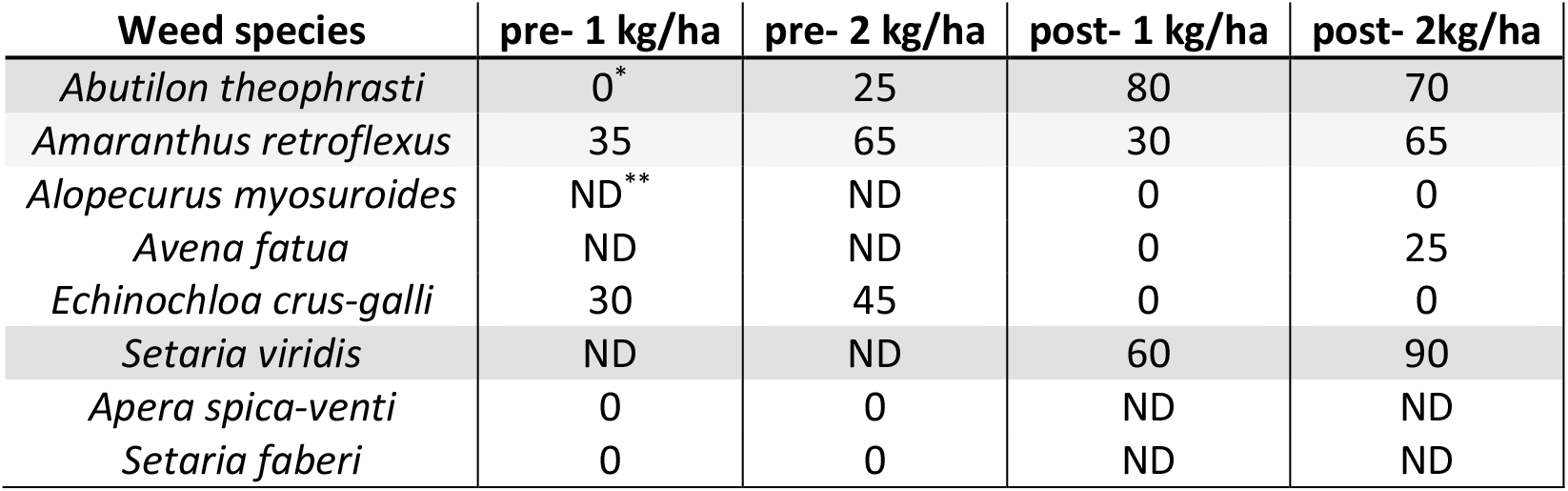
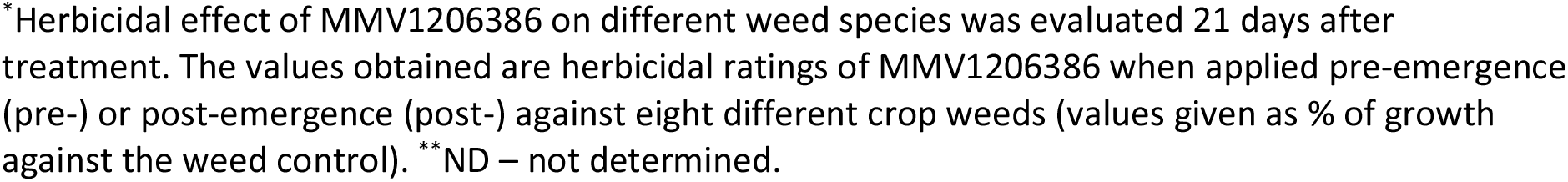
MMV1206386 is strongly herbicidal against *Abutilon theophrasti* and *Setaria viridis*.

| Weed species | pre- 1 kg/ha | pre- 2 kg/ha | post- 1 kg/ha | post- 2kg/ha |
| --- | --- | --- | --- | --- |
| *Abutilon theophrasti* | 0* | 25 | 80 | 70 |
| *Amaranthus retroflexus* | 35 | 65 | 30 | 65 |
| *Alopecurus myosuroides* | ND** | ND | 0 | 0 |
| *Avena fatua* | ND | ND | 0 | 25 |
| *Echinochloa crus-galli* | 30 | 45 | 0 | 0 |
| *Setaria viridis* | ND | ND | 60 | 90 |
| *Apera spica-venti* | 0 | 0 | ND | ND |
| *Setaria faberi* | 0 | 0 | ND | ND |

\*Herbicidal effect of MMV1206386 on different weed species was evaluated 21 days after treatment. The values obtained are herbicidal ratings of MMV1206386 when applied pre-emergence (pre-) or post-emergence (post-) against eight different crop weeds (values given as % of growth against the weed control). \*\*ND – not determined.

We found that MMV1206386 was strongly active as a post-emergence herbicide against *Abutilon theophrasti* (velvetleaf) and *Setaria viridis* (green bristlegrass). Moderate susceptibility was observed in the case of *A. retroflexus* whereas *E. crus-galli* was moderately sensitive to pre-emergence treatment (45% of control at 2,000 g/ha), but tolerant to post-emergence treatment. Three species, *A. fatua, A. spica-venti* and *S. faberi* were resistant to MMV1206386. Based on these data we can assume that MMV1206386 has the potential to control a wide range of species, and should be more successful if used as a post-emergence herbicide. Since a search of MMV1206386 structural analogues among known commercial herbicides did not find any structurally similar molecules, we thought MMV1206386 might have a novel mode of action. To determine this we obtained an industry-standard physiological profile, which includes 13 different assays and generates a profile that can be compared to commercial herbicides, for many their modes of action being known ^19, 26, 27^. Firstly, we tested MMV1206386 against small plants and cell suspension to see which symptoms are induced by treatment with the compound (**Figure 6A**).

**Figure 6.**
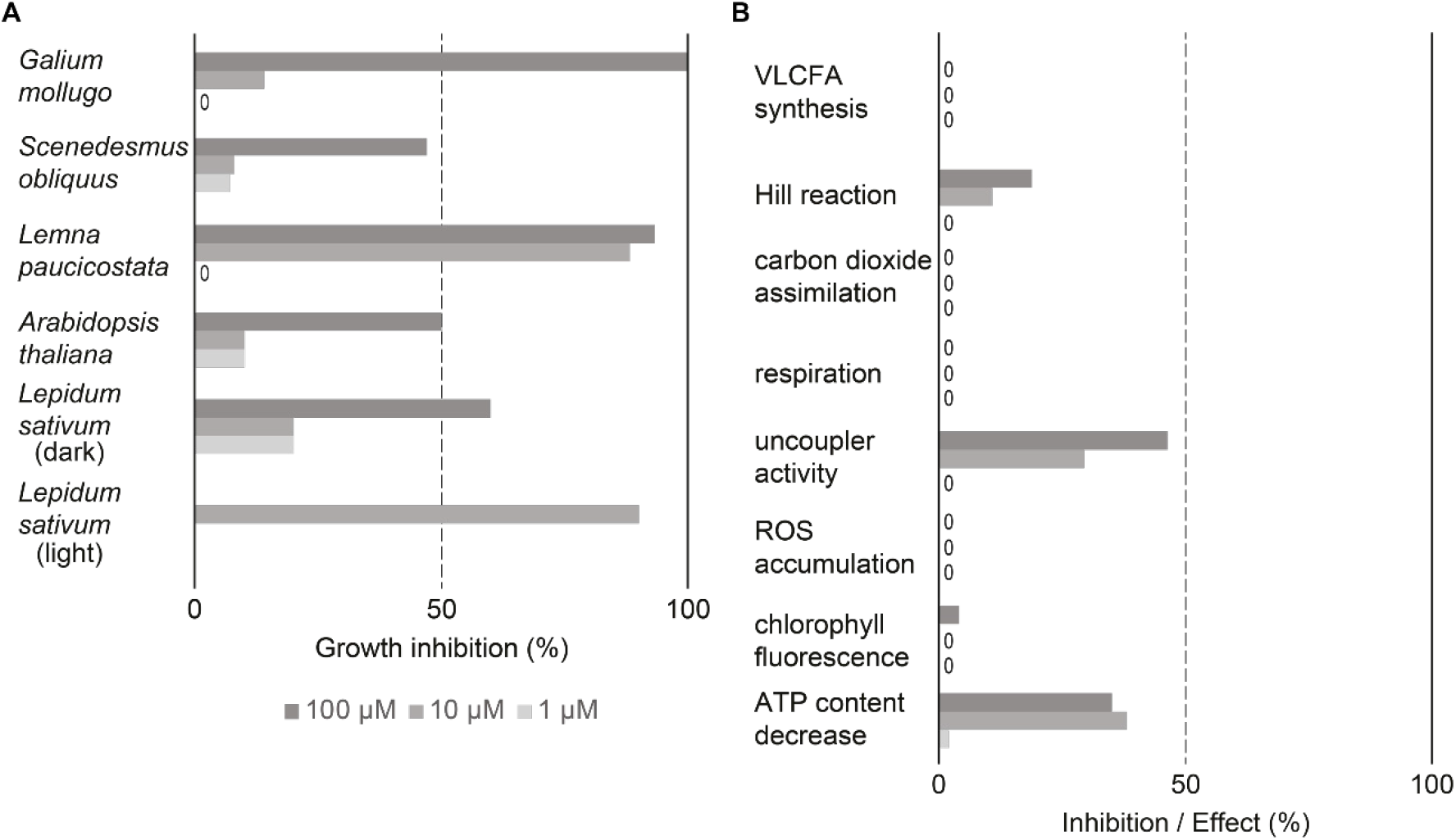
Evaluation of MMV1260386 by a BASF physiological profile^19^. (A) MMV1260386 strongly inhibited growth of both whole plants and cell suspensions. The herbicidal effect of MMV1206386 was noticeably high under light growth conditions. “0” – no inhibitory effect; (B) MMV1260386 had moderate uncoupler and Hill reaction inhibitor activities. VLCFA – very long chain fatty acid; ROS - reactive oxygen species.

MMV1206386 strongly inhibited cell division in heterotrophic *Galium mollugo* (cleaver) cell suspension culture at 100 μM. At lower concentrations cell division was only mildly affected. Growth of *Lemna paucicostata* (duckweed) was strongly inhibited when the compound was applied at 10 μM and 100 μM accompanied by rapid necrosis, reduced leaf growth and root growth inhibition. Treatment of plate-grown *A. thaliana* by MMV1206386 caused chlorosis, reduced leaf growth, root growth inhibition and hypocotyl swelling in seedlings, and these symptoms were similar to what was observed for soil-grown plants treated with MMV1206386. The germination inhibition assay with dark-grown *Lepidum sativum* (cress) showed that MMV1206386 had a moderate effect on germination rate. In contrast, when *L. sativum* plants were grown under light the inhibition of germination was more than 70% at all tested concentrations.

Compared to industry-standard physiological profiles of inhibitors of photosystem II, MMV1206386 mildly inhibited photosynthetic electron transport in photosystem II in isolated wheat thylakoids at 100 μM (**Figure 6B**). In the same linecontrast, chlorophyll fluorescence measured in *A. thaliana* was only slightly affected at 100 μM. MMV1206386 demonstrated robust or medium uncoupler activity: the electron transport activity dropped by 46% and 30% at 100 μM and 10 μM respectively upon treatment, as well as ATP levels decreased robustly. Interestingly, there was no effect on the cress very long chain fatty acid synthesis (at 1 μM), carbon dioxide assimilation (at 1000 μM), respiration (at 100 μM) or reactive oxygen species accumulation (at 10 μM and 100 μM). To find out what mode of action MMV1206386 had, this physiological profile was compared with the BASF database of physiological profiles representing all known modes of action for herbicides and reference compounds with standard modes of action. This comparison did not reveal any matches among fingerprint profiles for commercial herbicides, therefore suggested that MMV1206386 represents a novel mode of action.

## Discussion

Since the advent of Lipinski’s rules for oral drugs in 1997 several studies have analysed the physico-chemical parameters of herbicides ^20–23, 28^. Where the underpinning datasets were made known, these consisted of commercial herbicides, so none of these studies considered initial herbicidal hits that then failed in foliar applications ^20, 22, 23^. Only the study by Clarke considered data from both successful and unsuccessful herbicidal leads ^21^. Unfortunately, as the identities of their training set was suppressed, no subsequent analysis could use these data.

To create a dataset containing positive and negative data we completed screening of the 400-compound Malaria Box against *A. thaliana* on agar plates and then tested a sub-set against soil-grown plants. The reason to choose the Malaria Box was the fact that it was partially screened in previous studies and yielded many herbicidal hits, so screening the rest of the library would expand the training set ^18, 19^. Screening the entire MMV400 increased the number of actives tested on soil and agar from 9 to 39 compounds plants, enabling statistical analyses like Fisher’s exact test and two sample t-test.

To describe molecular properties of herbicides, previous studies used Lipinski’s ‘rule of five’ itself or with minor corrections ^20–22^. The original ‘rule of five’ employed a few simple descriptors such as partition coefficient (logP), molecular weight, number of hydrogen bond donors and acceptors and rotatable bond count to predict oral bioavailability of a drug candidate ^24^. These parameters do not consider ionisation properties, polarity or solubility, so to better characterise the physico-chemical parameters of herbicidal and non-herbicidal molecules we used an extended set of descriptors ^23^. The set included all Lipinski’s descriptors as well as: distribution coefficient (logD); characterises distribution of the dominant ionisation form at given pH, solubility coefficient (logS); characterises solubility of molecules, polar surface area, aromatic atom proportion and formal charge. Not surprisingly, compounds active against soil-grown plants as well as compounds active only against plate-grown plants fit within characteristic limits of physico-chemical properties for commercial herbicides. The logP of antimalarials active against soil-grown plants had a low average value of logP that did not differ significantly from commercial herbicides, whereas the antimalarials active only against agar-grown plants had significantly higher logP value.

Another parameter that differed was formal charge. Soil-active antimalarials were mostly neutral or negatively charged. Although positive charge facilitates absorption by roots or leaves ^29, 30^, it might impede translocation of herbicide molecules. This is similar to what has been observed for nanoparticles ^31^ and fertilizers ^32^, where in contrast negatively charged and neutral particles have increased translocation ^33^. In addition, organic soils are in general possess negatively charged materials and so positively charged herbicides are readily bound, limiting pre-emergent efficacy ^34^.

Our hypothesis was a weighted scoring system for herbicide-likeness could allow *in silico* screening of libraries and find new herbicides. The only published scoring system for herbicide-likeness was one proposed by Avram *et al.* in 2014 ^35^ where quantitative estimates of drug-likeness based on the use of distribution functions were used to create a continuous ranking model. Although it worked well describing herbicide-likeness of molecules deposited in the AgroSAR patent database, it did not consider formal charge and did not discriminate between the different descriptors, being used, according to their importance. Moreover, using a continuous distribution fraction for describing discrete parameters such as hydrogen donor and acceptor count, although convenient, is not appropriate for discrete parameters.

The scoring system described here takes into account the importance of logP and formal charge for herbicide-likeness and scores discrete and continuous parameters differently. After the scoring system for herbicide-likeness was validated using the subset of Malaria Box compounds that were tested against soil-grown plants we applied it to a compound library of 631 antimalarials active against liver-stage *P. falciparum* ^25^. We chose this library as it consists of compounds with high antiprotozoal activity, so we expected a high proportion to be herbicidal, but the set would be structurally different from the blood stage antimalarials in the Malaria Box.

For the library targeted against liver-stage *P. falciparum*, eleven molecules scored 17 or more of a maximal possible 18 points, however only six were readily available. The only molecule to exhibit high herbicidal activity was MMV1206386, and a search did not reveal any herbicides with similar chemical structure. Structure-activity studies revealed critical aspects of its structure, and physiological profiling suggested that MMV1206386 has a new mode of action with an unusual combination of affected processes. While MMV1206386 demonstrated uncoupler activity, the respiration and carbon dioxide assimilation processes were not affected, although uncoupling is normally reflected in these processes. The combination of symptoms induced by MMV1206386 treatment in *A. thaliana* did not match with any of more than 2,700 tested by BASF before. There were three matches (of more than 10,000 compounds) for the symptoms observed in *Lemna paucicostata* treated by MMV1206386, however the physiological profiles did not match those obtained for MMV1206386. The combination of structural uniqueness, distinct physiological profiles and unique combination of symptoms support the hypothesis that MMV1206386 has a new mode of action.

## Conclusion

By comparing failed and successful hits from the Malaria Box it was revealed how different molecular properties had different impacts on the chance of herbicidal activity. Having a suitable partition coefficient and formal charge appeared essential, whereas other parameters could range more widely. Based on these findings we developed a weighted scoring system for herbicide-likeness and used it to select top-scoring compounds from a large compound library of liver-stage effective antimalarial leads. One molecule (MMV1206386) had efficiency against soil-grown plants comparable to commercial herbicides and no close structural analogues in use and its physiological profile indicate a new mode of action. Thus, the weighted rules appear to be a useful tool when using high throughput screening approaches in the discovery of new herbicides.

## Methods

### Safety comment

No unexpected or unusually high safety hazards were encountered.

### Updating the Gandy et al. (2015) herbicide database

A database of commercial herbicides compiled in 2015 contained 334 compounds ^23^ and as a prelude to this study, we updated this database. Dimethylarsinic acid, DSMA (disodium methyl arsenate) and MSMA (monosodium methyl arsenate) were removed as they are rarely or no longer used due to toxicity ^36^. Acrolein, which is used as a commercial algicide, was removed because the database is focused on herbicides applied against soil-grown weeds. Metolachlor, a racemic mixture of S-metolachlor and R-metolachlor, was excluded because it duplicated physico-chemical properties of S-metolachlor that are already present in the database. To check if we missed any herbicides in the first iteration we screened through “Alan Wood’s Pesticides Common Names Compendium” and “Pesticide Properties Data Base by the University of Hertfordshire” and found 27 commercial herbicides not in the original database ^36, 37^. These include four 4-hydroxyphenylpyruvate dioxygenase inhibitors (fenquinotrione, tioclorim, tolpyralate, tefuryltrione); five that affect protoporphyrinogen oxidase (flumipropyn, fluorodifen, tiafenacil, bencarbazone, trifludimoxazin); three photosystem II inhibitors (ametridione, dipropetryn, eglinazine); two synthetic auxins (4-chlorophenoxyacetic acid, halauxifen); and six herbicides from different groups, including the acetyl-CoA carboxylase inhibitor haloxyfop-P, the acetolactate synthase inhibitor monosulfuron, 1-deoxy-D-xylulose 5-phosphate synthase inhibitor bixlozone, a phytoene desaturase inhibitor metflurazon, an auxin transport inhibitor thidiazuron, and an inhibitor of long chain fatty acid synthesis terbuchlor. Seven of the newly added herbicides have unknown modes of action (benzipram, florpyrauxifen, haloxydine, isopolinate, pyribambenz-isopropyl, pyribambenz-propyl, trifopsime). In addition, a few herbicides have been introduced to market since 2015 and so were also added. These three herbicides each possess a novel mode of action: an inhibitor of pyruvate dehydrogenase (clacyfos), an inhibitor of homogentisate solanesyltransferase (cyclopyrimorate) and an inhibitor of dihydroorotate dehydrogenase (tetflupyrolimet). The SMILES codes of the newly added compounds were used to propagate their details and the new database (**Supporting Dataset 1**).

### Calculation of physico-chemical properties of the Malaria Box compounds

To describe physico-chemical properties of the Malaria Box compounds a set of physico-chemical descriptors was generated. These included solubility coefficient (logS), partition coefficient (logP), distribution (logD) coefficient, molar mass, proportion of aromatic atoms, polar surface area, rotatable bond count, hydrogen bond donor count, hydrogen bond acceptor count and formal charge. The parameter values were calculated using calculator plugins included in Marvin Suite program package v. 20.19 (ChemAxon). Formal charge, hydrogen bond donor count, hydrogen bond acceptor count, were calculated for the major tautomeric form at pH 7.4.

### Herbicidal activity assay on plates

To assess herbicidal activity of the Malaria Box compounds they were tested against agar-grown *Arabidopsis thaliana*. A 2.5 μL aliquot of either an 8 mM solution of antimalarial in DMSO or pure DMSO (negative control) was placed into a well of a sterile transparent 96-well plate prior to pouring of 250 μL of molten MS-agar medium that contained 1% of agar, 4 g/L of Murashige-Skoog, 10 g/L of glucose and 0.3% 2-(N-morpholino) ethanesulfonic acid (v/v), pH 5.7. After the agar solidified roughly 30-40 ethanol-sterilised *A. thaliana* (Col-0) seeds were sown onto the surface as 25 μL of seed suspension in 0.1% agar. Prior to the experiment the seeds were stratified for three days to synchronise germination. After the surface of agar dried the plates were covered with lids and sealed with porous tape, transferred to a growth chamber and left to grow under long-day illumination (16 h light/8 h dark, 136 μmol/m^2^/sec) at 26°C and 60% relative humidity. After sixteen days, lids were removed and images taken.

### Herbicidal activity assay on soil

To assess herbicidal activity ~30 *A. thaliana* Col-0 seeds were sown in 63 × 63 × 59 mm pots consisting of Irish peat pre-wet before sowing. Seeds were treated for 3 days in the dark at 4°C to synchronise germination and then grown in a chamber at 22°C, with 60% relative humidity and in a 16 h light / 8 h dark photoperiod. Antimalarial compounds and control herbicide oryzalin were initially dissolved in DMSO at 20 mg/mL and further diluted in water containing 0.02% surfactant (Brushwet, SST Australia, v/v) prior to treatments. Another herbicide control, glyphosate was diluted in the same manner, but the original 20 mg/mL stock was prepared in water. DMSO at a concentration of 2% (v/v) was used as a negative control. Seeds or seedlings were treated as previously detailed ^19^ with 500 μL of 0, 25, 50, 100, 200 or 400 mg/L solutions of each compound.

Pre-emergence treatments were done as trays were moved into their first long day, whereas post-emergence treatments were conducted three and six days after germination. Seedlings were grown for 16 days after treatment before photos were taken.

### Physiological profiling of MMV1206386

To determine the mode of action of MMV1206386 the effect of the compound on different physiological processes was studied according to protocols previously described ^26, 27^. The influence of MMV1206386 on cell growth was studied in cell suspensions of phototrophic green algae *Scenedesmus obliquus*. The effect of MMV1206386 on Hill reaction rate was assessed in isolated wheat (*Triticum aestivum*) chloroplasts. Carbon assimilation and oxygen consumption was studied in heterotrophic *Galium mollugo* cell suspensions. Oxidative phosphorylation uncoupler activity and accumulation of reactive oxygen species were measured in *Lemna paucicostata* root tissue. Toluidine-blue staining of *Lepidium sativum* hypocotyls was used for the detection of inhibition of very long chain fatty acid synthesis. Additionally the effect of MMV1206386 on *A. thaliana* seedling morphology, chlorophyll fluorescence and ATP content was evaluated.

## Acknowledgement

The authors thank Simone Huber, Sarina Ruehm and Stefanie Zimmermann at BASF for performing the physiological profile. The authors thank Philippe Hervé and Bruce Lee from Nexgen Plants for fruitful discussions. K.V.S. is supported by Research Training Program Fee Offset and International Student Research Training Program Stipend. J.S.M. was supported in part by an Australian Research Council (ARC) Future Fellowship (FT120100013) and J.H. by an ARC Discover Early Career Researcher Award DE180101445. This work was supported in part by ARC grant DP190101048 to J.S.M., K.A.S. and J.H.

## Supporting Information

**Supporting Figure 1**: Contains collated images for the agar plate germination assay of Malaria Box compounds.

**Supporting_Dataset_1.zip**: Contains the database of physico-chemical properties and chemical structures of 360 commercial herbicides.

**Supporting_Dataset_2.zip**: Contains the database of physico-chemical properties, chemical structures and scores for herbicide likeness of 400 antimalarials from the Malaria Box.

**Supporting_Dataset_3.zip**: Contains the database of physico-chemical properties, chemical structures and scores for herbicide likeness of 631 liver-stage antimalarials.

---

**Supporting Figure 1.**
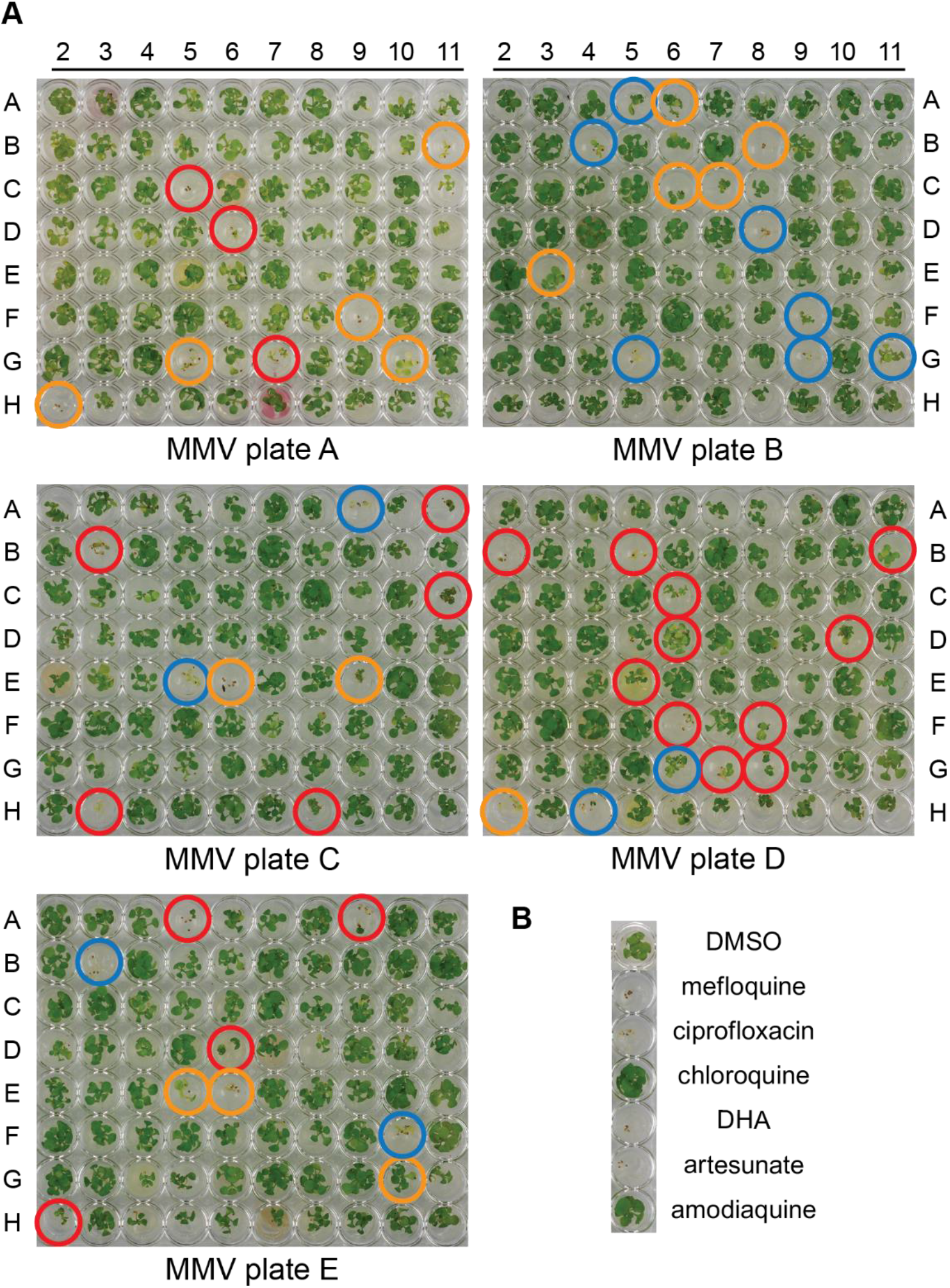
Agar plate herbicidal activity of antimalarial compounds. (A) Activity of the MMV400 against *A. thaliana* at 80 μM. Orange circles indicate the compounds active against soil-grown plants, red circles indicate plate-only active compounds, blue circles denote plate-active compounds that were not tested for activity against soil-grown plants; (B) Positive controls at 80 μM, DHA – dihydroartemisinin, DMSO was used as a negative control at 1% (vol) concentration.

