## Supplementary figures and images for "Refining physico-chemical rules for herbicides using an antimalarial library"

### 314-40-9.png

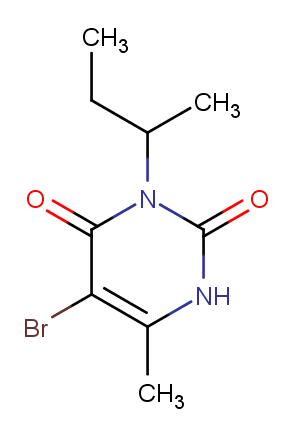

### 330-54-1.png

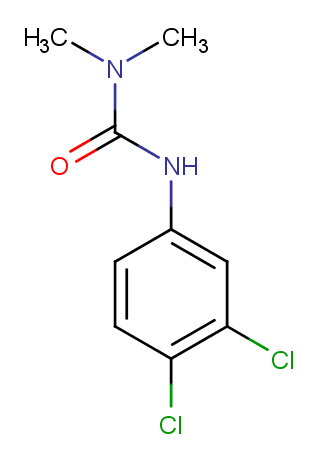

### 330-55-2.png

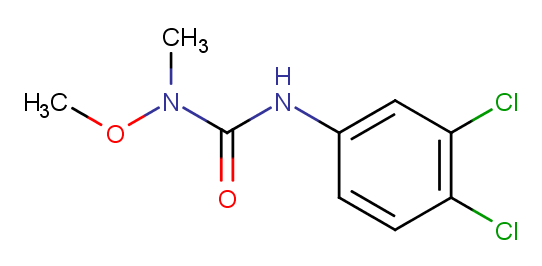

### 1582-09-8.png

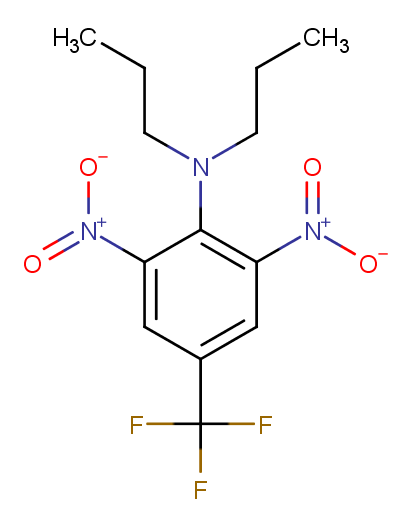

### 1610-18-0.png

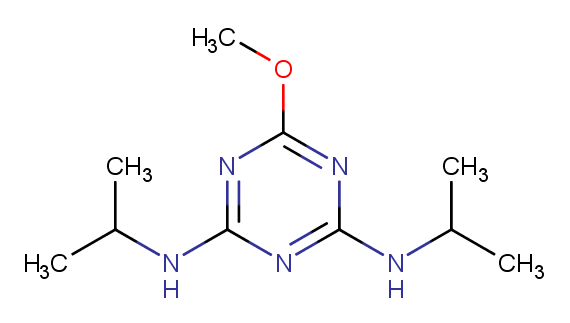

### 1689-83-4.png

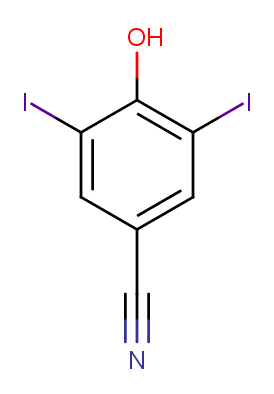

### 1689-84-5.png

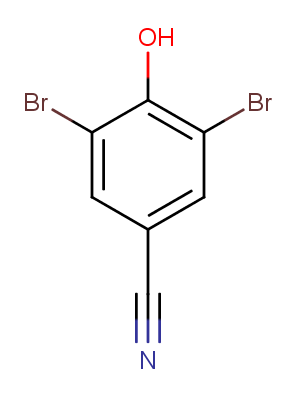

### 1698-60-8.png

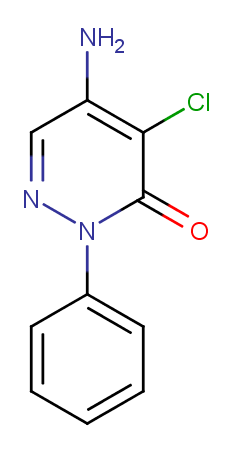

### 1702-17-6.png

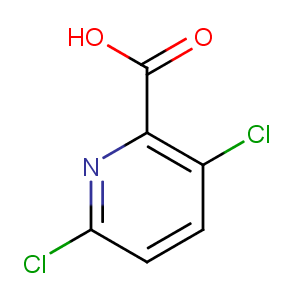

### 1746-81-2.png

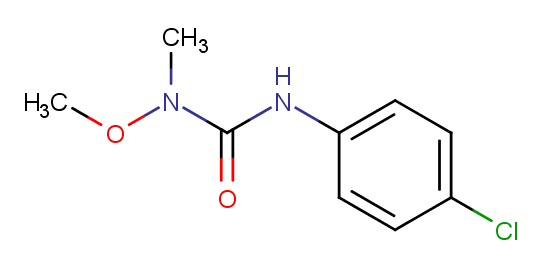

### 1836-75-5.png

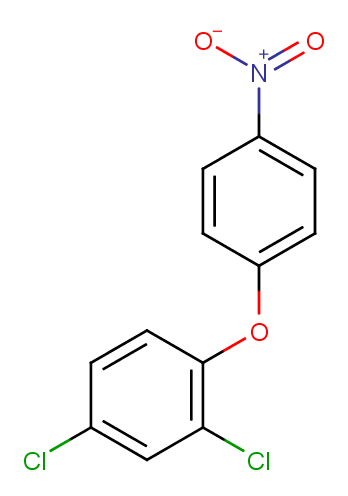

### 1861-32-1.png

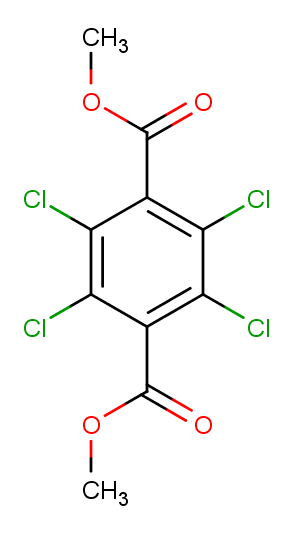

### 1861-40-1.png

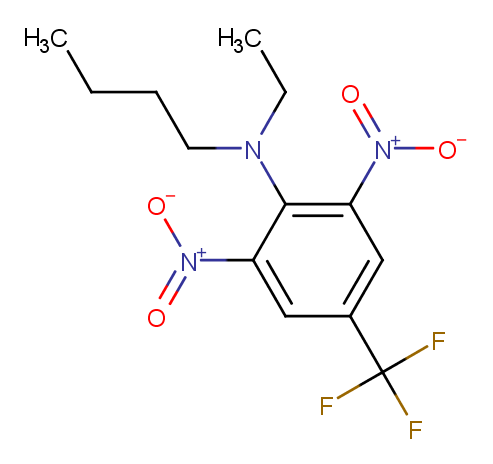

### 1910-42-5.png

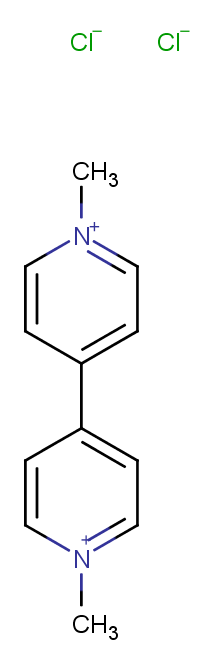

### 1912-24-9.png

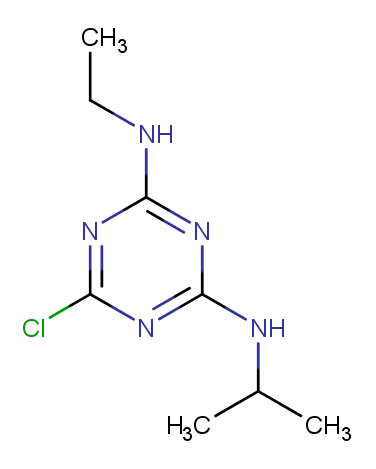

### 1912-26-1.png

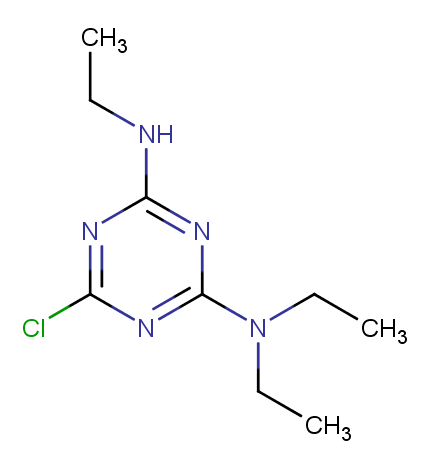

### 1918-00-9.png

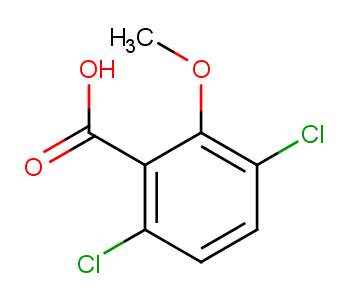

### 1918-02-1.png

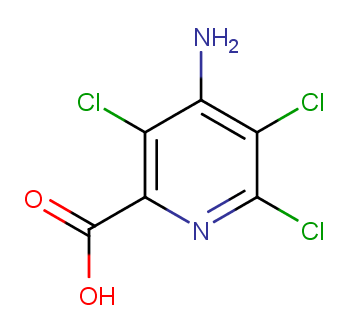

### 1918-11-2.png

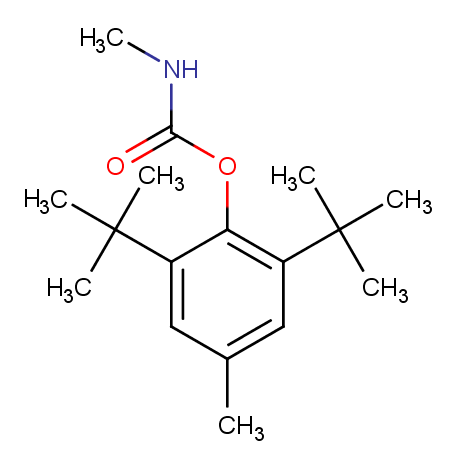

### 1918-13-4.png

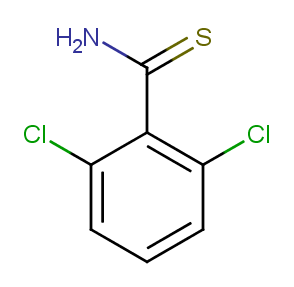

### 1918-16-7.png

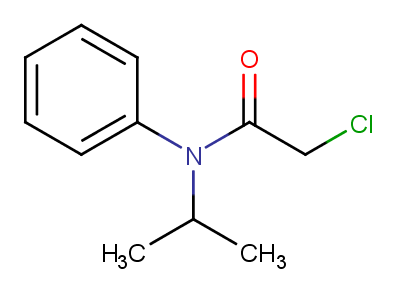

### 1929-77-7.png

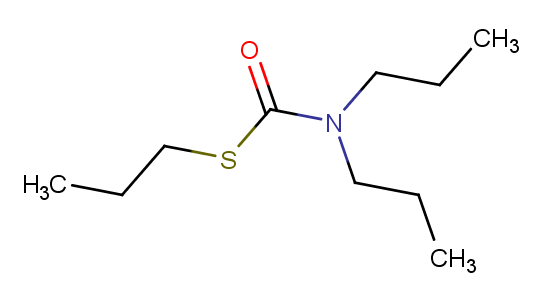

### 1982-47-4.png

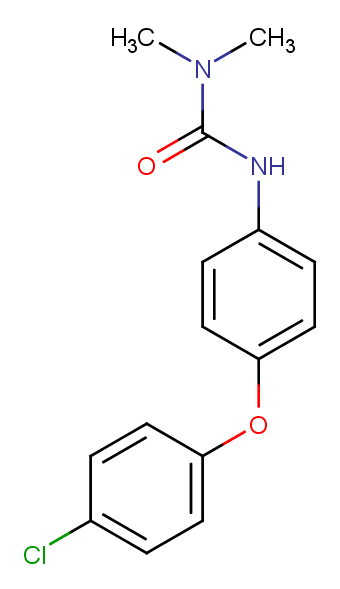

### 1982-49-6.png

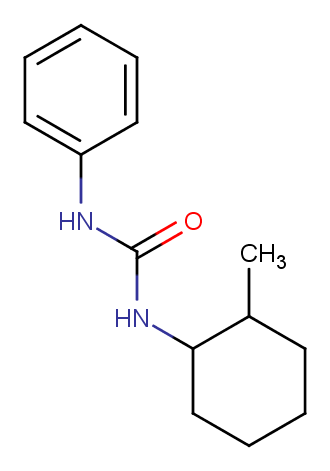

### 2008-41-5.png

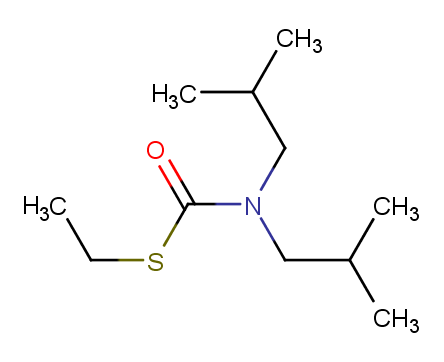

### 2163-80-6.png

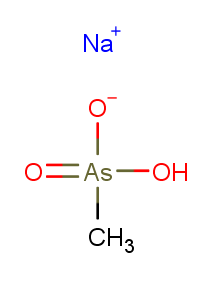

### 2164-08-1.png

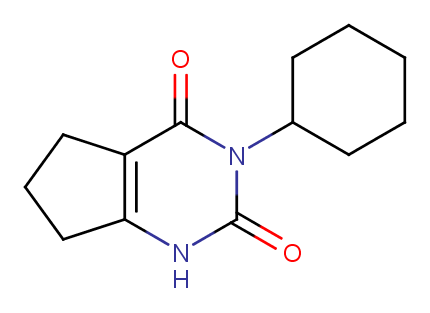

### 2164-17-2.png

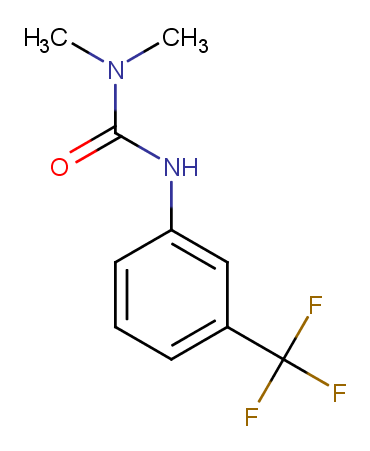

### 2212-67-1.png

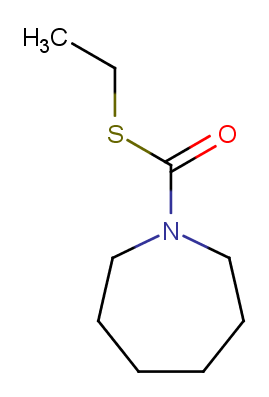

### 2303-16-4.png

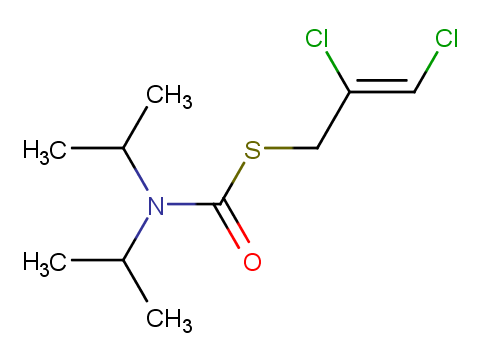

### 2303-17-5.png

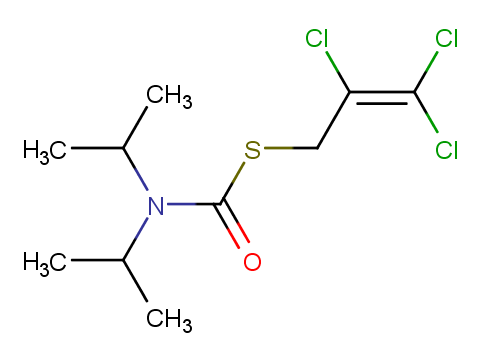

### 2307-68-8.png

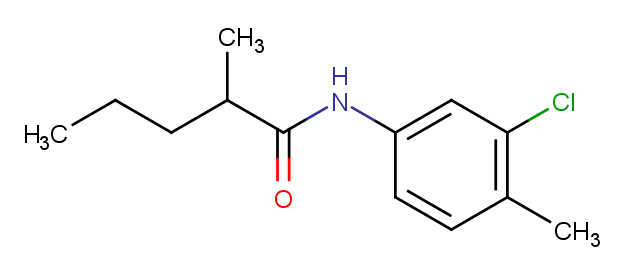

### 2464-37-1.png

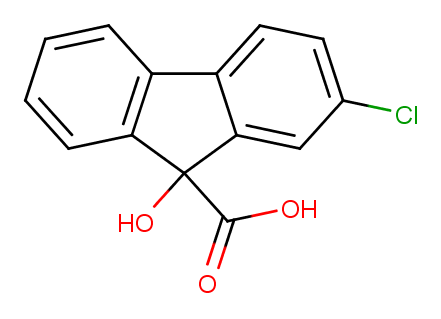

### 2693-61-0.png

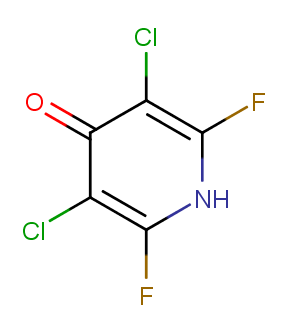

### 2797-51-5.png

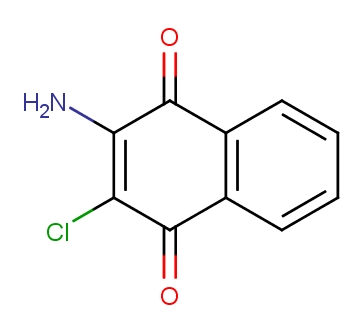

### 2941-55-1.png

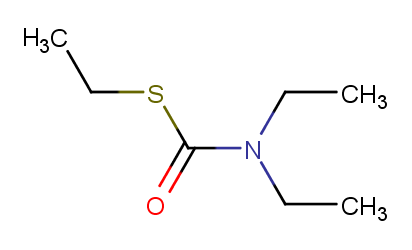

### 3060-89-7.png

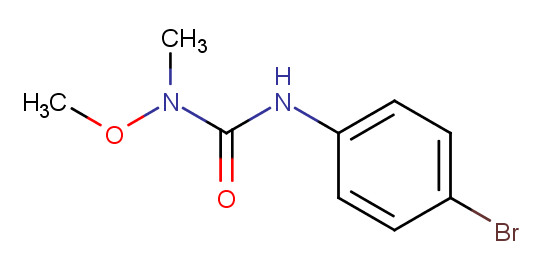

### 3134-70-1.png

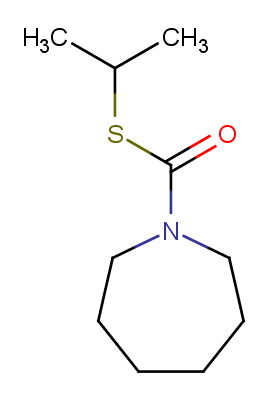

### 3337-71-1.png

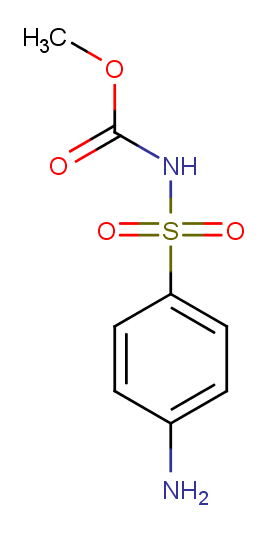

### 15972-60-8.png

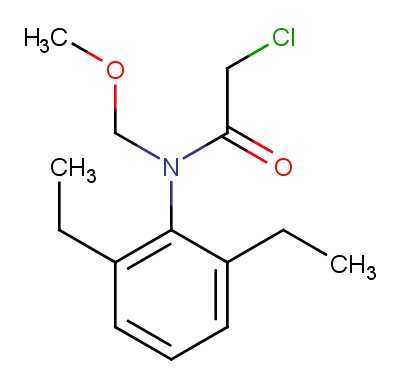

### 16118-49-3.png

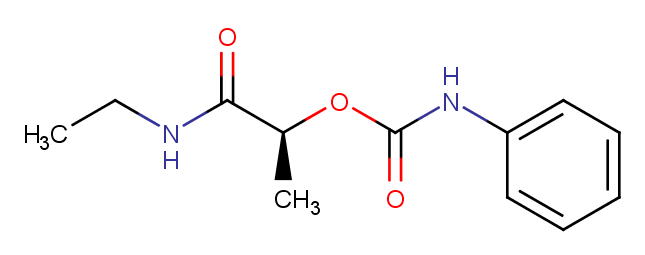

### 18691-97-9.png

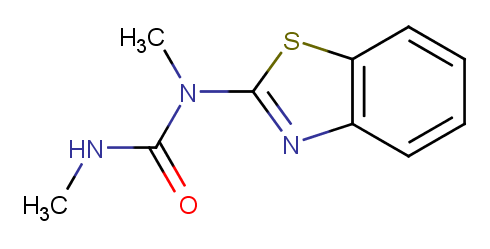

### 19044-88-3.png

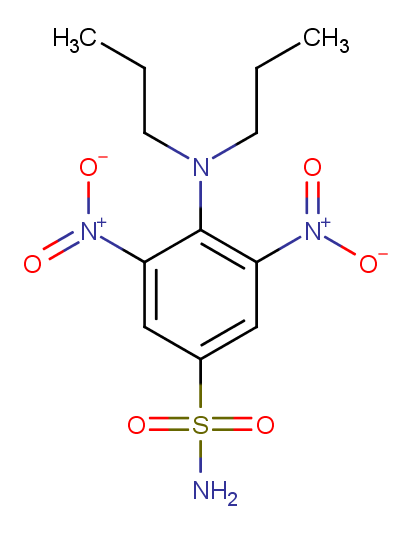

### 19666-30-9.png

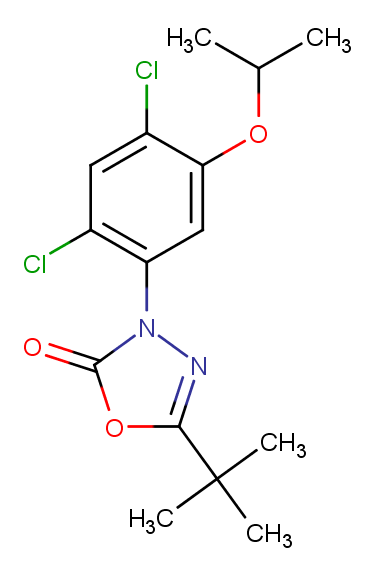

### 19937-59-8.png

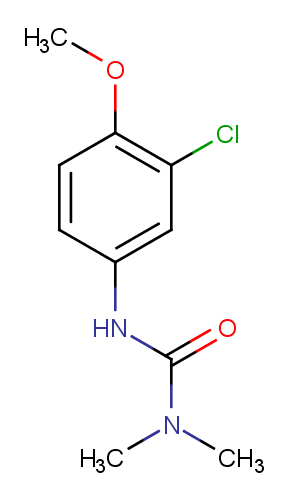

### 20354-26-1.png

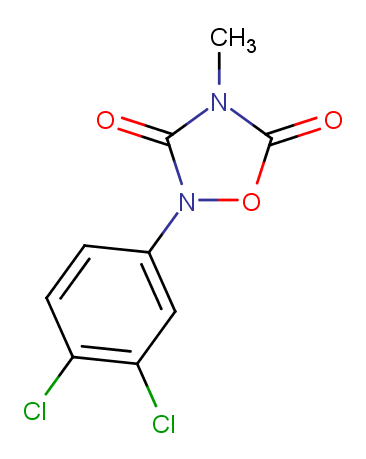

### 21087-64-9.png

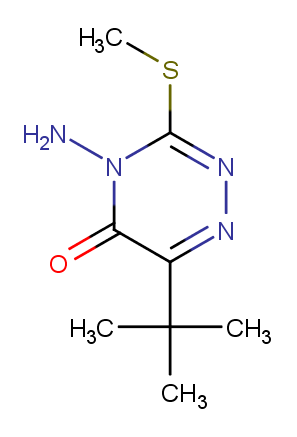

### 21725-46-2.png

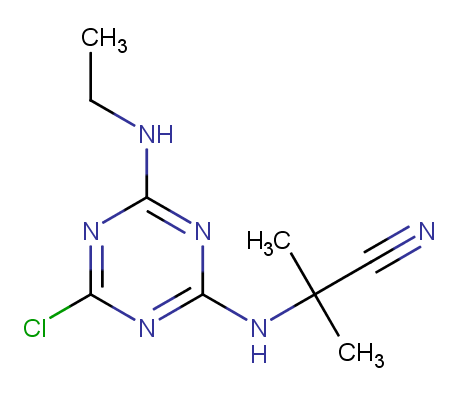

### 22936-75-0.png

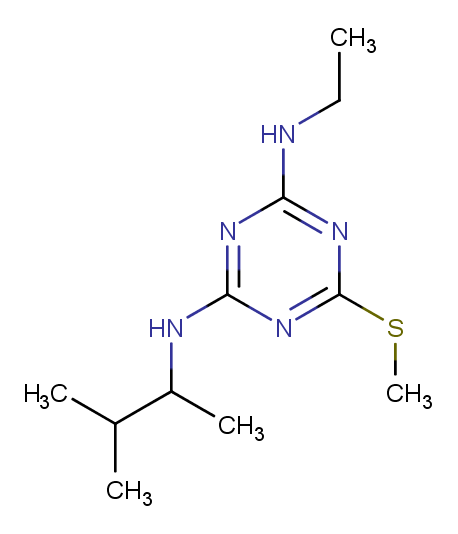

### 23184-66-9.png

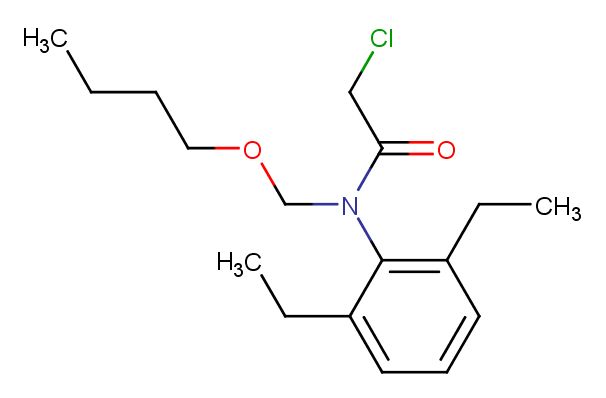

### 23576-23-0.png

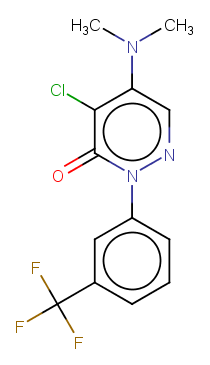

### 23950-58-5.png

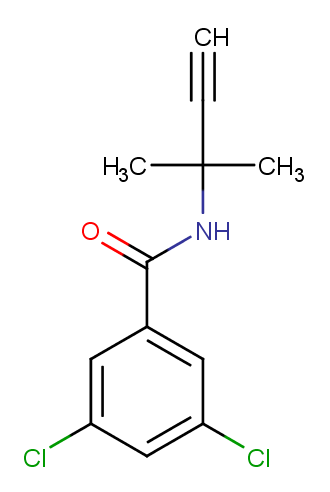

### 24151-93-7.png

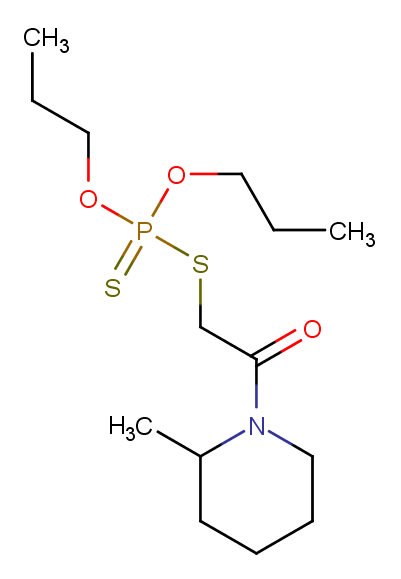

### 25057-89-0.png

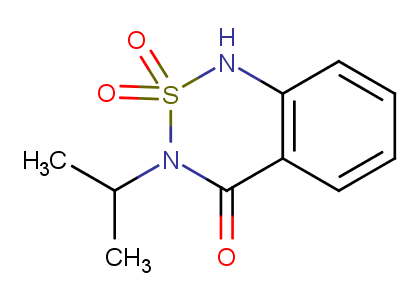

### 25059-80-7.png

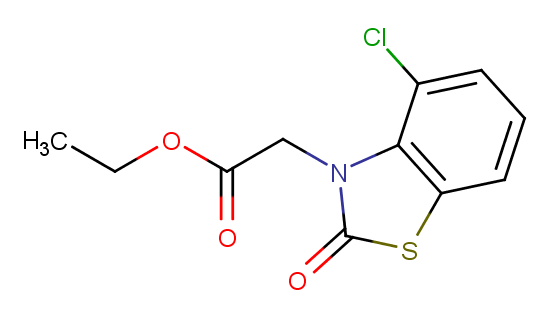

### 25319-90-8.png

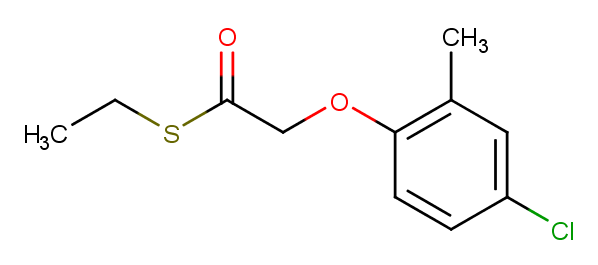

### 25366-23-8.png

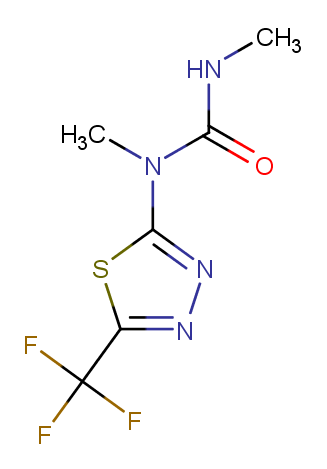

### 26225-79-6.png

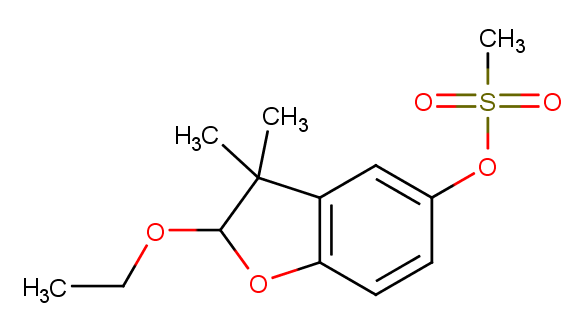

### 26399-36-0.png

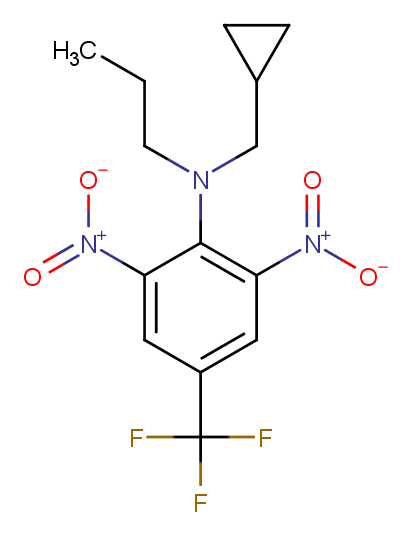

### 27314-13-2.png

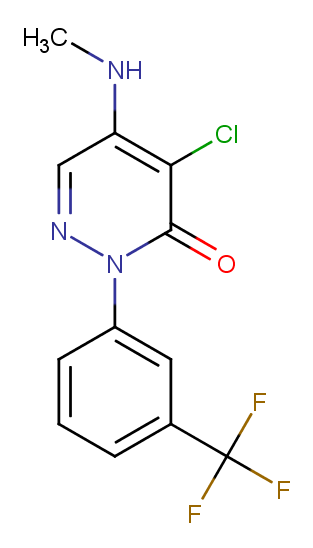

### 28249-77-6.png

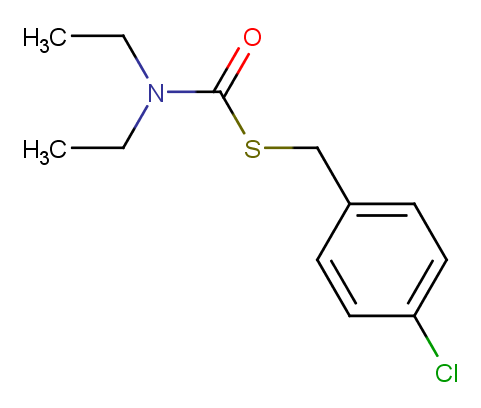

### 29091-05-2.png

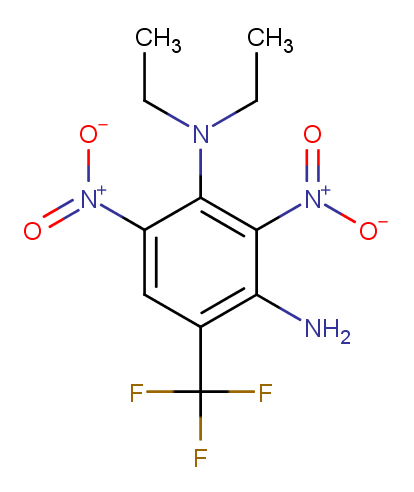

### 29091-21-2.png

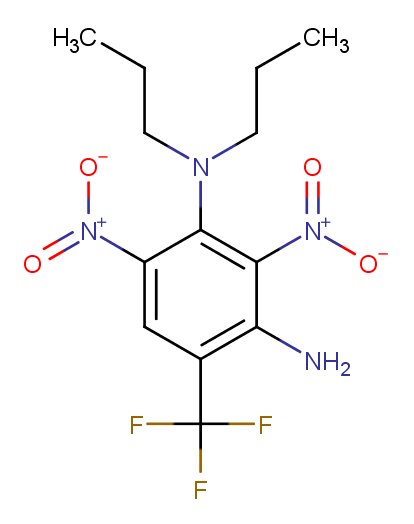

### 30043-49-3.png

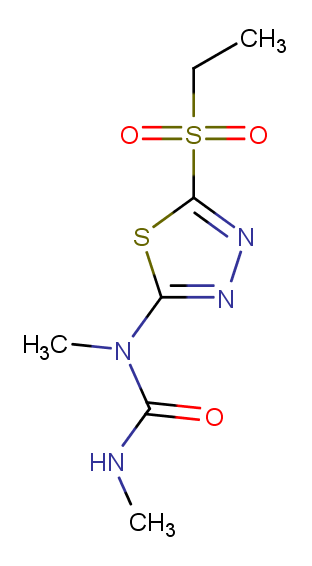

### 32861-85-1.png

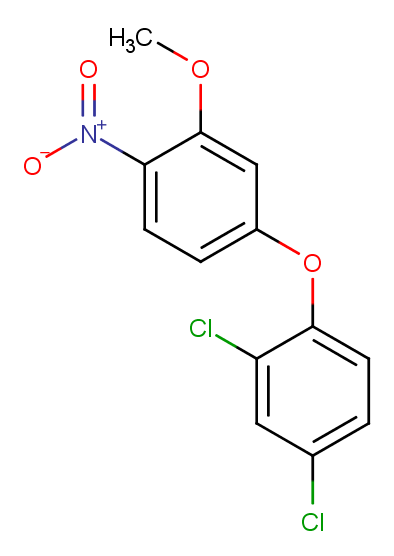

### 33245-39-5.png

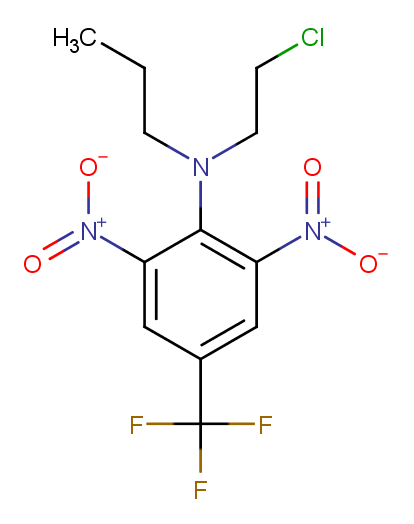

### 33629-47-9.png

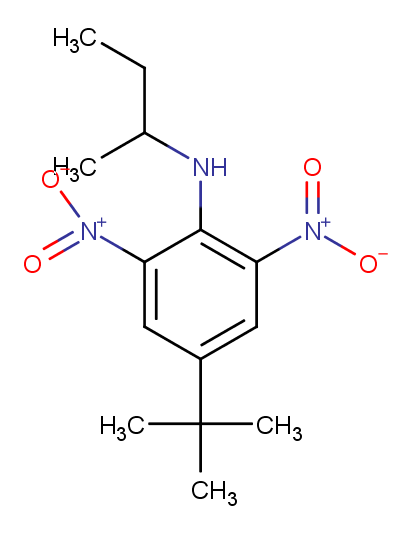

### 33693-04-8.png

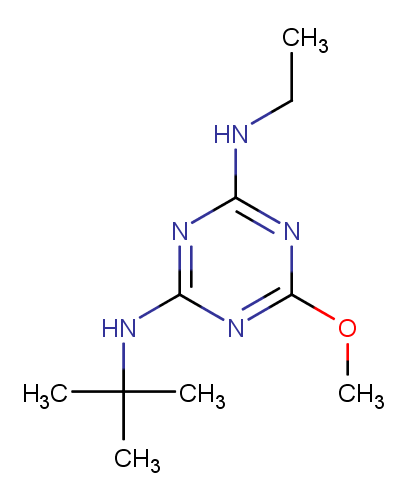

### 33820-53-0.png

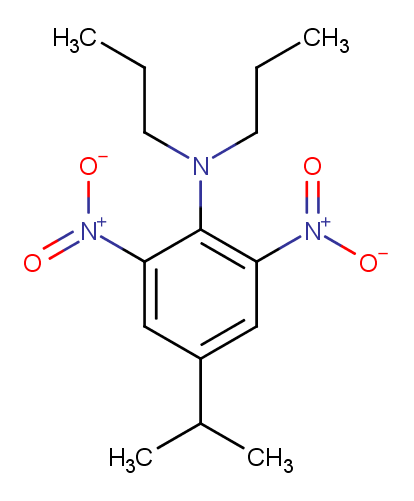

### 34014-18-1.png

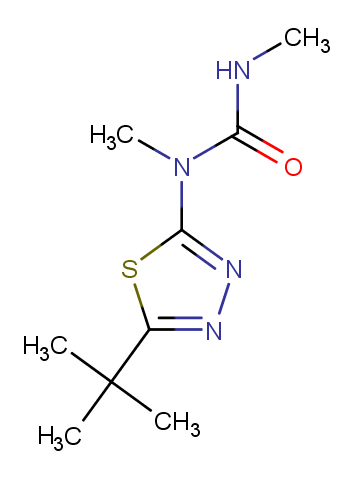

### 34123-59-6.png

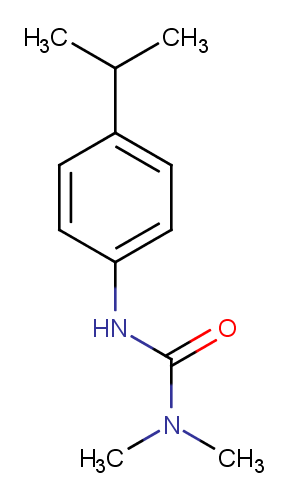

### 34205-21-5.png

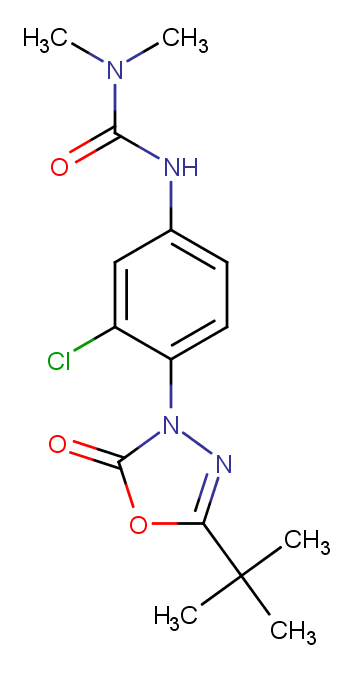

### 34256-82-1.png

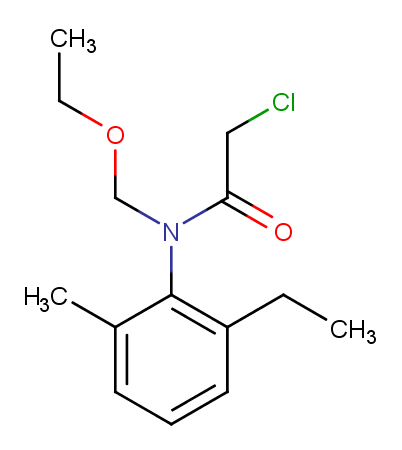

### 34622-58-7.png

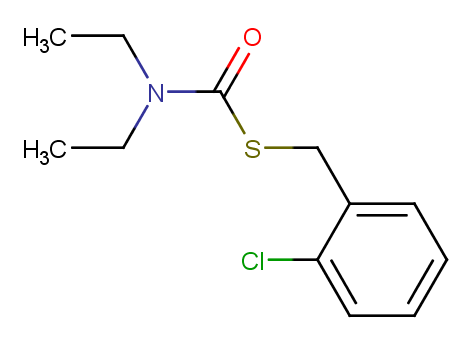

### 35256-85-0.png

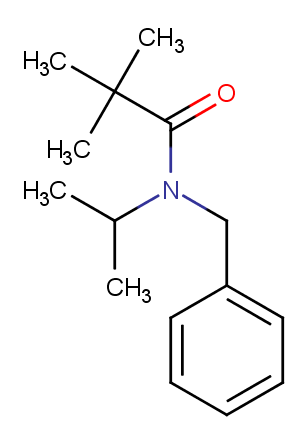

### 156963-66-5.png

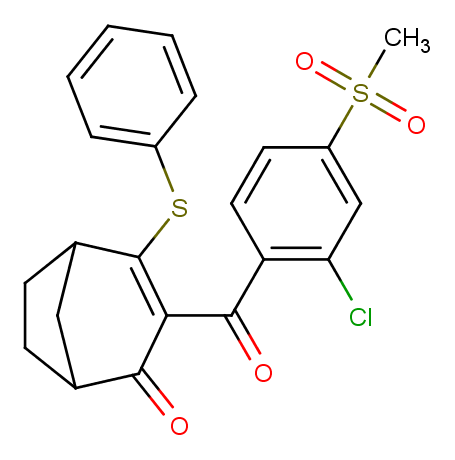

### 158237-07-1.png

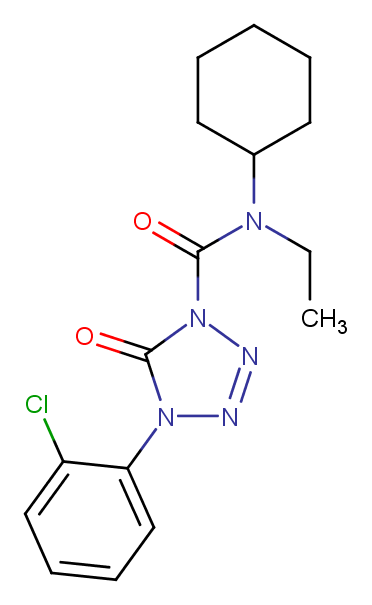

### 158353-15-2.png

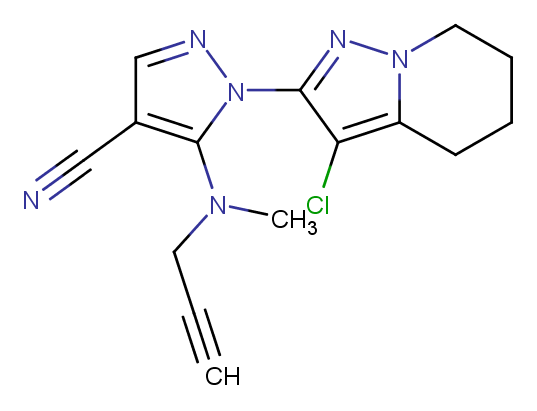

### 158755-95-4.png

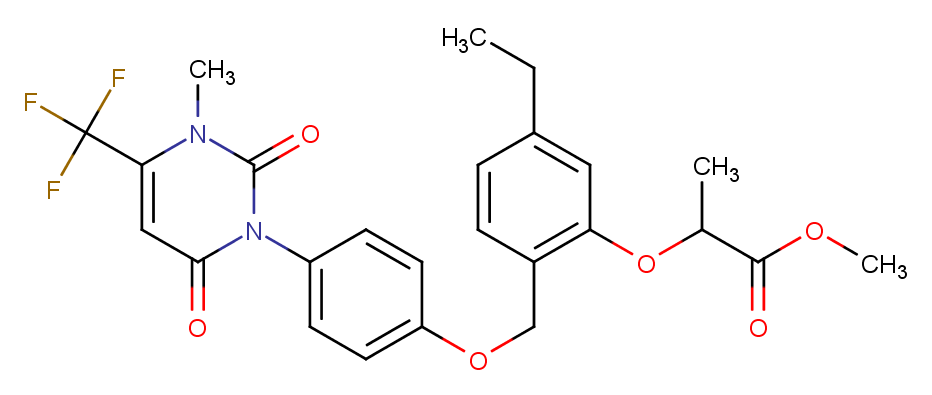

### 168088-61-7.png

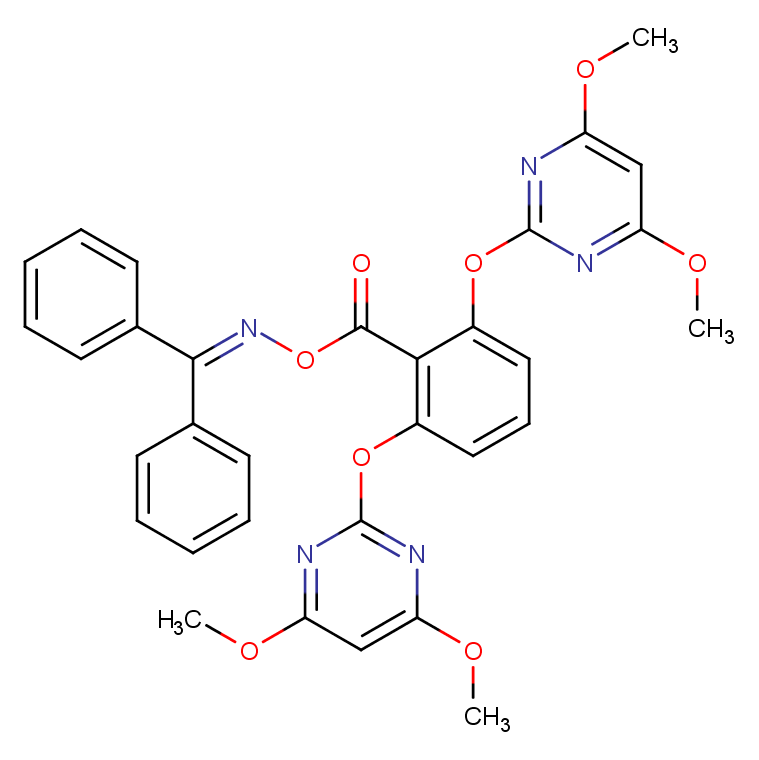

### 173159-57-4.png

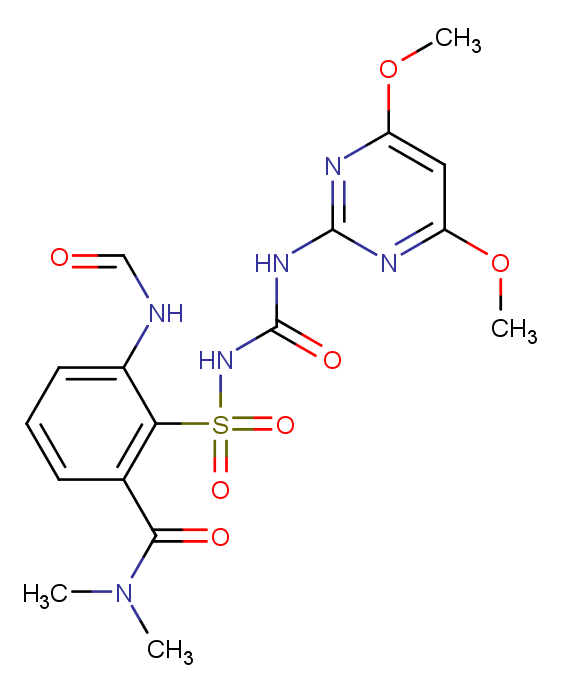

### 173980-17-1.png

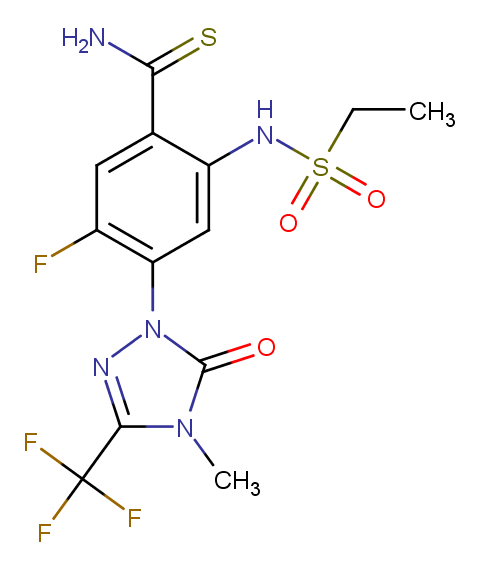

### 174514-07-9.png

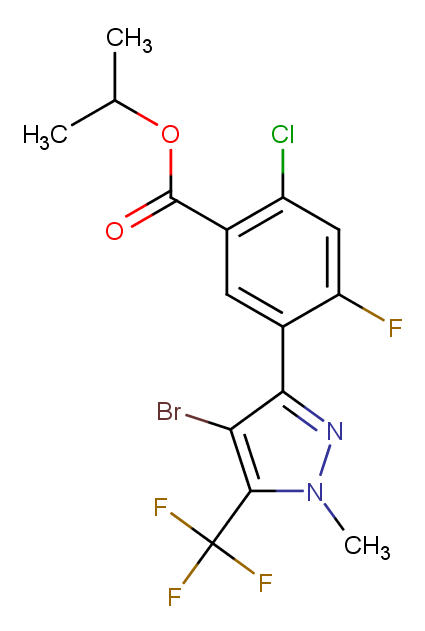

### 181274-15-7.png

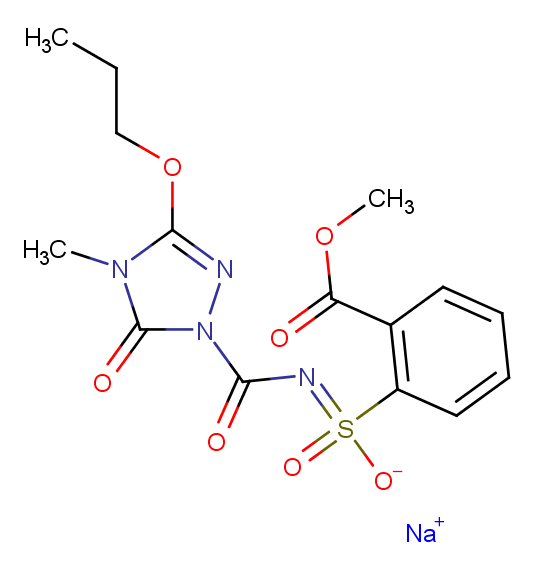

### 181274-17-9.png

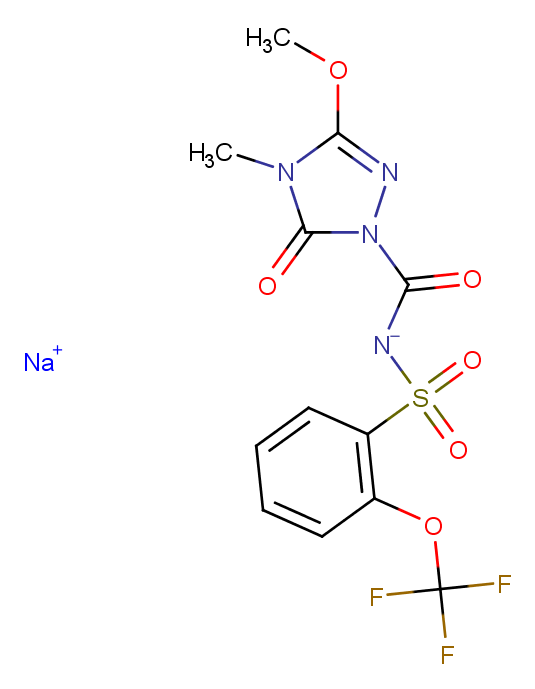

### 185119-76-0.png

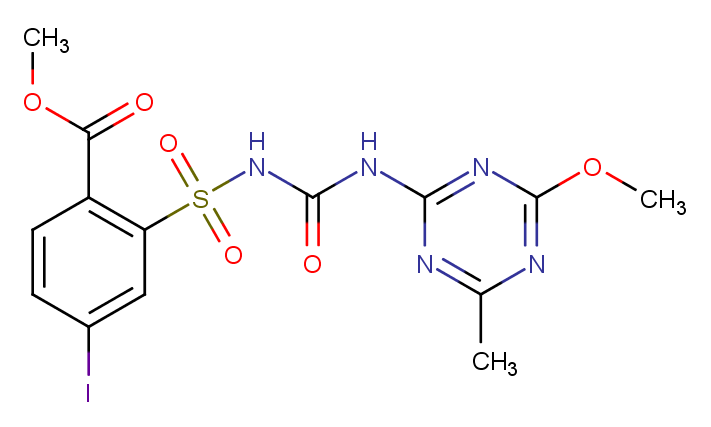

### 188489-07-8.png

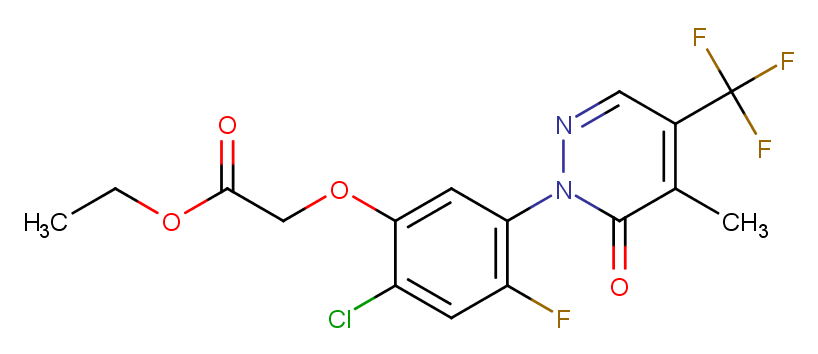

### 190314-43-3.png

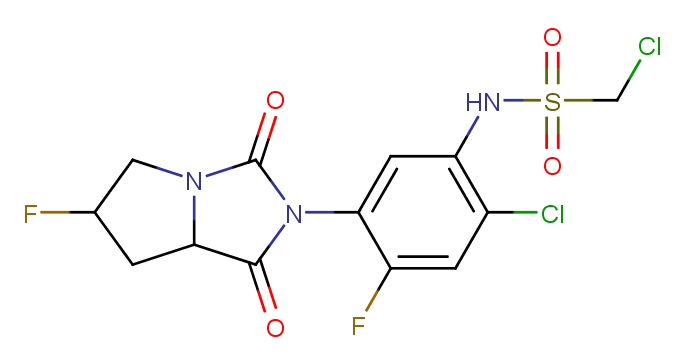

### 200509-41-7.png

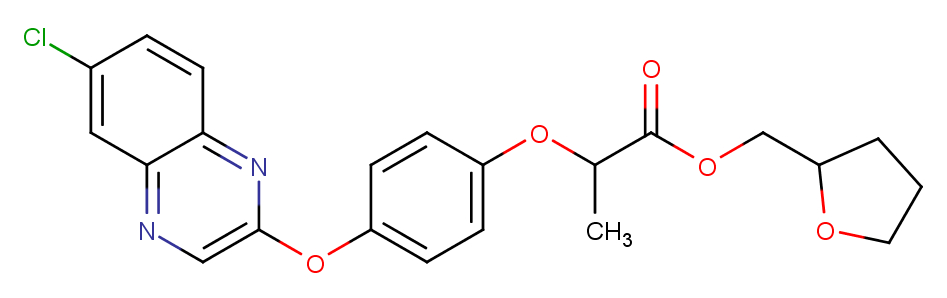

### 210631-68-8.png

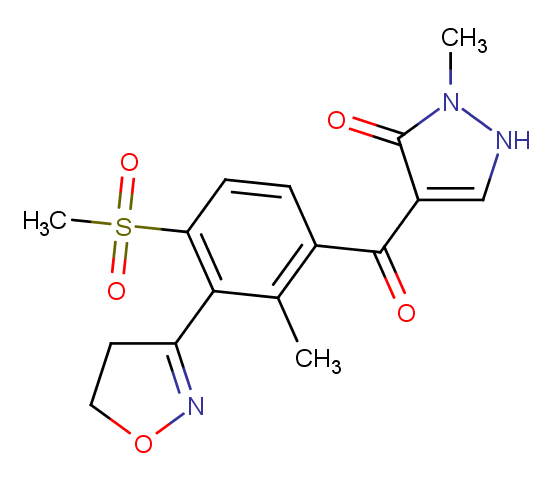

### 212201-70-2.png

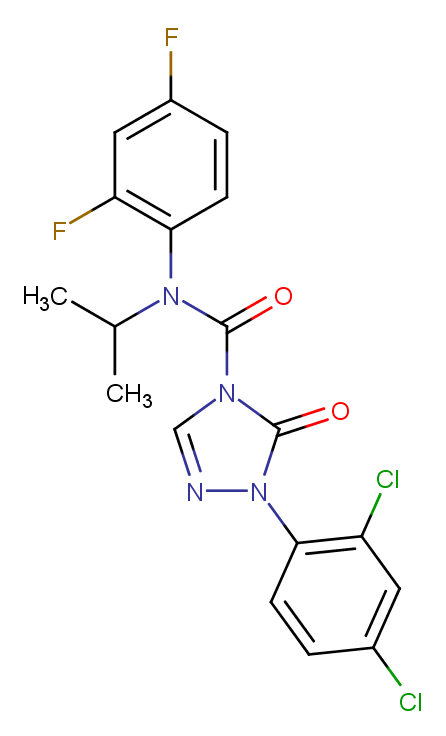

### 213464-77-8.png

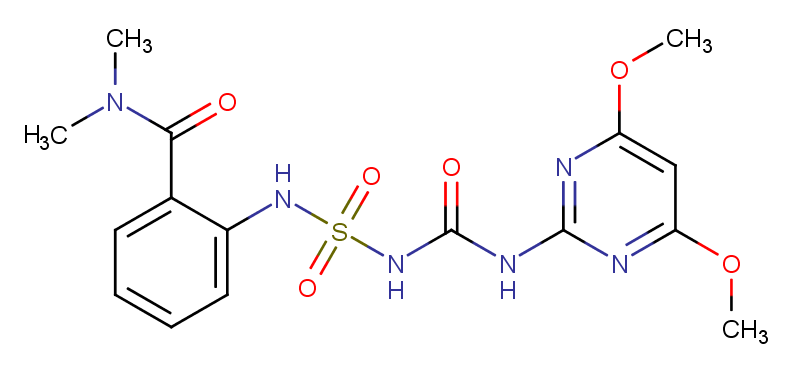

### 215655-76-8.png

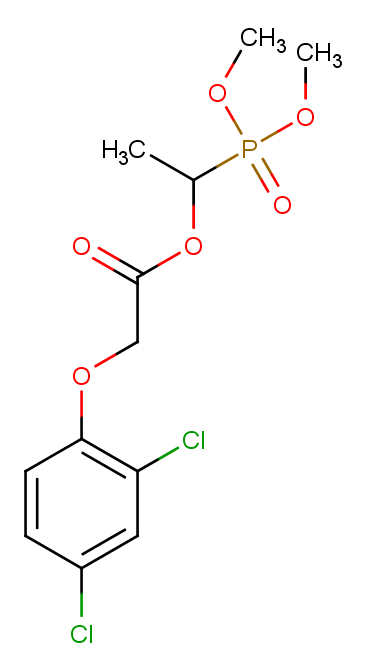

### 219714-96-2.png

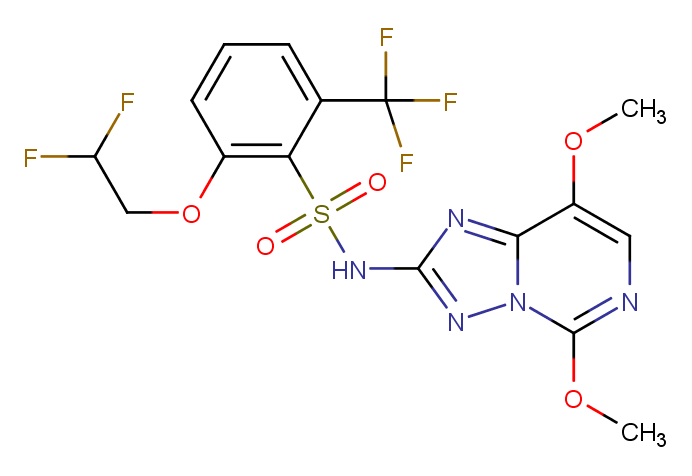

### 221205-90-9.png

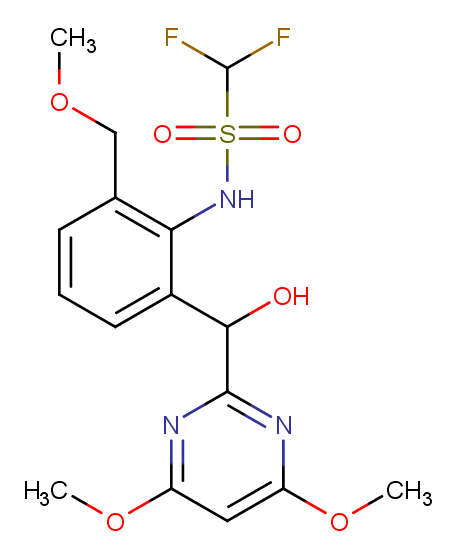

### 243973-20-8.png

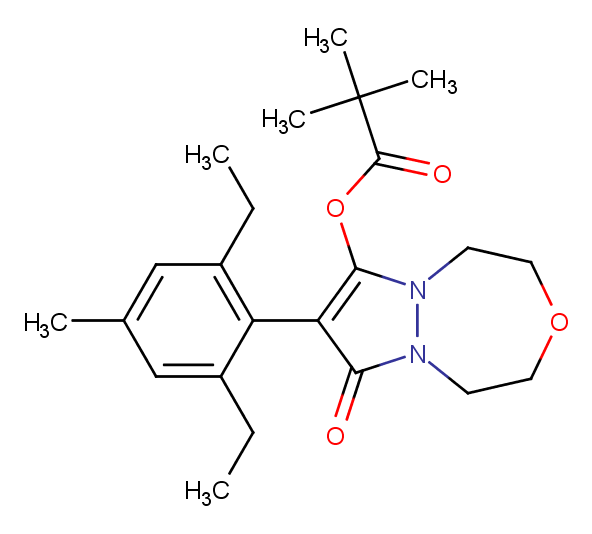

### 335104-84-2.png

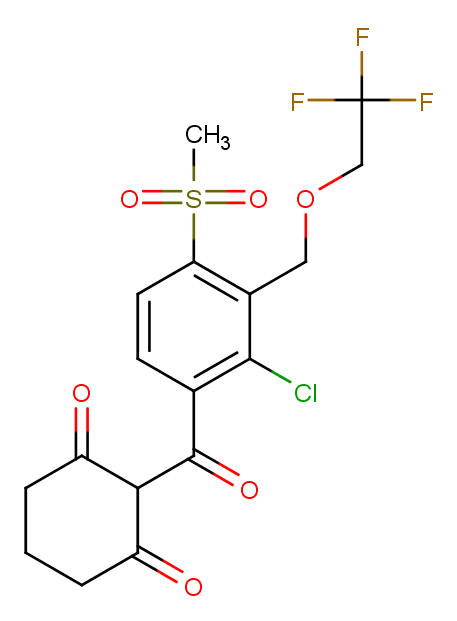

### 352010-68-5.png

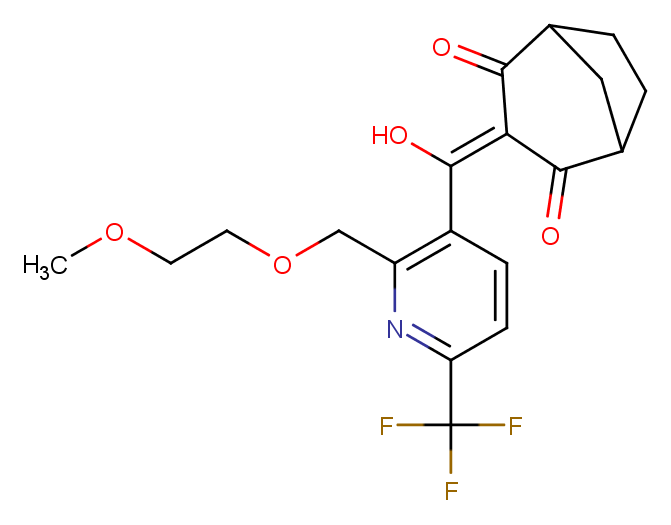

### 2053901-33-8.png

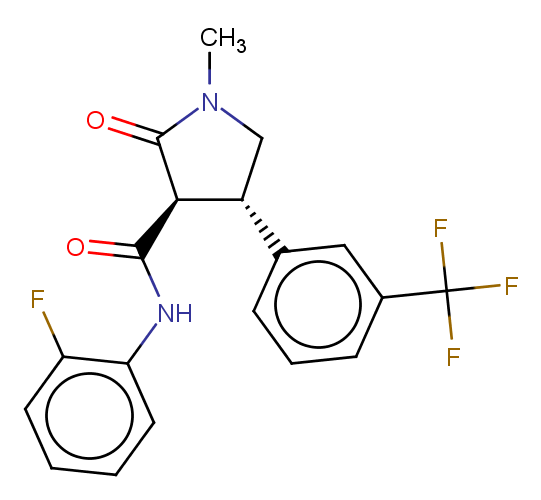

### MMV000248.png

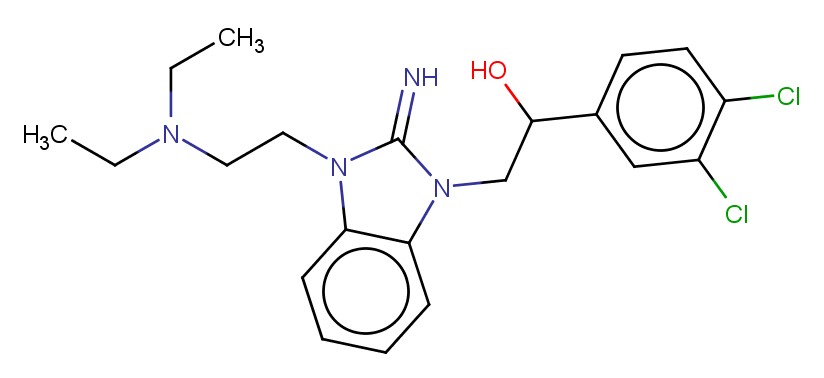
