## Supplementary figures and images for "Refining physico-chemical rules for herbicides using an antimalarial library"

### 101-21-3.png

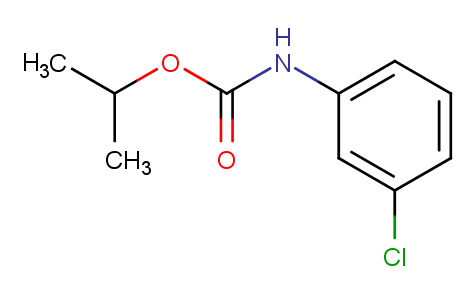

### 101-42-8.png

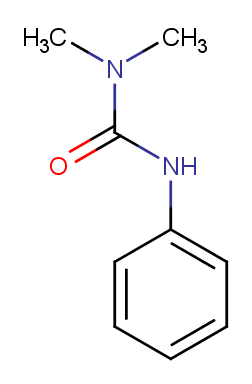

### 107-02-8.png

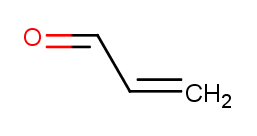

### 112-05-0.png

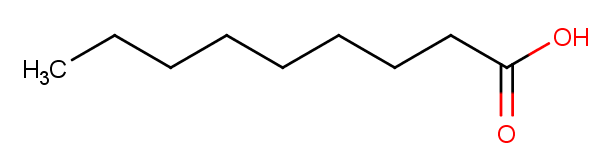

### 112-80-1.png

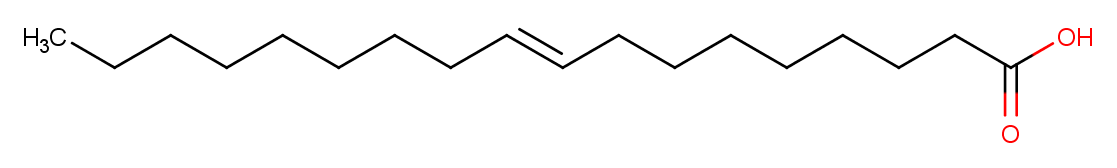

### 120-36-5.png

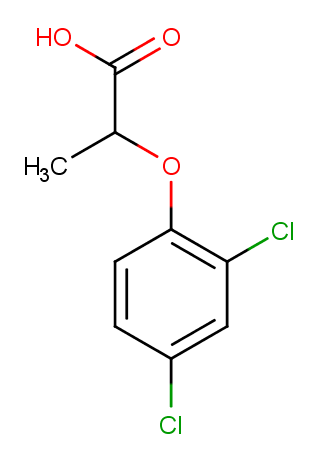

### 122-34-9.png

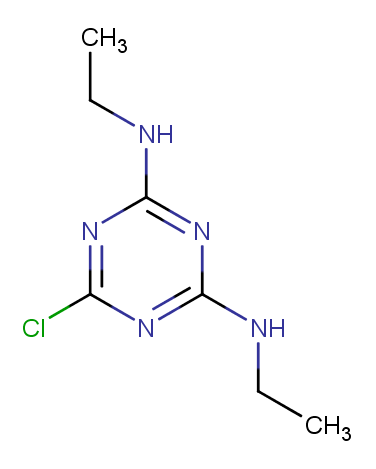

### 122-42-9.png

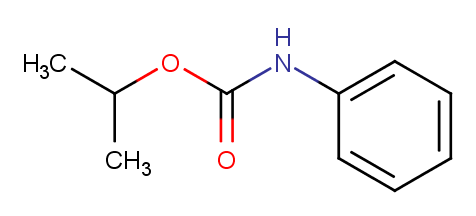

### 122-88-3.png

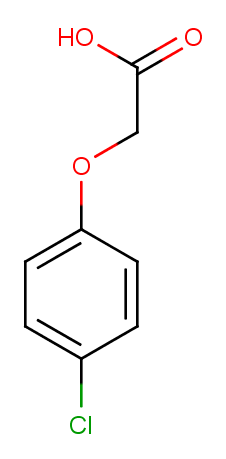

### 132-66-1.png

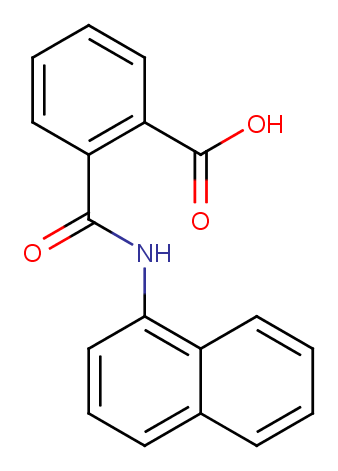

### 133-90-4.png

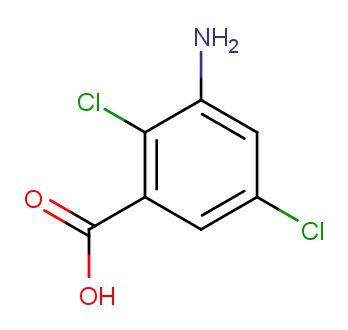

### 139-40-2.png

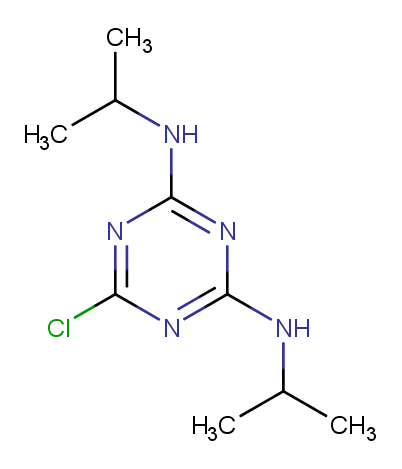

### 144-21-8.png

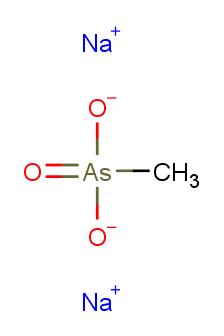

### 144-54-7.png

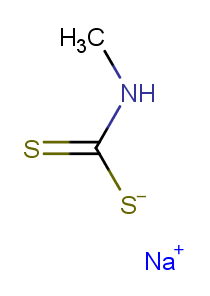

### 145-73-3.png

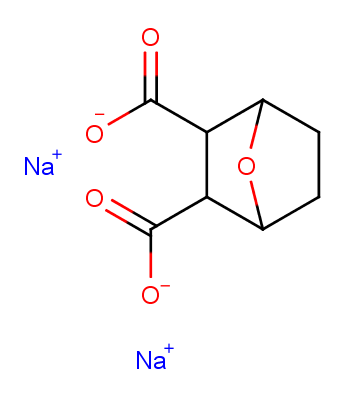

### 150-68-5.png

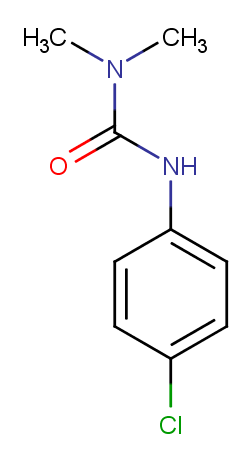

### 1014-69-3.png

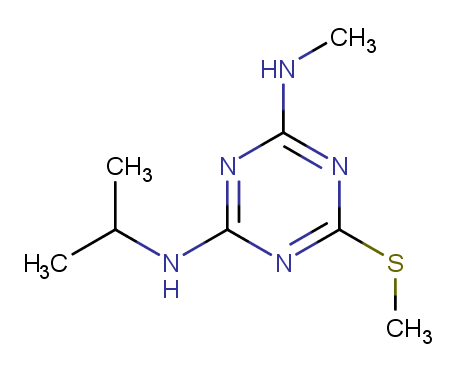

### 1014-70-6.png

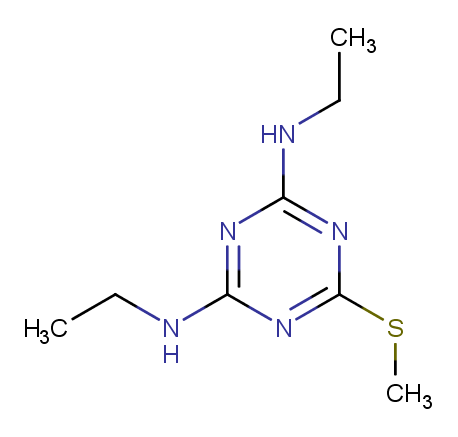

### 1071-83-6.png

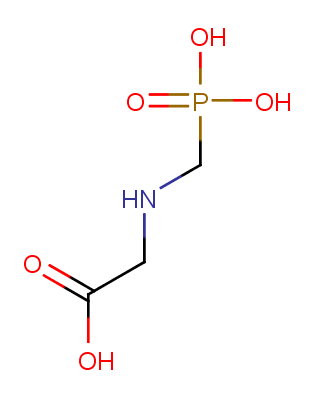

### 1114-71-2.png

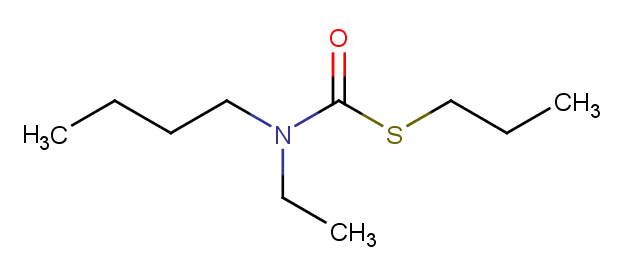

### 1134-23-2.png

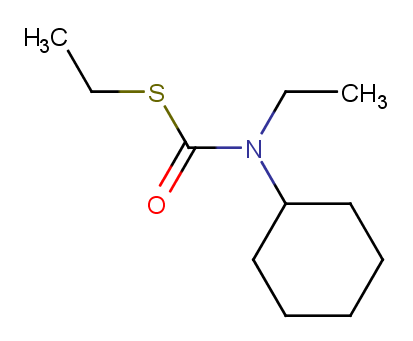

### 1194-65-6.png

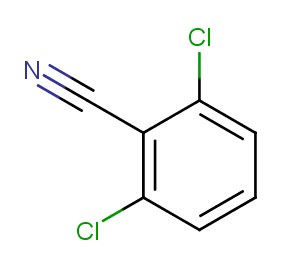

### 1420-07-1.png

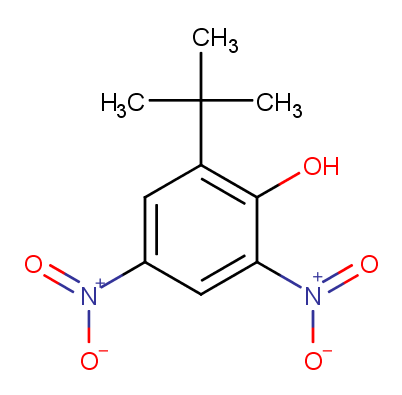

### 13181-17-4.png

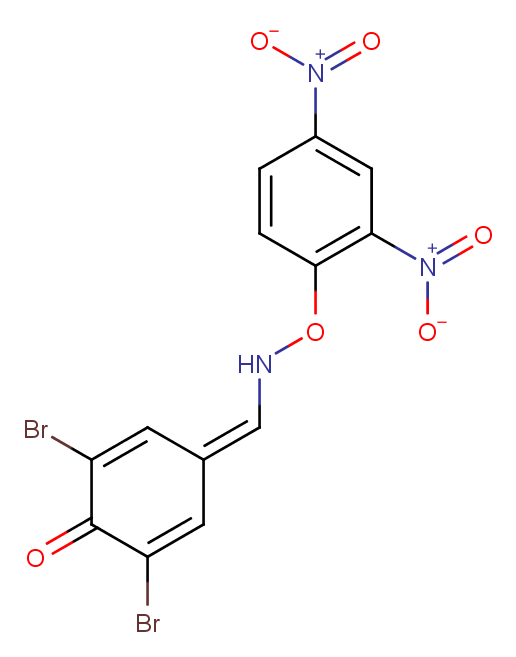

### 13360-45-7.png

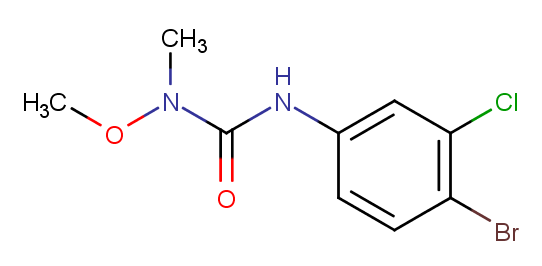

### 13684-56-5.png

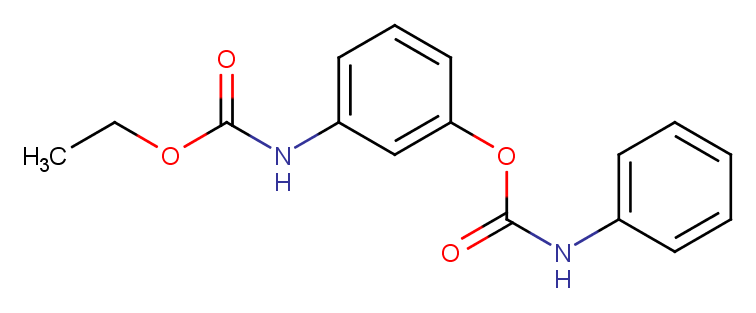

### 13684-63-4.png

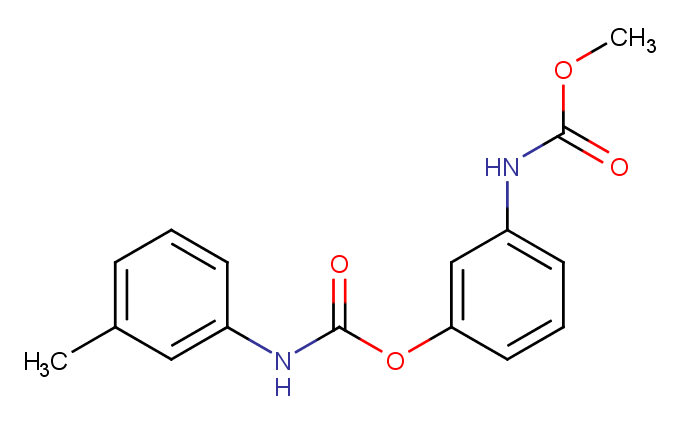

### 15299-99-7.png

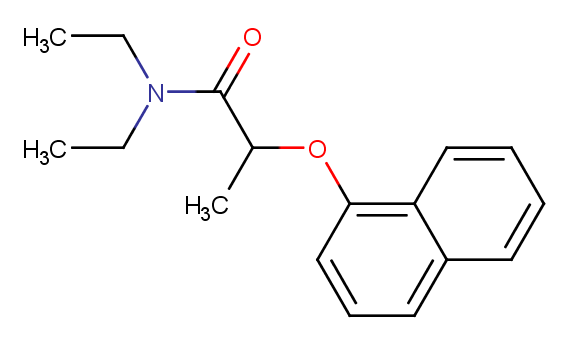

### 15545-48-9.png

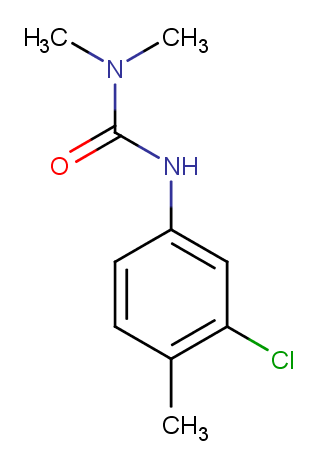

### 100646-51-3.png

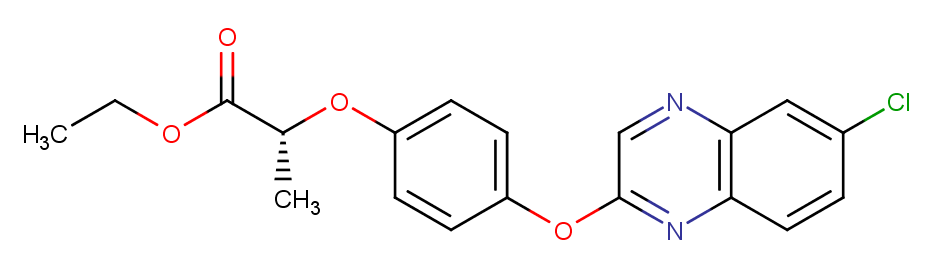

### 100784-20-1.png

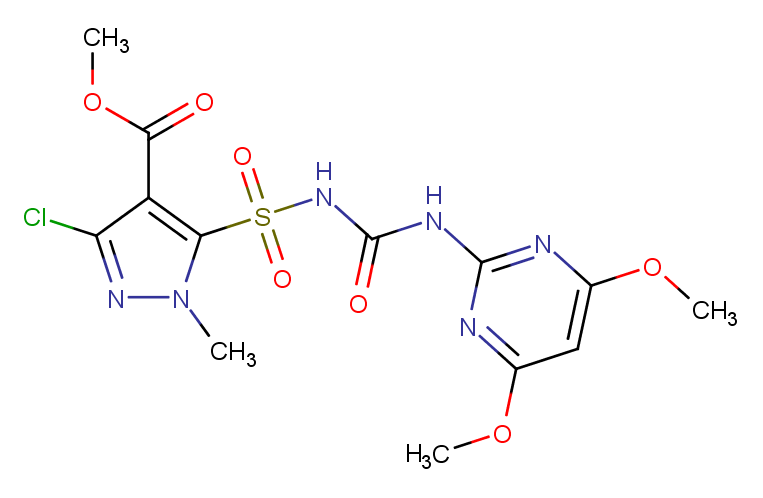

### 101200-48-0.png

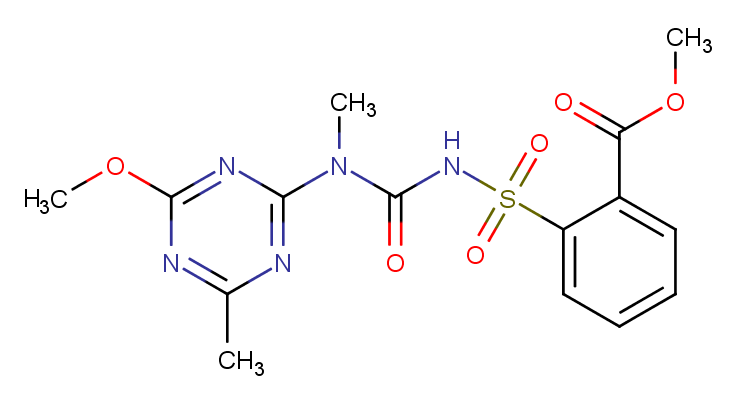

### 101205-02-1.png

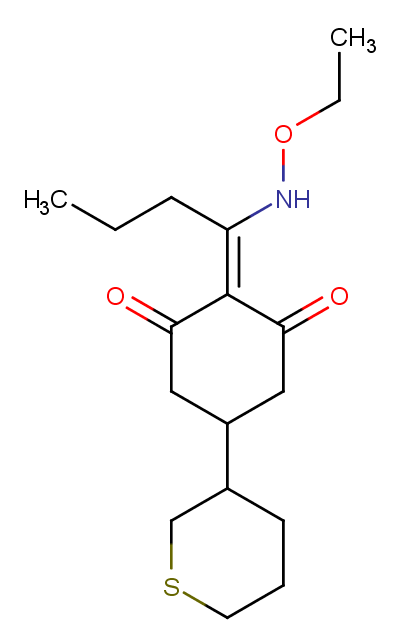

### 103361-09-7.png

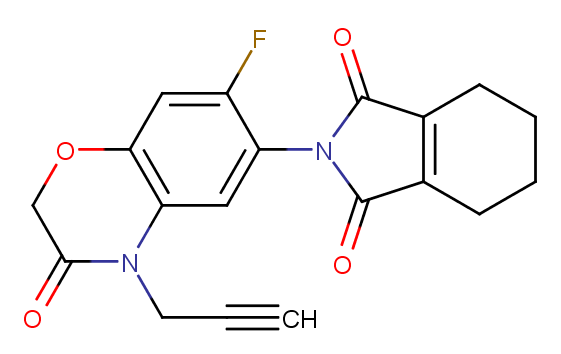

### 104040-78-0.png

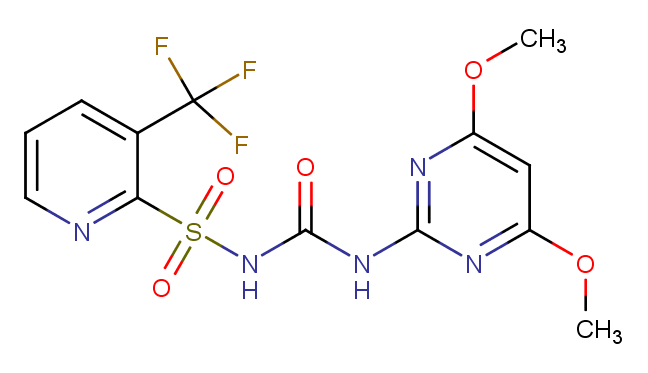

### 104098-48-8.png

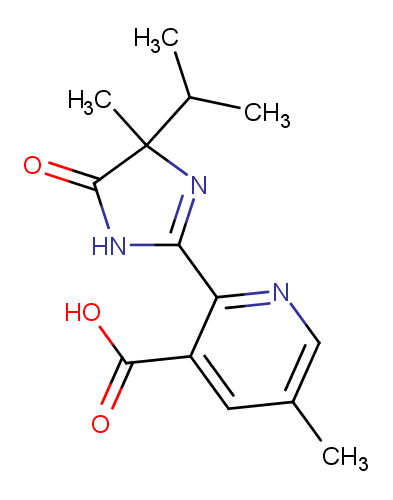

### 104206-82-8.png

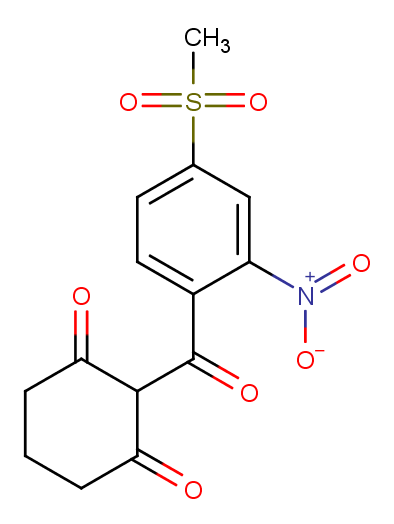

### 104770-29-8.png

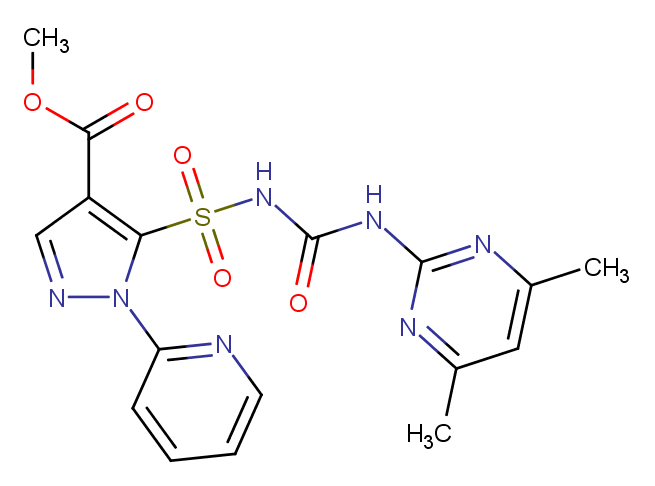

### 105512-06-9.png

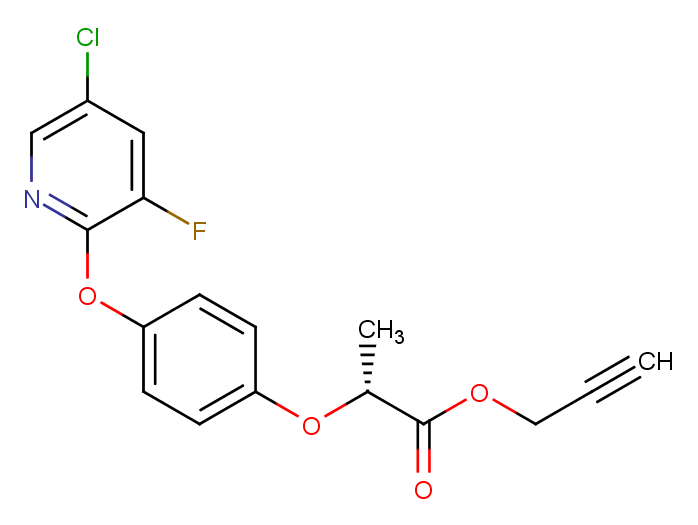

### 106700-29-2.png

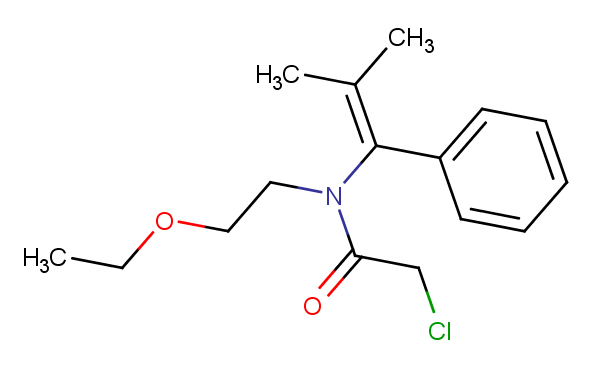

### 109293-97-2.png

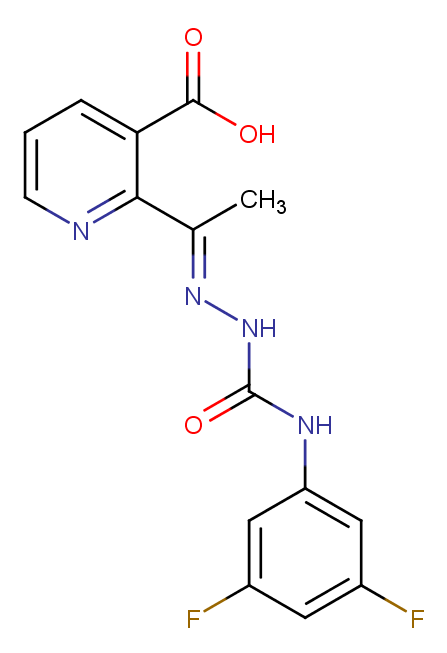

### 110956-75-7.png

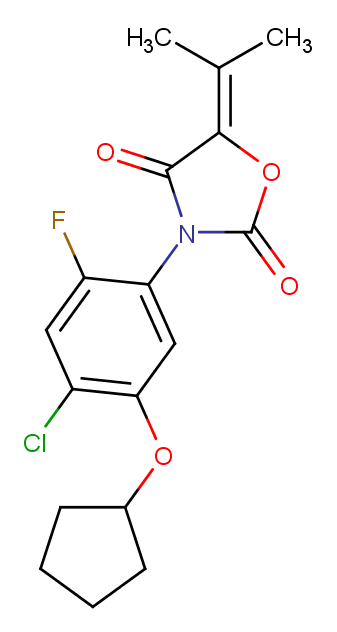

### 111479-05-1.png

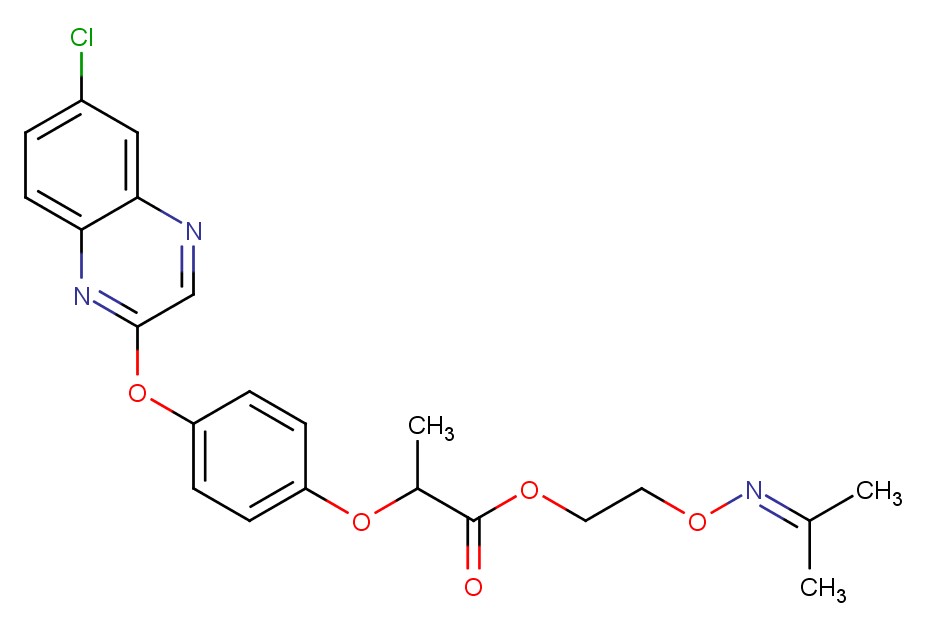

### 111578-32-6.png

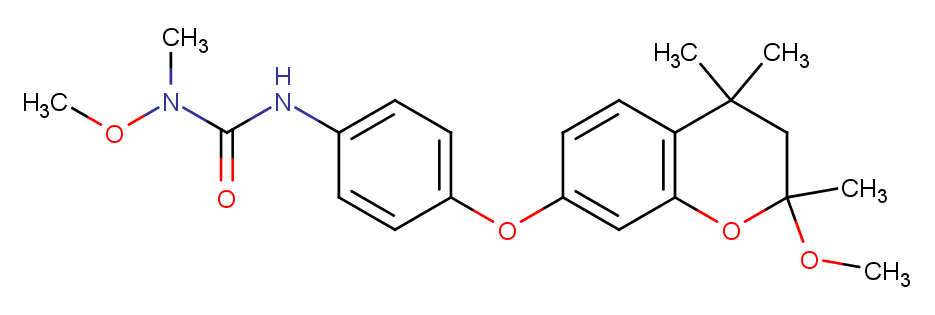

### 111991-09-4.png

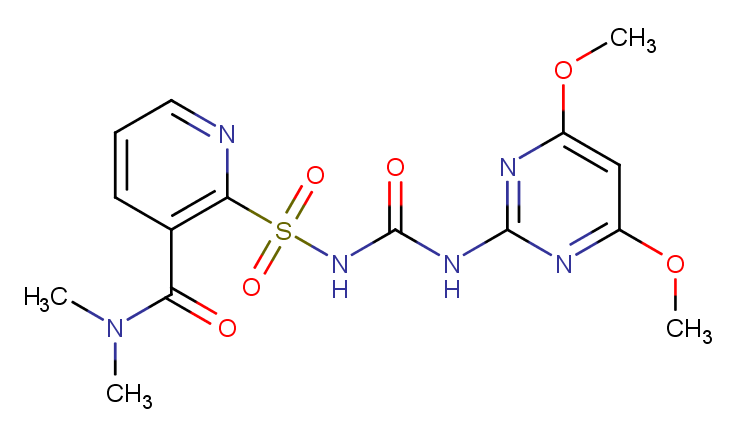

### 113614-08-7.png

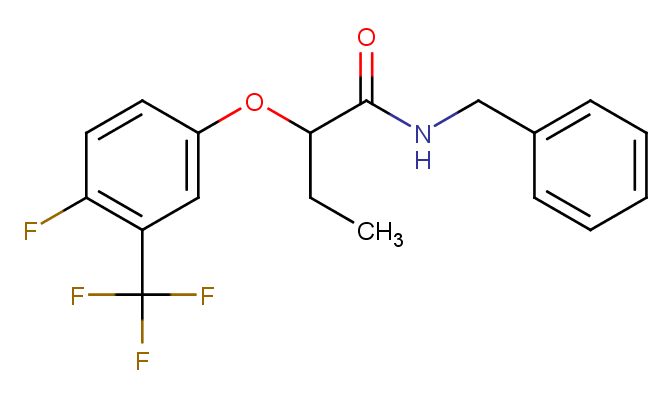

### 114311-32-9.png

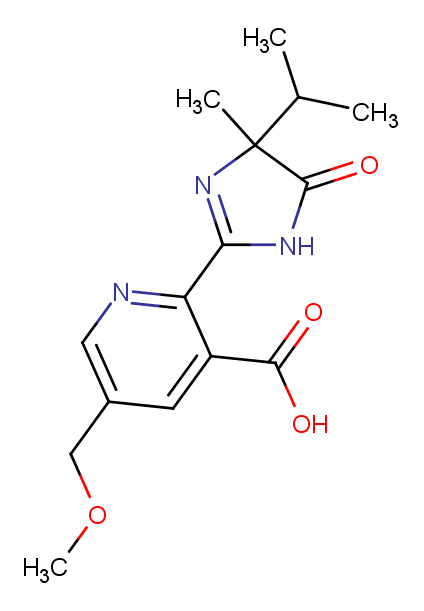

### 114420-56-3.png

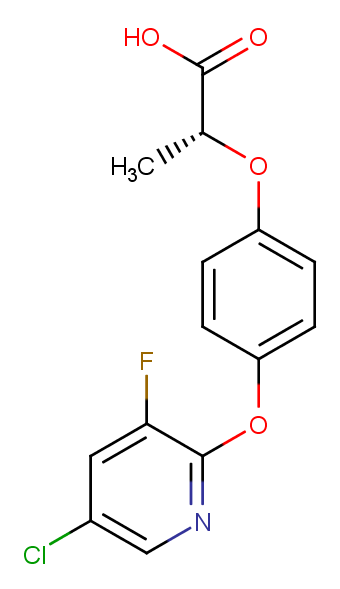

### 117337-19-6.png

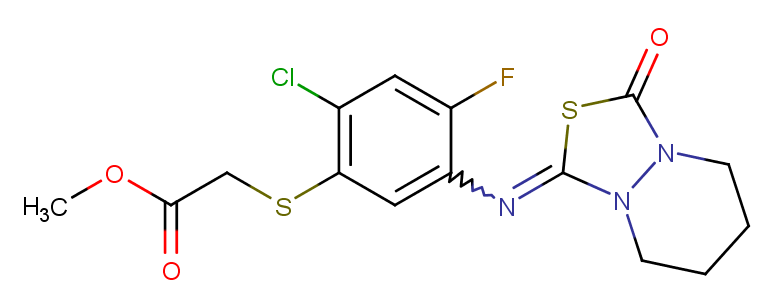

### 117718-60-2.png

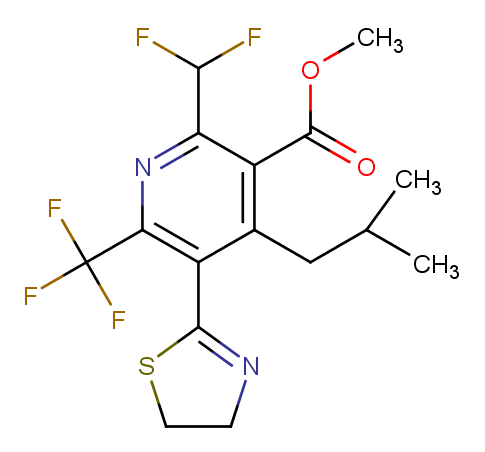

### 119126-15-7.png

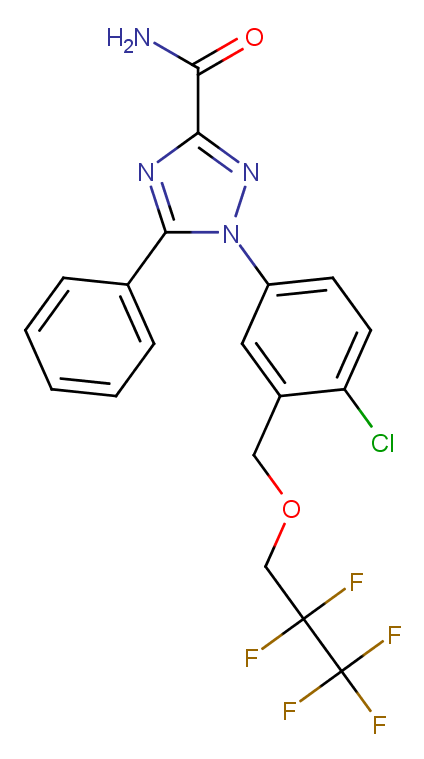

### 120162-55-2.png

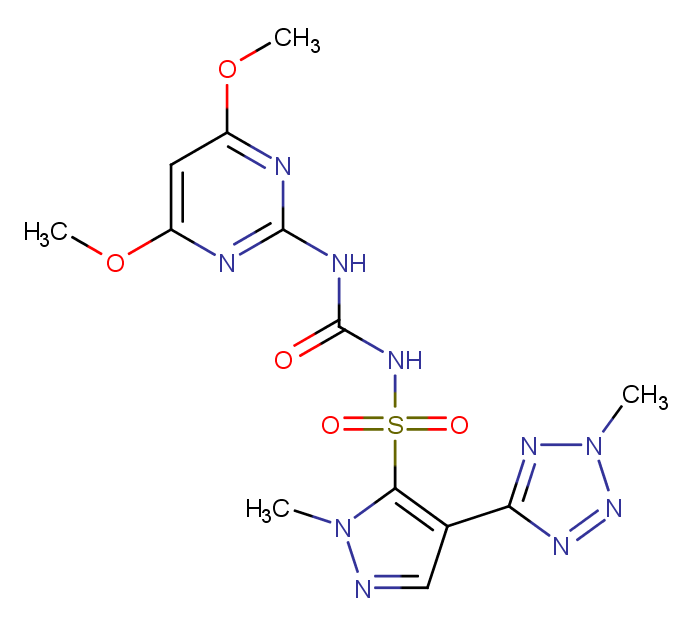

### 120890-70-2.png

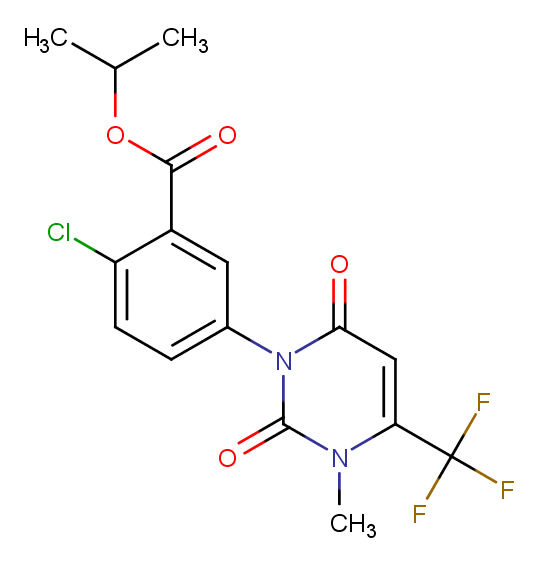

### 120923-37-7.png

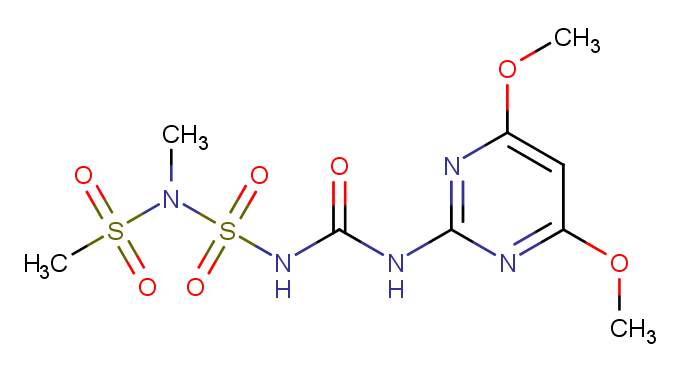

### 122008-85-9.png

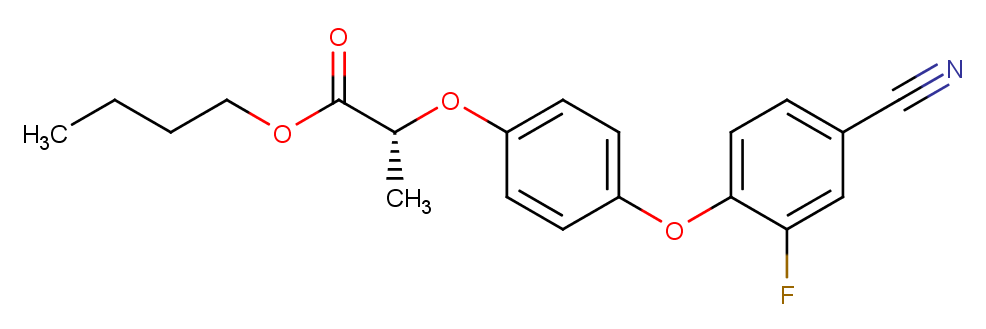

### 122548-33-8.png

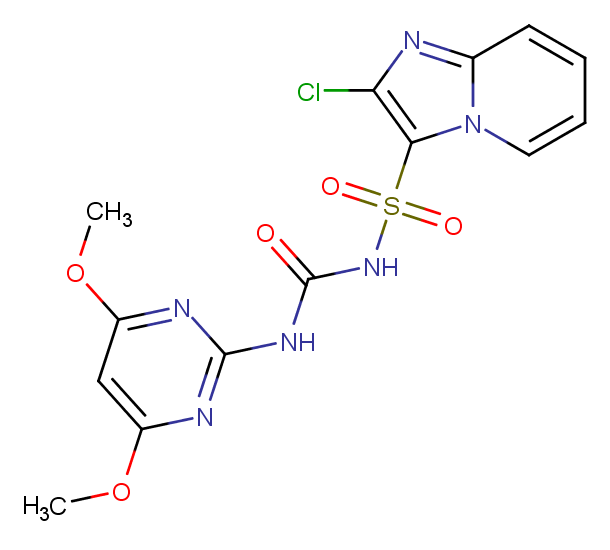

### 122836-35-5.png

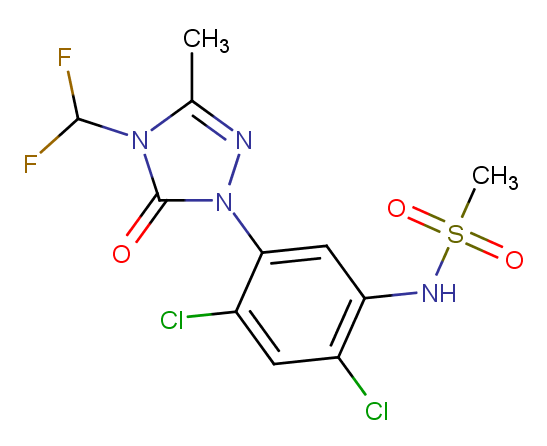

### 122931-48-0.png

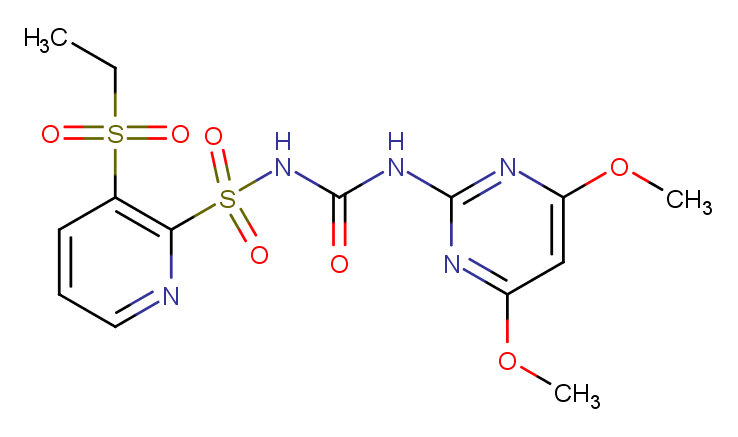

### 123249-43-4.png

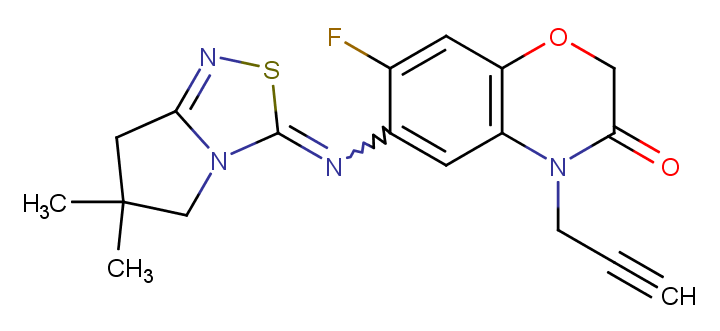

### 123342-93-8.png

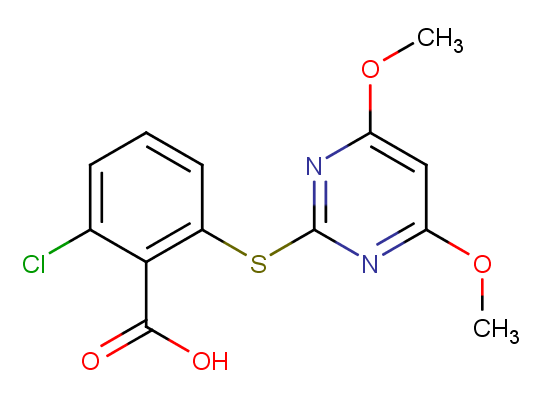

### 125306-83-4.png

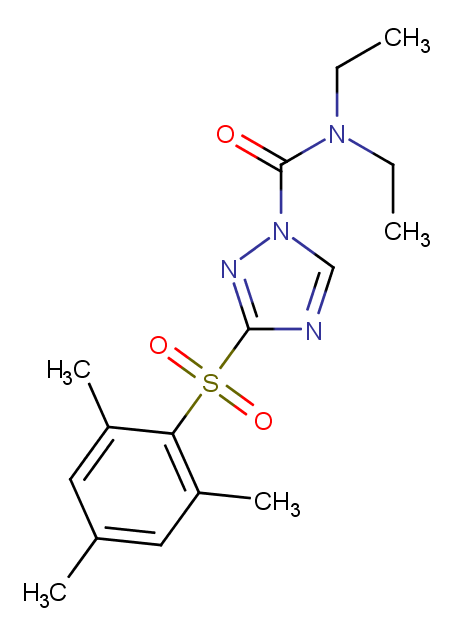

### 125401-75-4.png

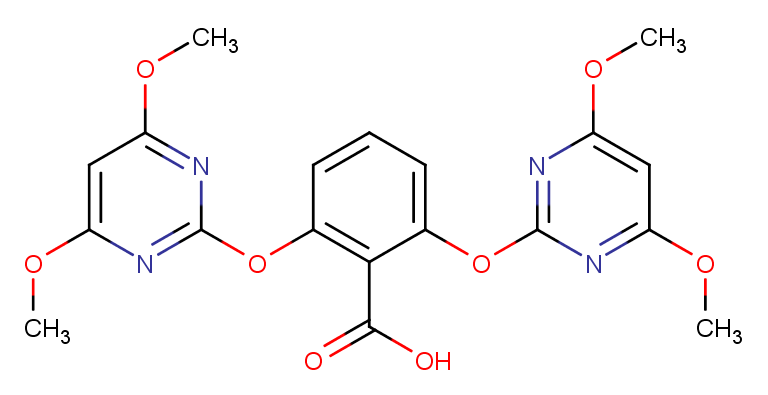

### 126801-58-9.png

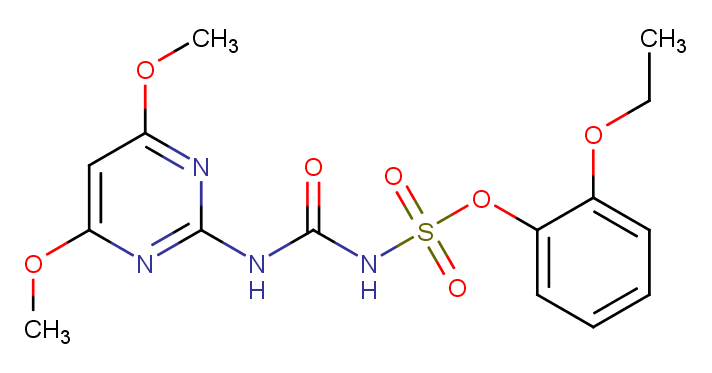

### 128639-02-1.png

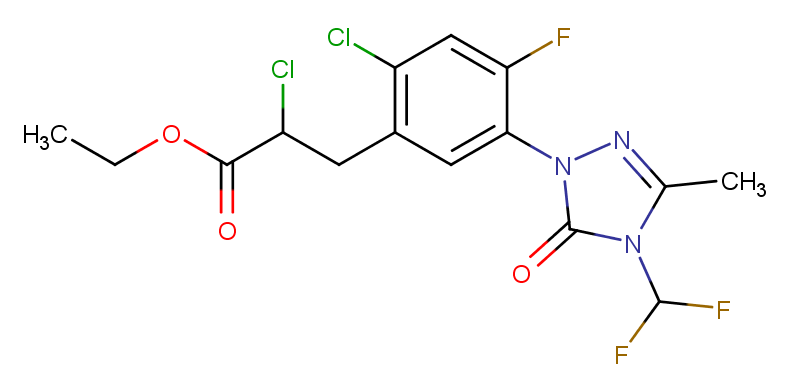

### 129630-19-9.png

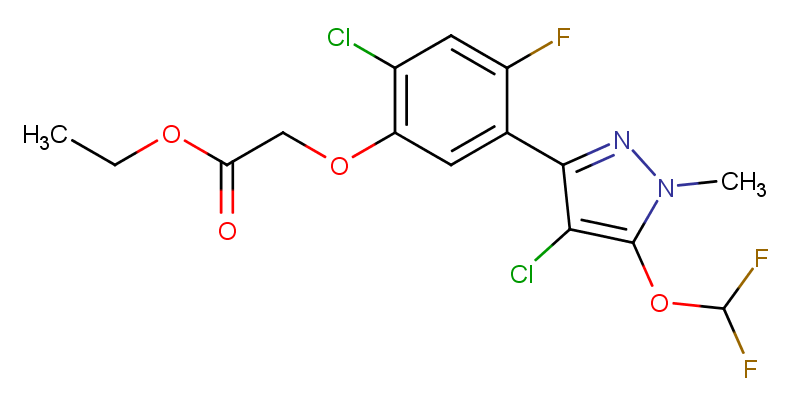

### 129909-90-6.png

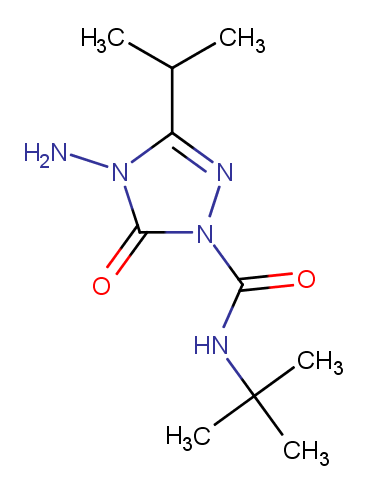

### 131086-42-5.png

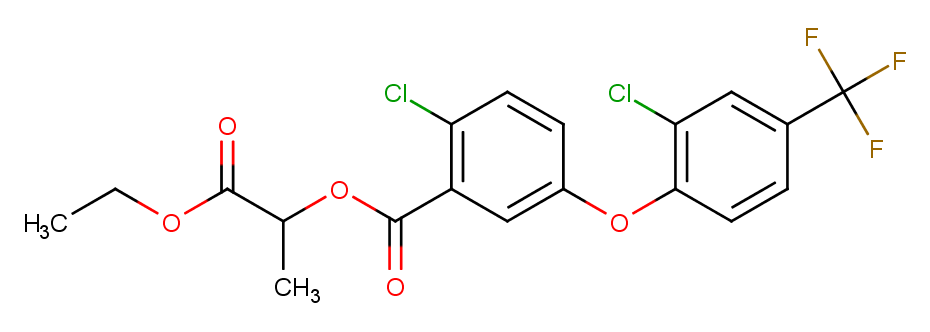

### 131475-57-5.png

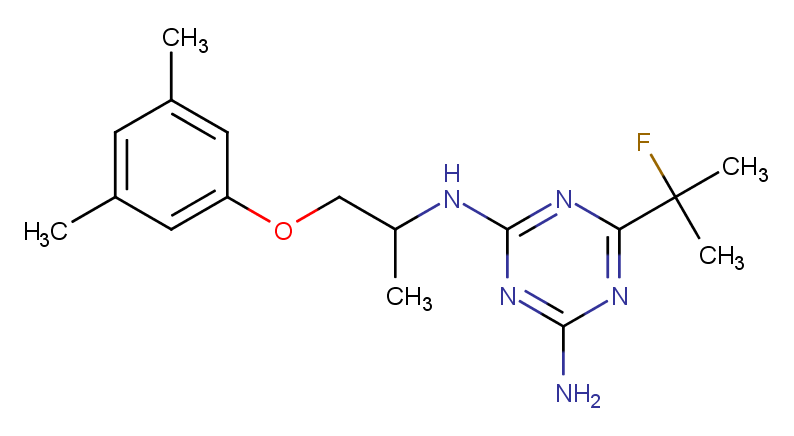

### 133220-30-1.png

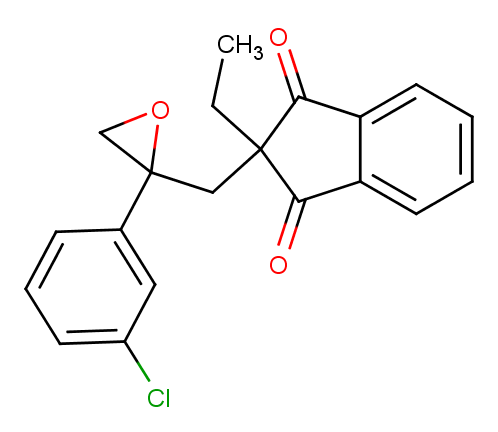

### 134019-30-0.png

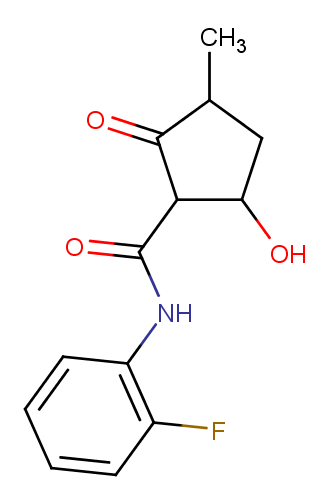

### 134605-64-4.png

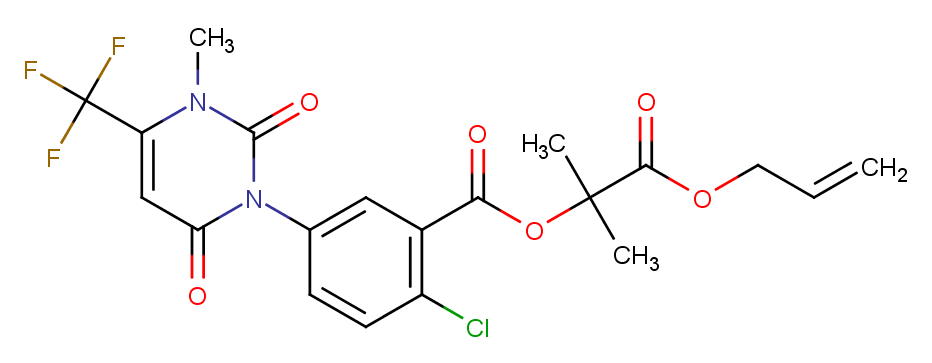

### 135186-78-6.png

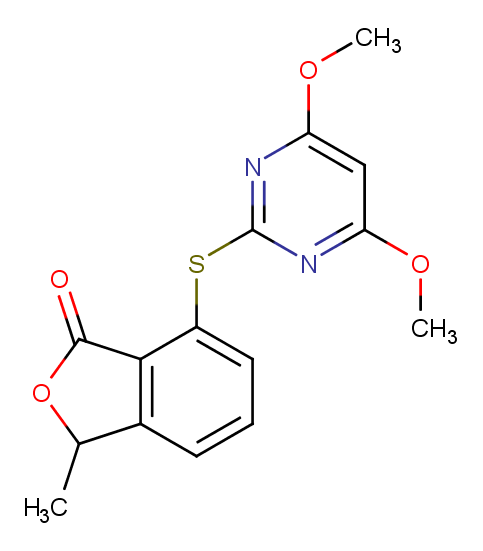

### 135990-29-3.png

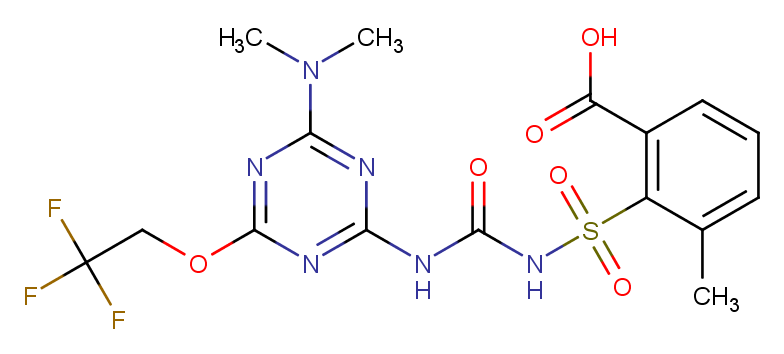

### 136849-15-5.png

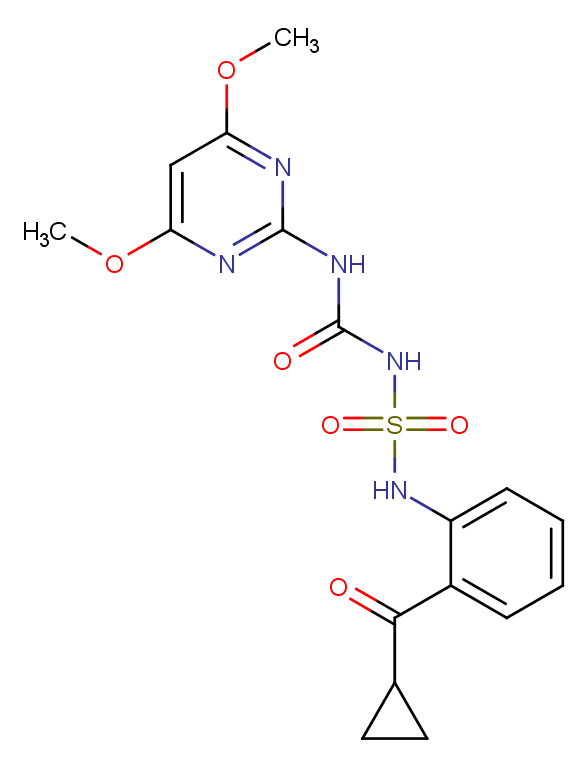

### 137641-05-5.png

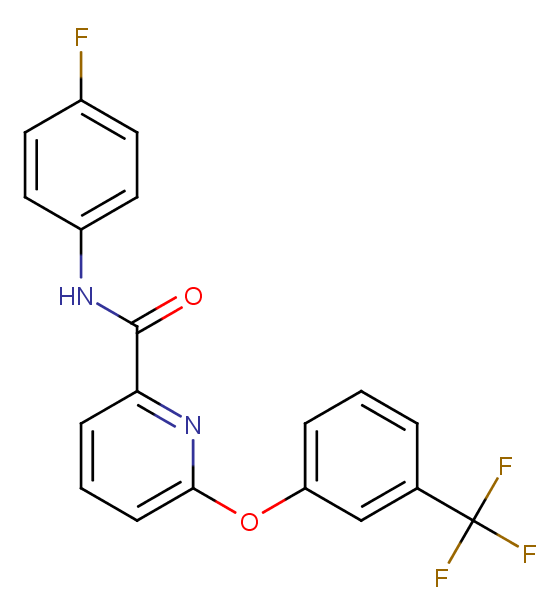

### 138164-12-2.png

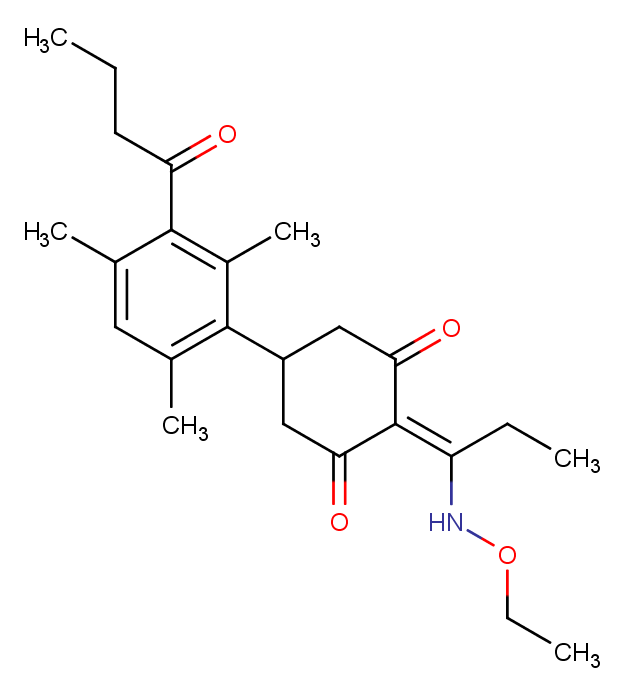

### 139001-49-3.png

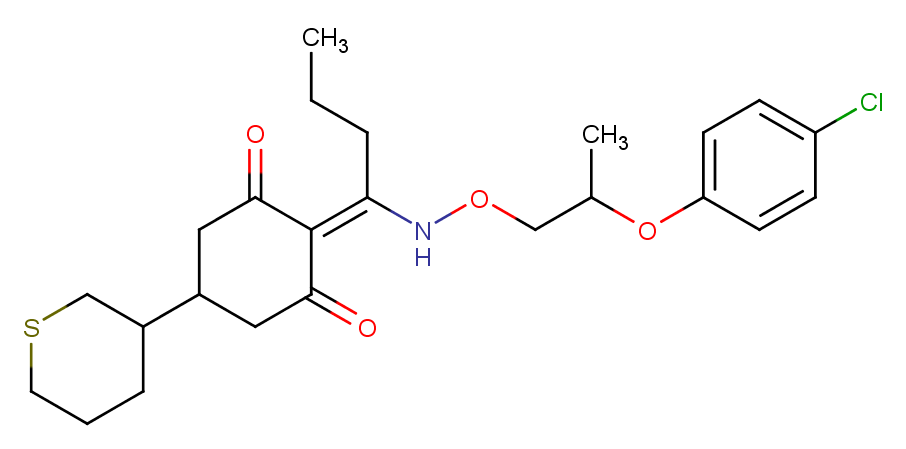

### 139528-85-1.png

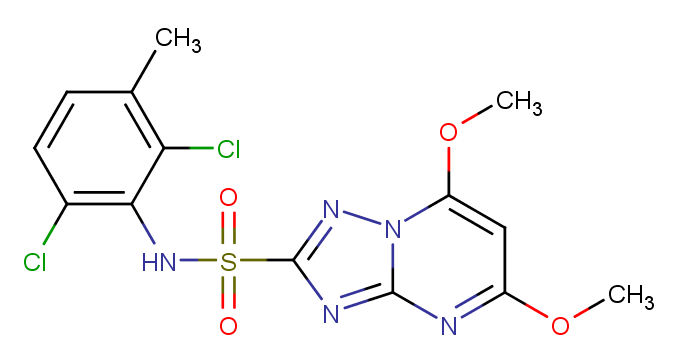

### 141112-06-3.png

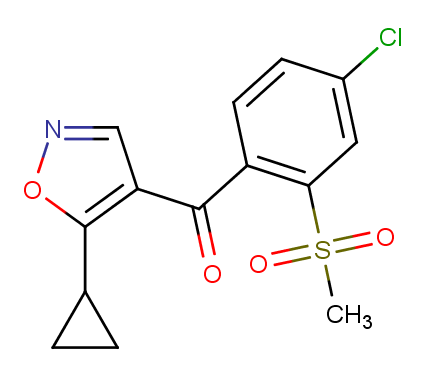

### 141112-29-0.png

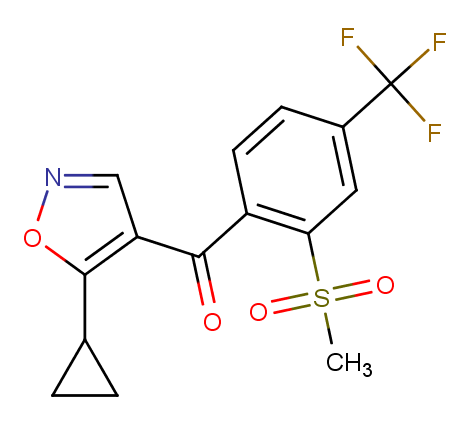

### 141776-32-1.png

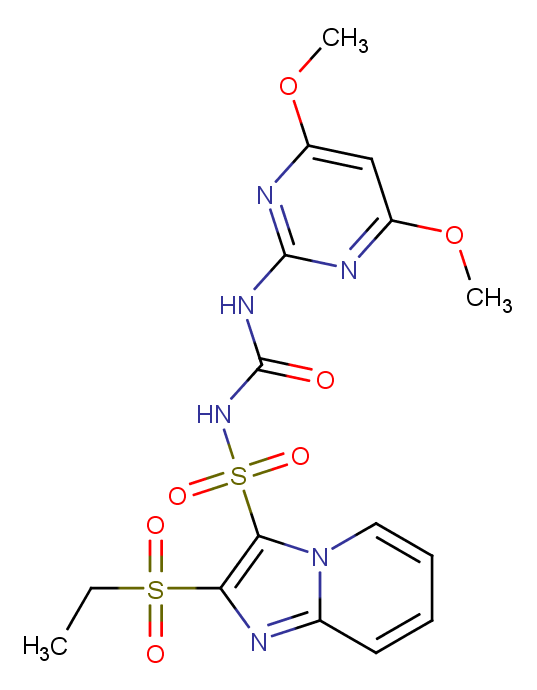

### 142459-58-3.png

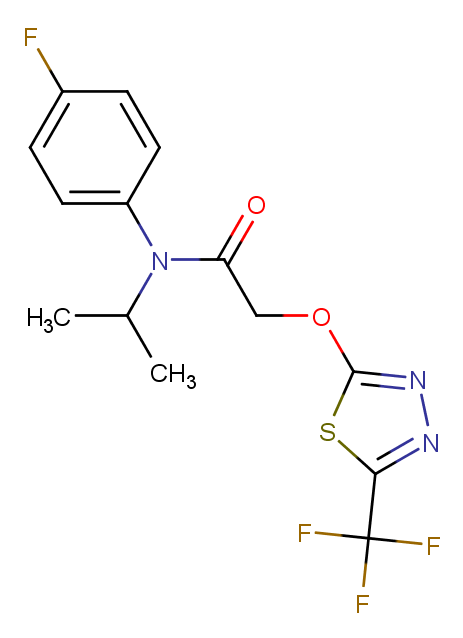

### 142469-14-5.png

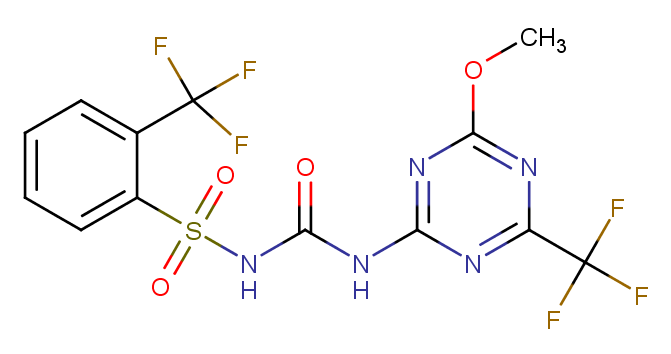

### 142891-20-1.png

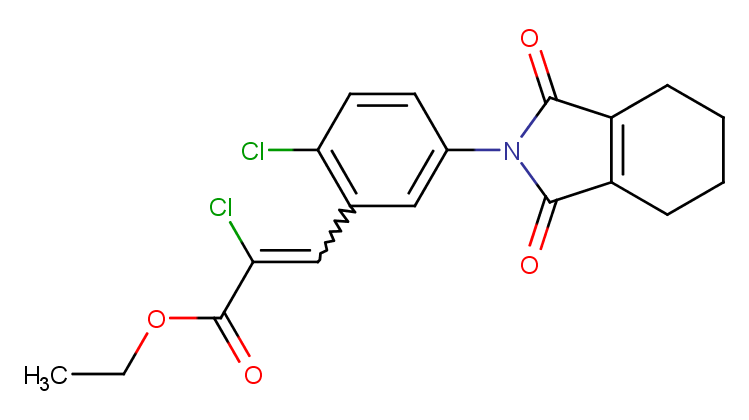

### 144651-06-9.png

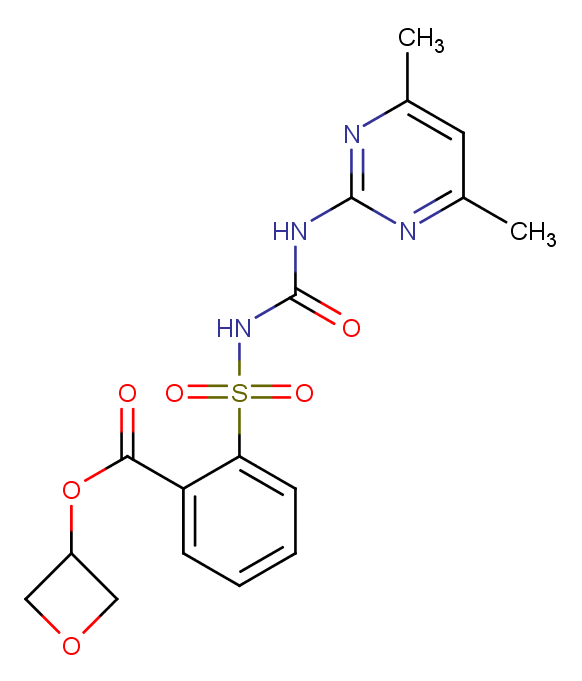

### 144740-53-4.png

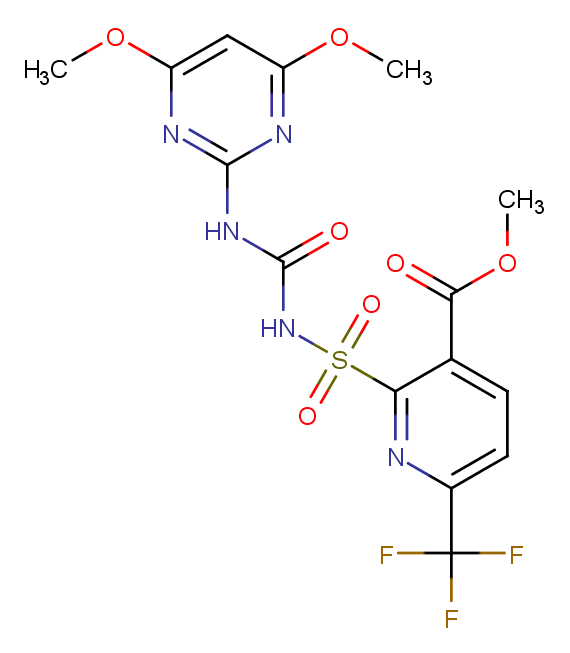

### 145099-21-4.png

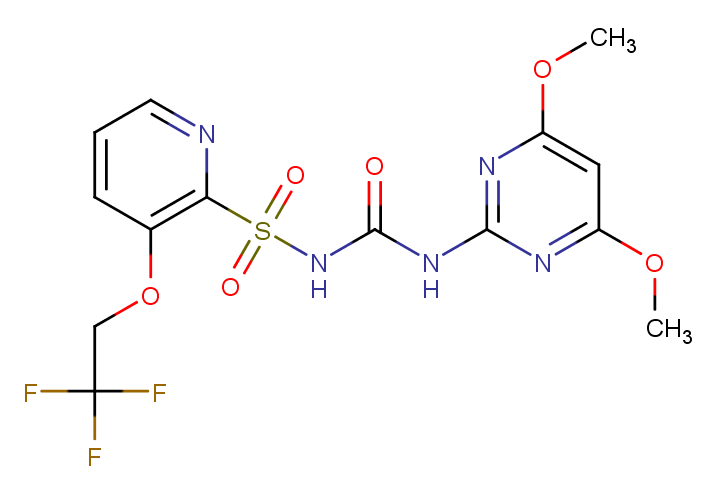

### 145701-21-9.png

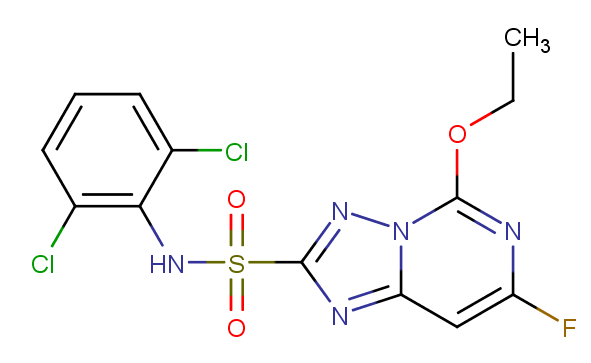

### 145701-23-1.png

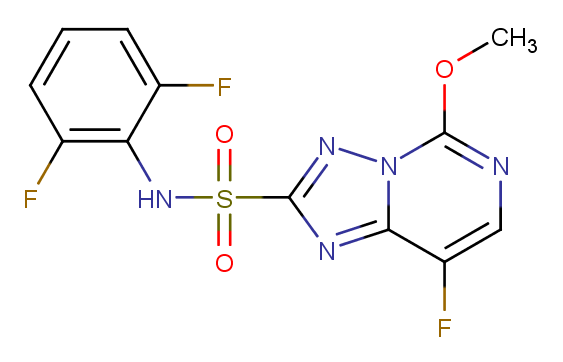

### 147150-35-4.png

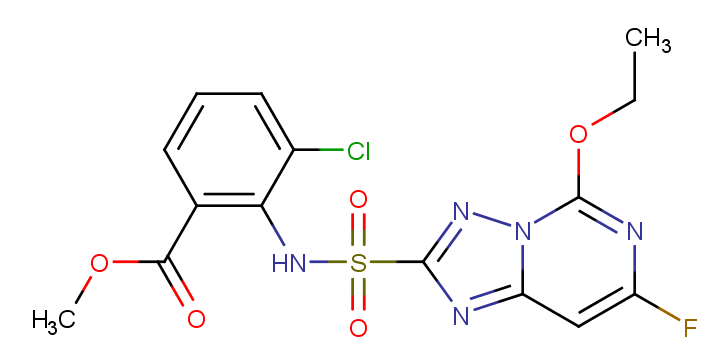

### 147411-69-6.png

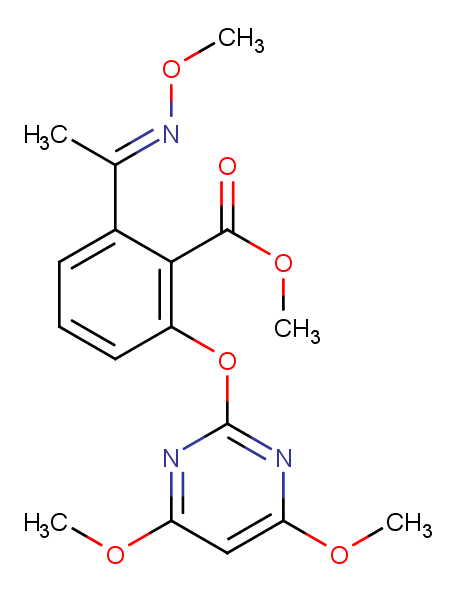

### 149979-41-9.png

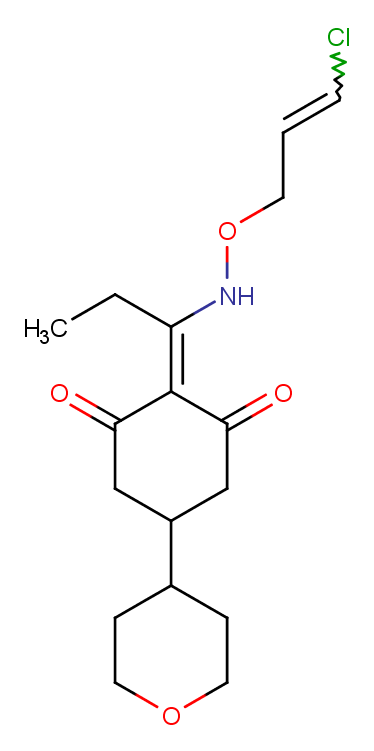

### 150114-71-9.png

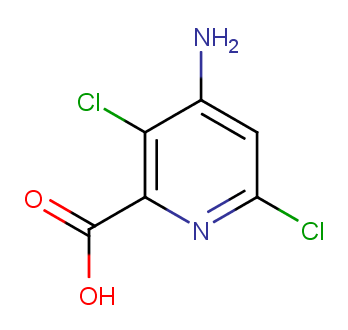

### 153197-14-9.png

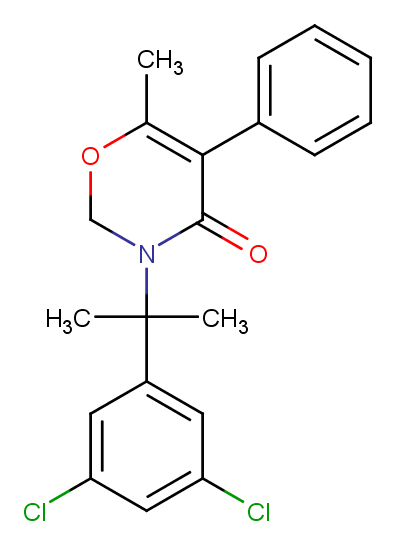

### 155860-63-2.png

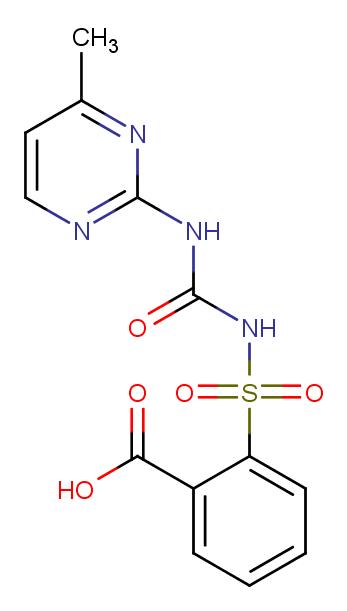

### 1101132-67-5.png

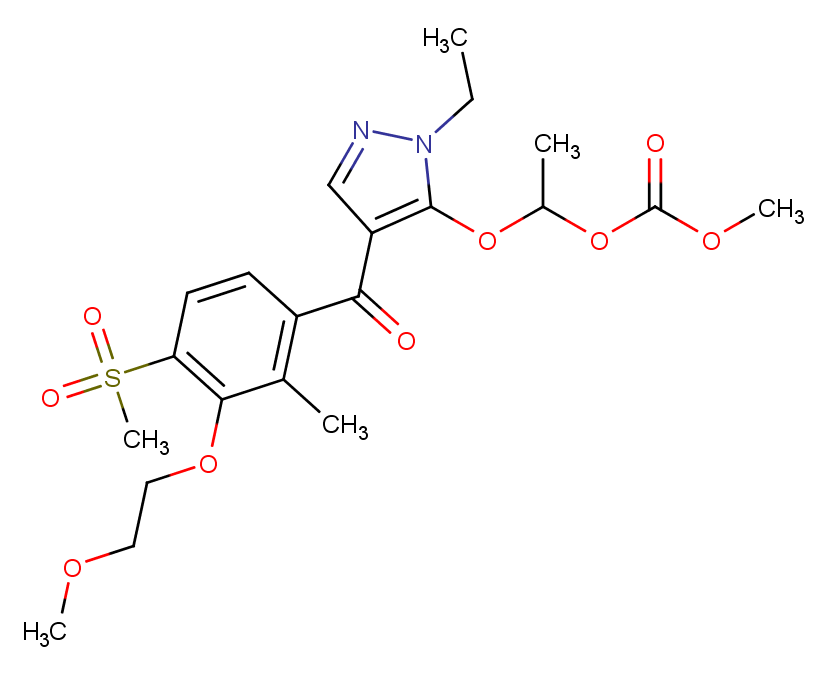

### 1144097-22-2.png

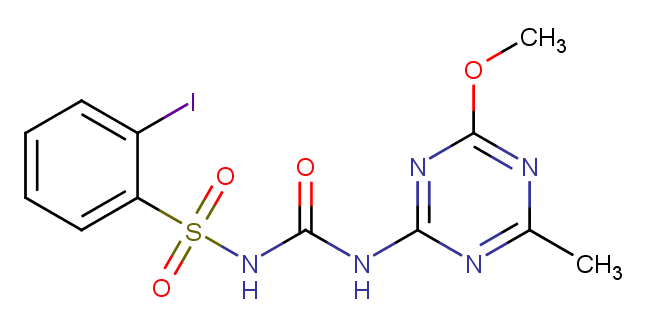

### 1220411-29-9.png

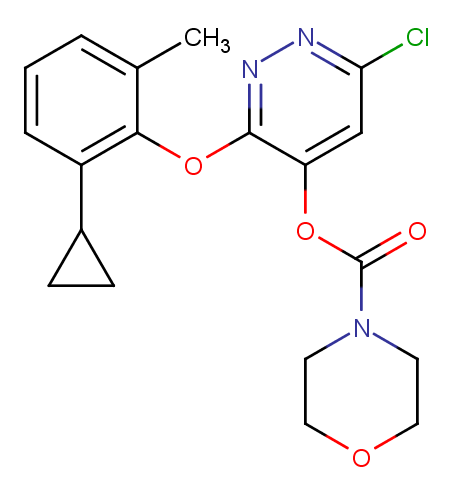

### 1258836-72.png

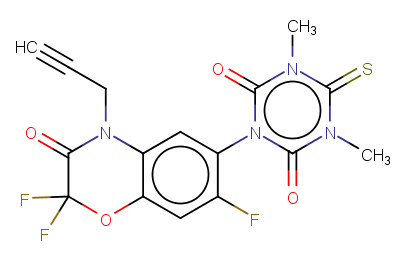

### 1342891-70-6.png

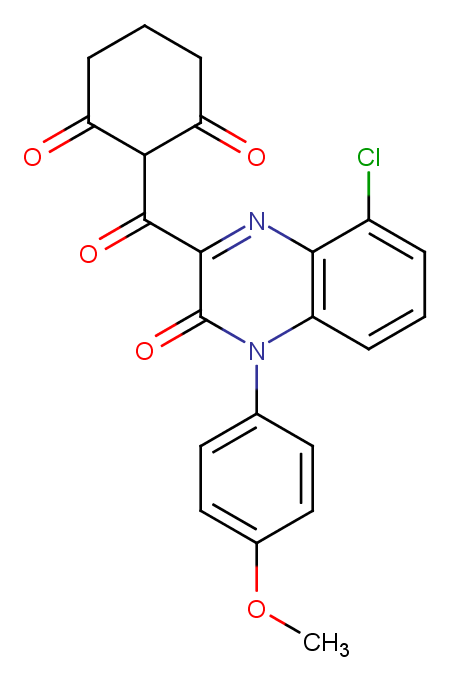
