## Supplementary figures and images for "Refining physico-chemical rules for herbicides using an antimalarial library"

### MMV000052.png

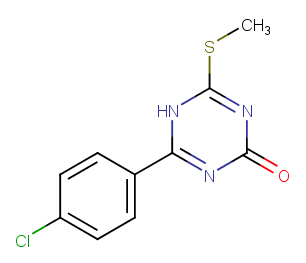

### MMV000187.png

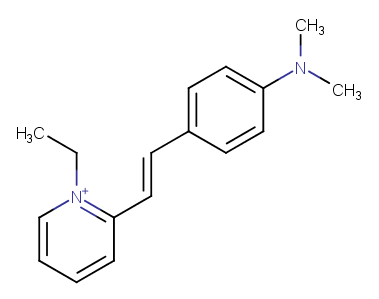

### MMV007374.png

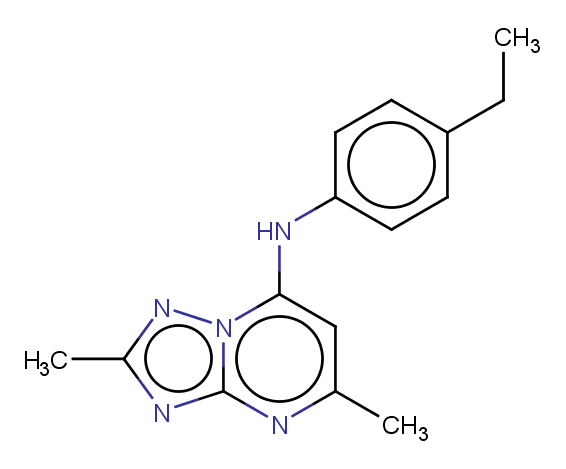

### MMV007384.png

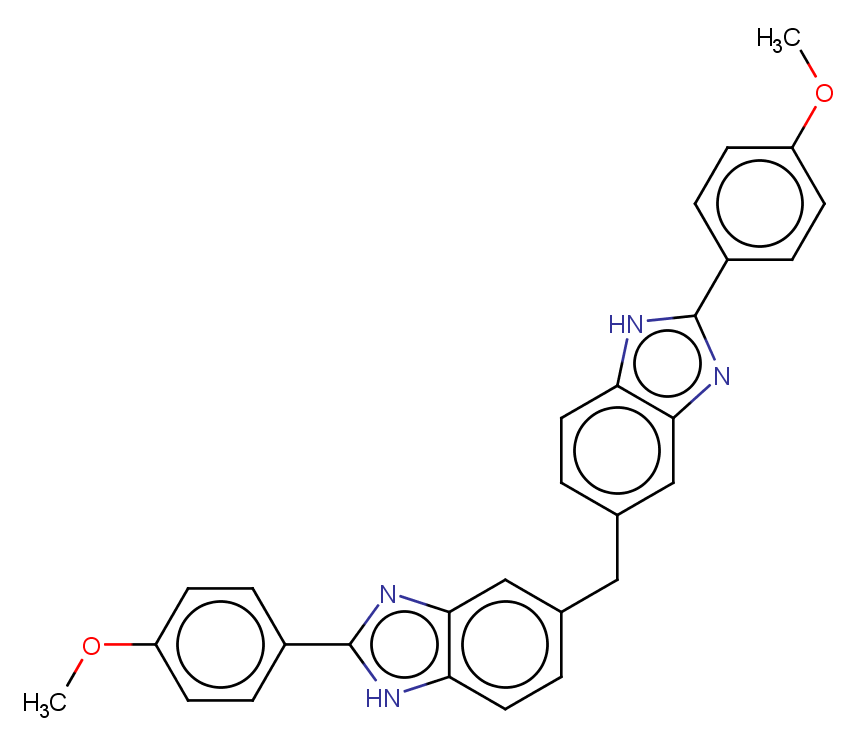

### MMV007396.png

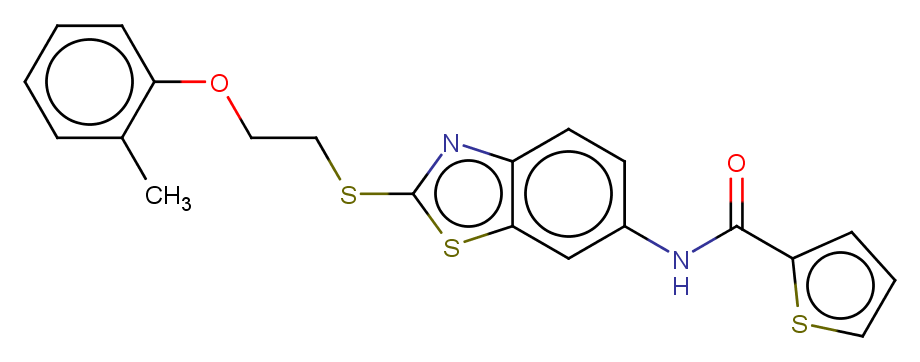

### MMV007430.png

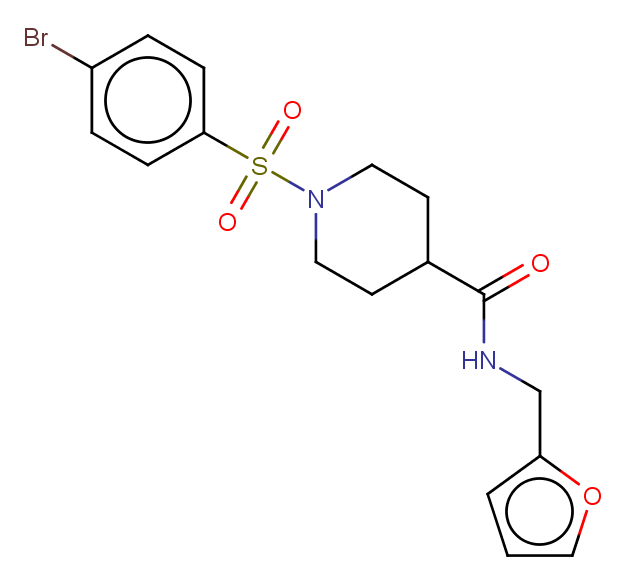

### MMV007474.png

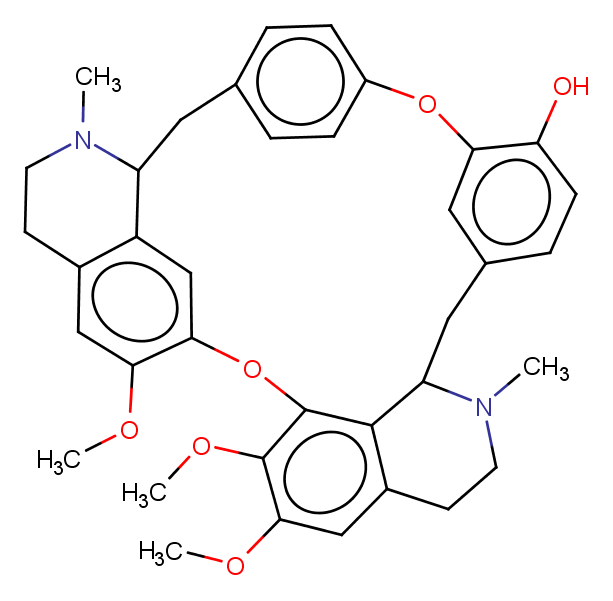

### MMV007557.png

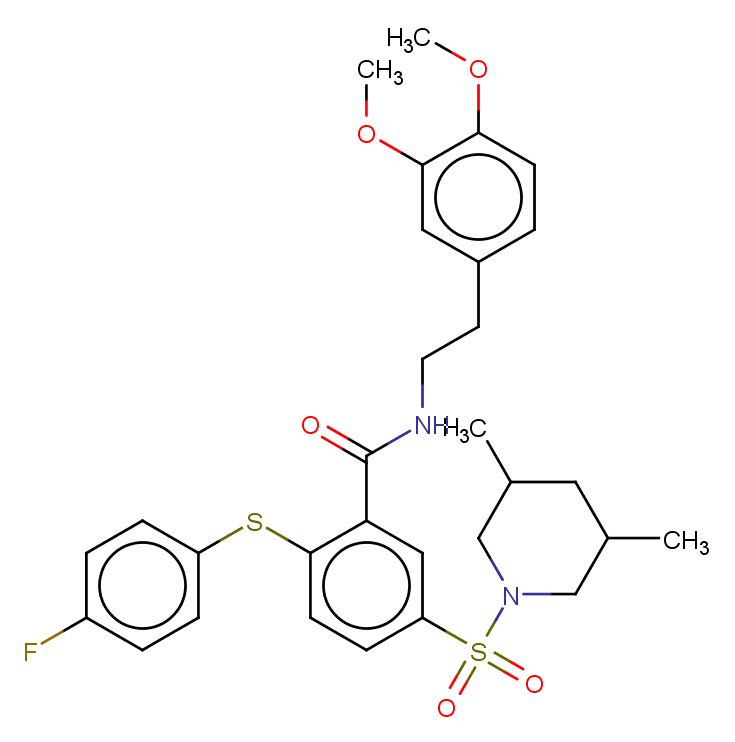

### MMV007564.png

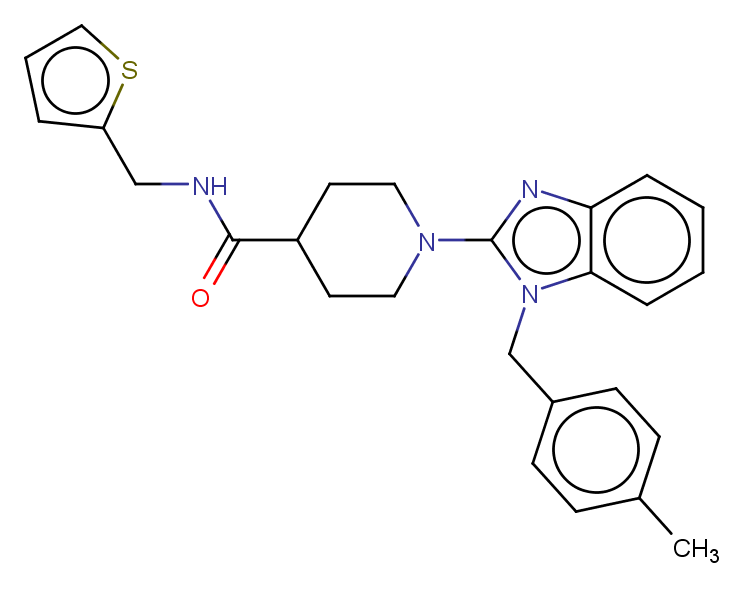

### MMV007571.png

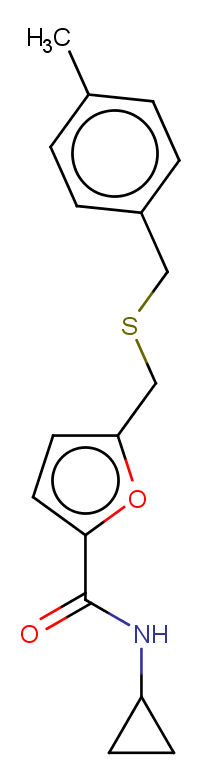

### MMV007574.png

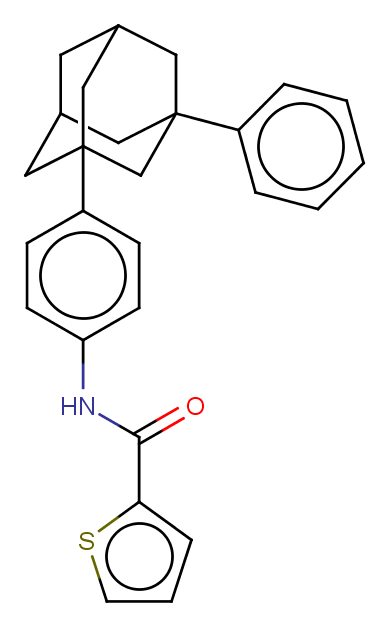

### MMV007577.png

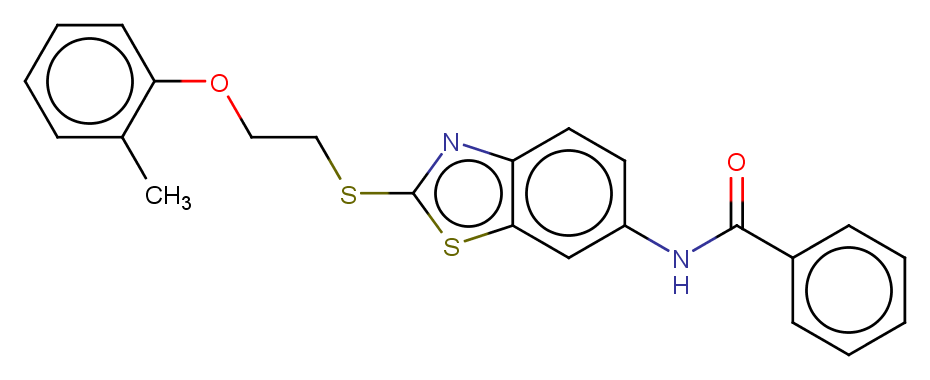

### MMV007591.png

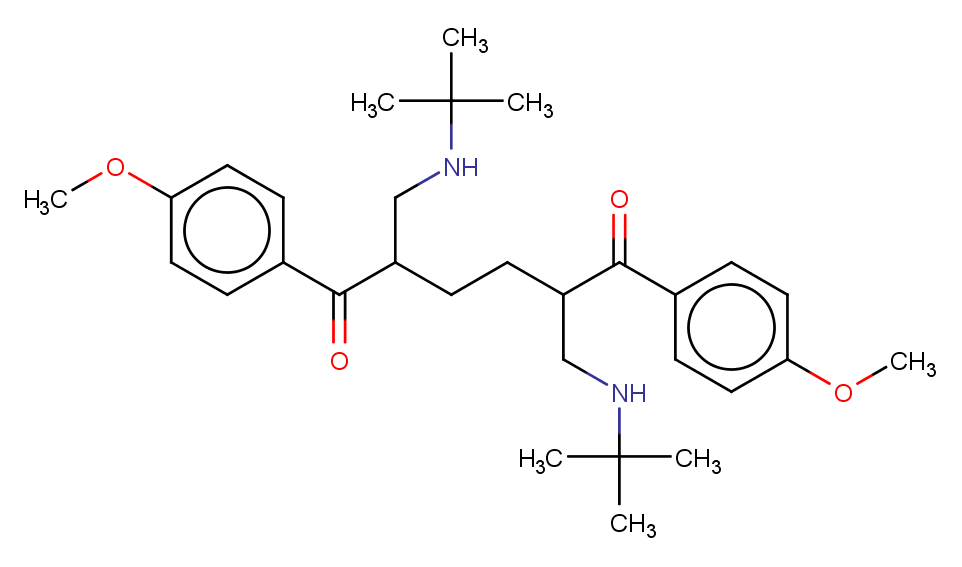

### MMV007617.png

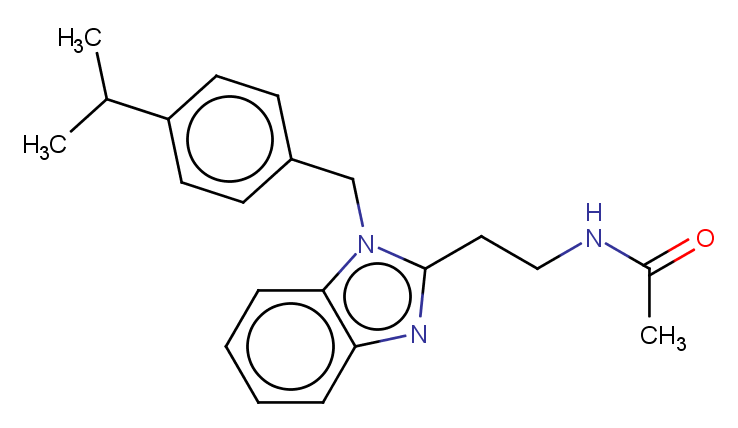

### MMV007654.png

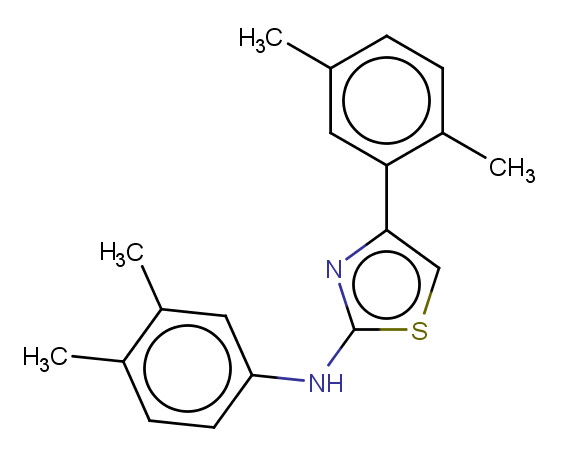

### MMV007686.png

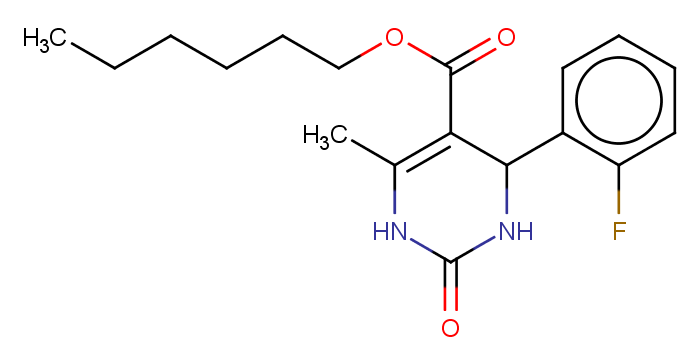

### MMV007695.png

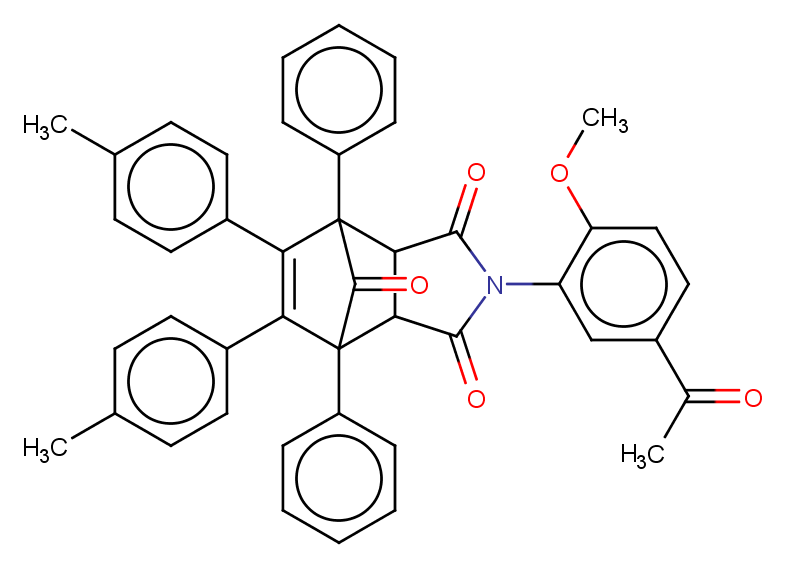

### MMV007764.png

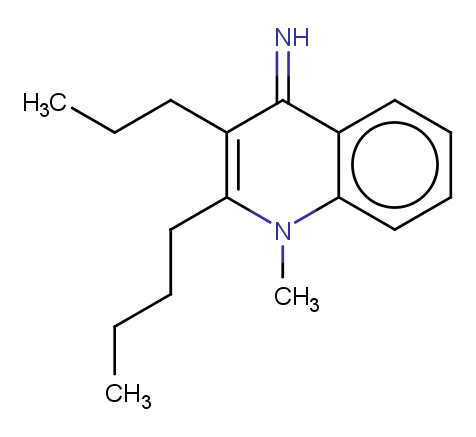

### MMV007791.png

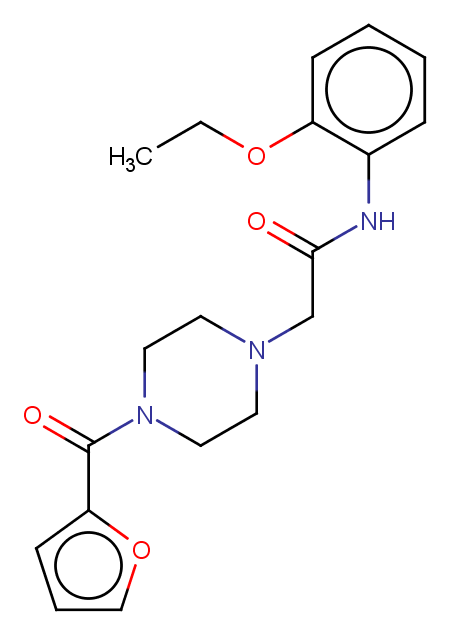

### MMV007808.png

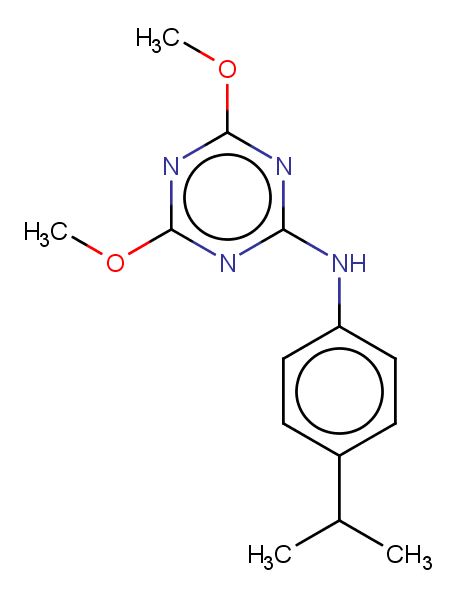

### MMV007839.png

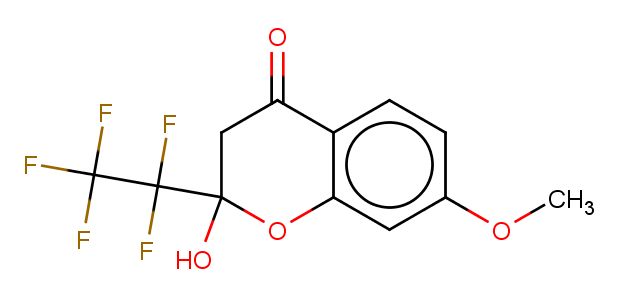

### MMV007875.png

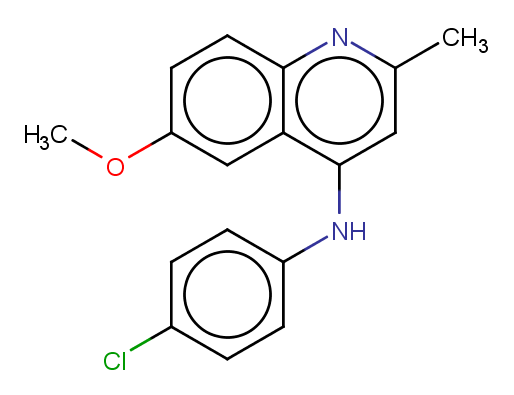

### MMV007881.png

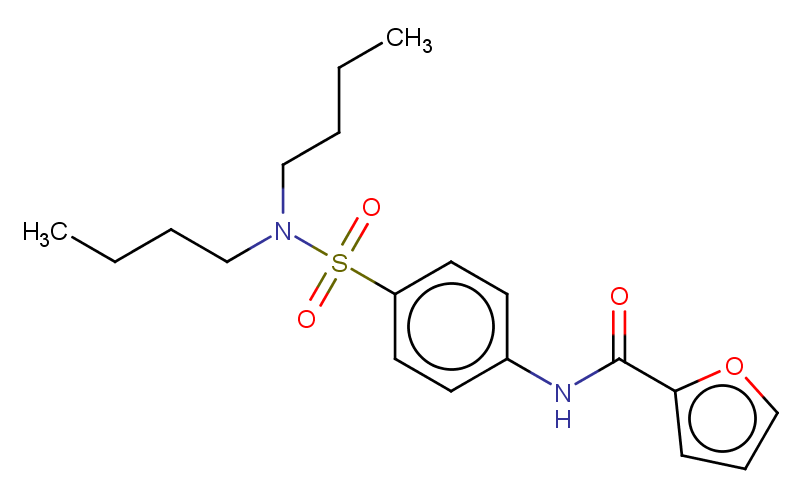

### MMV007906.png

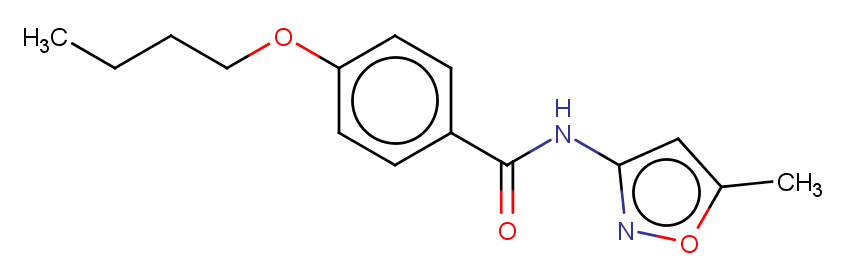

### MMV007907.png

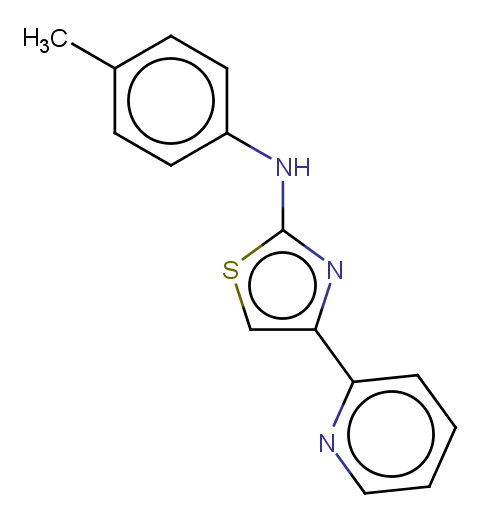

### MMV007977.png

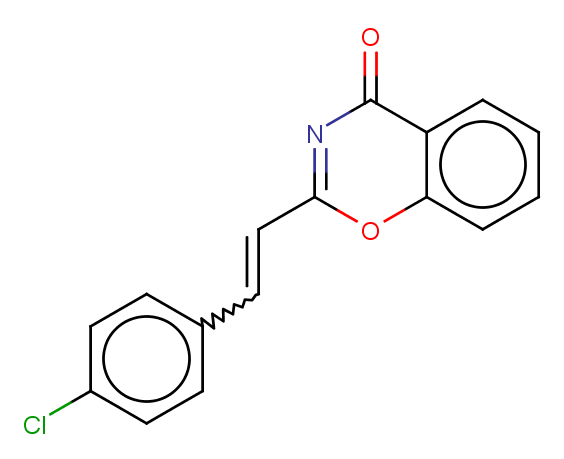

### MMV007978.png

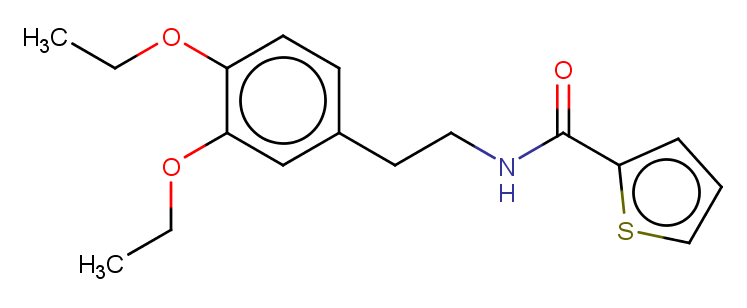

### MMV008127.png

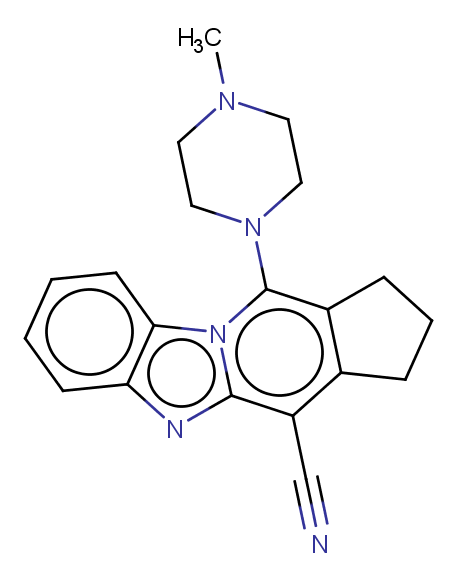

### MMV008138.png

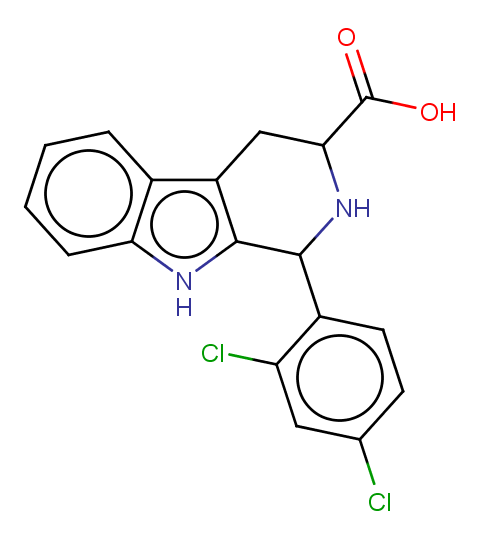

### MMV008149.png

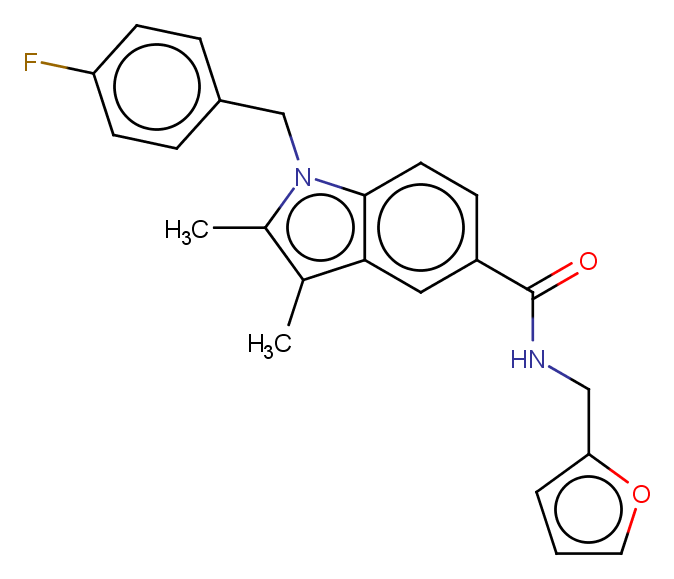

### MMV008160.png

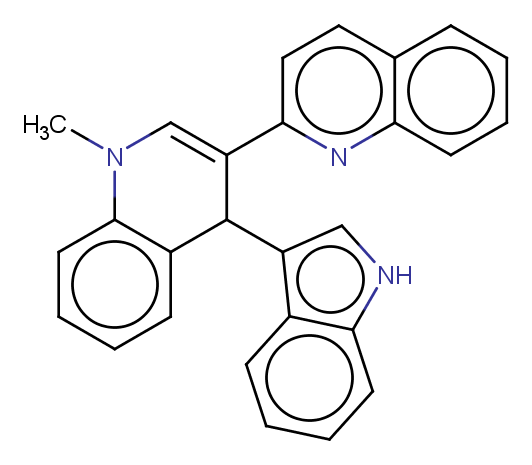

### MMV008173.png

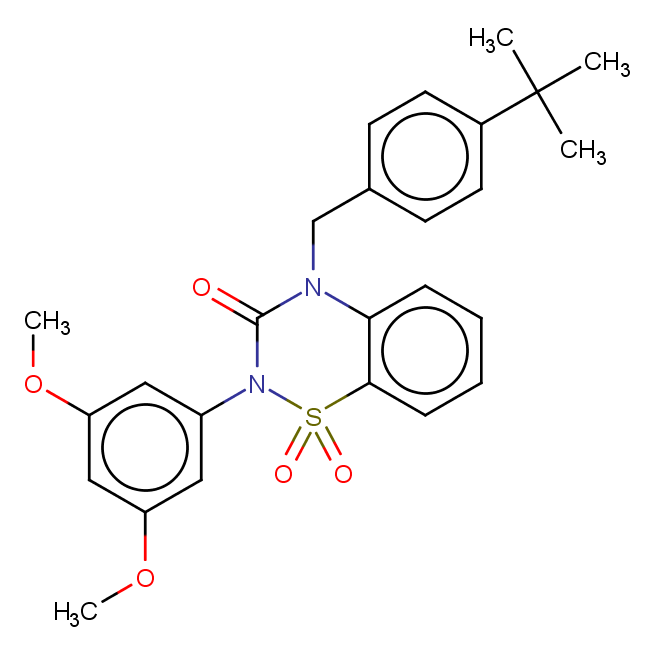

### MMV008212.png

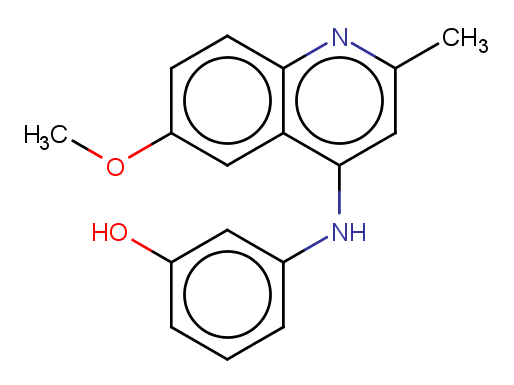

### MMV008270.png

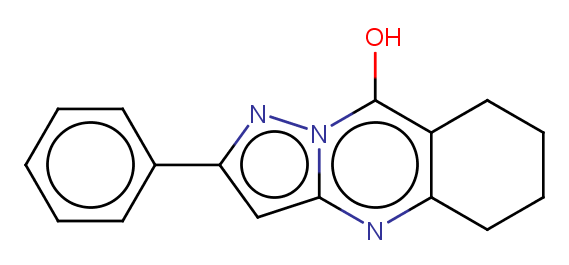

### MMV008294.png

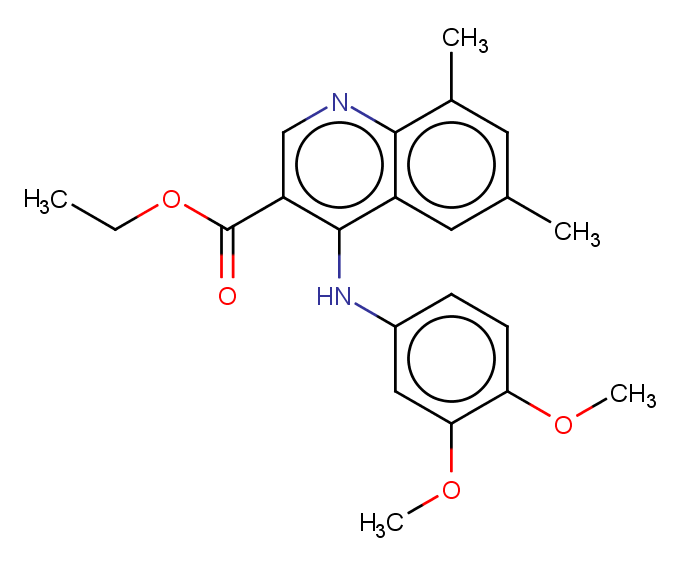

### MMV008416.png

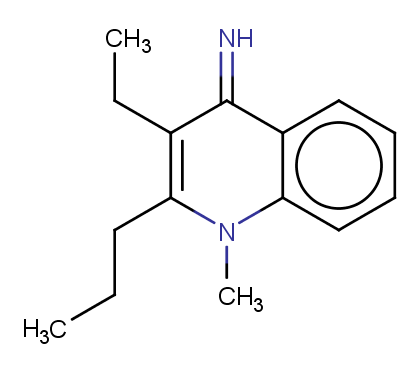

### MMV008455.png

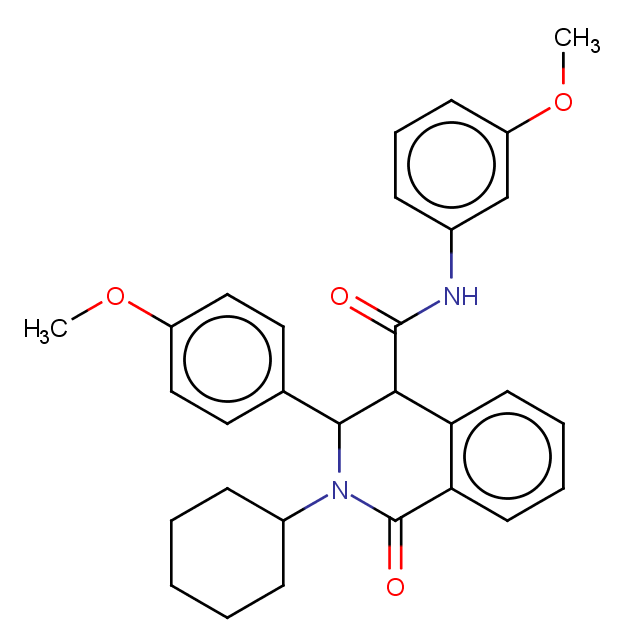

### MMV008474.png

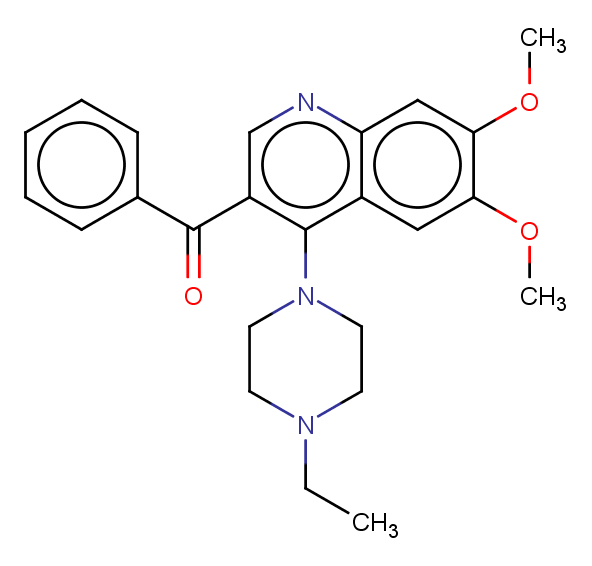

### MMV008829.png

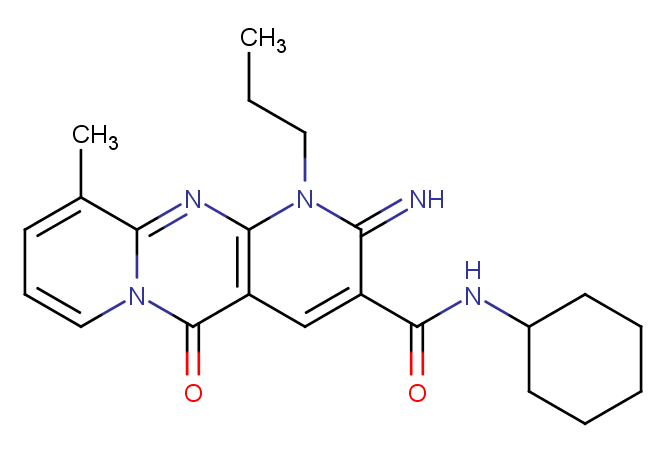

### MMV008956.png

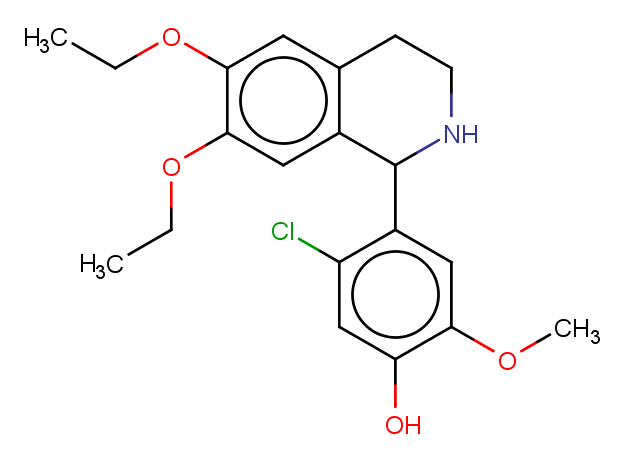

### MMV009015.png

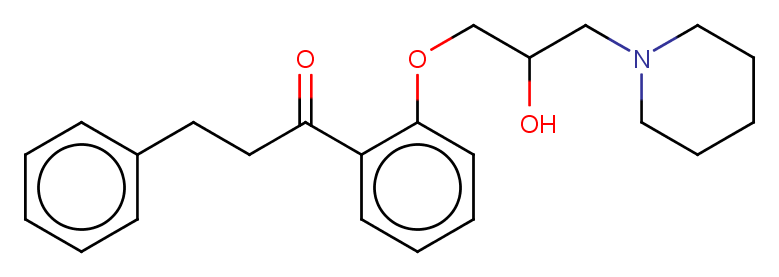

### MMV009060.png

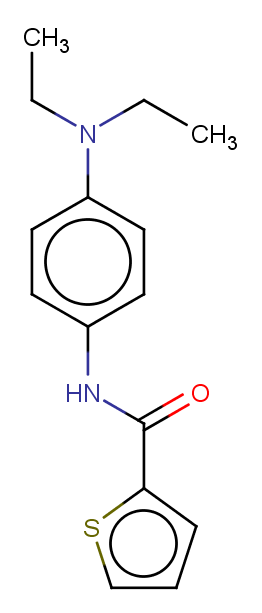

### MMV009063.png

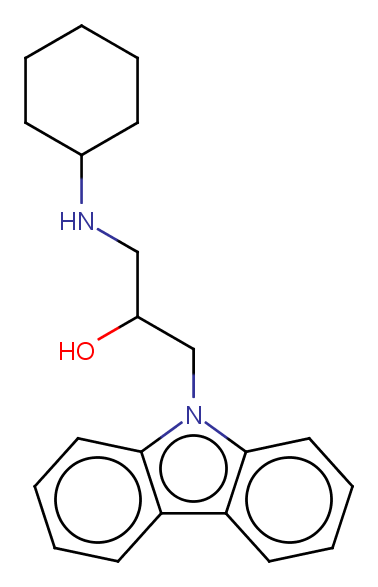

### MMV009085.png

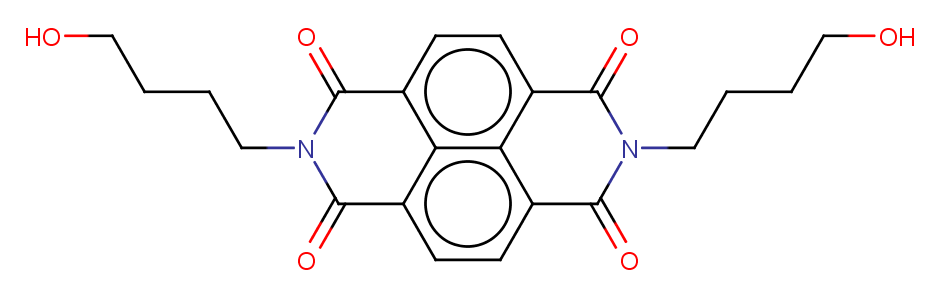

### MMV009108.png

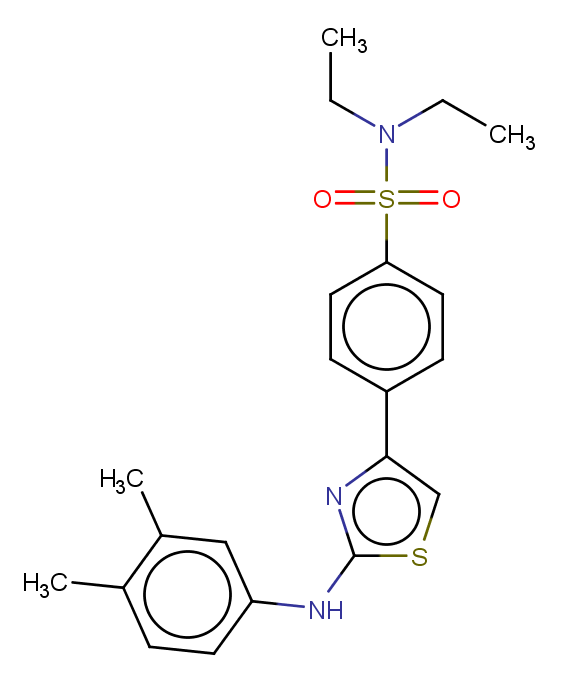

### MMV009127.png

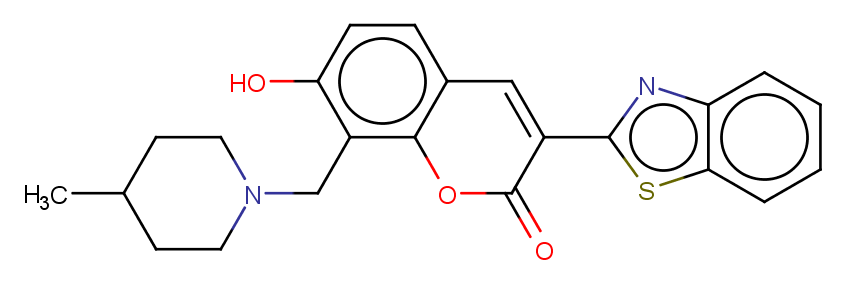

### MMV011099.png

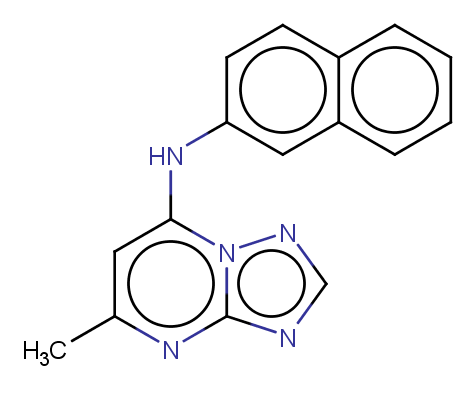

### MMV011256.png

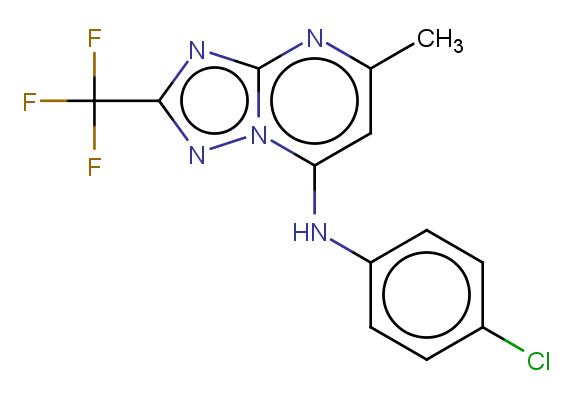

### MMV011259.png

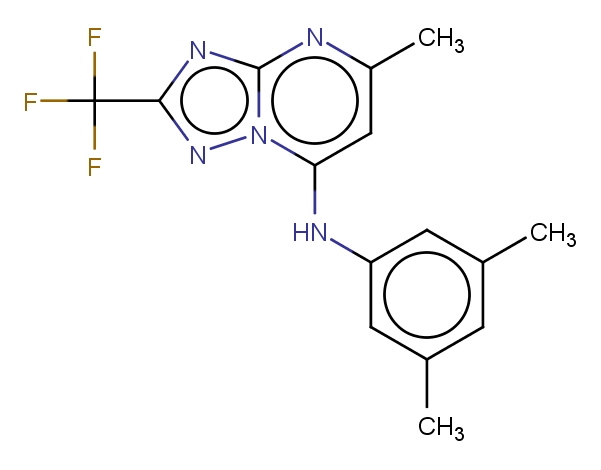

### MMV011436.png

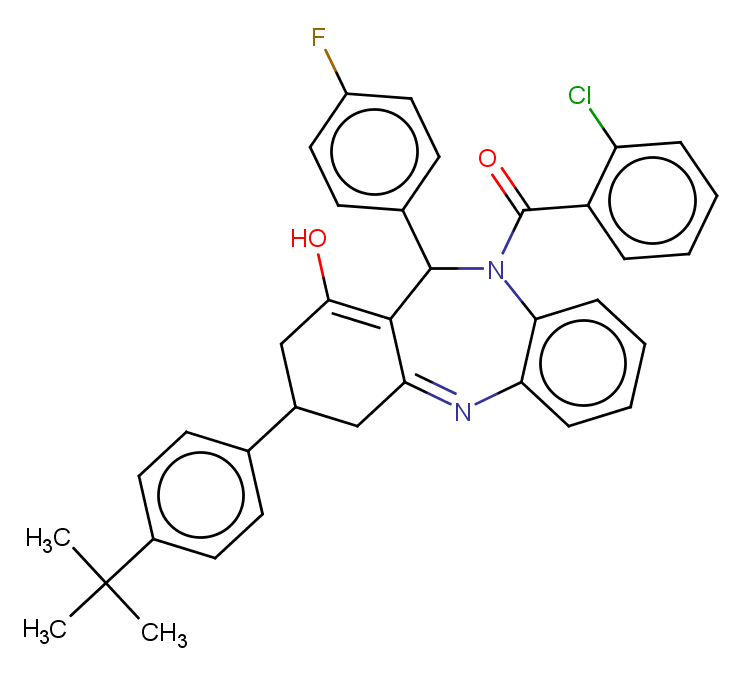

### MMV011438.png

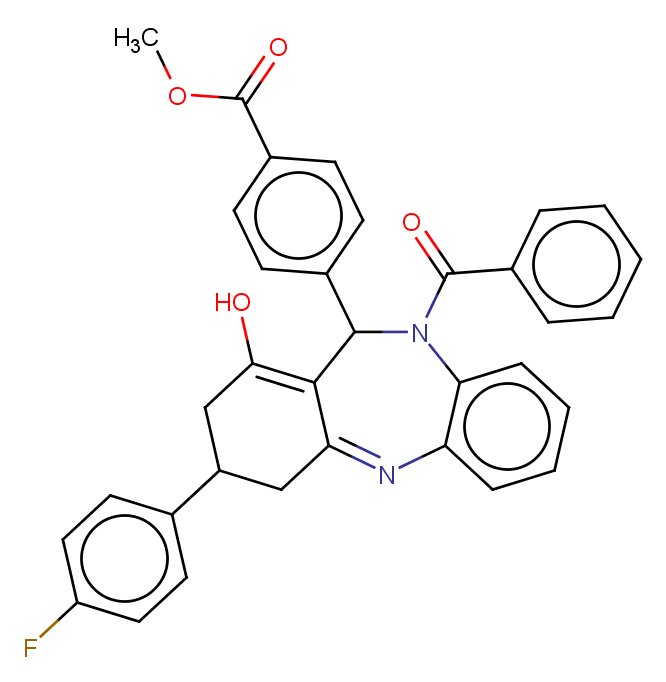

### MMV011522.png

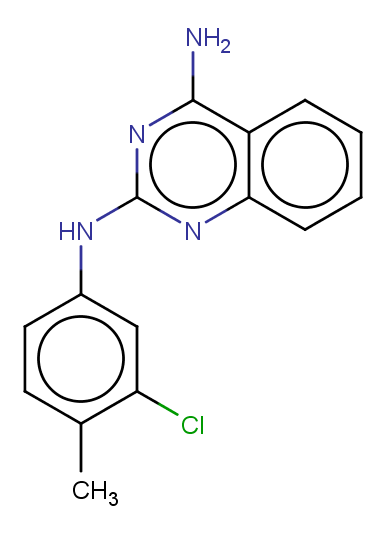

### MMV011567.png

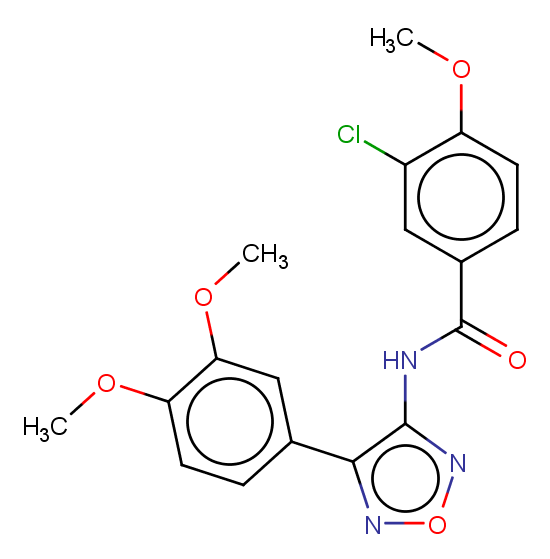

### MMV011576.png

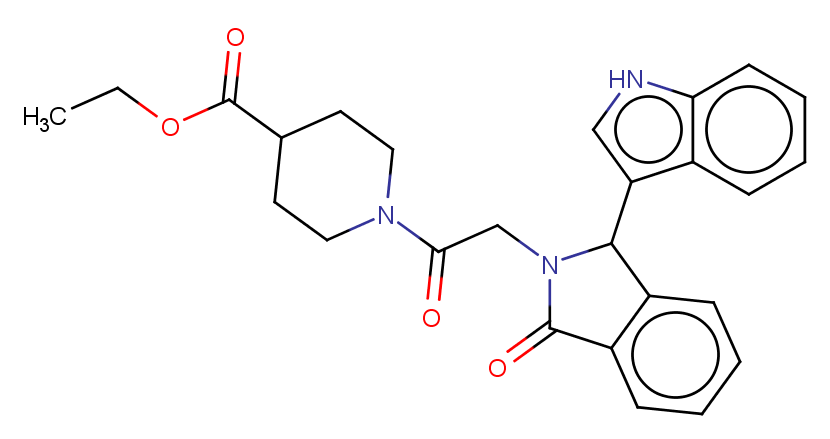

### MMV011795.png

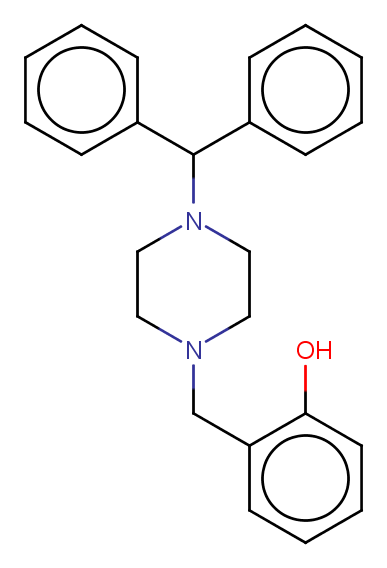

### MMV011832.png

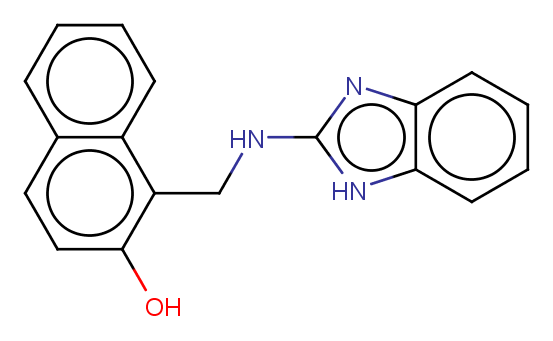

### MMV011895.png

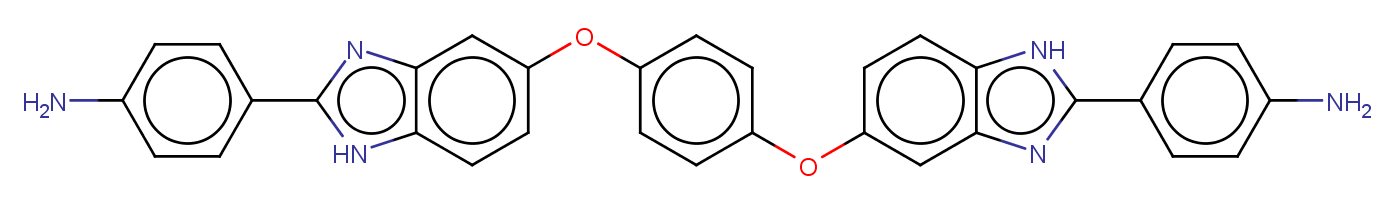

### MMV011944.png

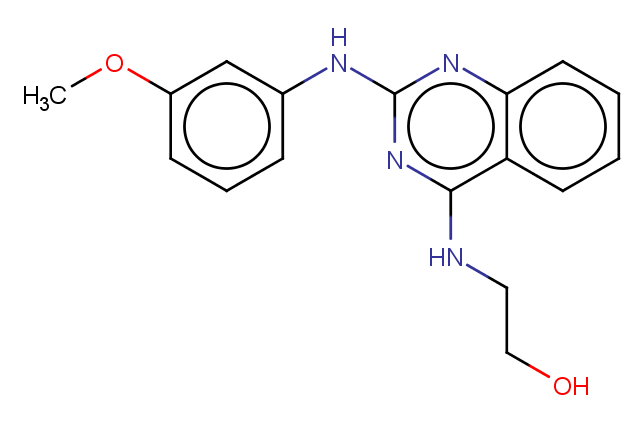

### MMV018984.png

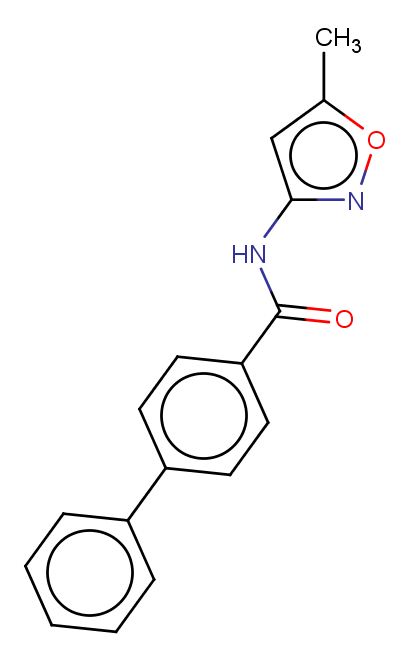

### MMV019017.png

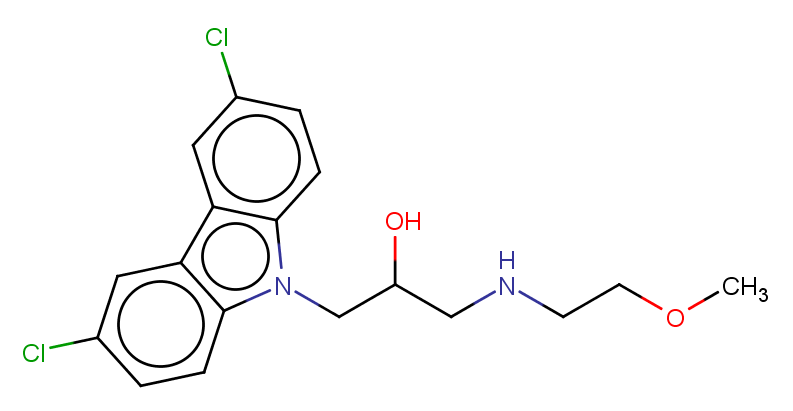

### MMV019064.png

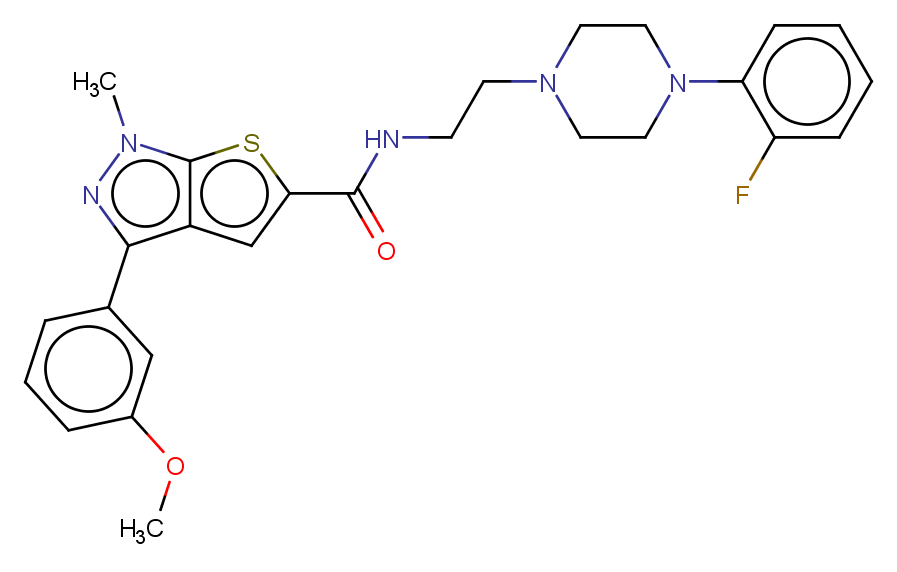

### MMV019066.png

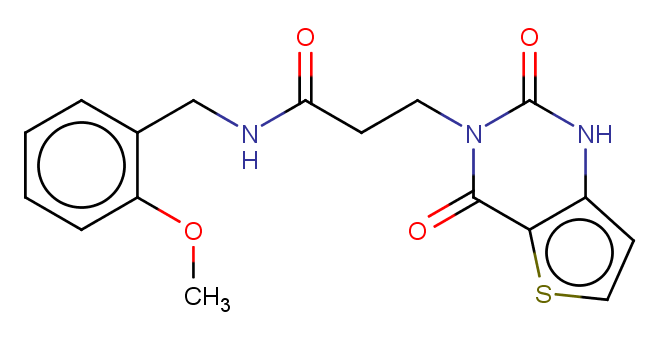

### MMV019074.png

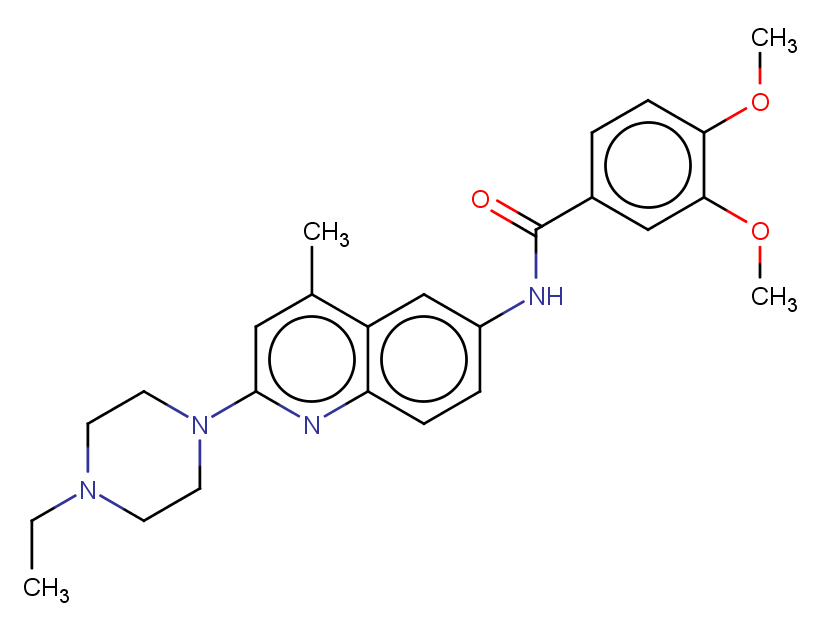

### MMV019110.png

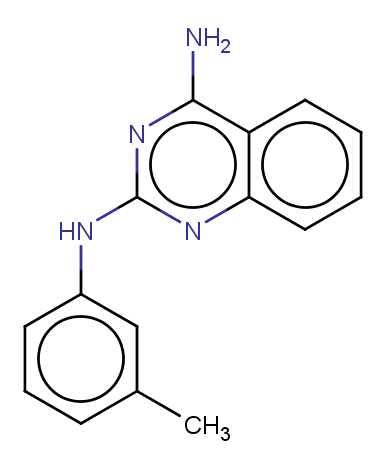

### MMV019124.png

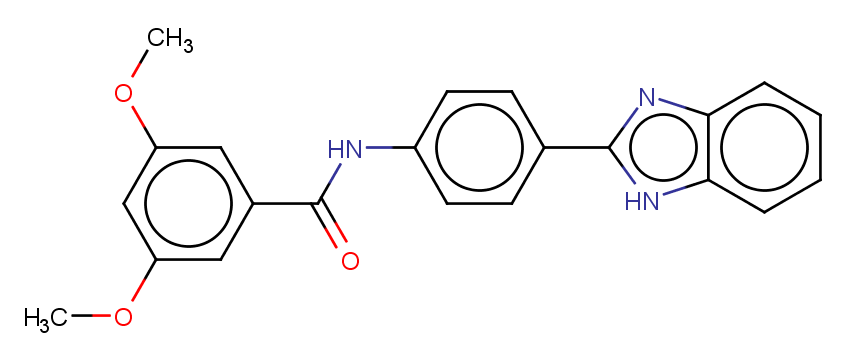

### MMV019127.png

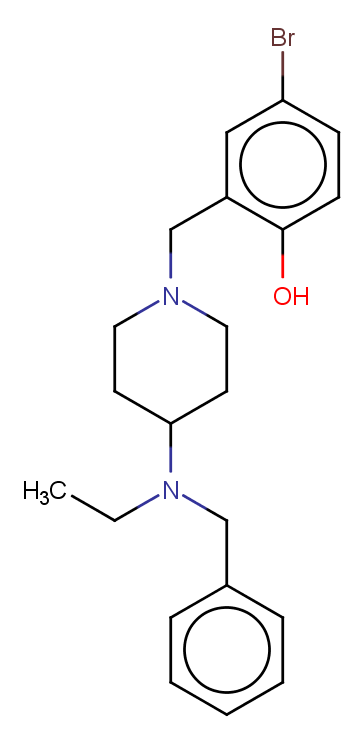

### MMV019199.png

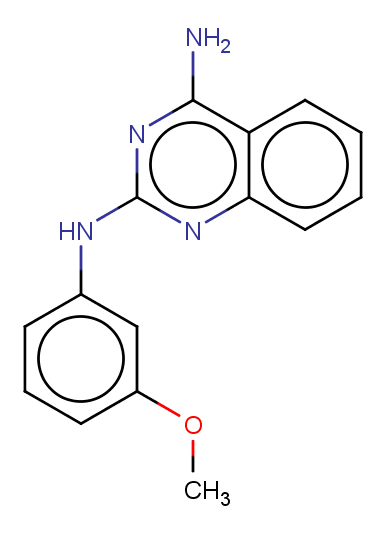

### MMV019202.png

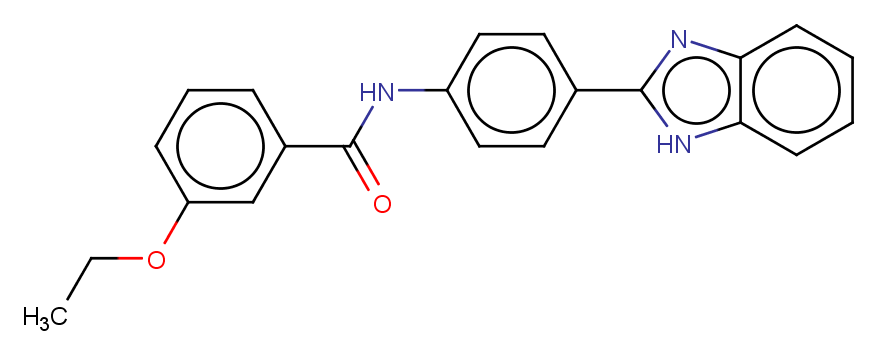

### MMV019241.png

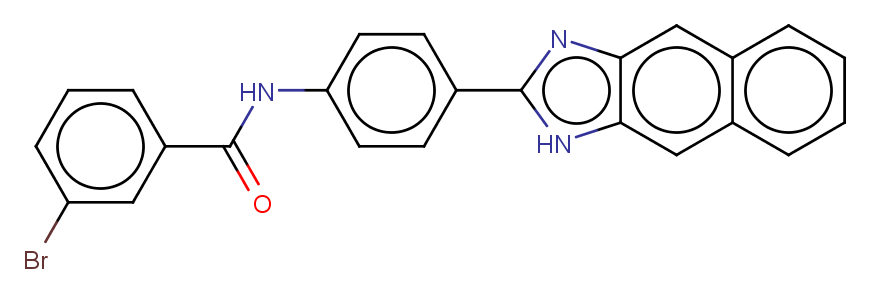

### MMV019258.png

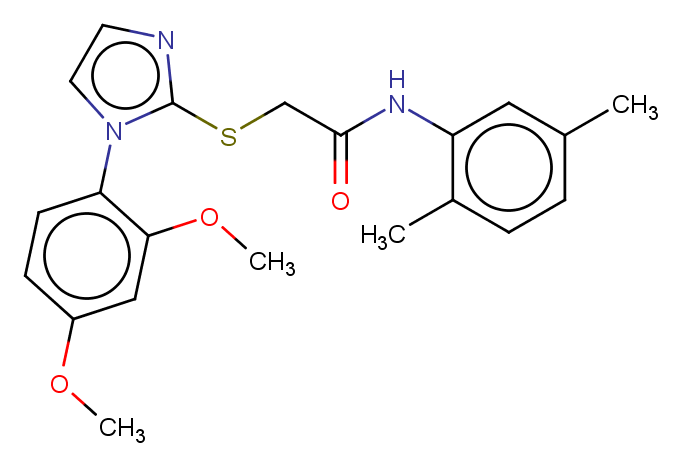

### MMV019266.png

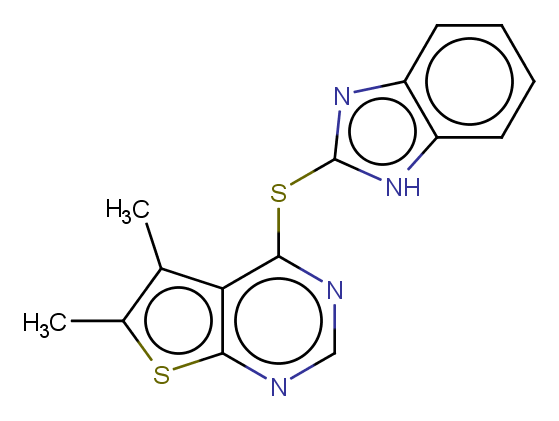

### MMV019313.png

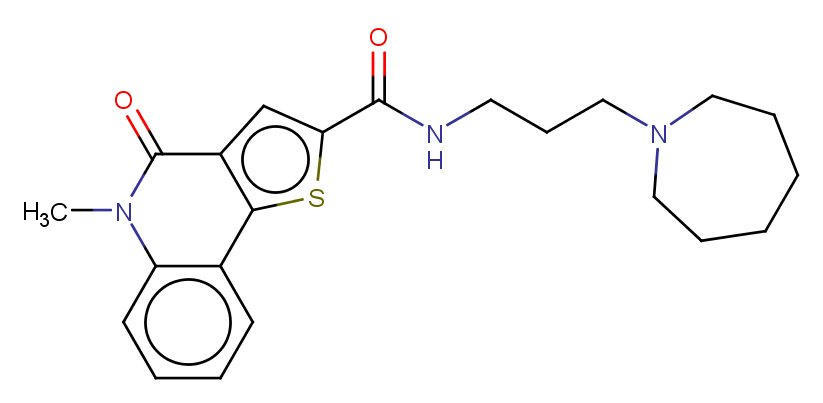

### MMV019406.png

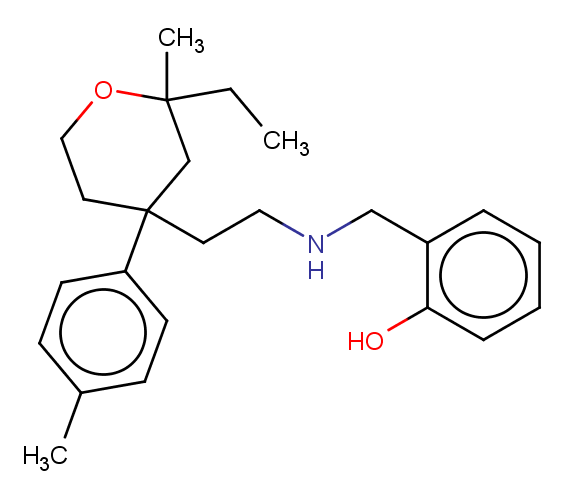

### MMV019555.png

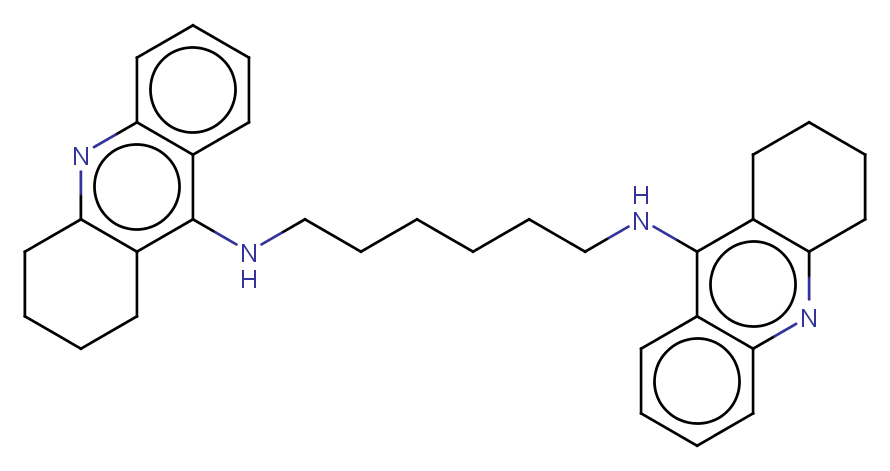

### MMV019662.png

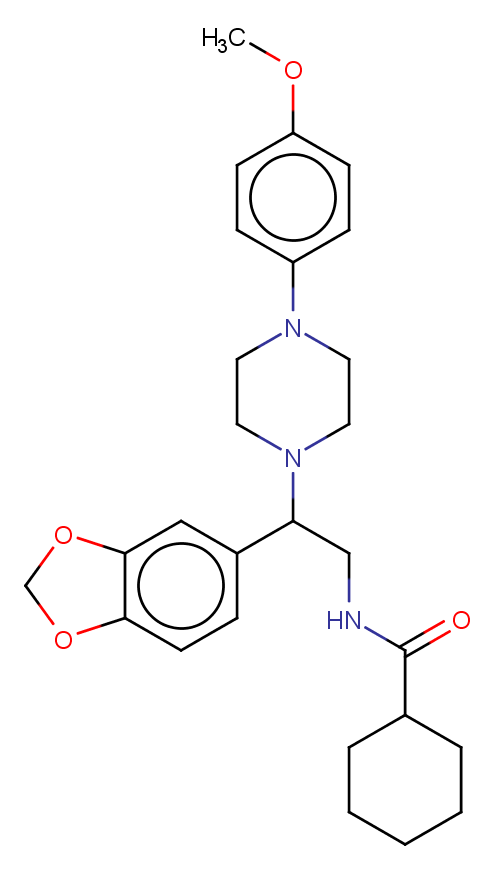

### MMV019670.png

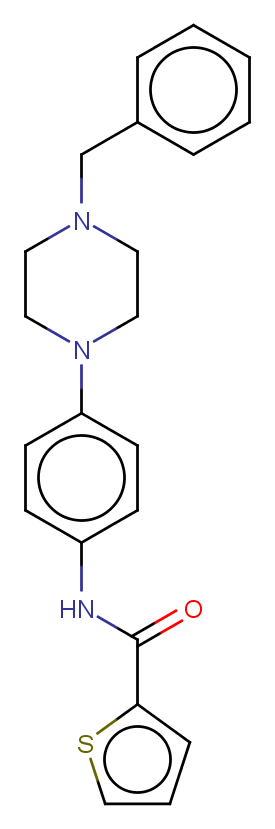

### MMV019690.png

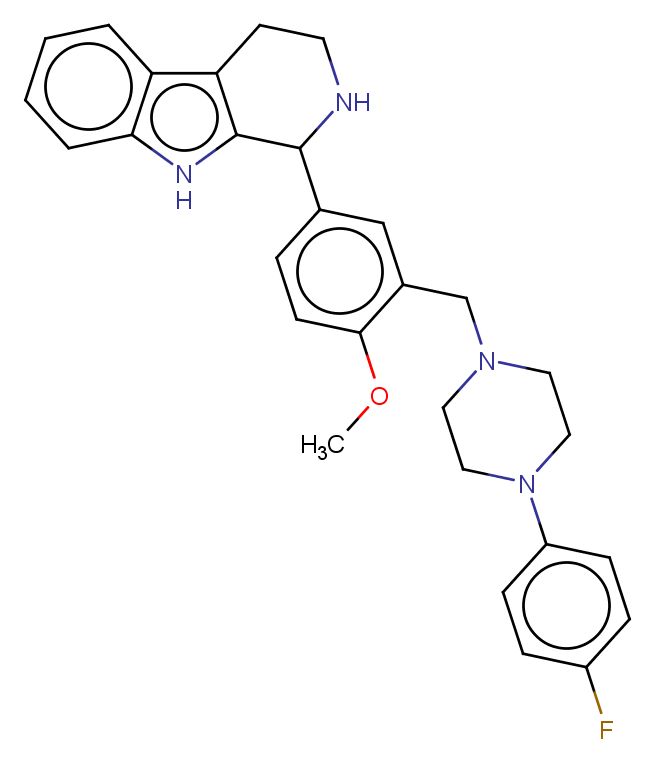

### MMV019700.png

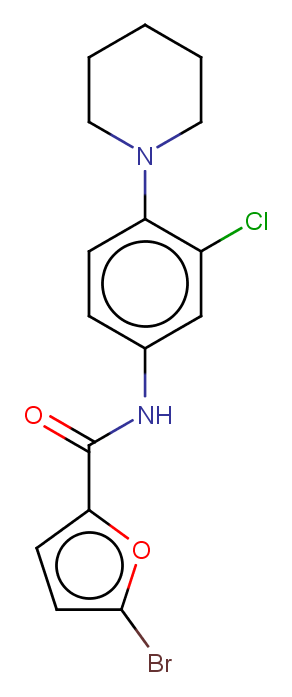

### MMV019738.png

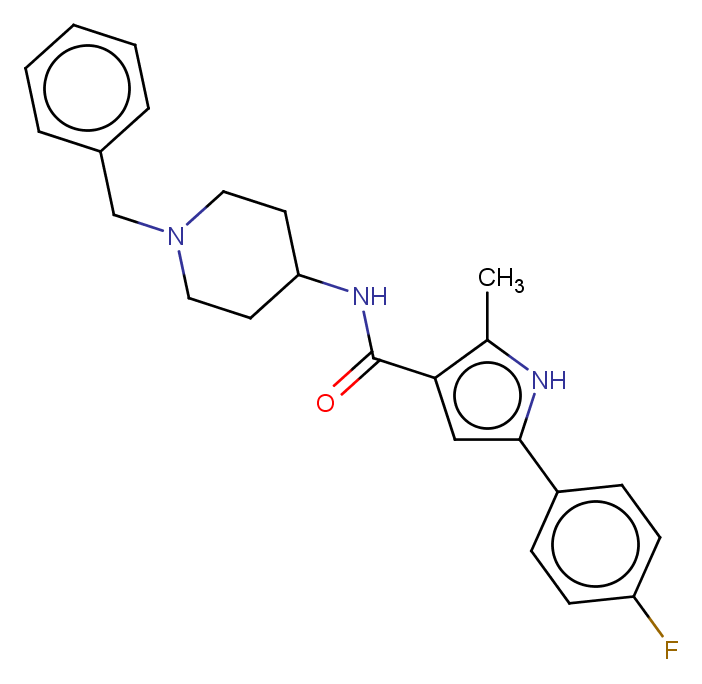

### MMV019741.png

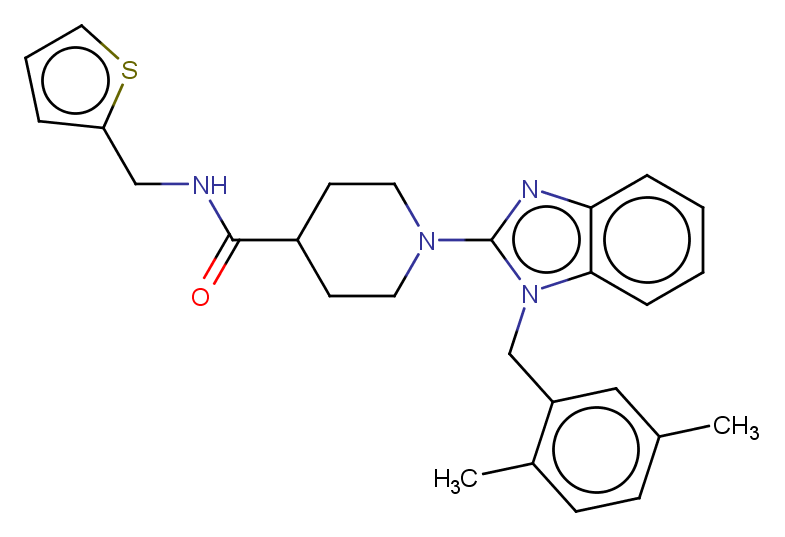

### MMV019746.png

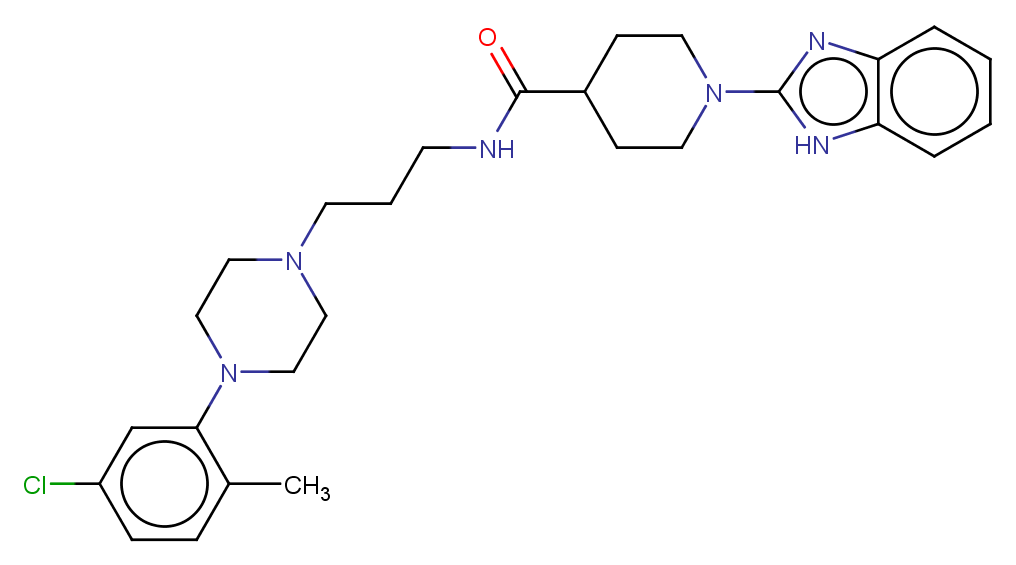

### MMV019758.png

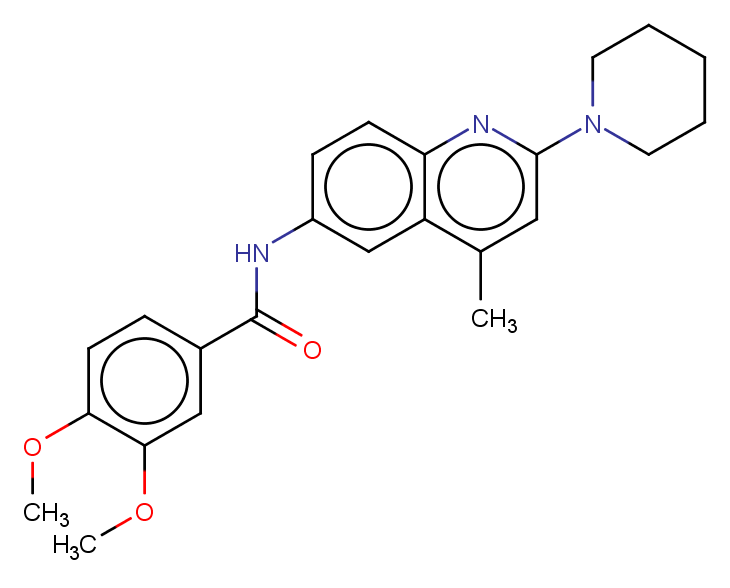

### MMV019762.png

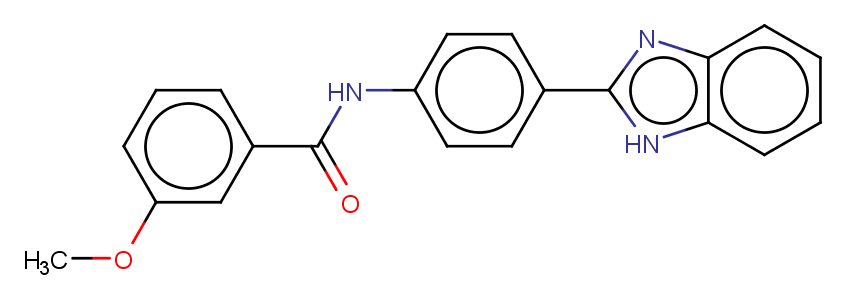

### MMV019780.png

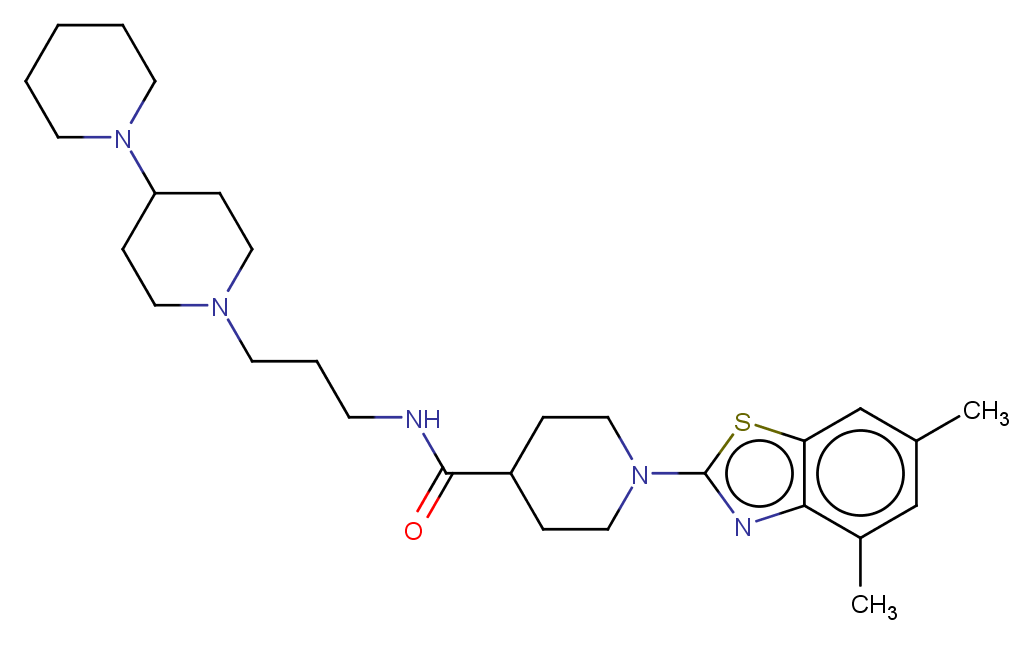

### MMV019871.png

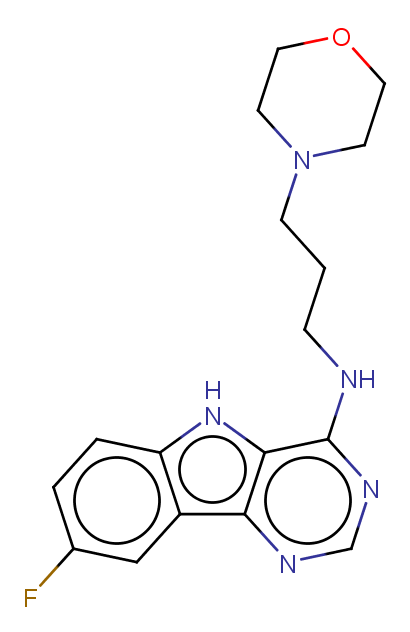

### MMV019881.png

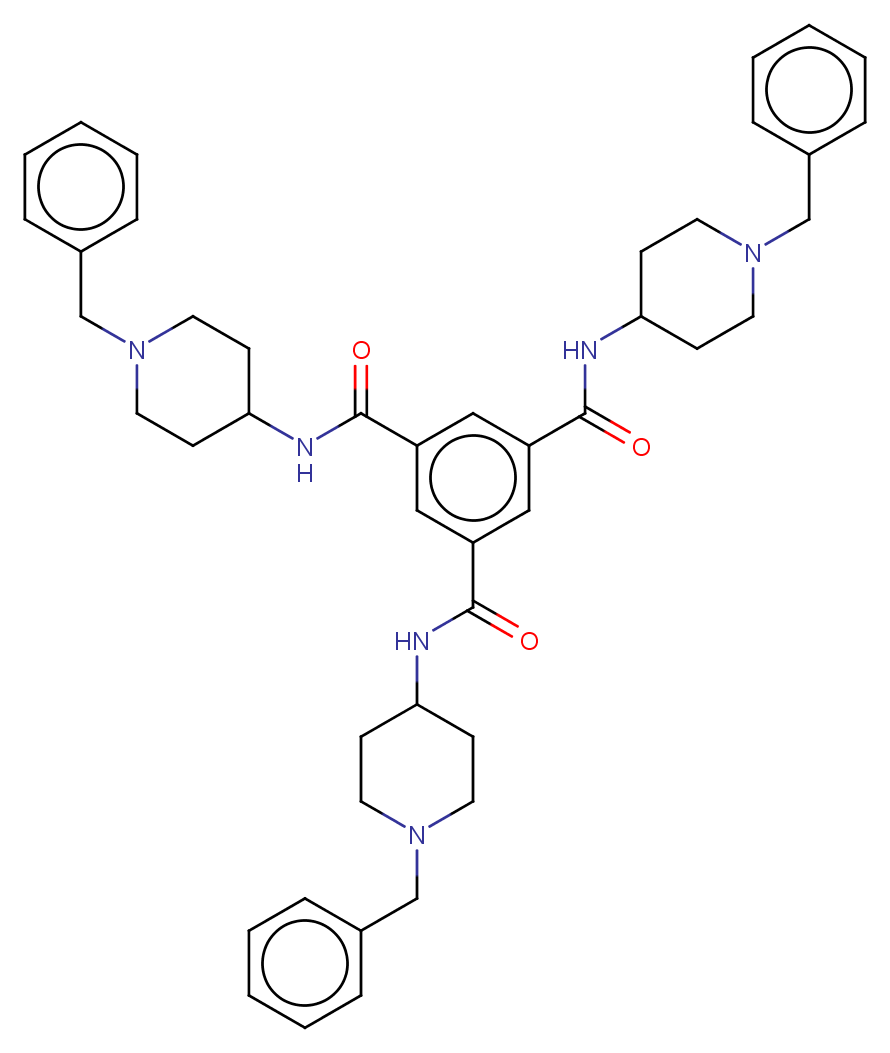

### MMV019918.png

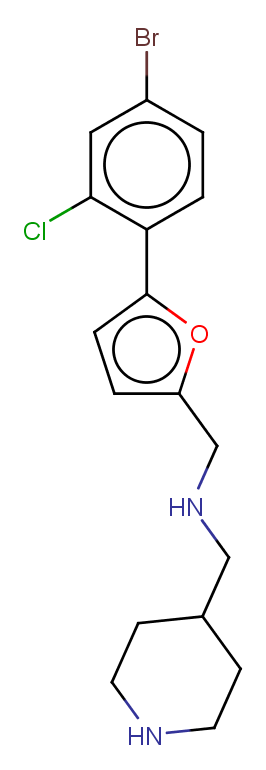

### MMV019995.png

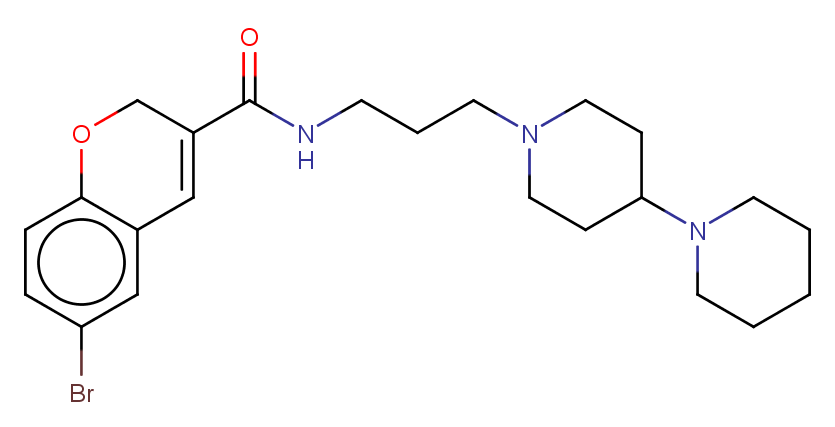

### MMV020243.png

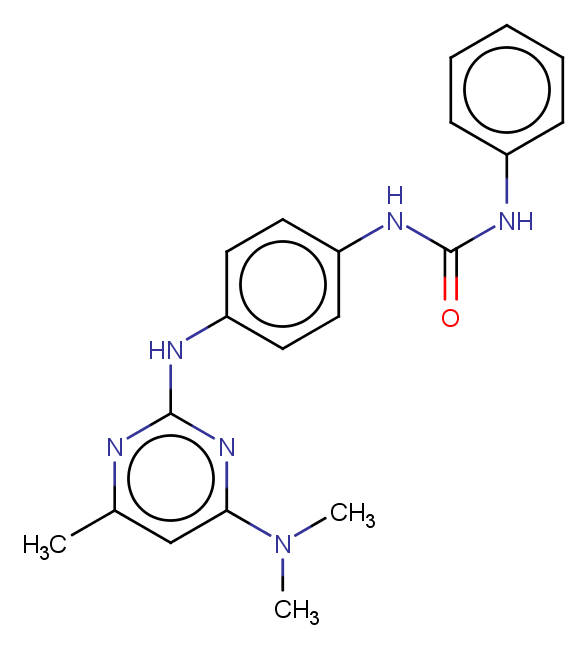

### MMV020275.png

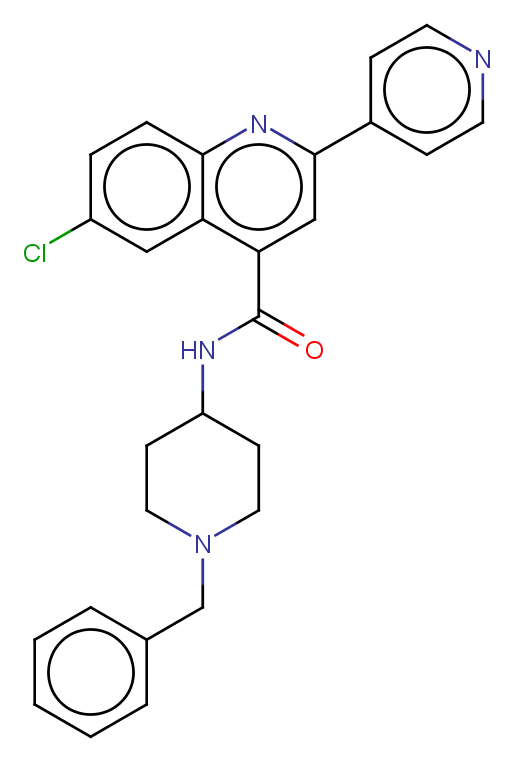

### MMV020403.png

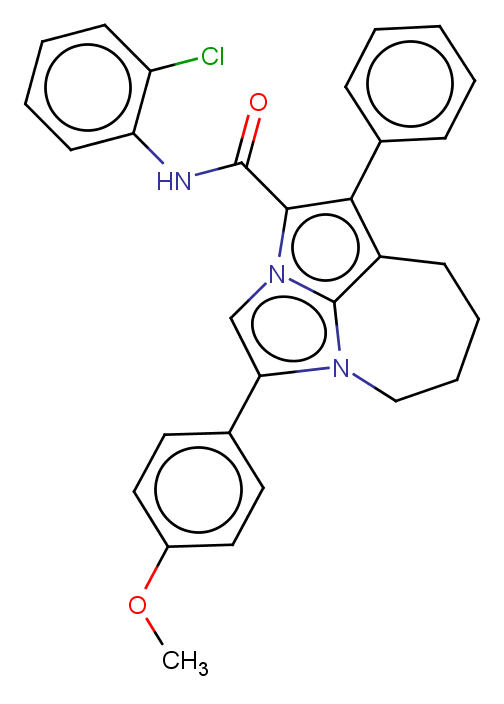

### MMV020439.png

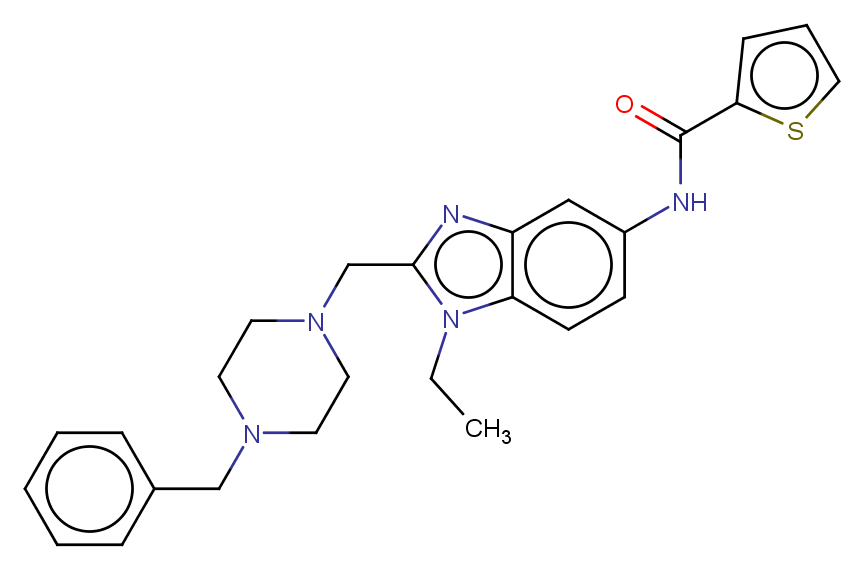

### MMV020490.png

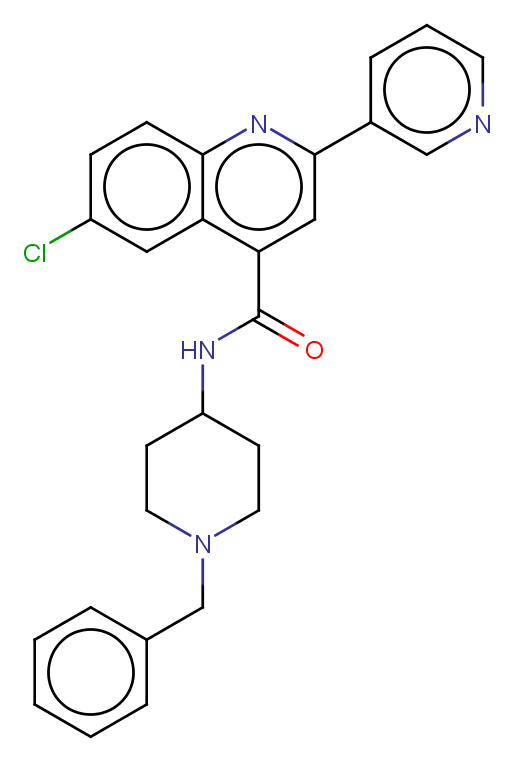

### MMV020492.png

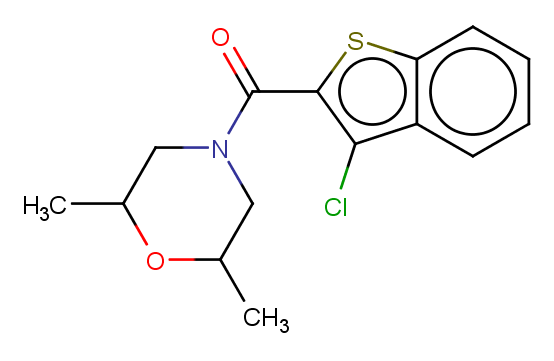

### MMV020500.png

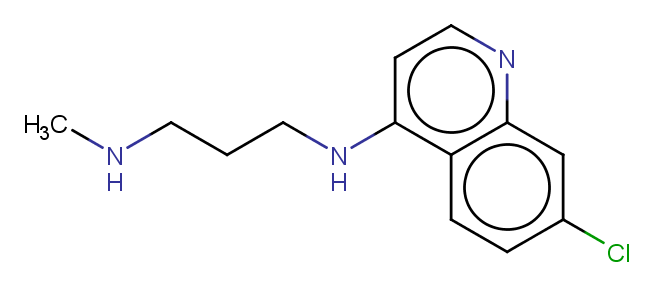

### MMV020505.png

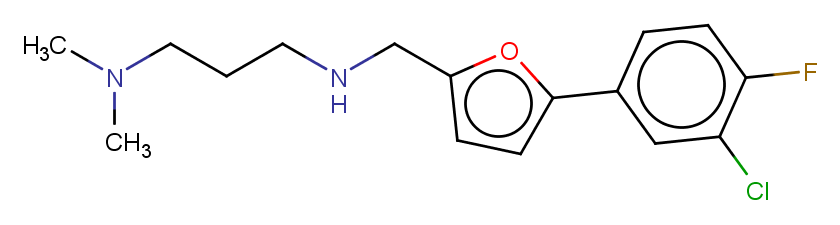

### MMV020548.png

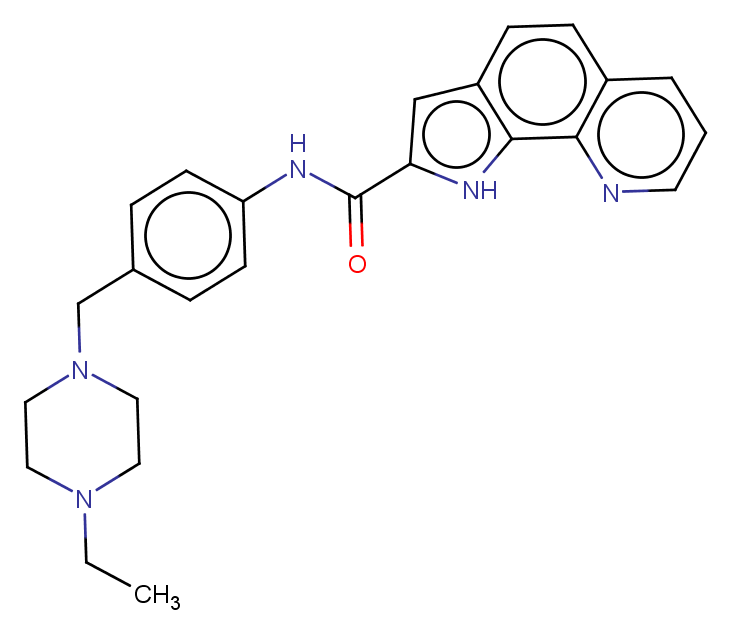

### MMV020549.png

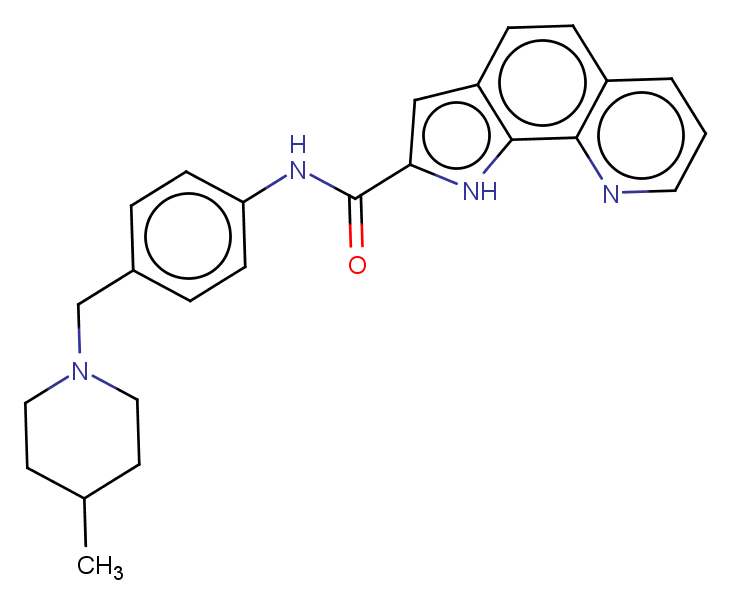

### MMV020651.png

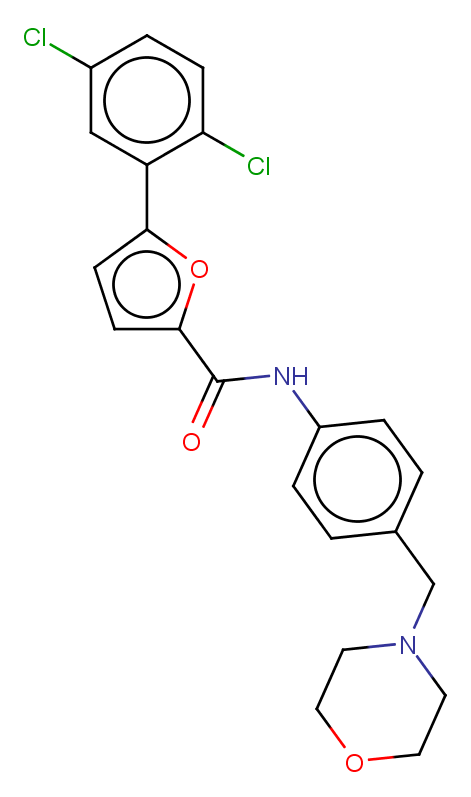

### MMV020654.png

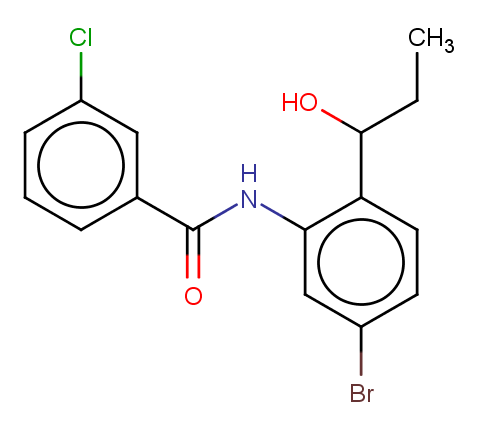
