## Supplementary figures and images for "Refining physico-chemical rules for herbicides using an antimalarial library"

### MMV000304.png

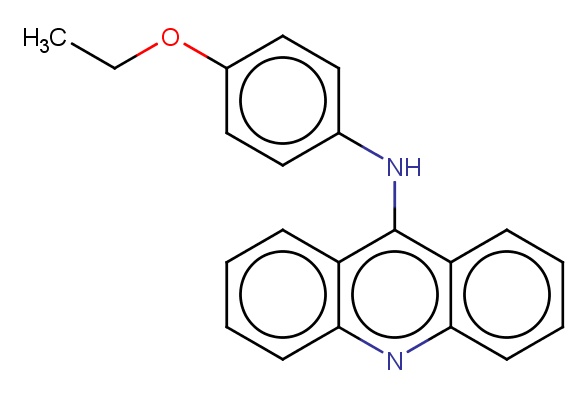

### MMV000326.png

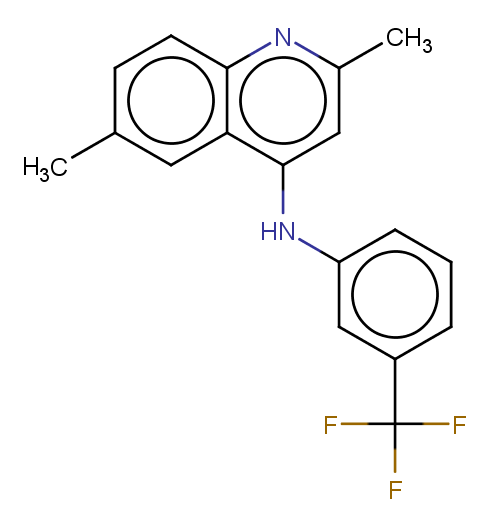

### MMV000340.png

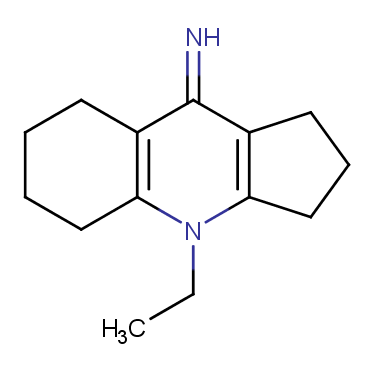

### MMV000356.png

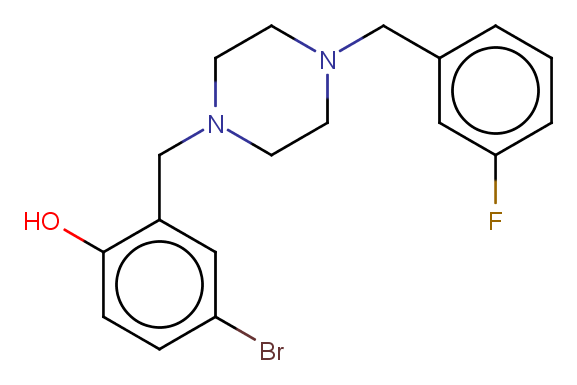

### MMV000442.png

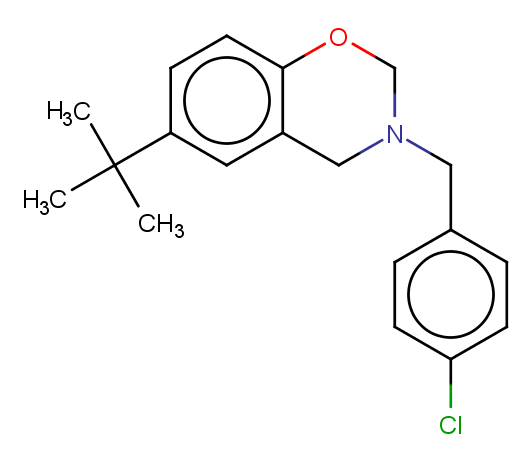

### MMV000443.png

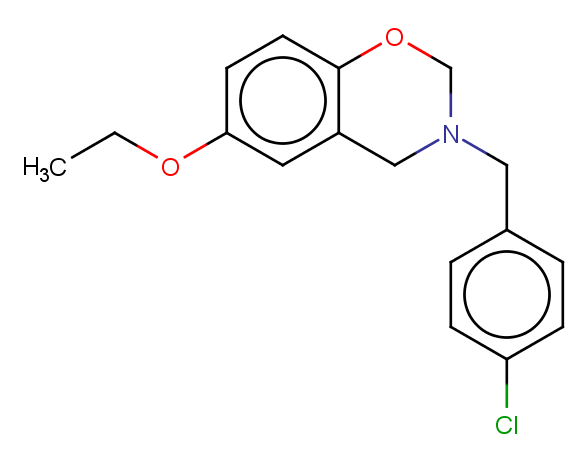

### MMV000444.png

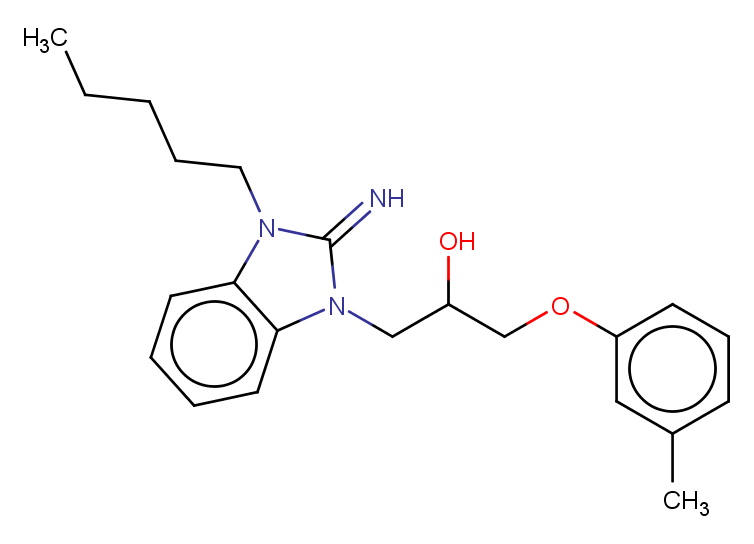

### MMV000445.png

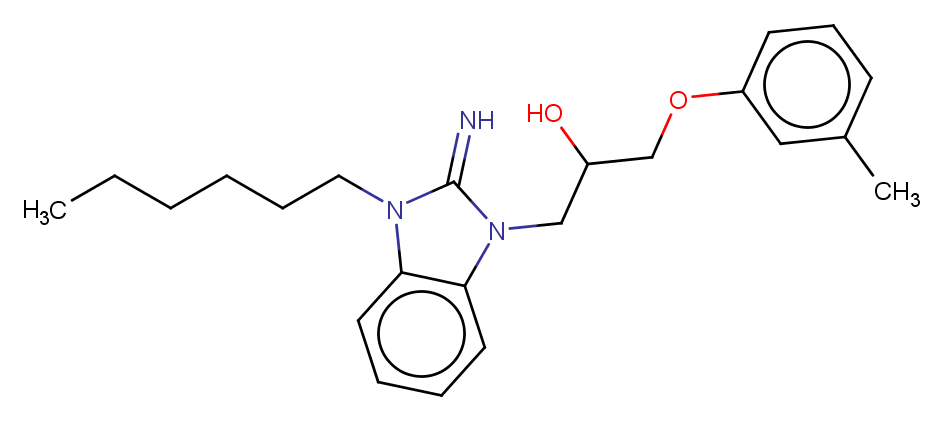

### MMV000448.png

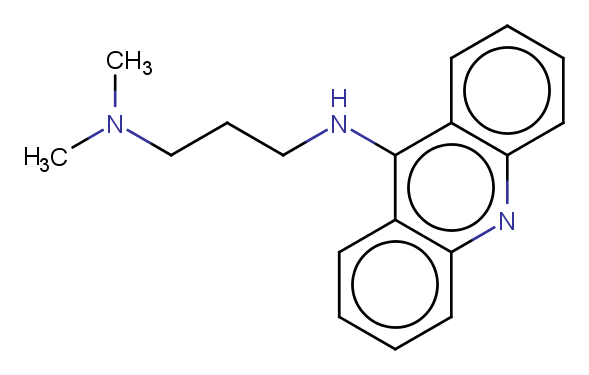

### MMV000478.png

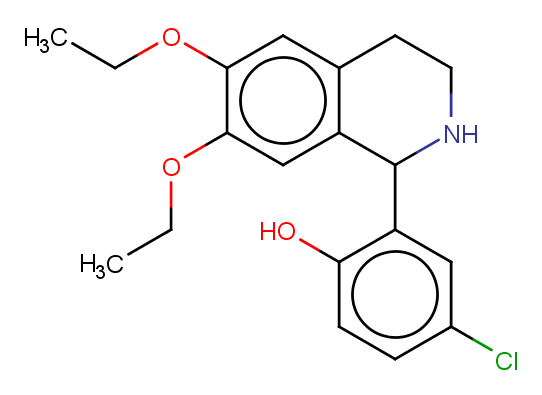

### MMV000481.png

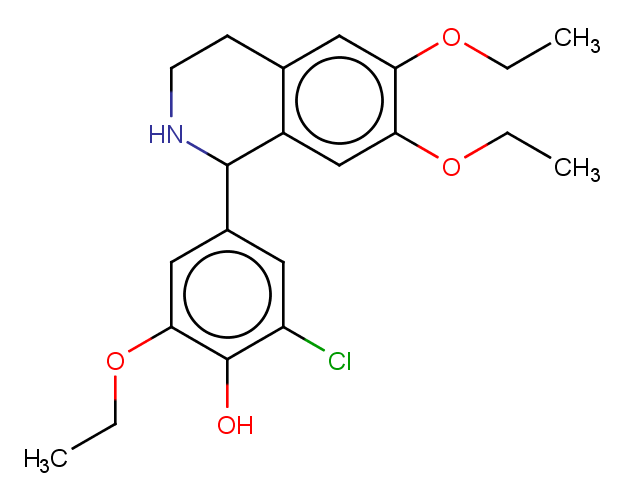

### MMV000483.png

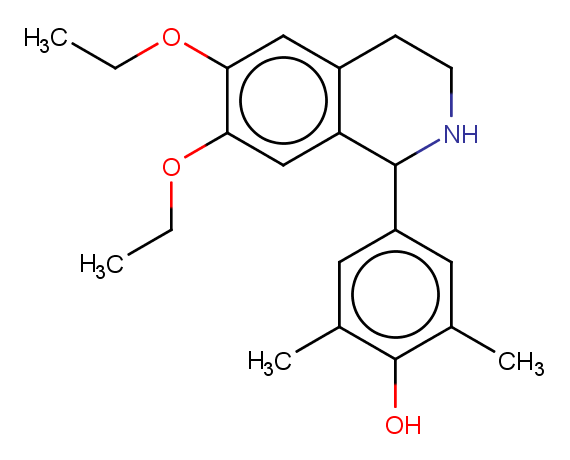

### MMV000498.png

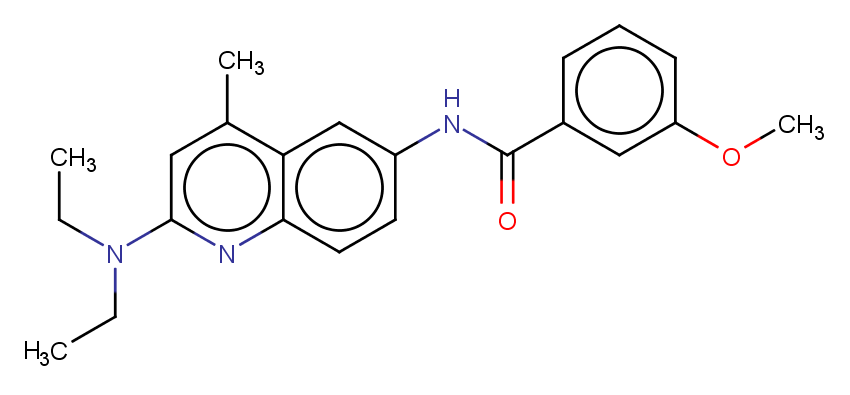

### MMV000561.png

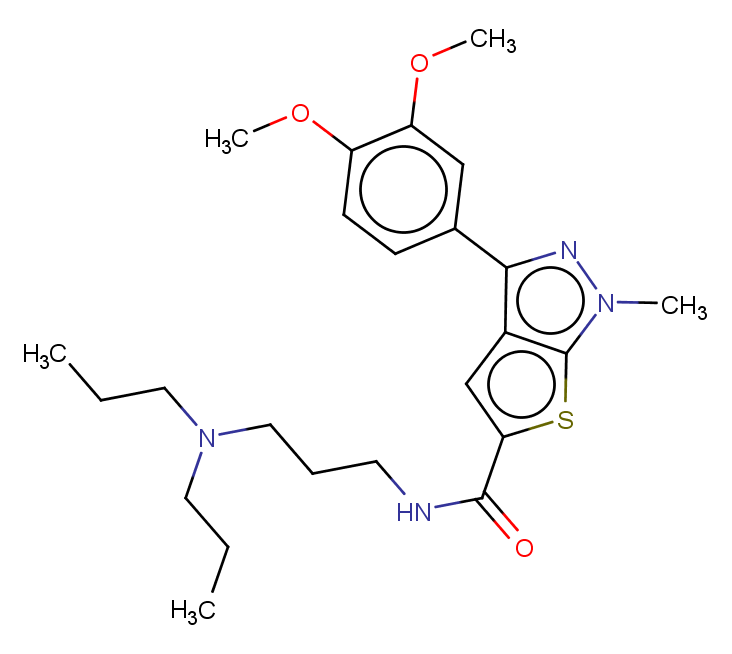

### MMV000563.png

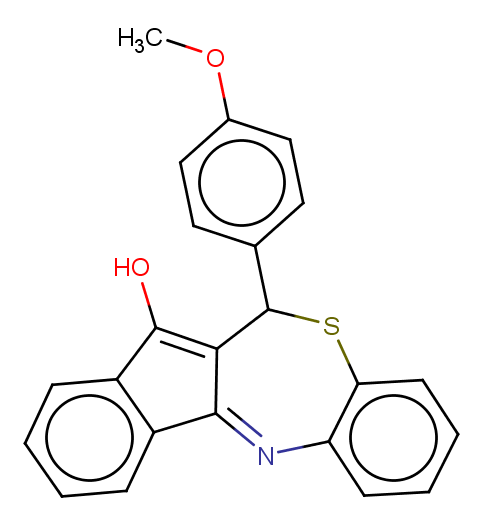

### MMV000570.png

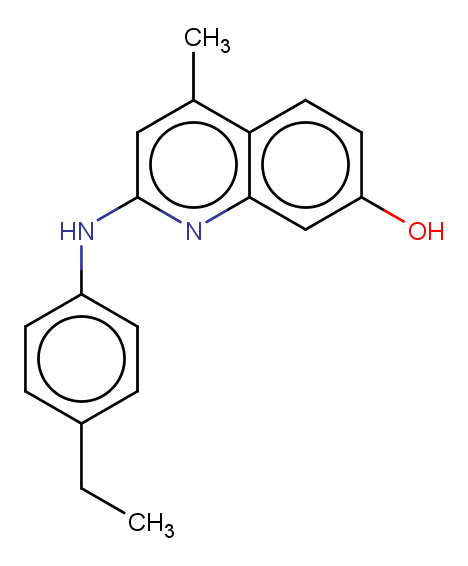

### MMV000604.png

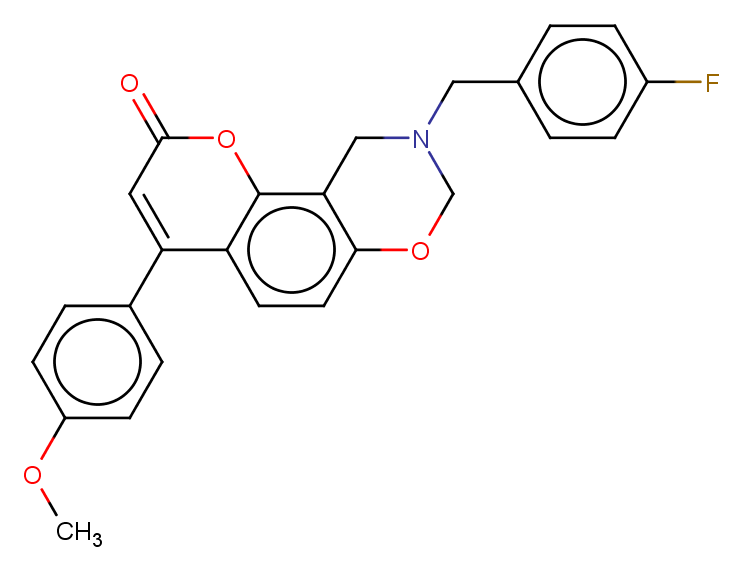

### MMV000617.png

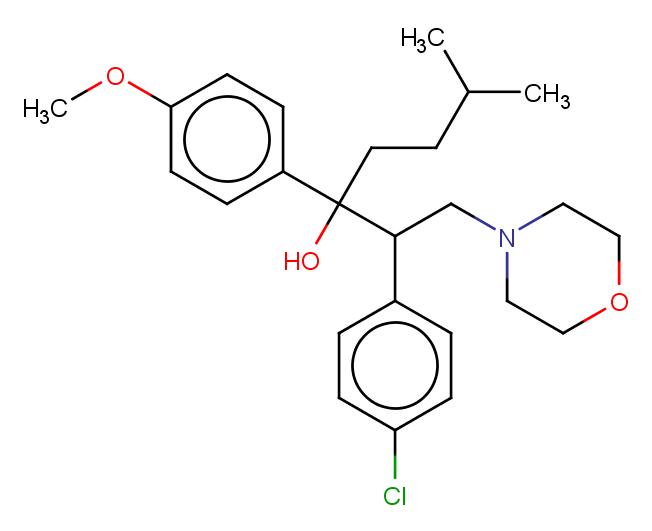

### MMV000619.png

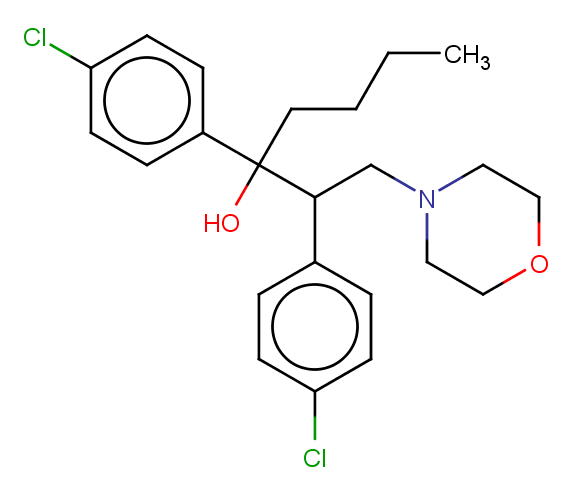

### MMV000620.png

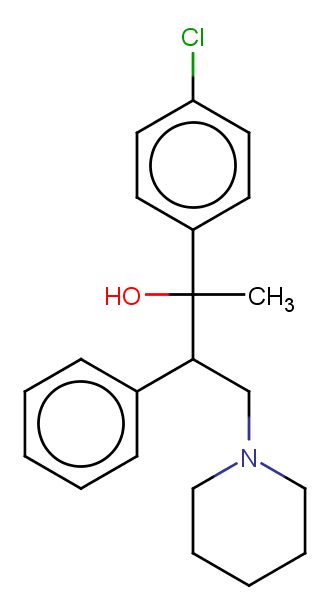

### MMV000621.png

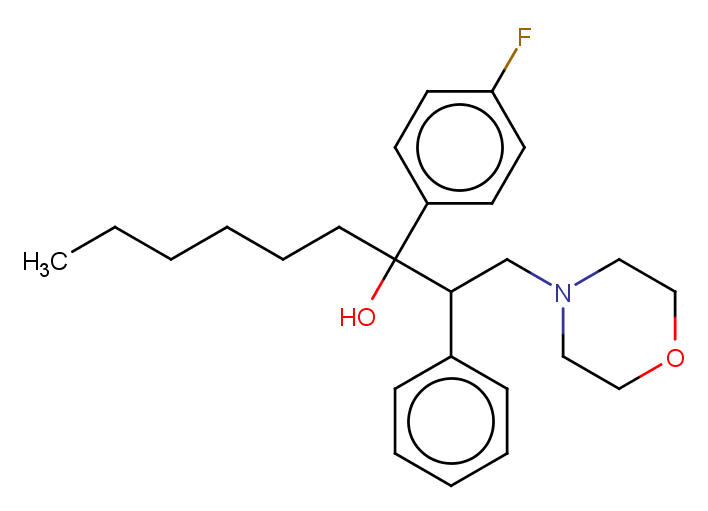

### MMV000634.png

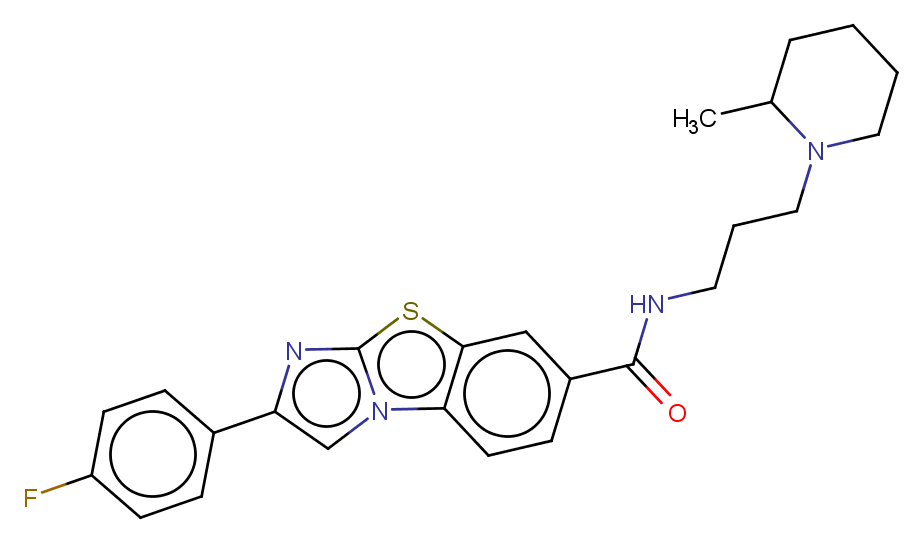

### MMV000642.png

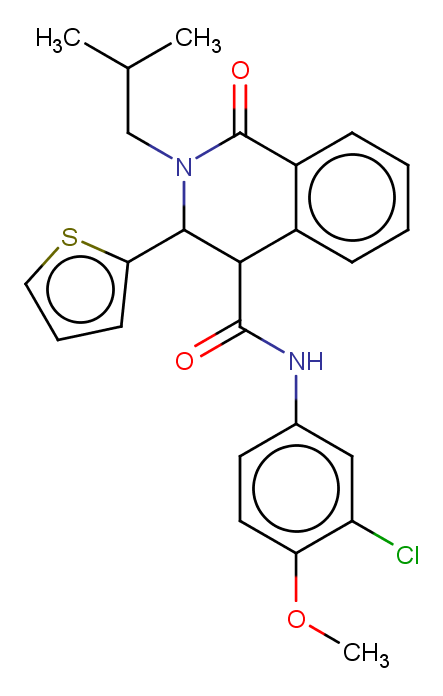

### MMV000648.png

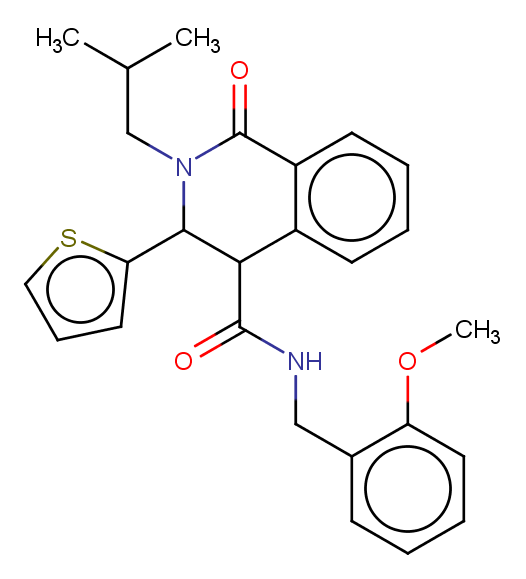

### MMV000653.png

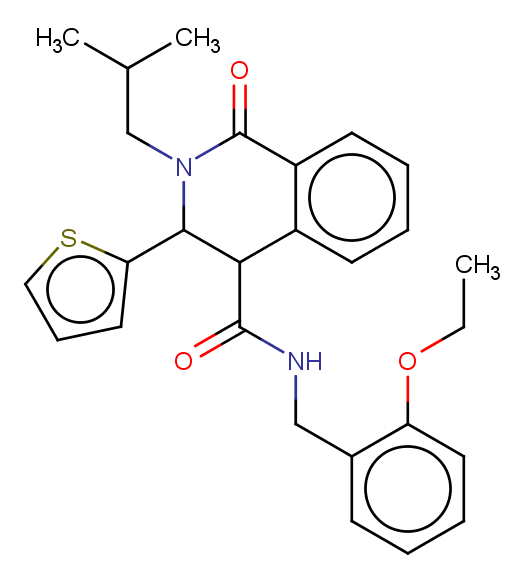

### MMV000662.png

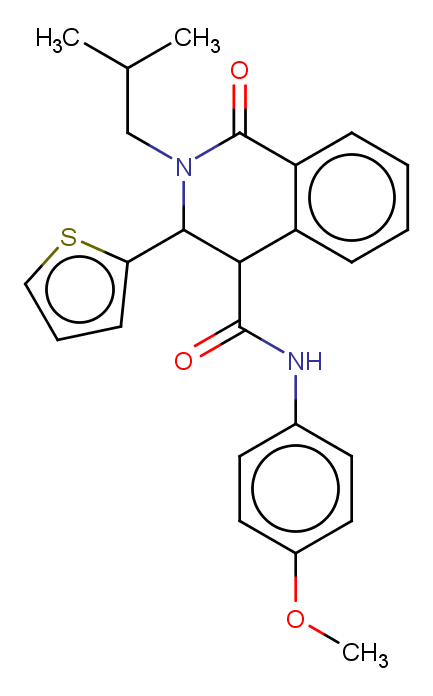

### MMV000699.png

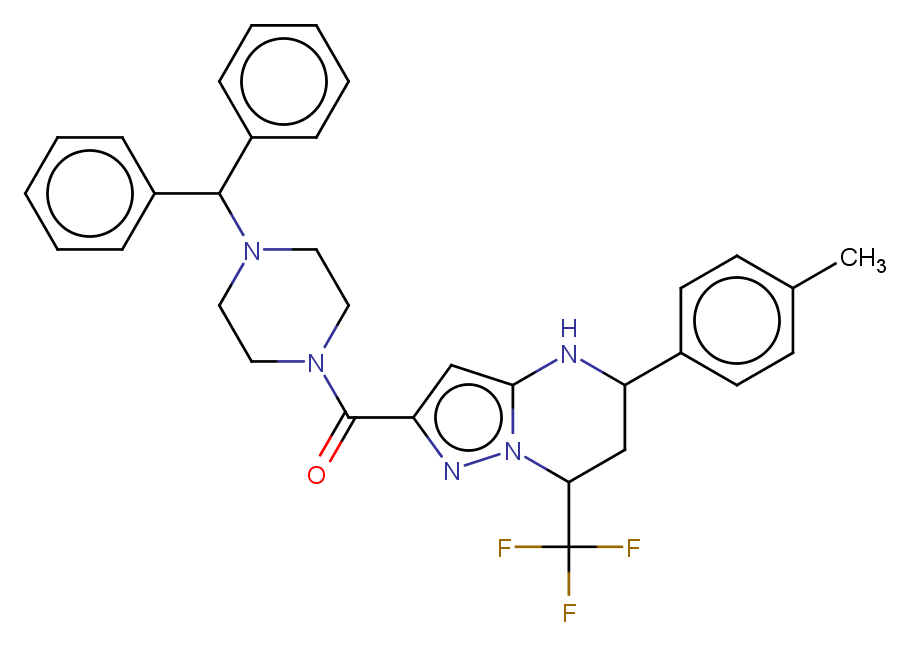

### MMV000704.png

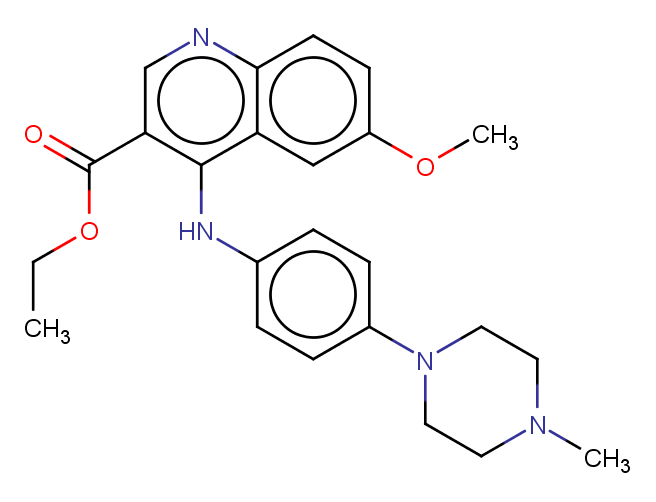

### MMV000720.png

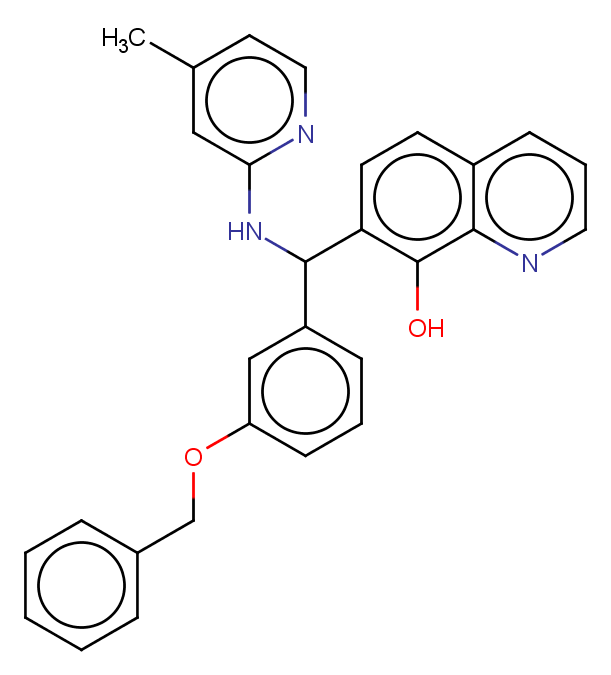

### MMV000753.png

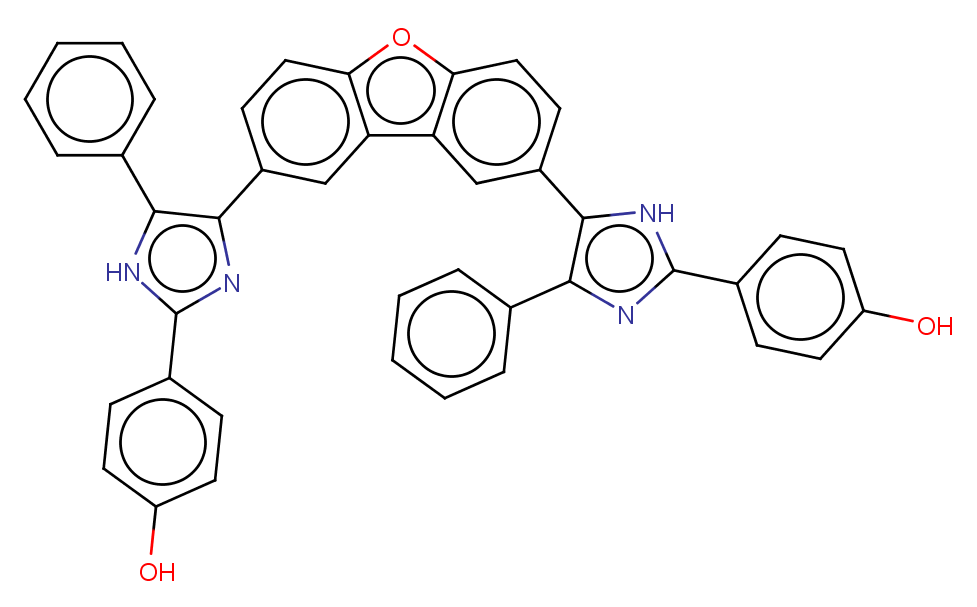

### MMV000760.png

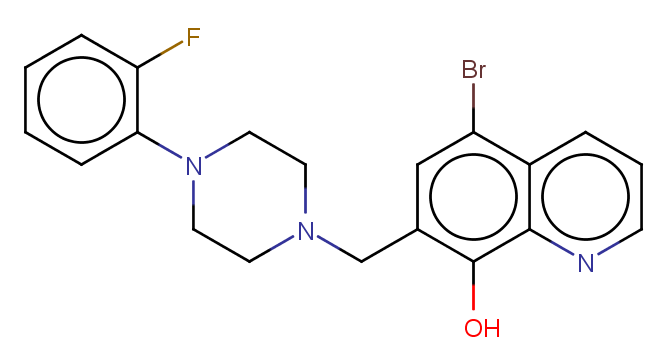

### MMV000787.png

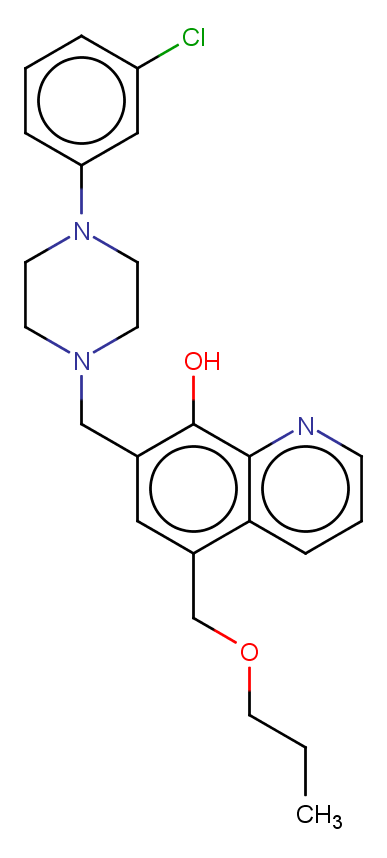

### MMV000788.png

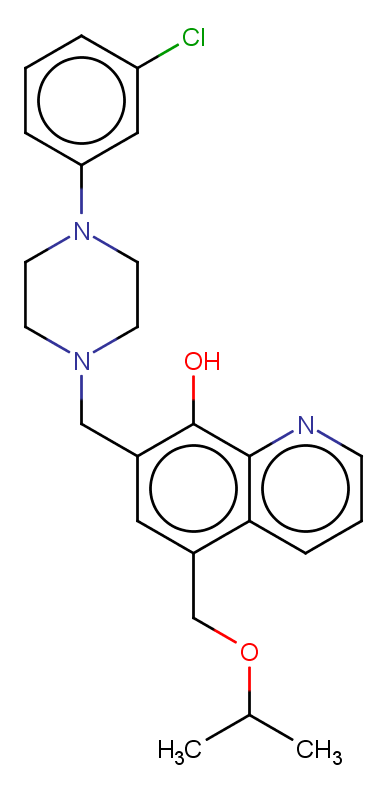

### MMV000839.png

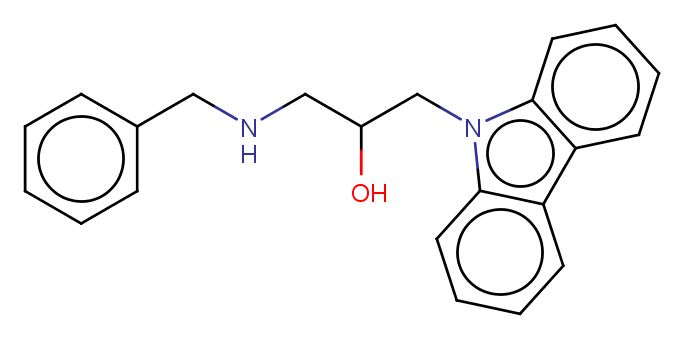

### MMV000848.png

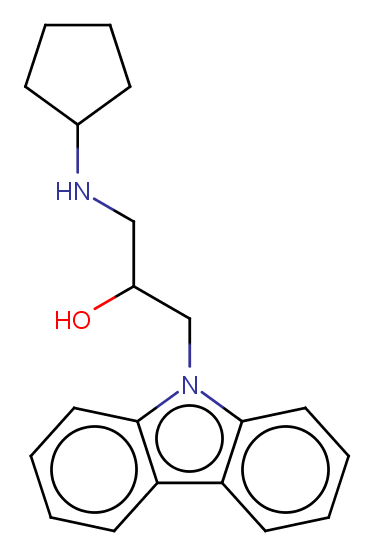

### MMV000911.png

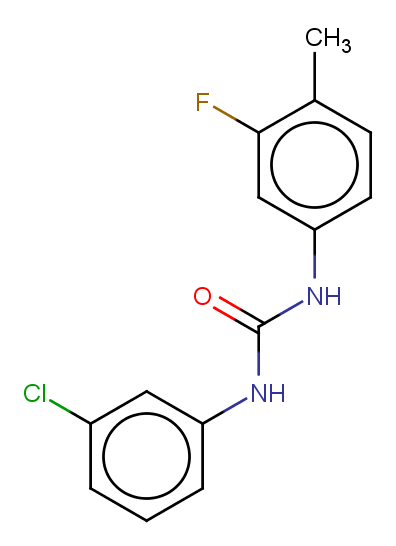

### MMV000917.png

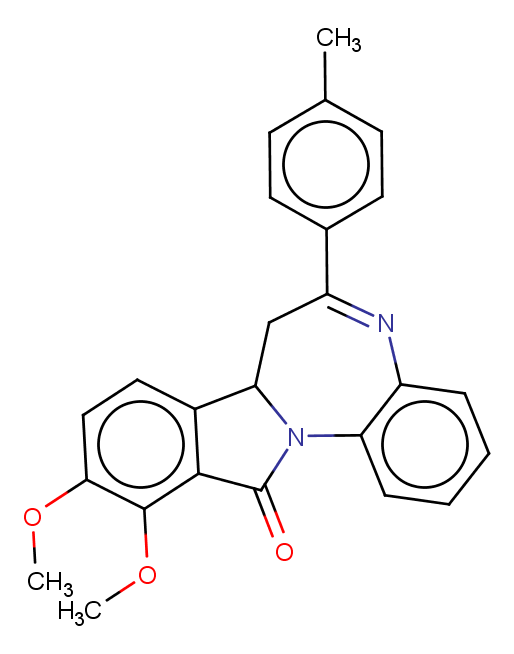

### MMV000963.png

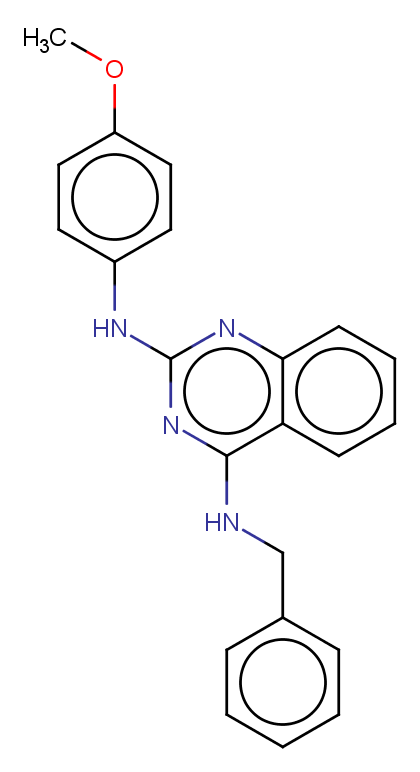

### MMV000972.png

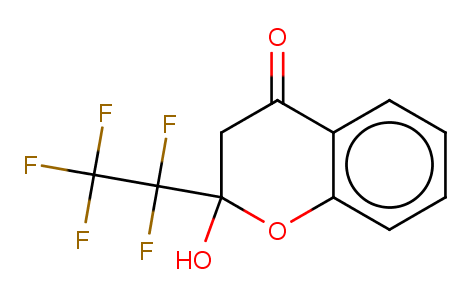

### MMV000986.png

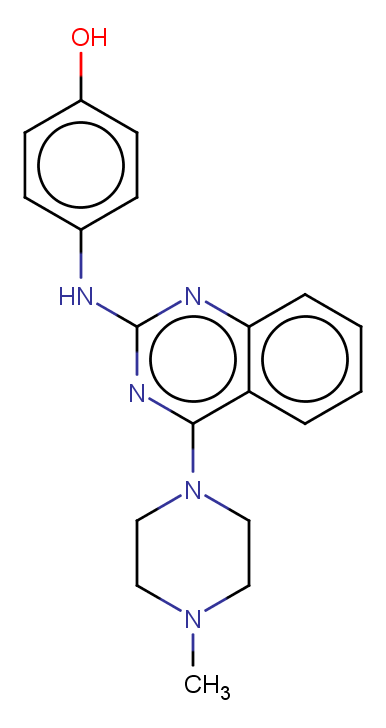

### MMV001038.png

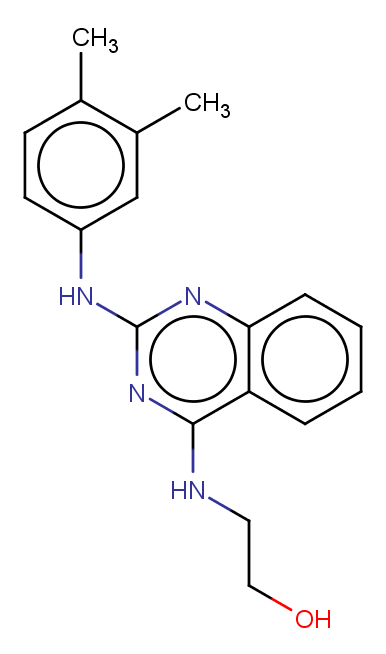

### MMV001041.png

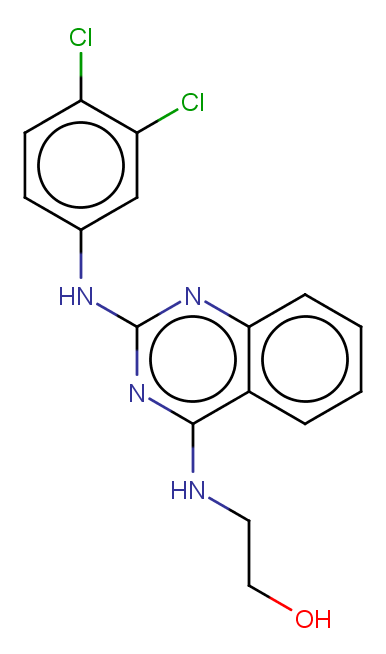

### MMV001049.png

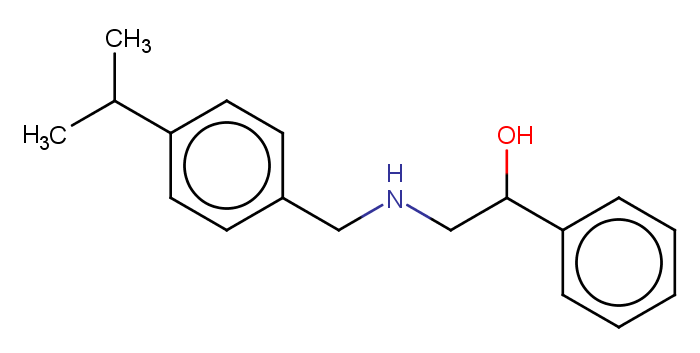

### MMV001230.png

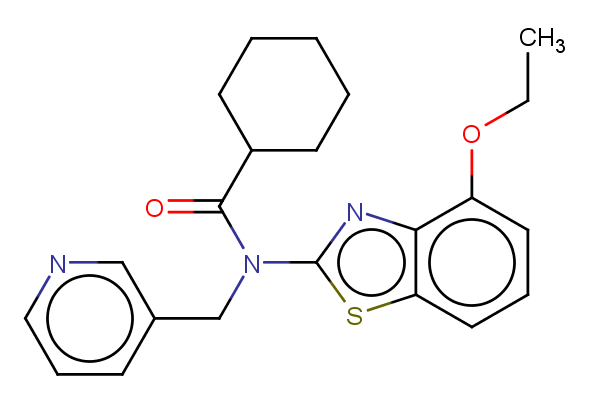

### MMV001239.png

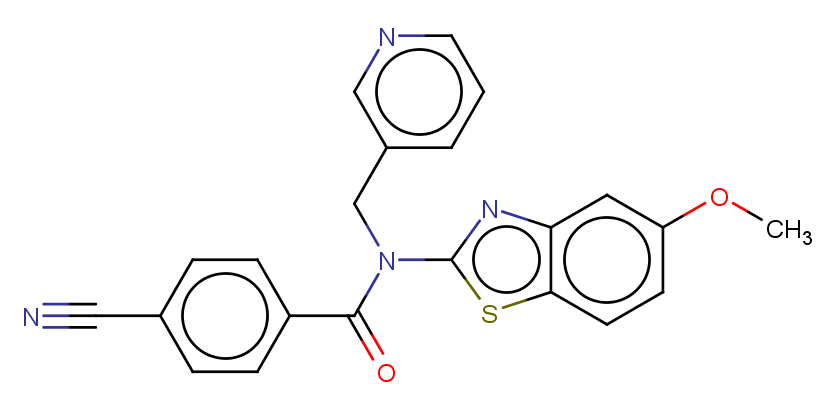

### MMV001241.png

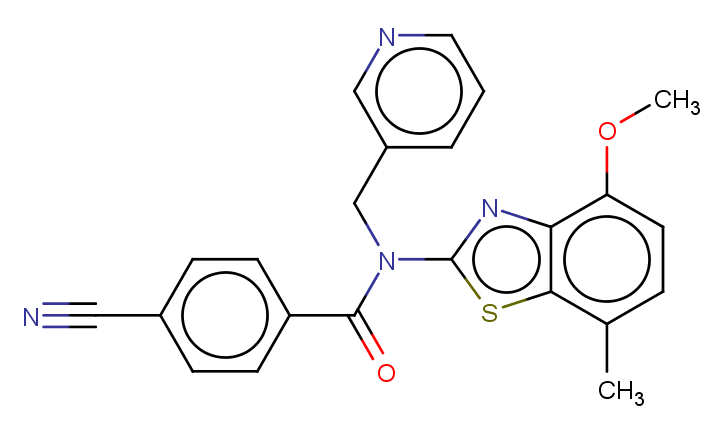

### MMV001246.png

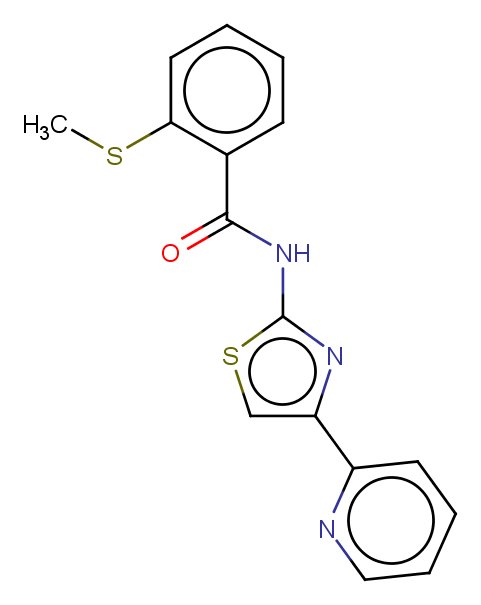

### MMV001255.png

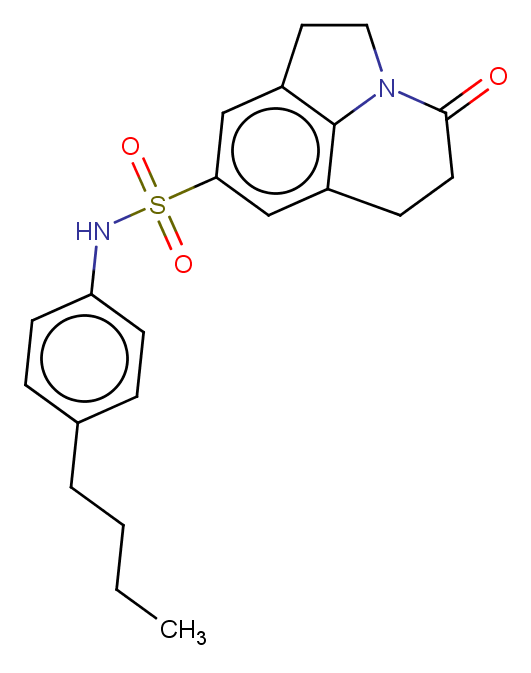

### MMV001318.png

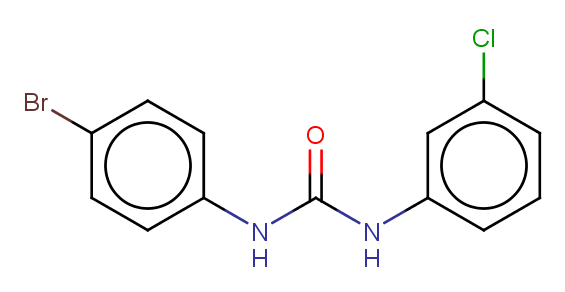

### MMV001344.png

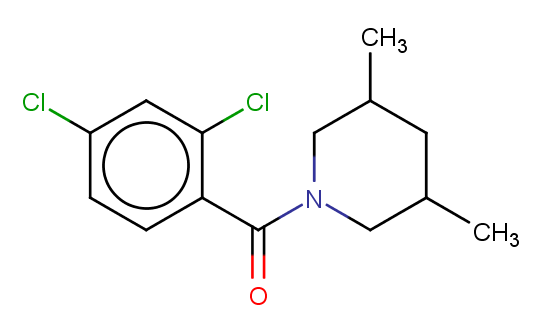

### MMV006087.png

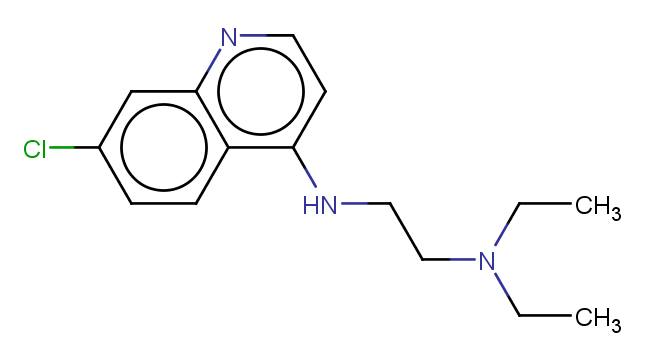

### MMV006169.png

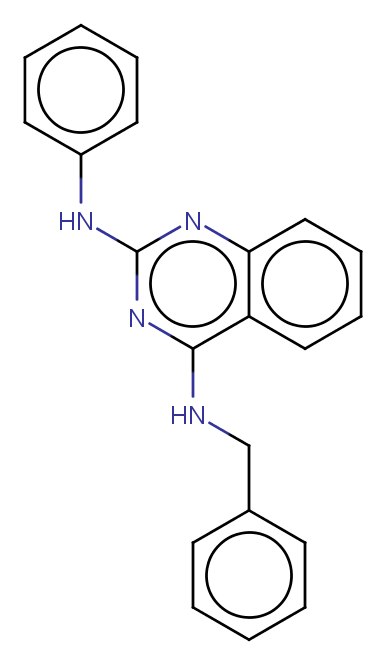

### MMV006172.png

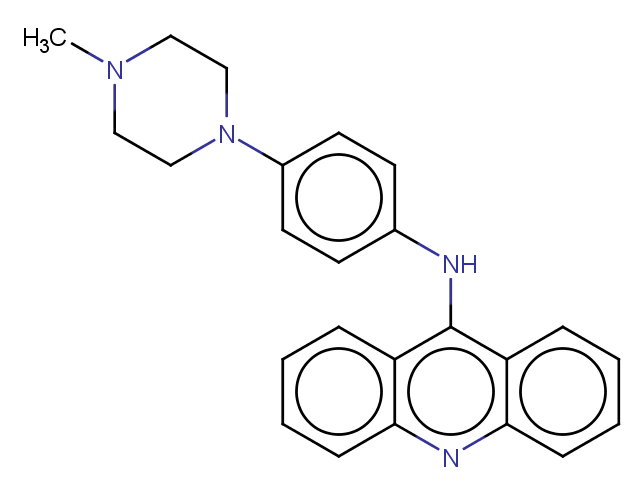

### MMV006188.png

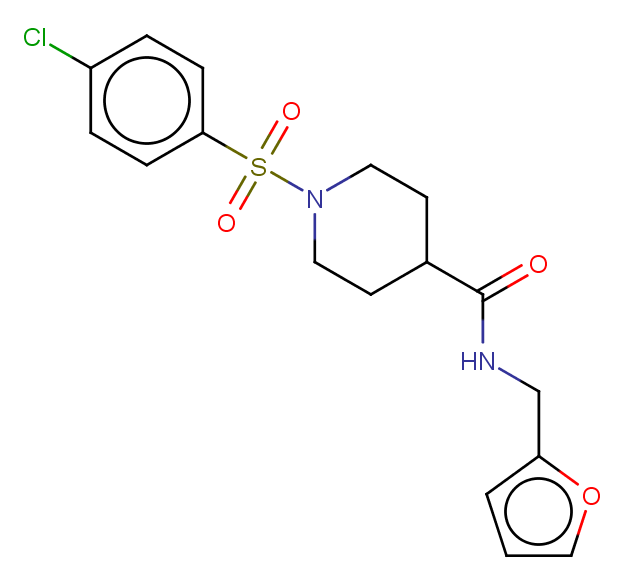

### MMV006203.png

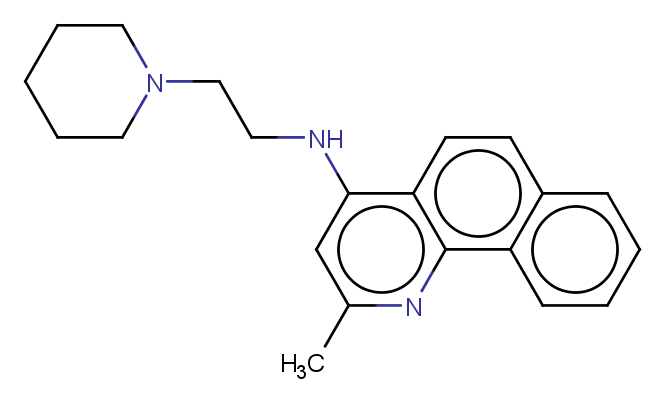

### MMV006250.png

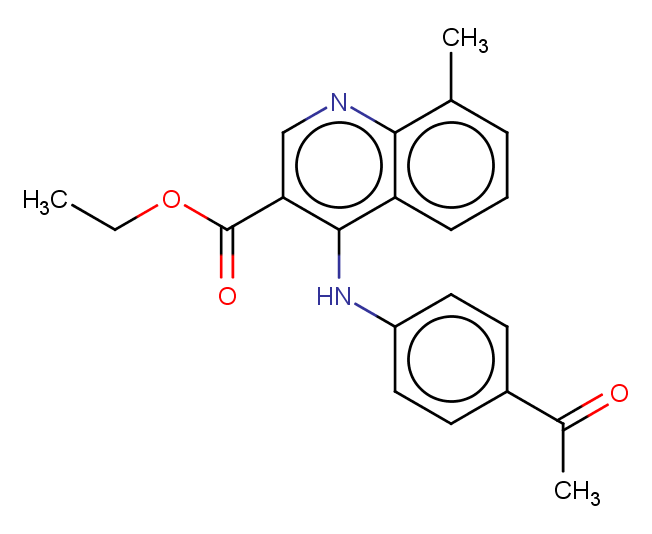

### MMV006278.png

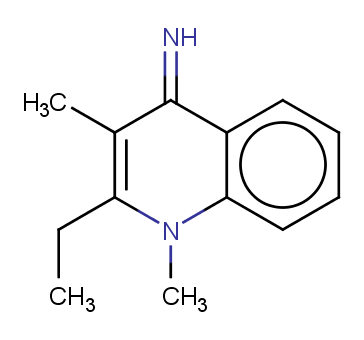

### MMV006303.png

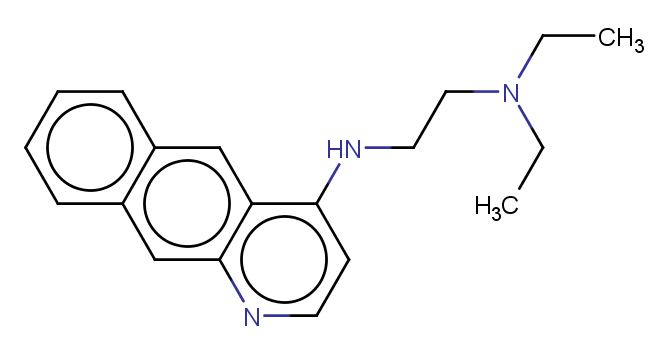

### MMV006309.png

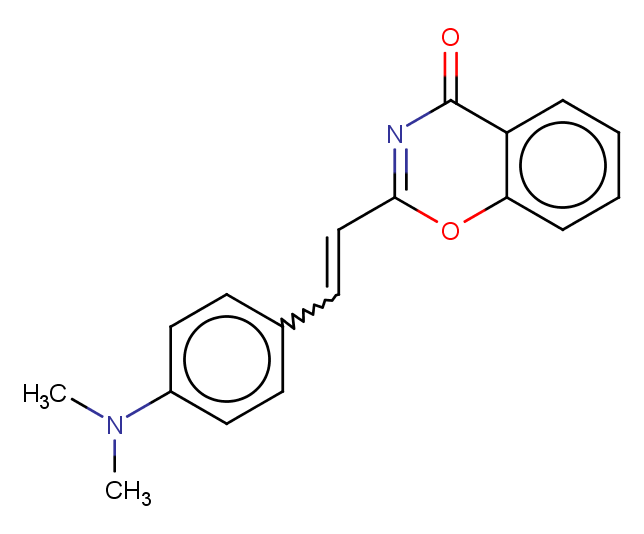

### MMV006319.png

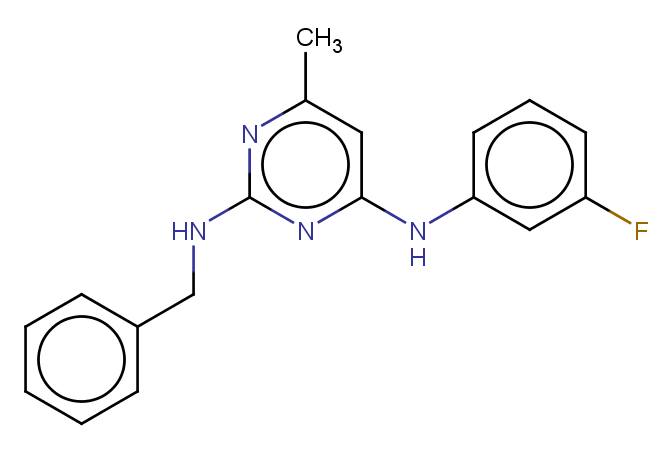

### MMV006389.png

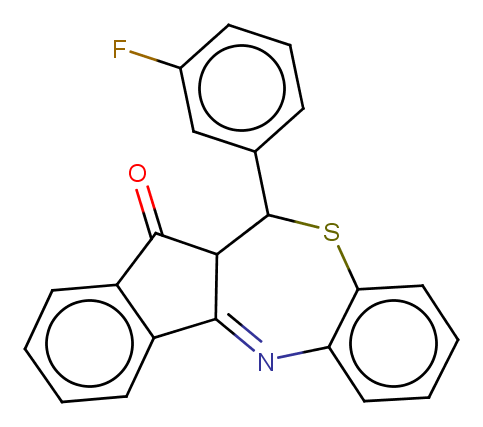

### MMV006427.png

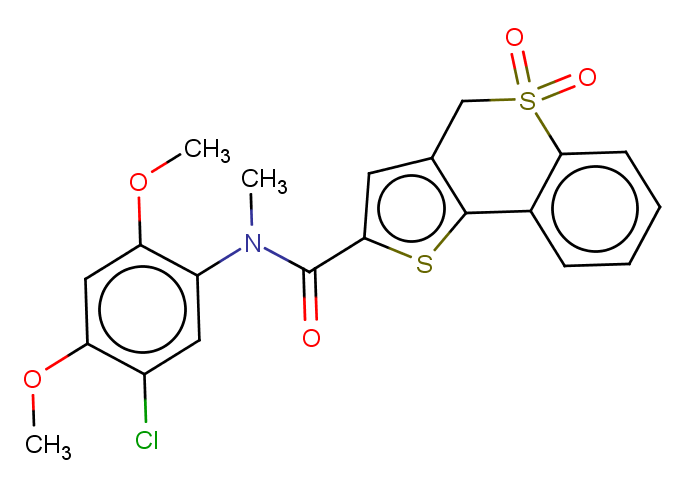

### MMV006429.png

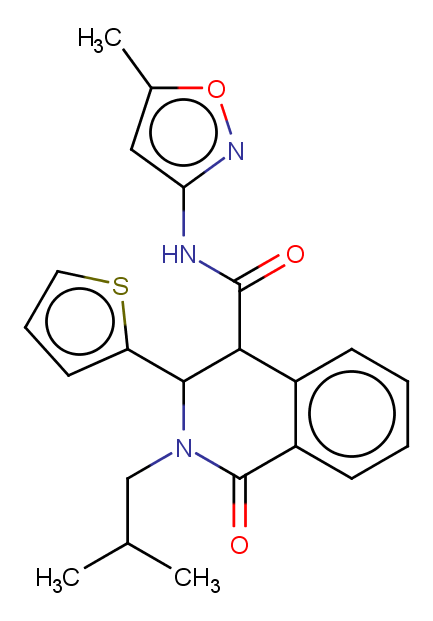

### MMV006455.png

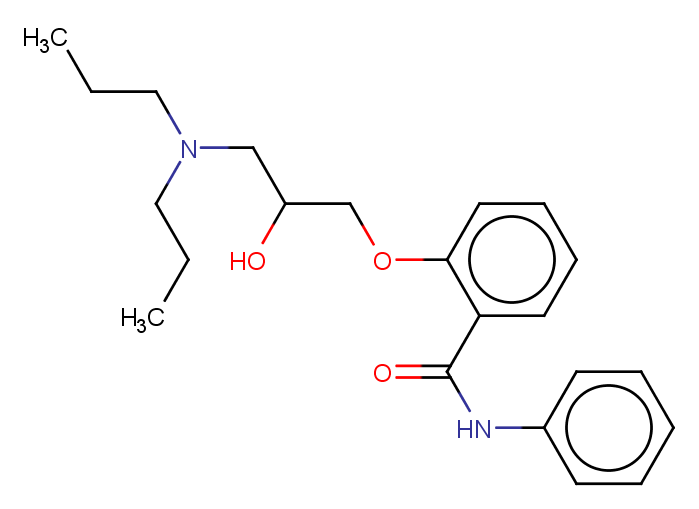

### MMV006457.png

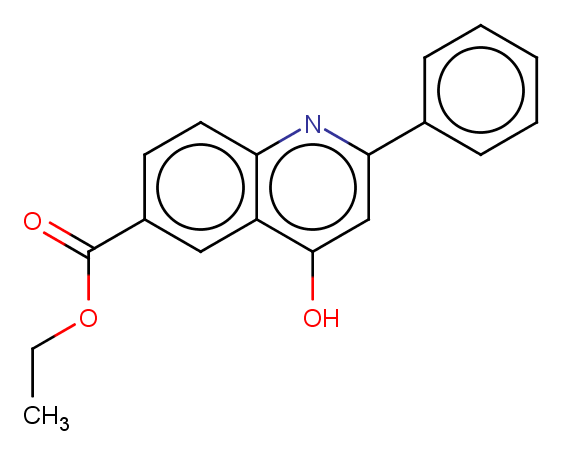

### MMV006513.png

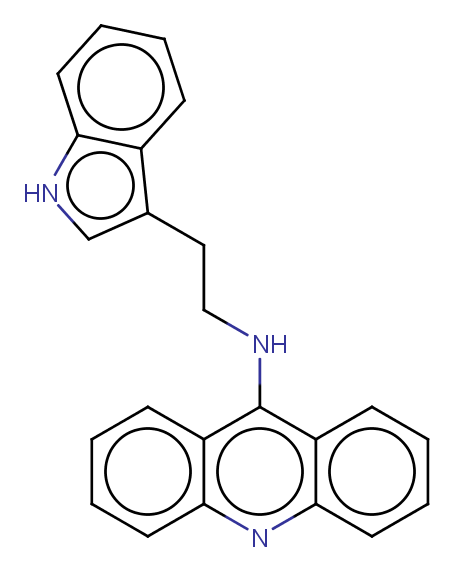

### MMV006522.png

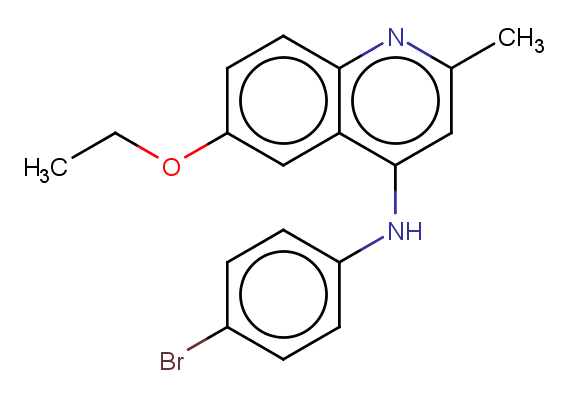

### MMV006545.png

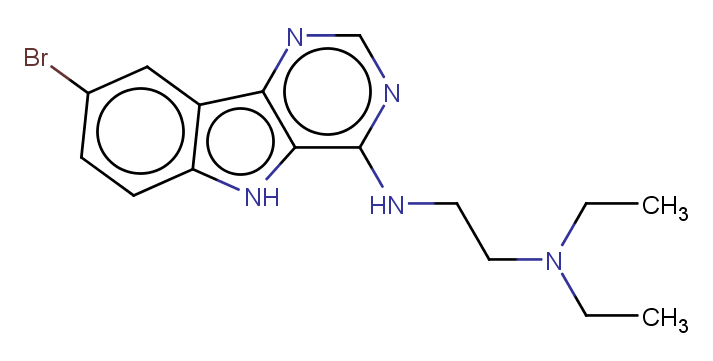

### MMV006558.png

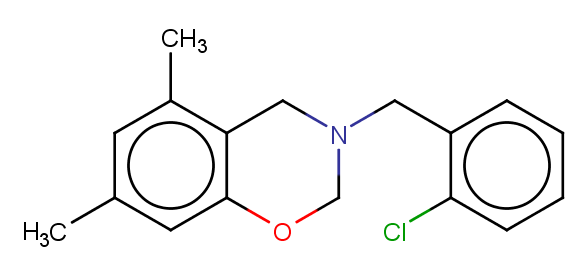

### MMV006587.png

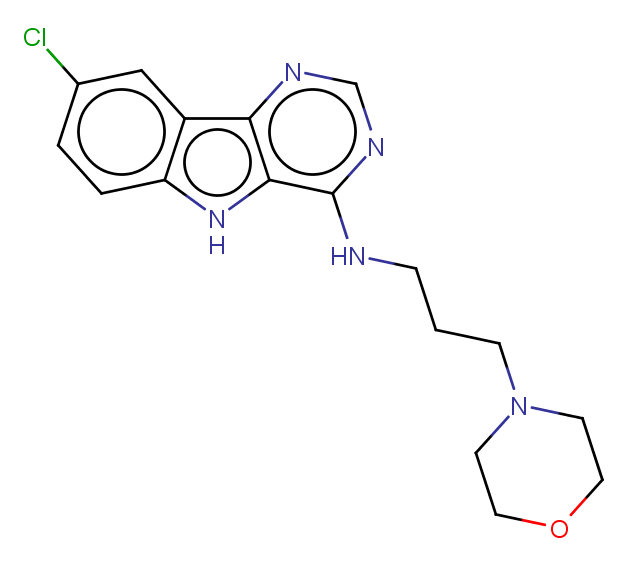

### MMV006656.png

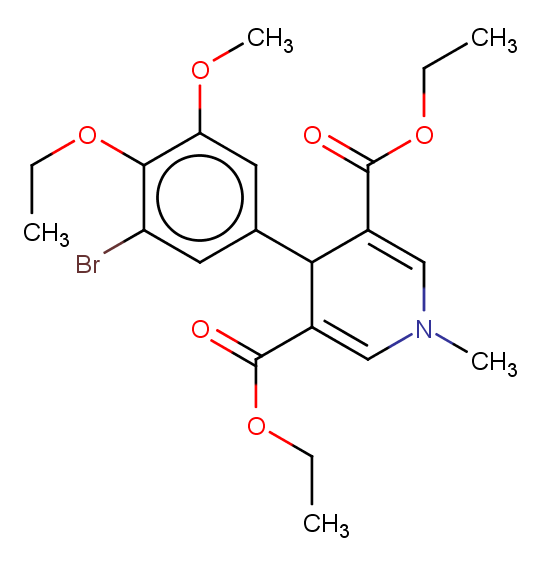

### MMV006704.png

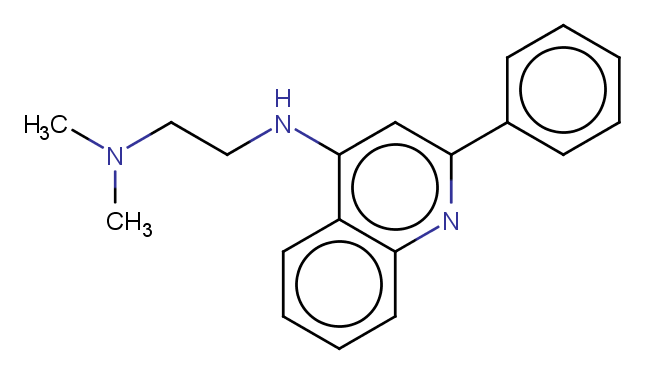

### MMV006706.png

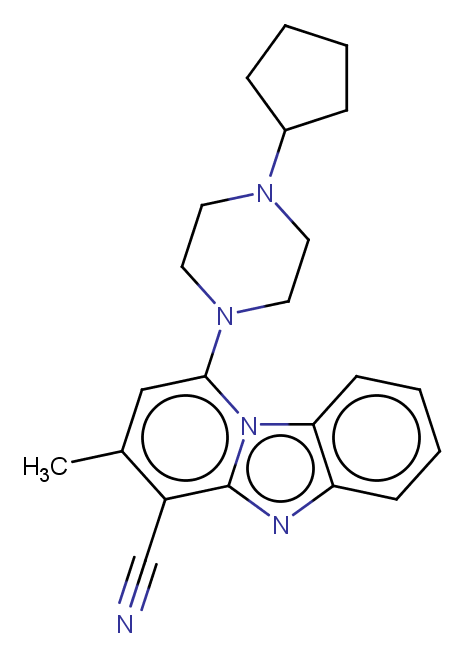

### MMV006753.png

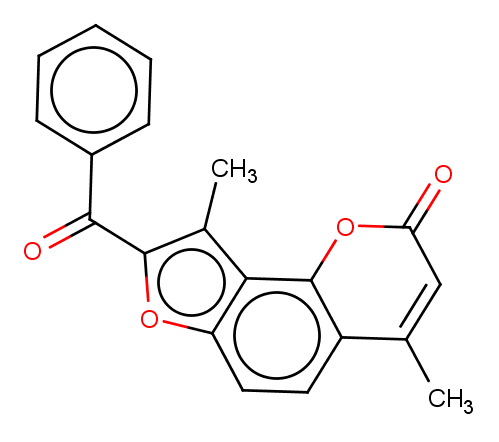

### MMV006764.png

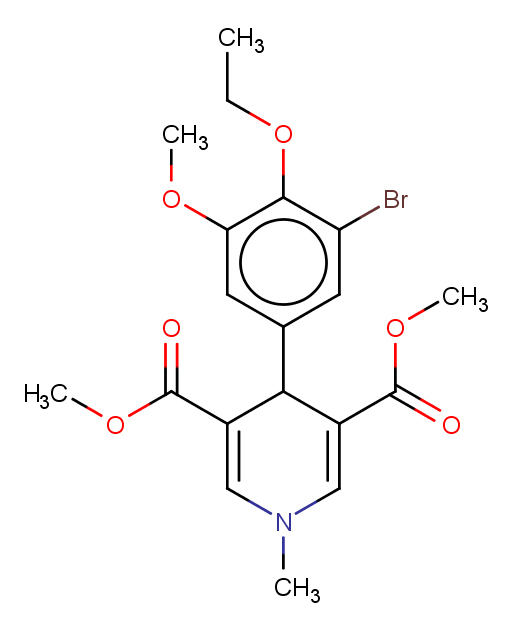

### MMV006767.png

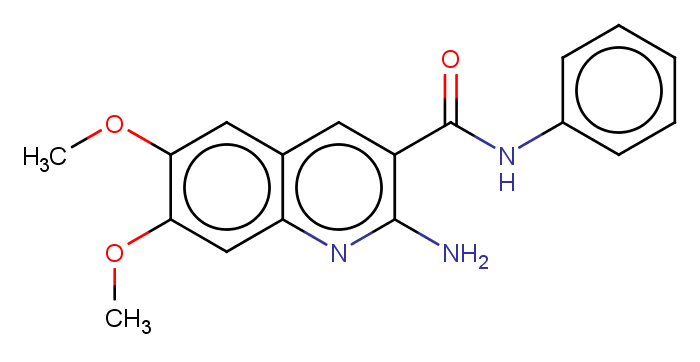

### MMV006787.png

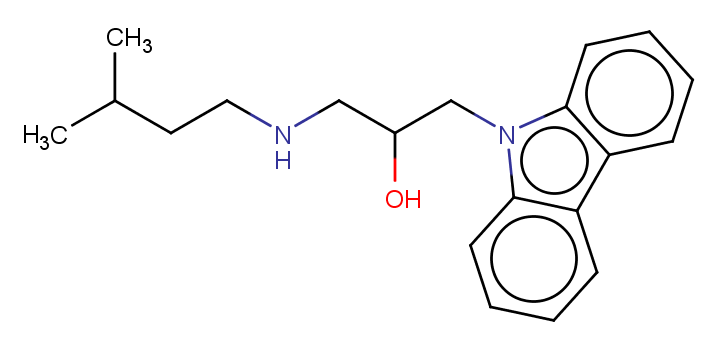

### MMV006820.png

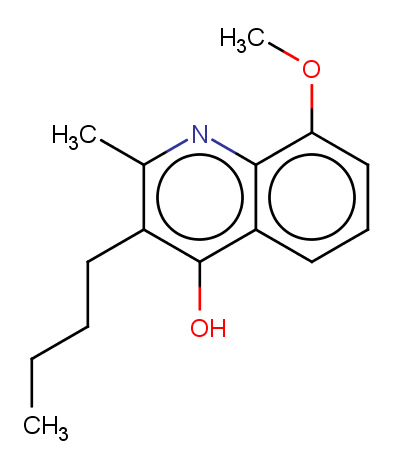

### MMV006825.png

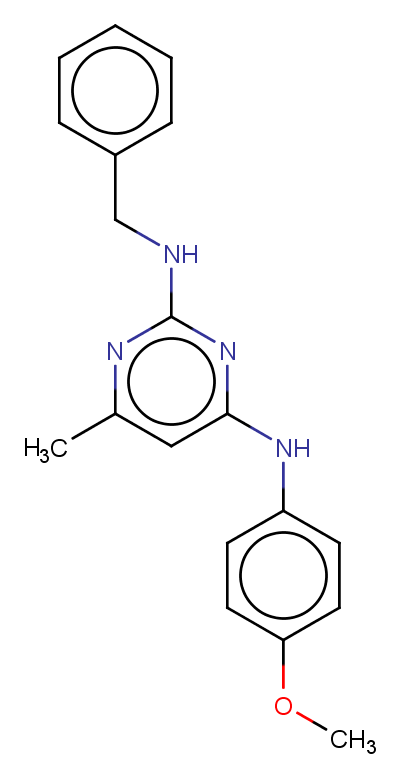

### MMV006861.png

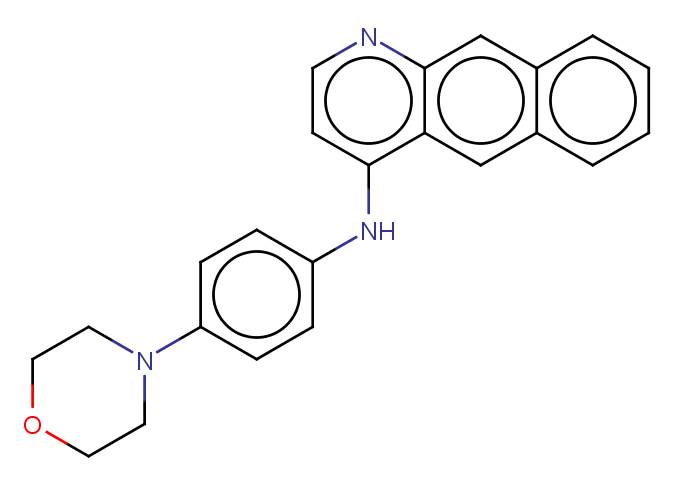

### MMV006882.png

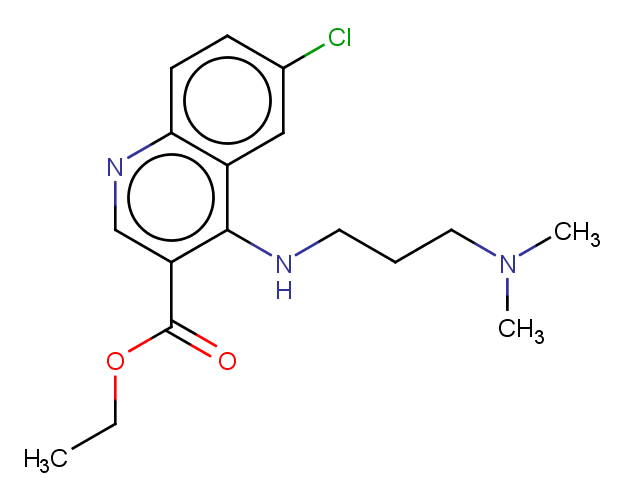

### MMV006913.png

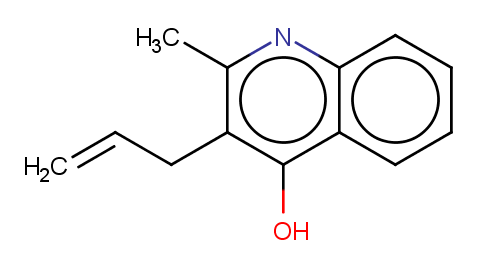

### MMV006937.png

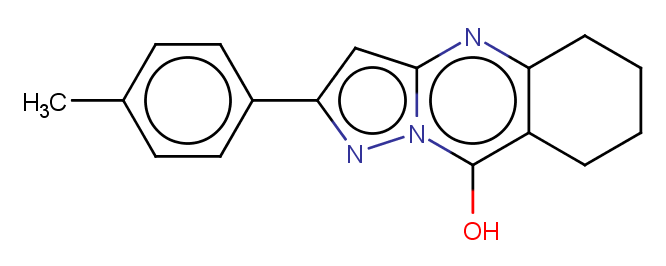

### MMV006962.png

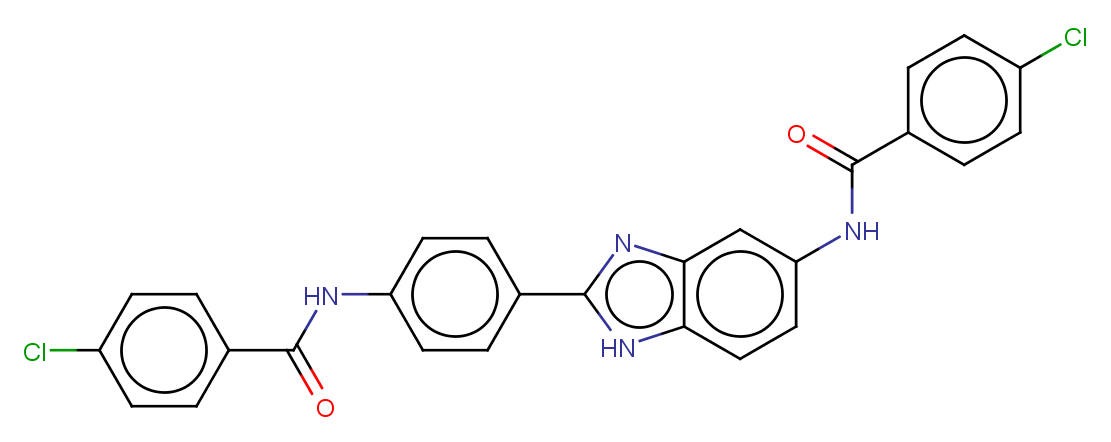

### MMV007020.png

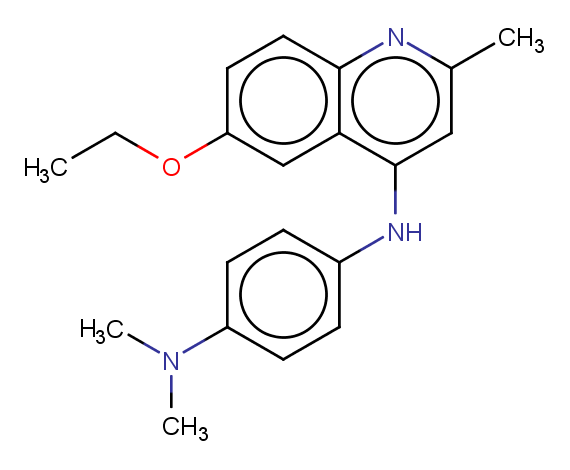

### MMV007041.png

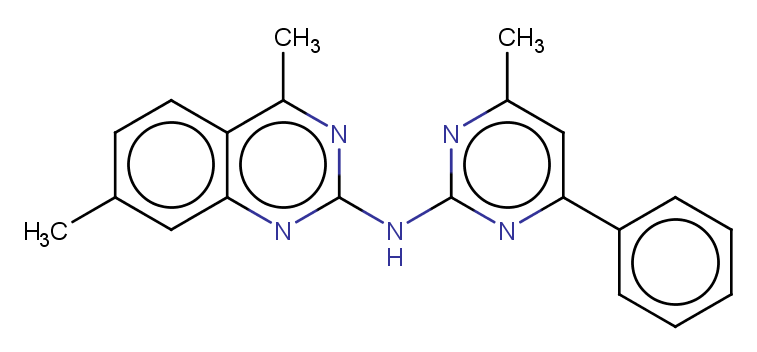

### MMV007092.png

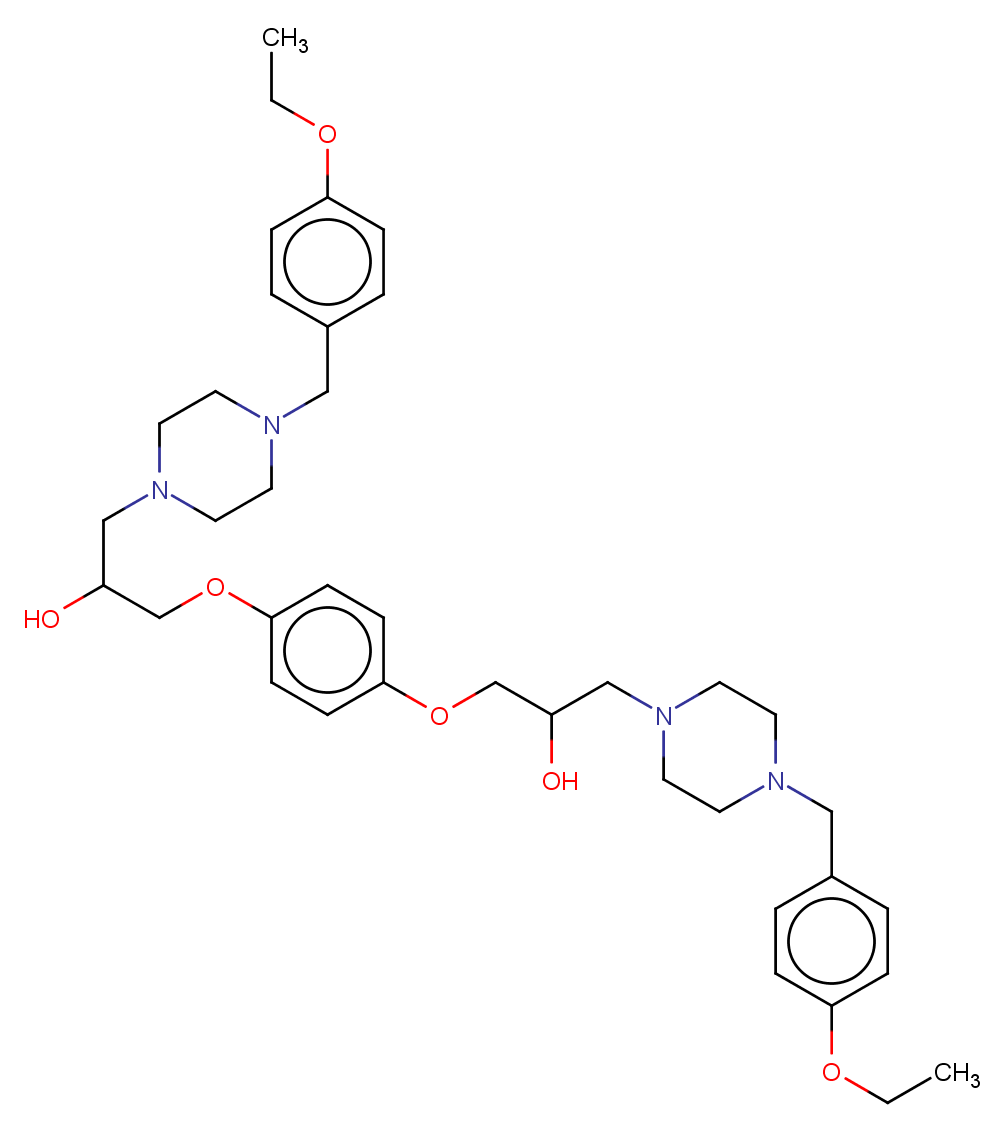

### MMV007113.png

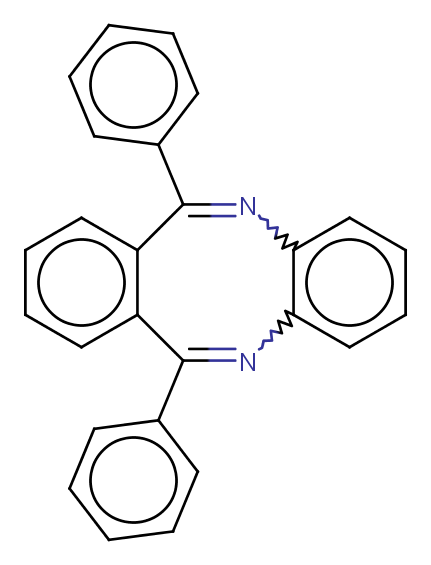

### MMV007116.png

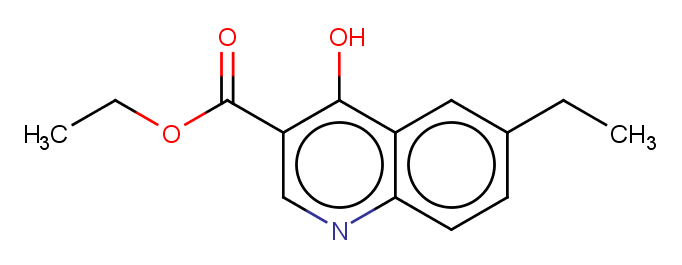

### MMV007127.png

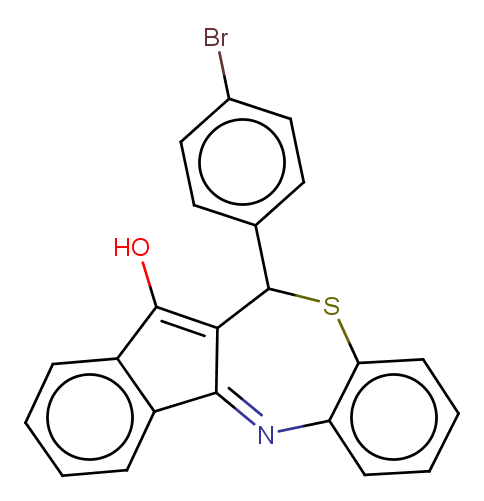

### MMV007160.png

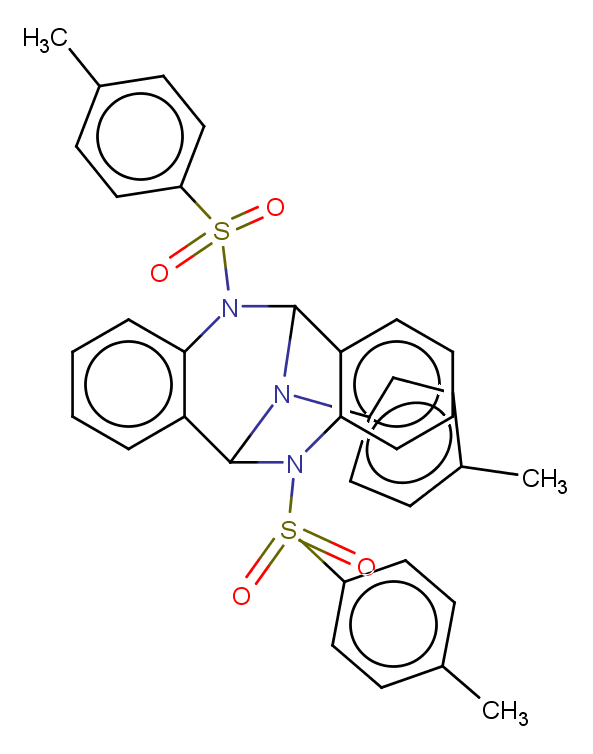

### MMV007181.png

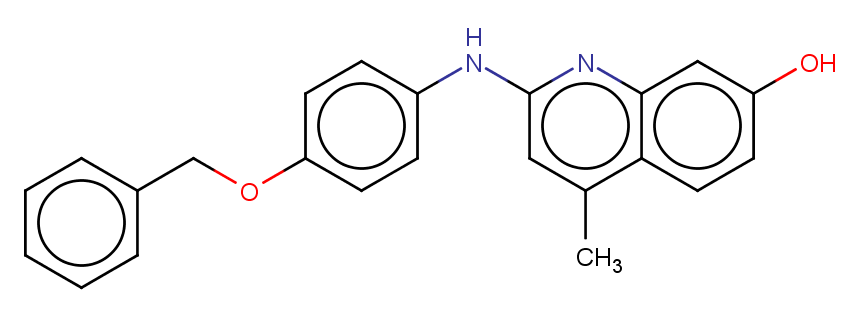

### MMV007199.png

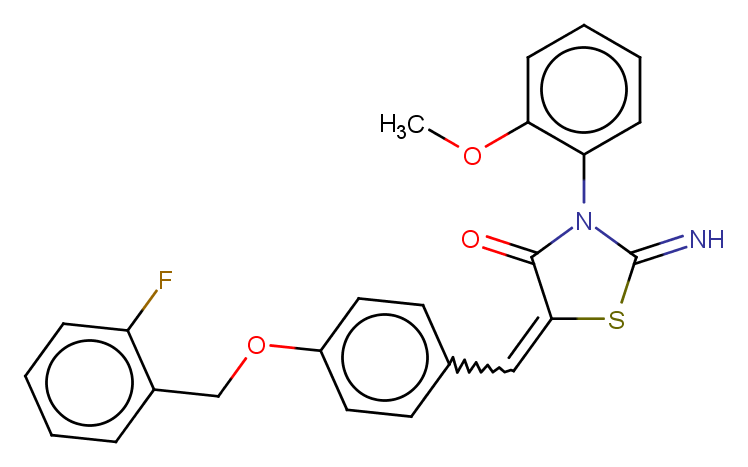

### MMV007208.png

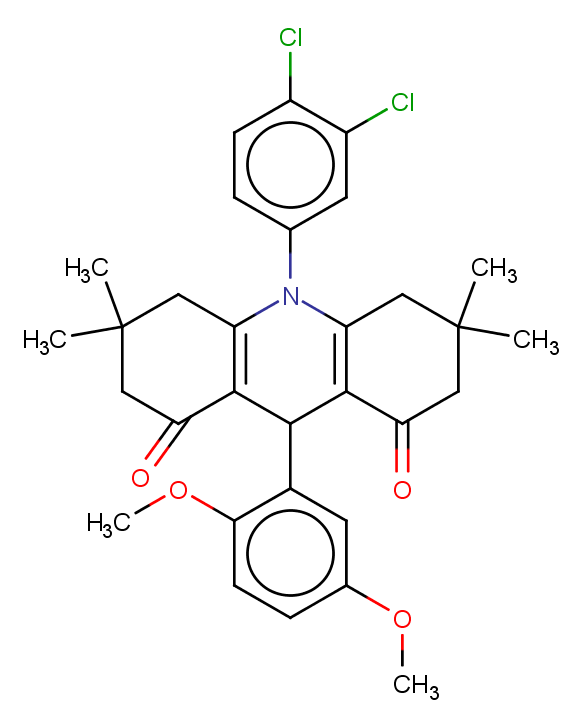

### MMV007224.png

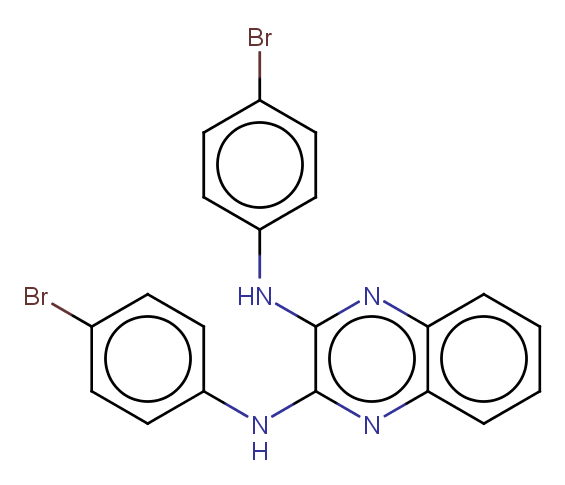

### MMV007228.png

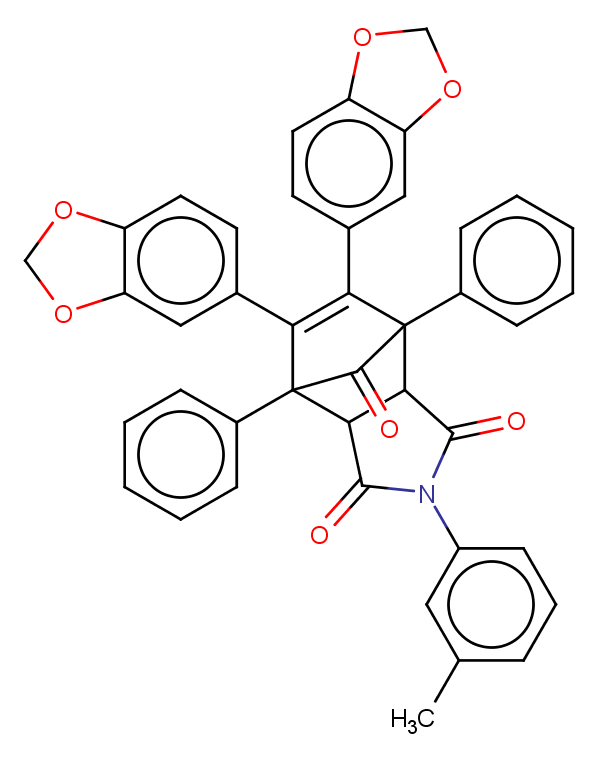

### MMV007273.png

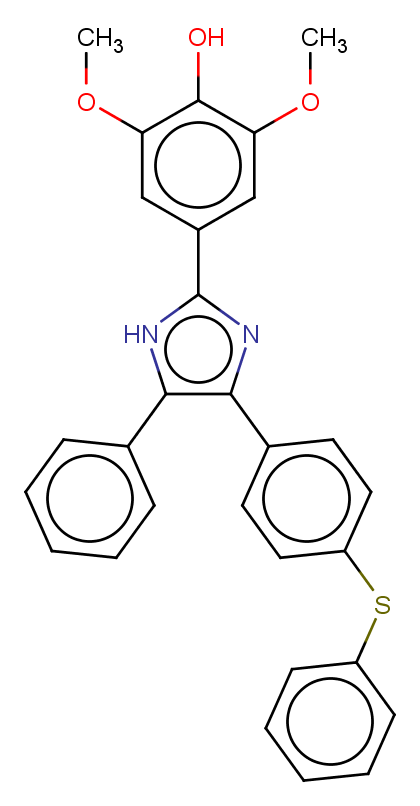

### MMV007275.png

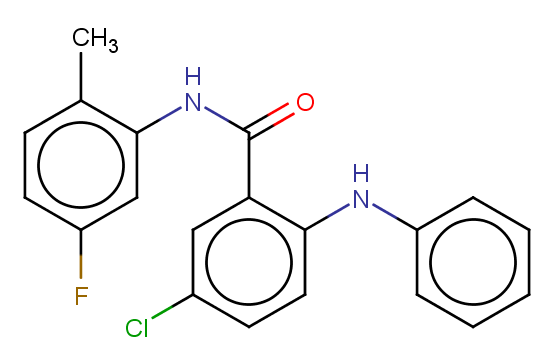

### MMV007285.png

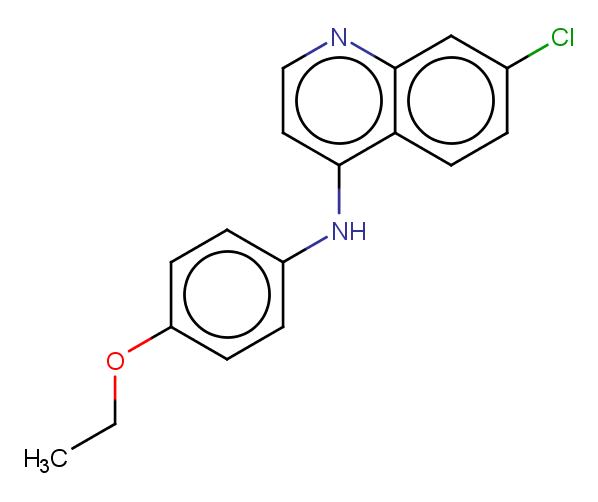

### MMV007363.png

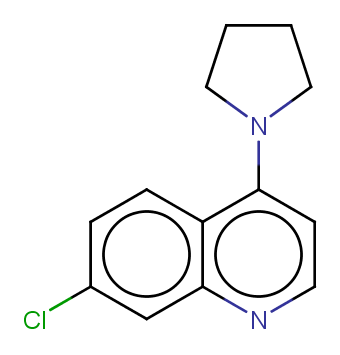
