## Supplementary figures and images for "Refining physico-chemical rules for herbicides using an antimalarial library"

### MMV1050520.png

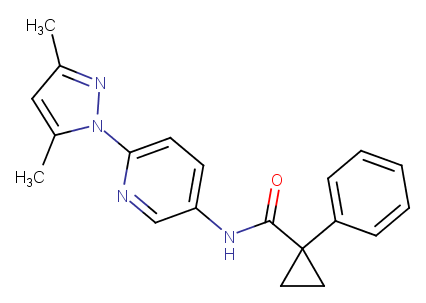

### MMV1051159.png

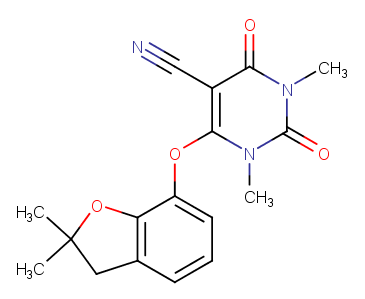

### MMV1052296.png

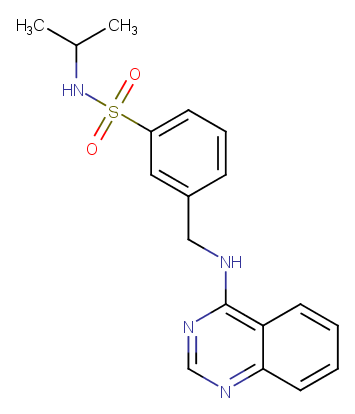

### MMV1055505.png

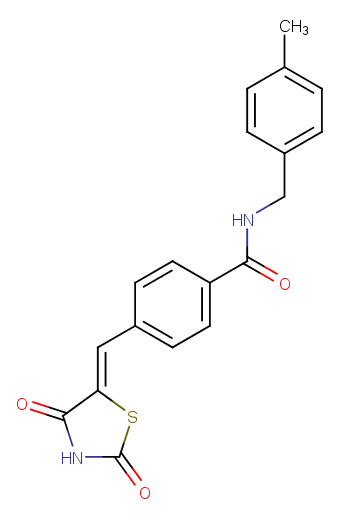

### MMV1057181.png

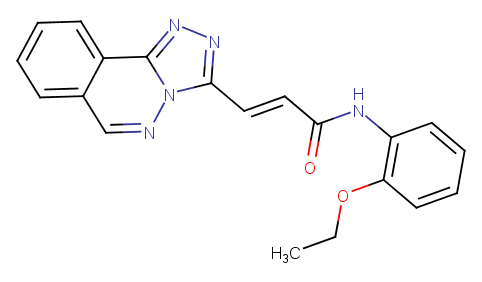

### MMV1058683.png

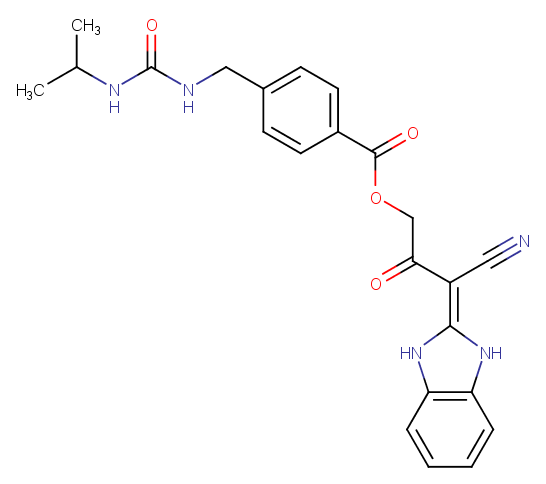

### MMV1065939.png

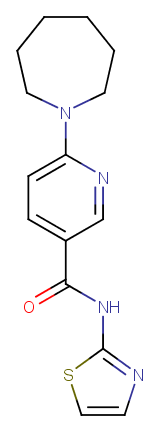

### MMV1068837.png

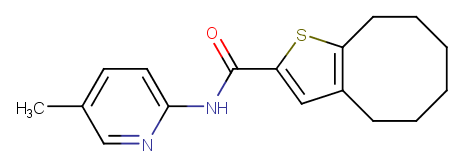

### MMV1068987.png

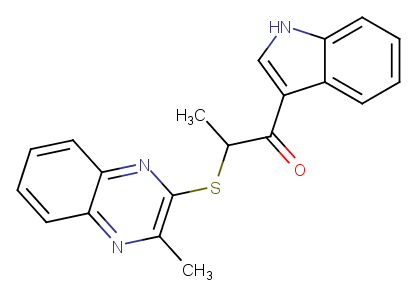

### MMV1069013.png

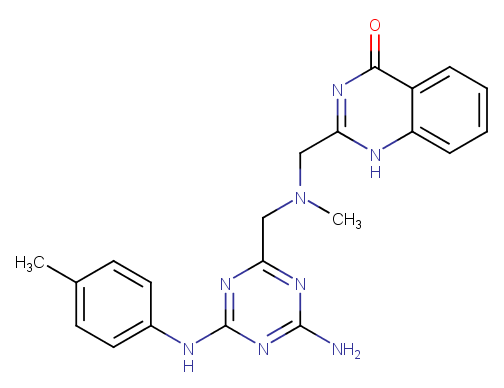

### MMV1069255.png

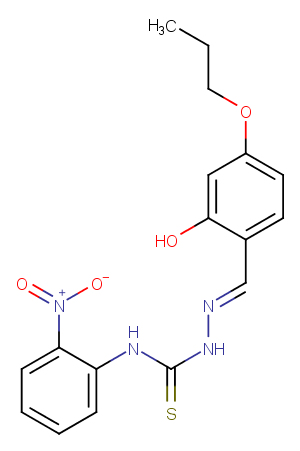

### MMV1069399.png

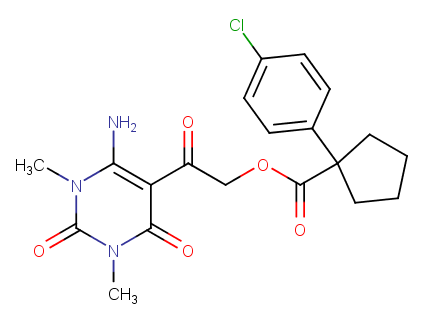

### MMV1069886.png

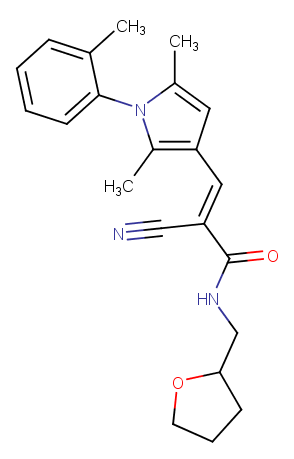

### MMV1070017.png

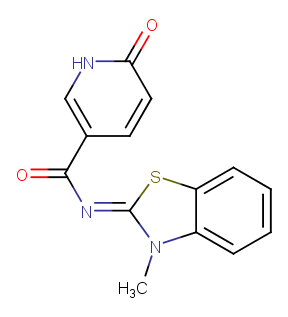

### MMV1070882.png

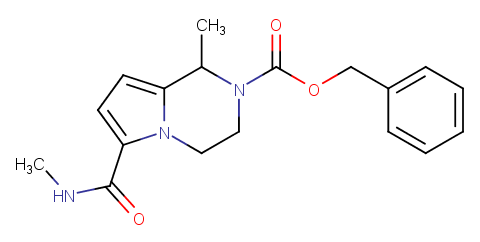

### MMV1071073.png

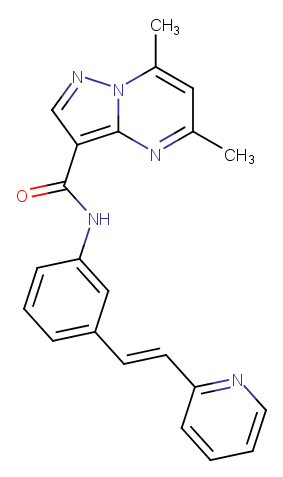

### MMV1071518.png

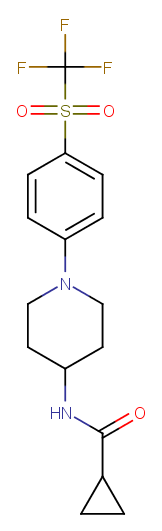

### MMV1071712.png

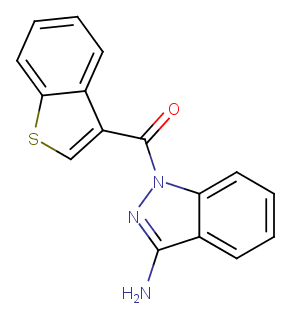

### MMV1071768.png

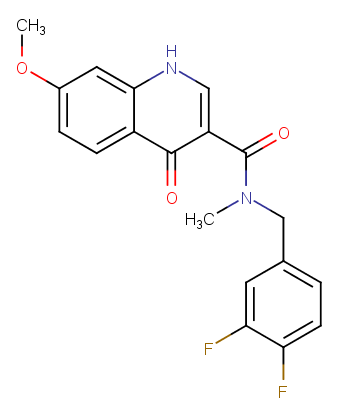

### MMV1072948.png

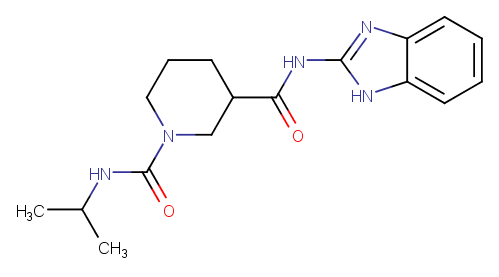

### MMV1075268.png

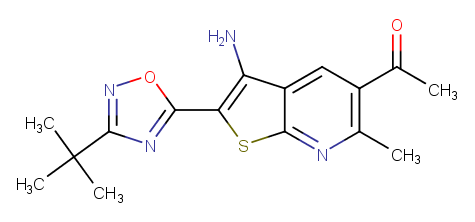

### MMV1076356.png

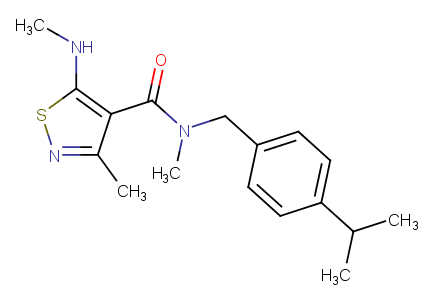

### MMV1078458.png

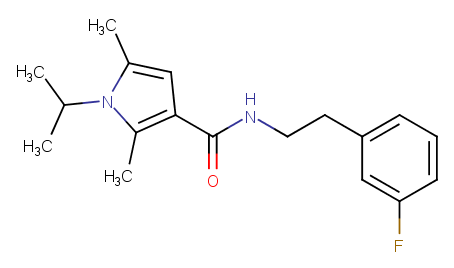

### MMV1079594.png

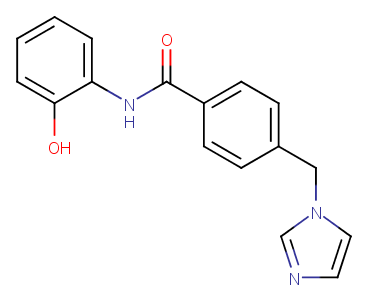

### MMV1081050.png

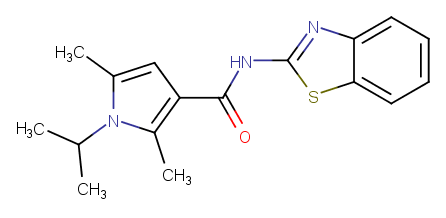

### MMV1081395.png

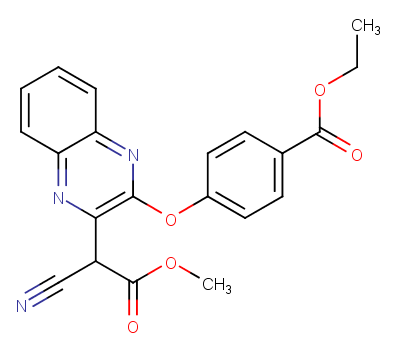

### MMV1083923.png

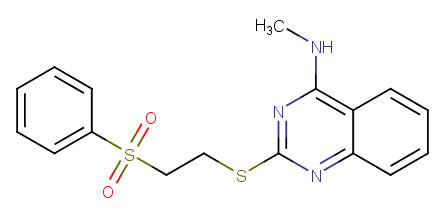

### MMV1084012.png

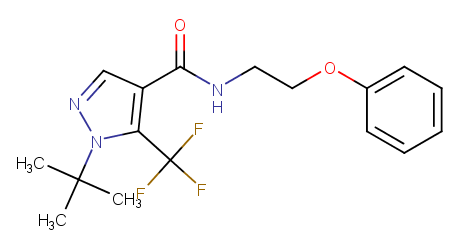

### MMV1084034.png

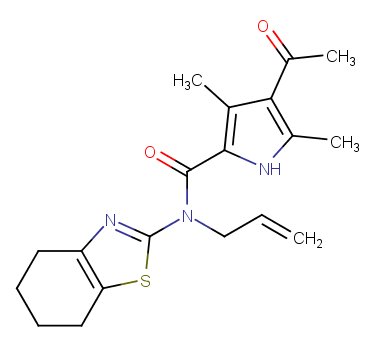

### MMV1084062.png

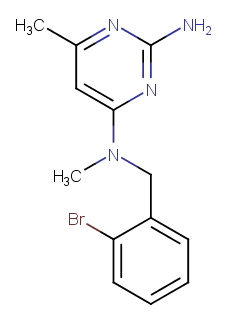

### MMV1084146.png

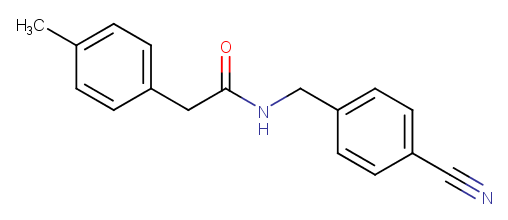

### MMV1085491.png

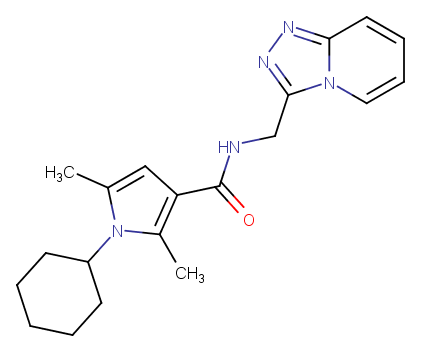

### MMV1086039.png

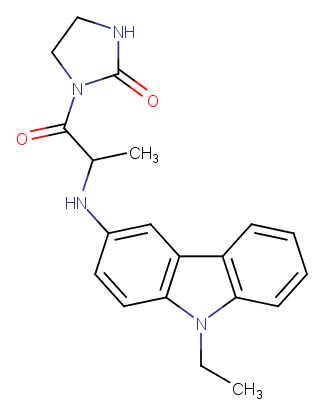

### MMV1086262.png

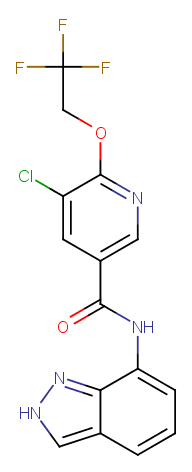

### MMV1086523.png

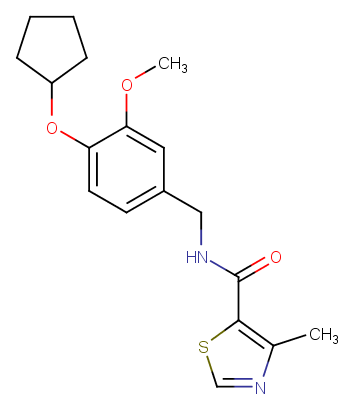

### MMV1087037.png

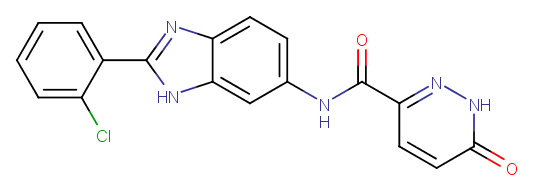

### MMV1087133.png

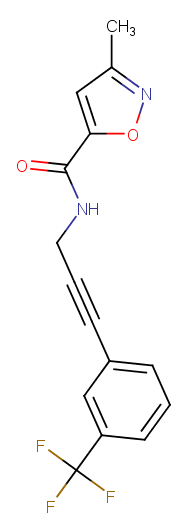

### MMV1088447.png

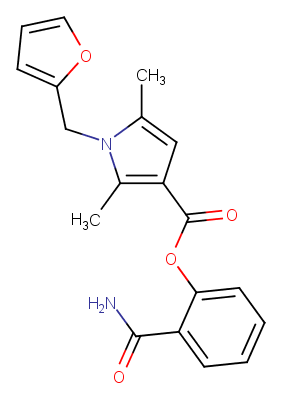

### MMV1088595.png

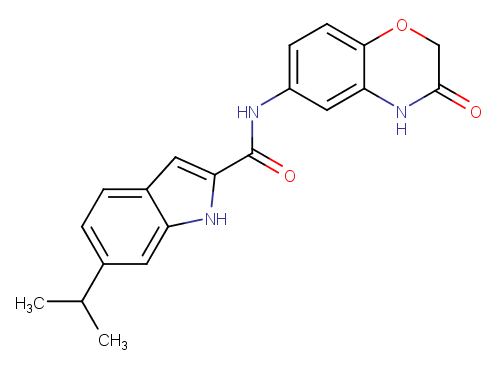

### MMV1088860.png

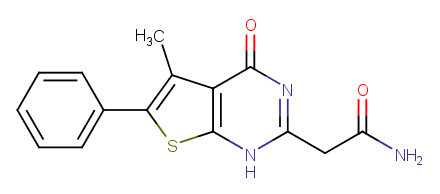

### MMV1090336.png

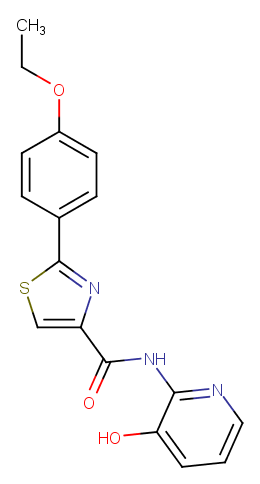

### MMV1090408.png

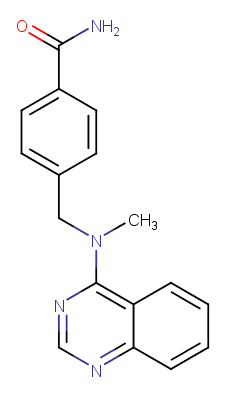

### MMV1090686.png

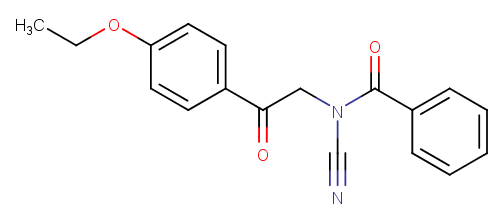

### MMV1091186.png

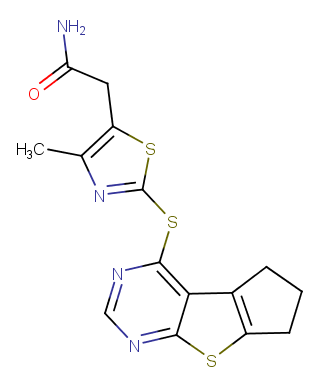

### MMV1091654.png

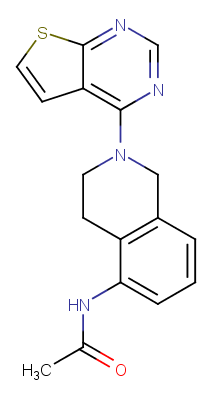

### MMV1092395.png

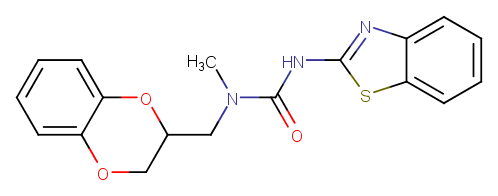

### MMV1092454.png

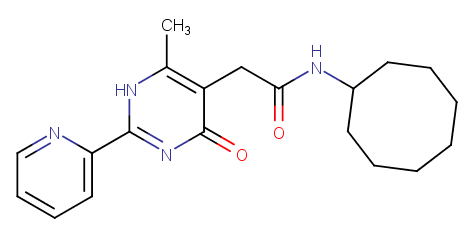

### MMV1092500.png

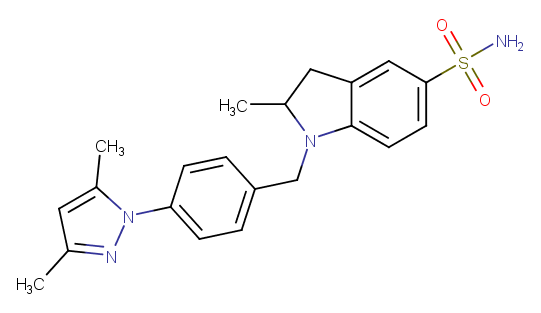

### MMV1092612.png

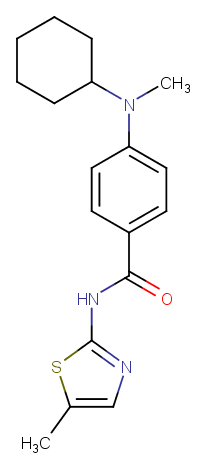

### MMV1092949.png

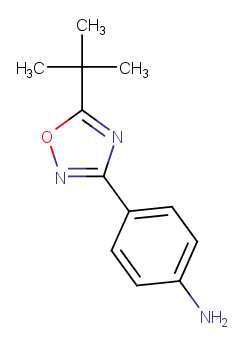

### MMV1097577.png

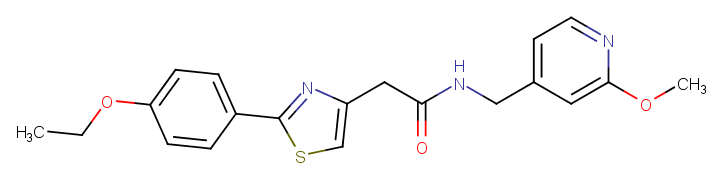

### MMV1097781.png

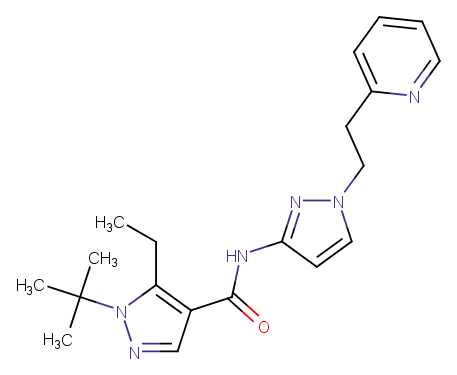

### MMV1097981.png

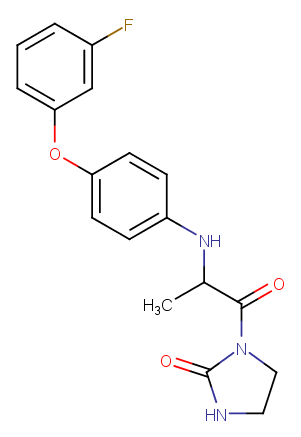

### MMV1098418.png

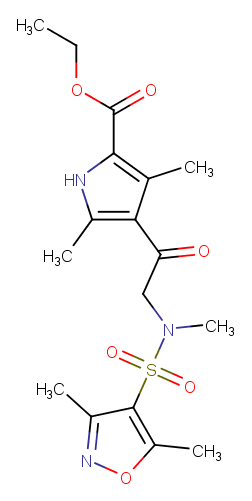

### MMV1098839.png

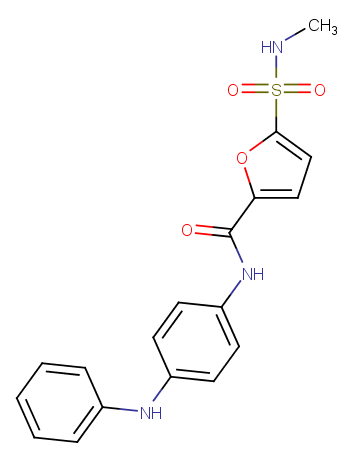

### MMV1100542.png

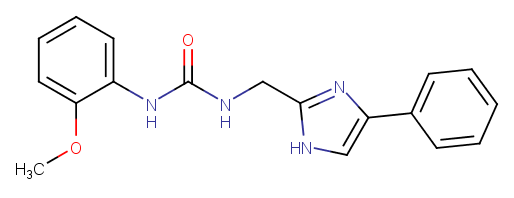

### MMV1100803.png

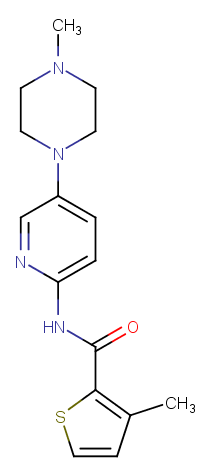

### MMV1101135.png

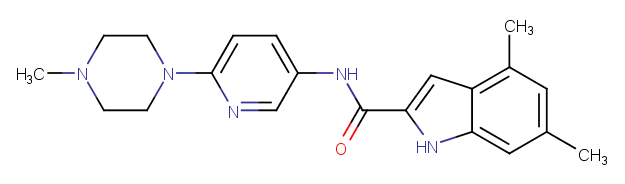

### MMV1102231.png

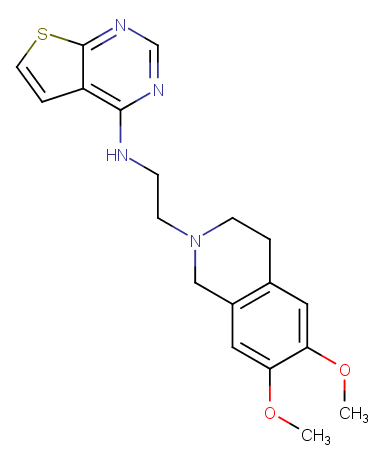

### MMV1102398.png

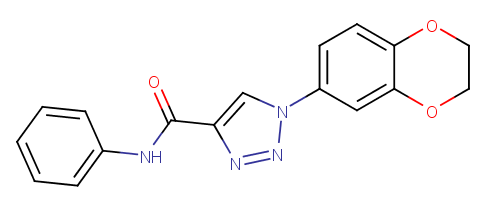

### MMV1102399.png

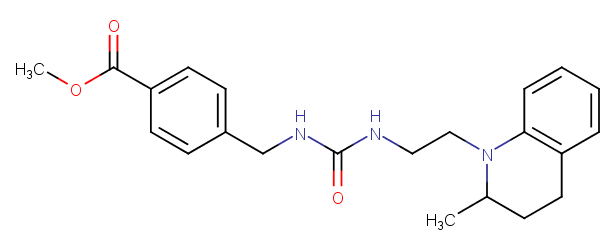

### MMV1102419.png

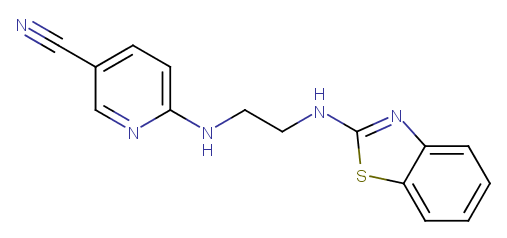

### MMV1102629.png

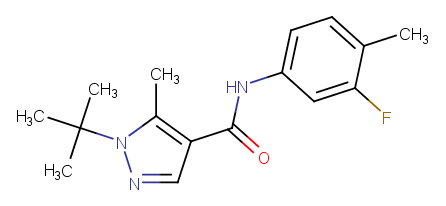

### MMV1102713.png

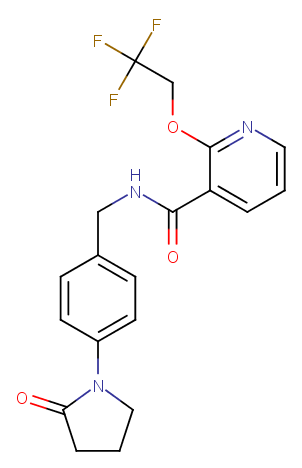

### MMV1102829.png

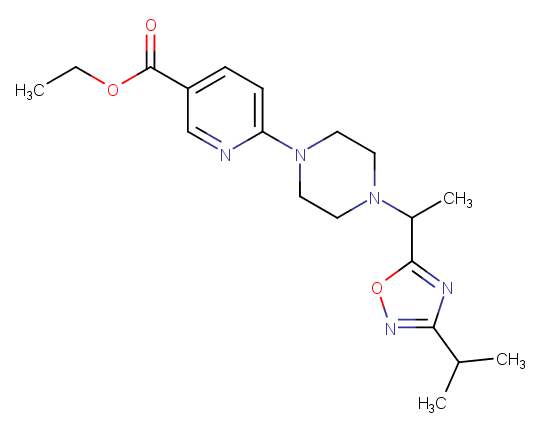

### MMV1102996.png

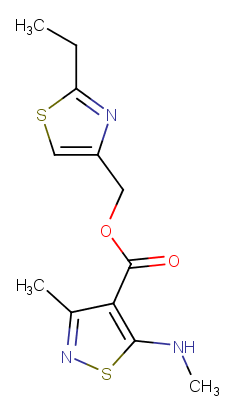

### MMV1103183.png

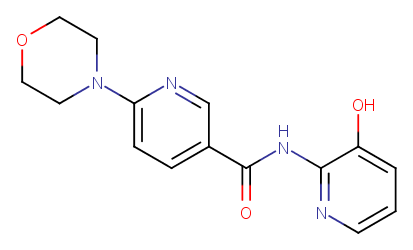

### MMV1103200.png

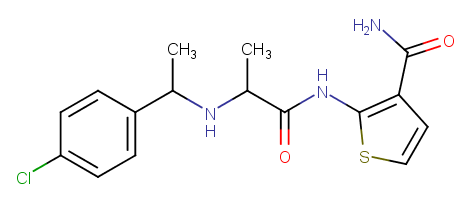

### MMV1103291.png

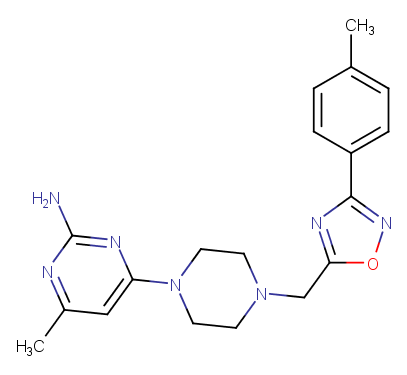

### MMV1103645.png

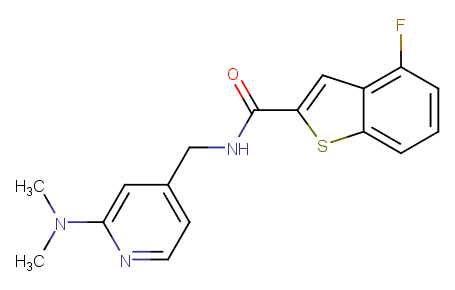

### MMV1104039.png

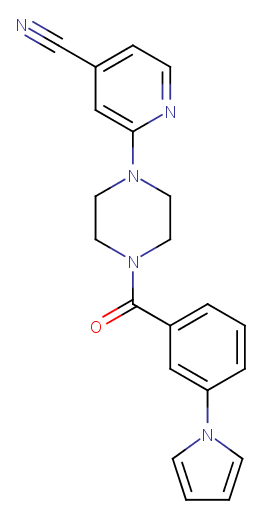

### MMV1104109.png

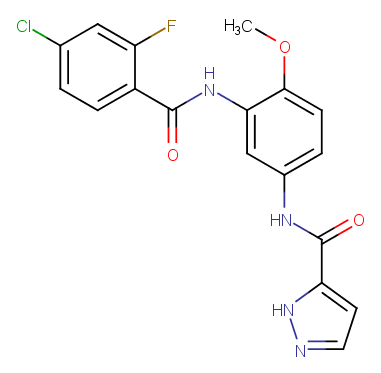

### MMV1151140.png

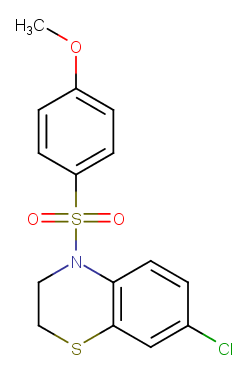

### MMV1162938.png

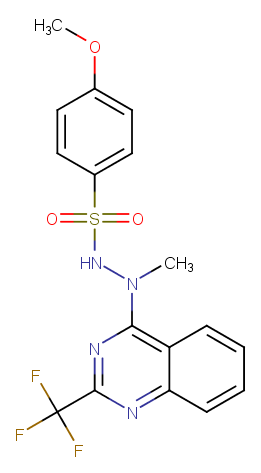

### MMV1164831.png

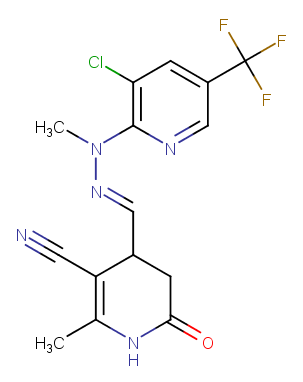

### MMV1167451.png

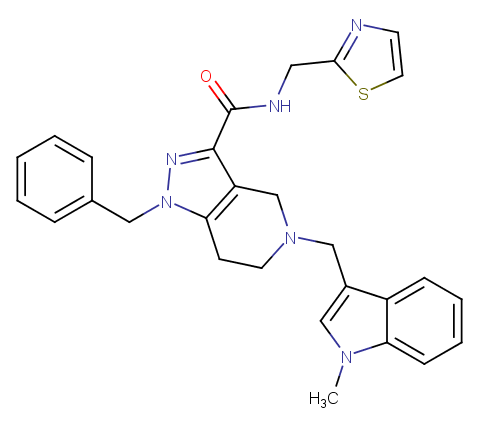

### MMV1167992.png

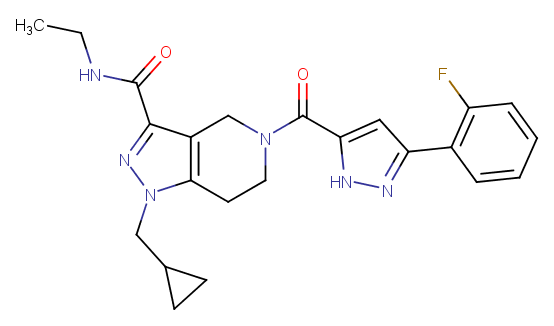

### MMV1168594.png

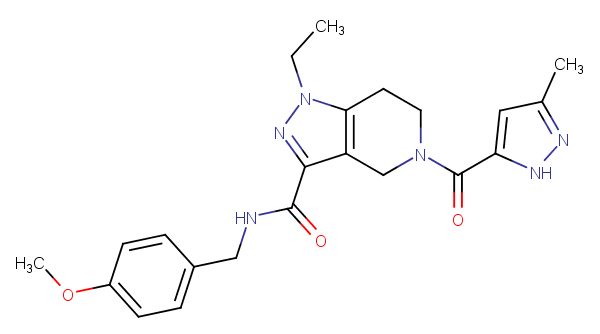

### MMV1168781.png

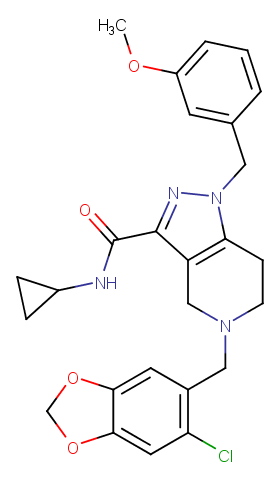

### MMV1168945.png

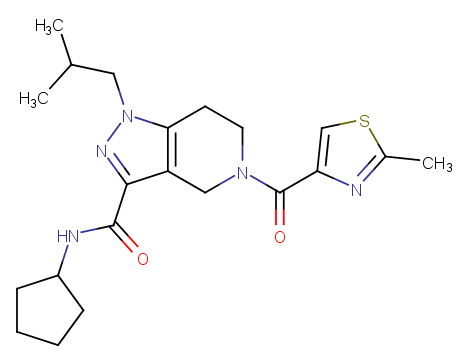

### MMV1169088.png

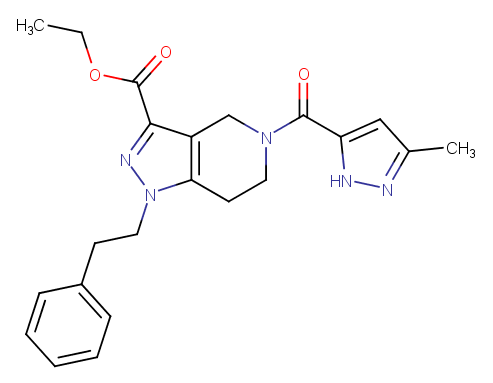

### MMV1169220.png

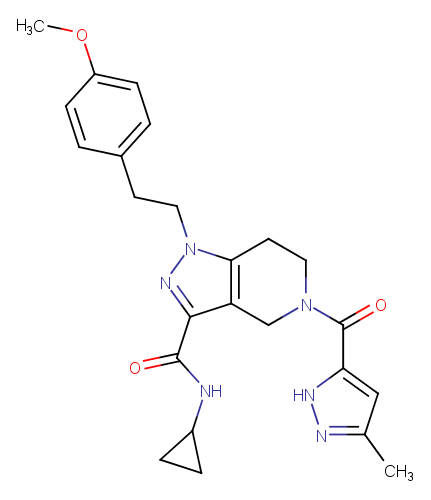

### MMV1169752.png

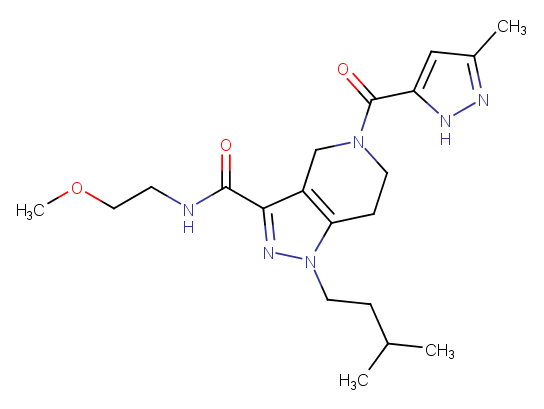

### MMV1169802.png

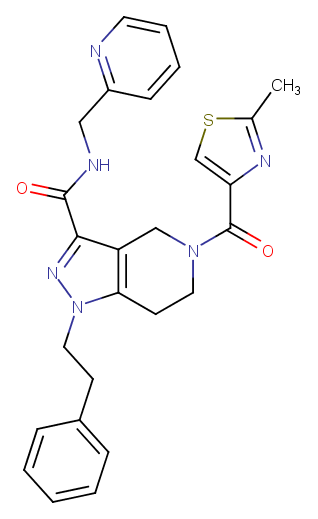

### MMV1170432.png

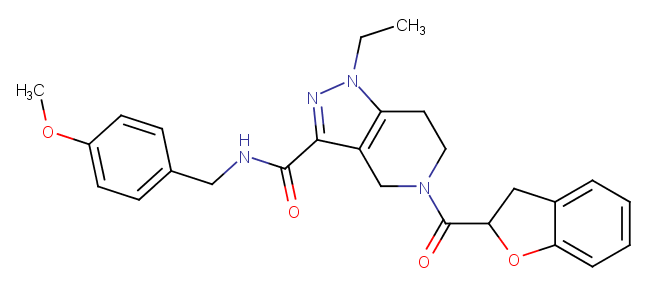

### MMV1171032.png

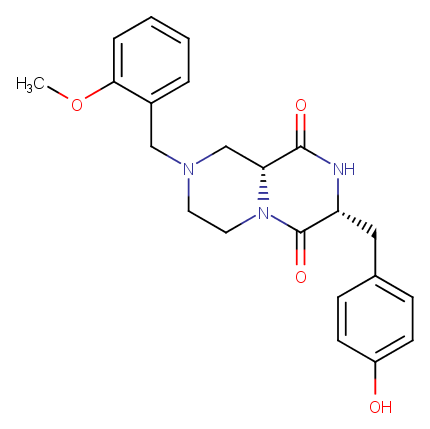

### MMV1174026.png

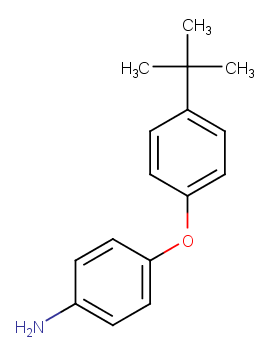

### MMV1174438.png

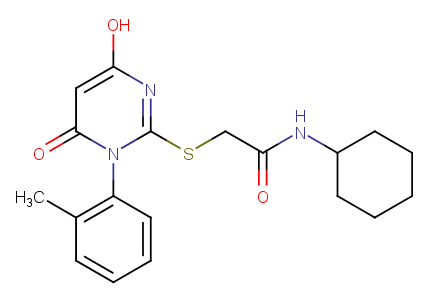

### MMV1174825.png

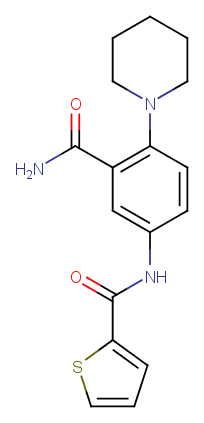

### MMV1174946.png

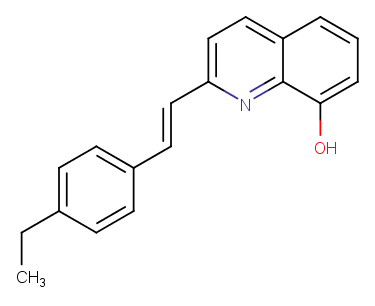

### MMV1175000.png

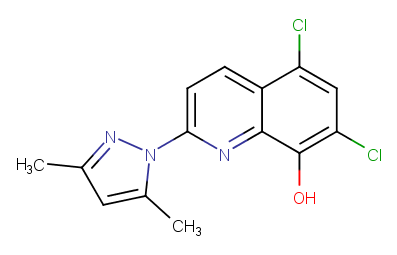

### MMV1175132.png

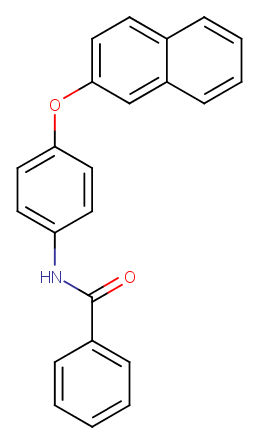

### MMV1175293.png

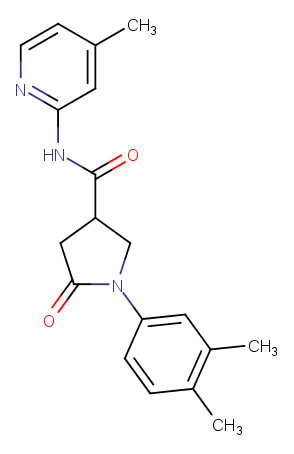

### MMV1175522.png

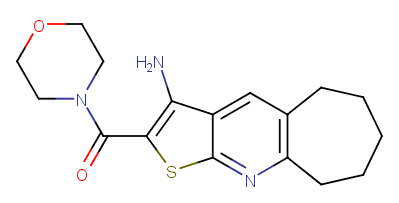

### MMV1177519.png

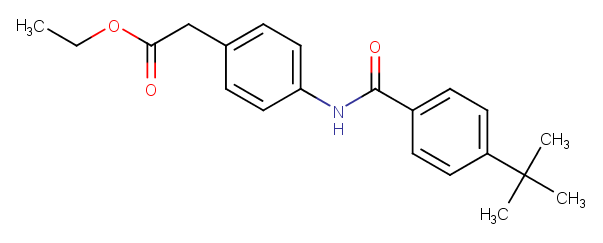

### MMV1178683.png

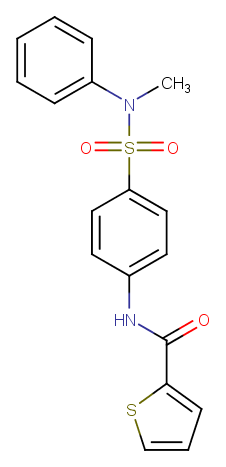

### MMV1179816.png

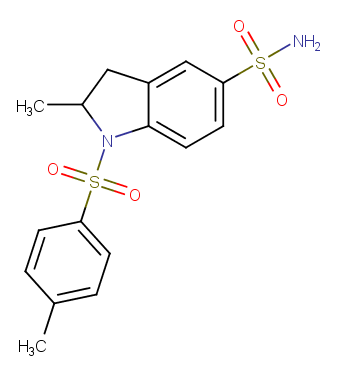
