## Supplementary figures and images for "Refining physico-chemical rules for herbicides using an antimalarial library"

### MMV007742.png

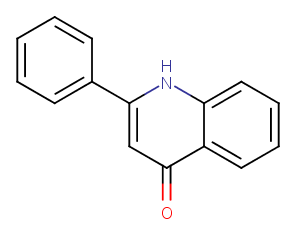

### MMV011573.png

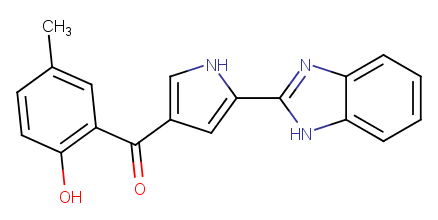

### MMV011772.png

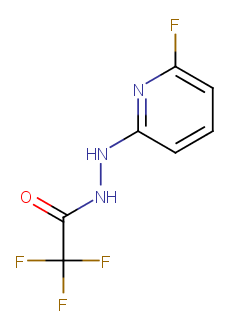

### MMV020795.png

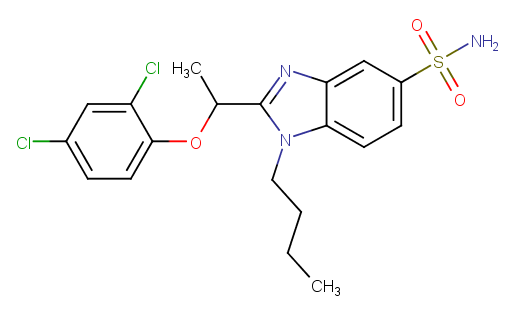

### MMV020978.png

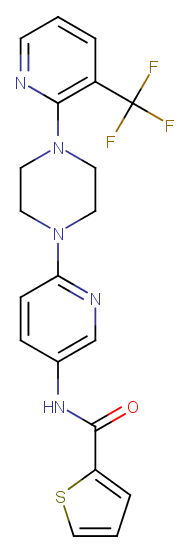

### MMV035664.png

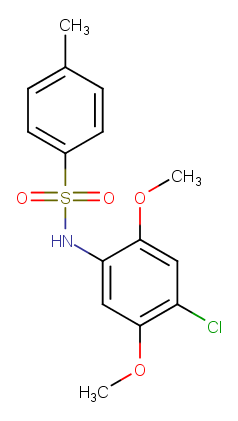

### MMV039455.png

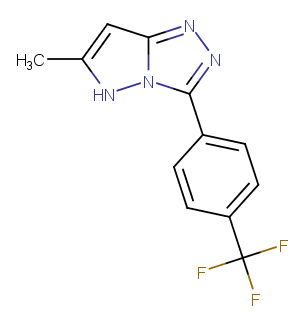

### MMV043084.png

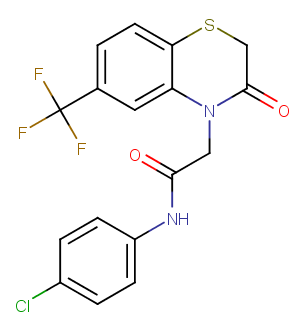

### MMV044470.png

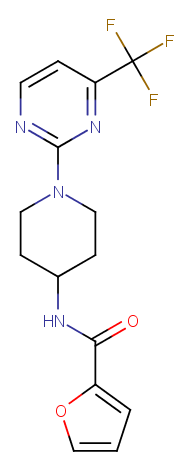

### MMV046844.png

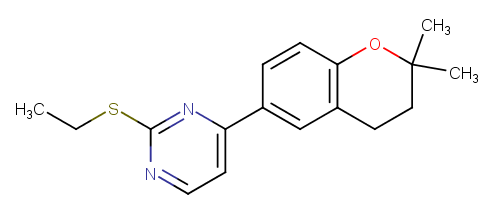

### MMV046853.png

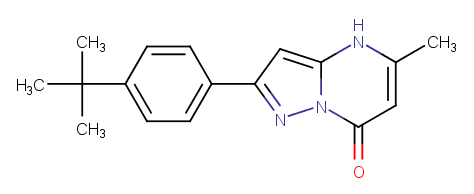

### MMV047179.png

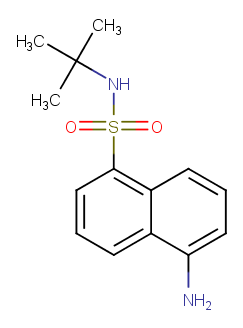

### MMV047558.png

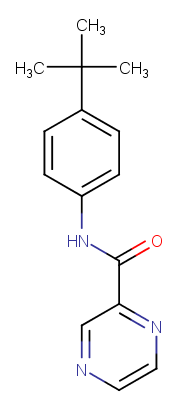

### MMV048884.png

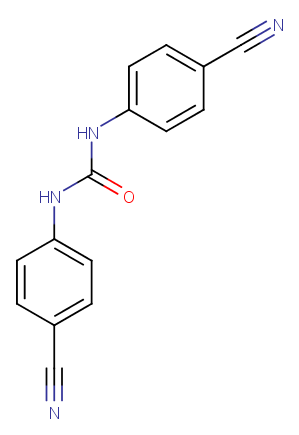

### MMV048973.png

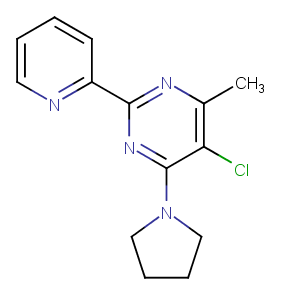

### MMV049229.png

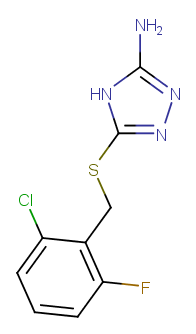

### MMV049749.png

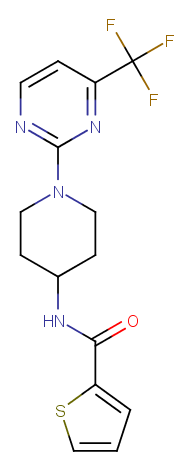

### MMV051436.png

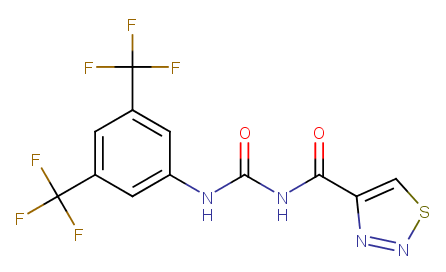

### MMV061540.png

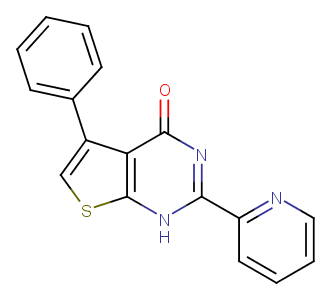

### MMV062850.png

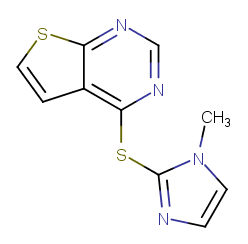

### MMV064286.png

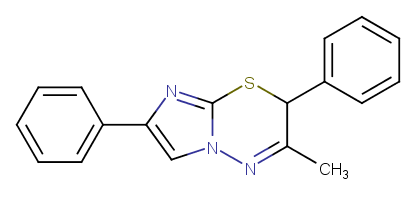

### MMV065310.png

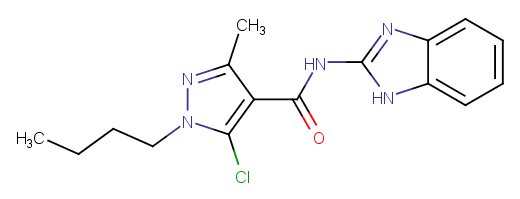

### MMV066171.png

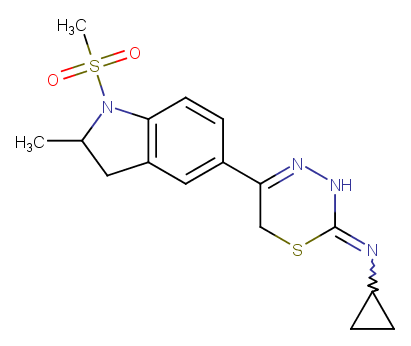

### MMV066594.png

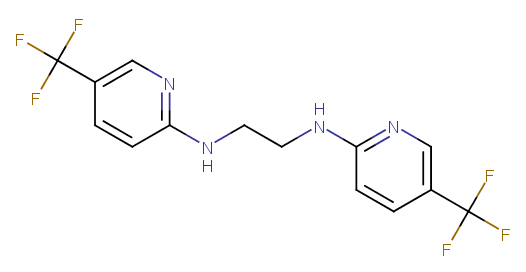

### MMV077871.png

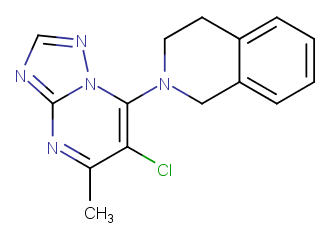

### MMV077993.png

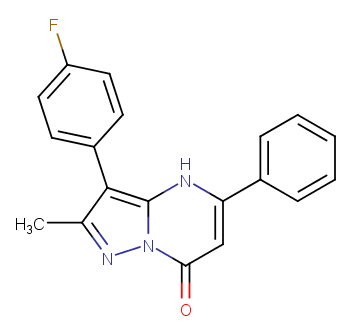

### MMV098914.png

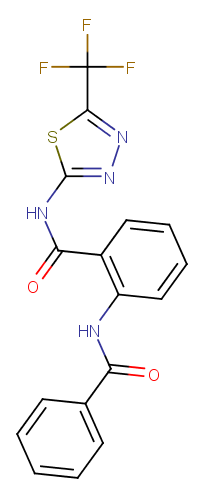

### MMV1005663.png

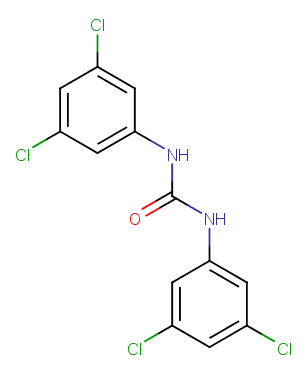

### MMV1005800.png

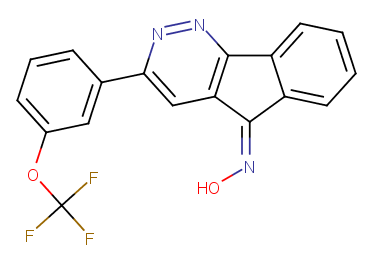

### MMV1006936.png

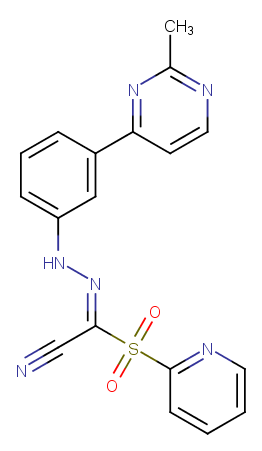

### MMV1006984.png

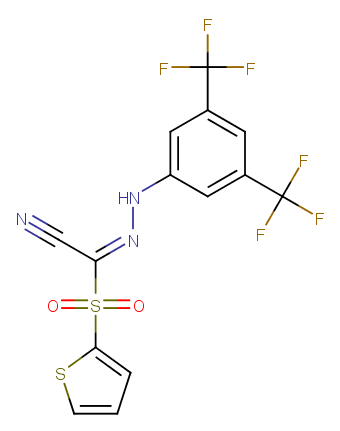

### MMV1007245.png

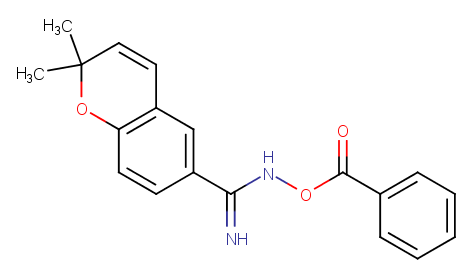

### MMV1008025.png

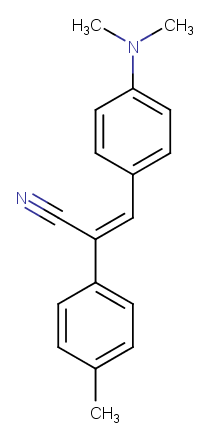

### MMV1008960.png

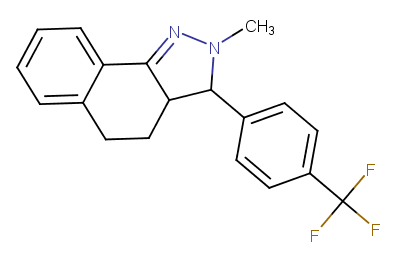

### MMV1009786.png

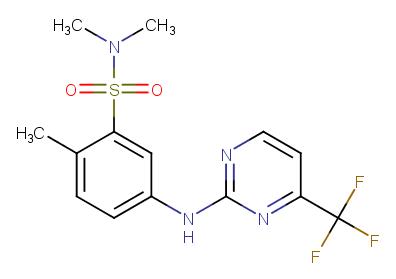

### MMV1009790.png

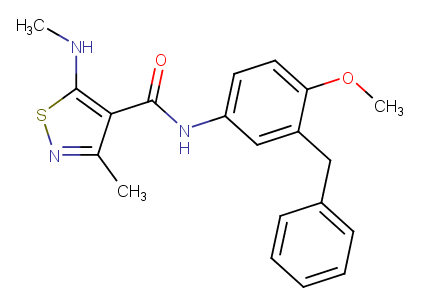

### MMV1009808.png

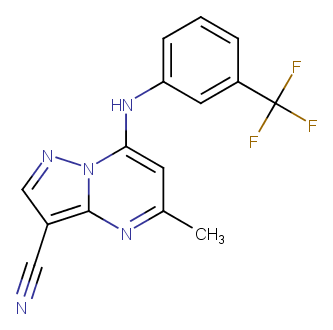

### MMV1009975.png

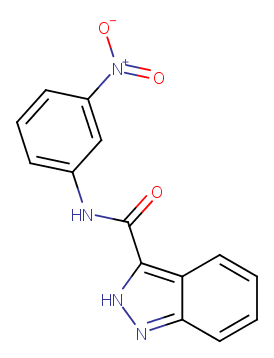

### MMV1010032.png

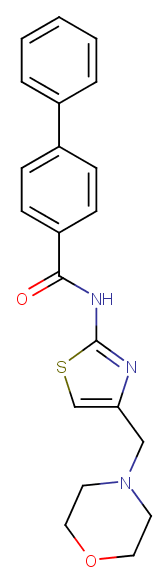

### MMV1010040.png

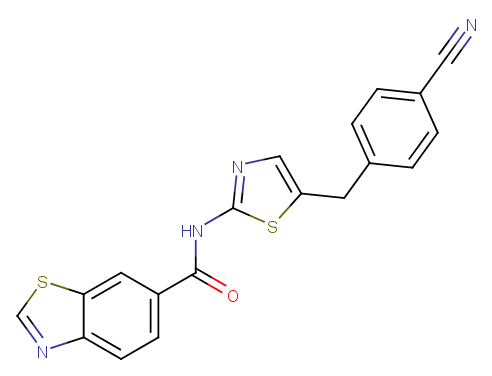

### MMV1010067.png

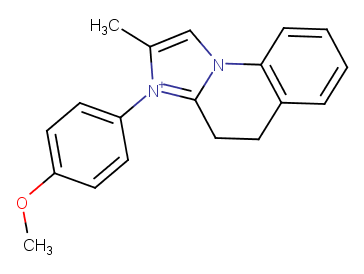

### MMV1010101.png

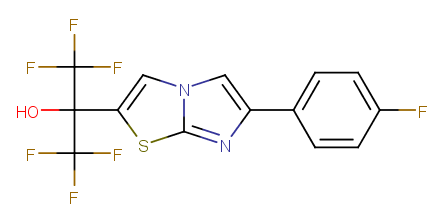

### MMV1010248.png

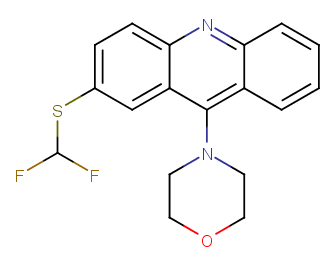

### MMV1010301.png

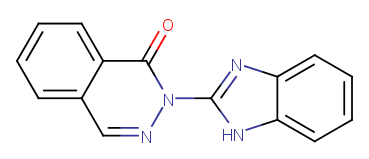

### MMV1010683.png

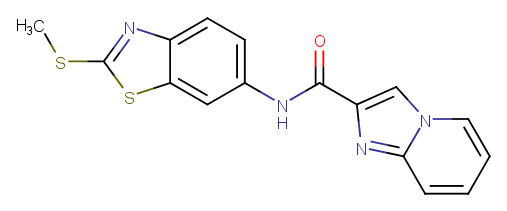

### MMV1013204.png

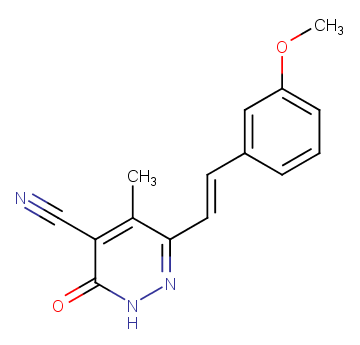

### MMV1013651.png

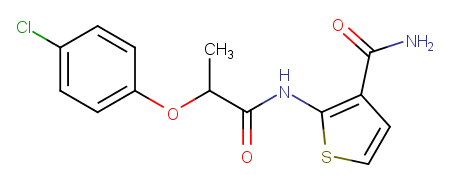

### MMV1013666.png

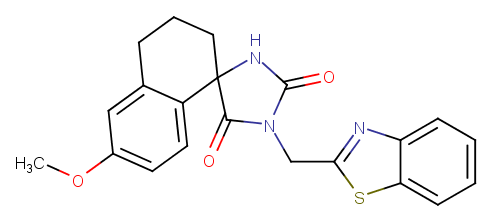

### MMV1013788.png

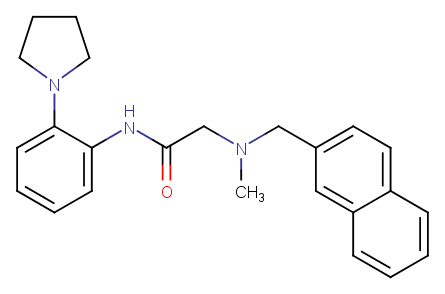

### MMV1014090.png

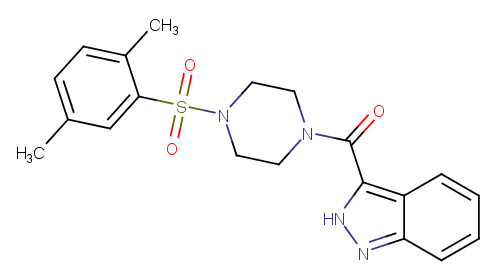

### MMV1014461.png

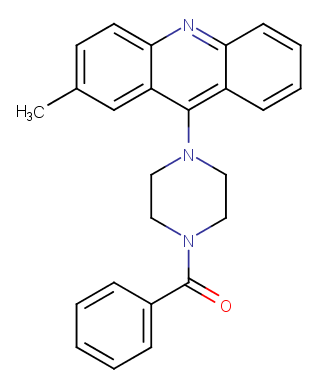

### MMV1014540.png

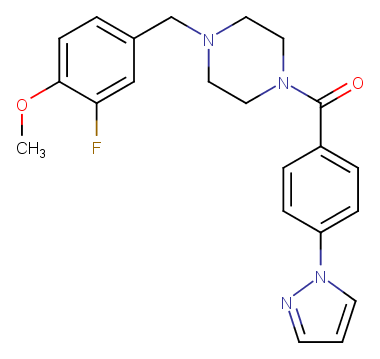

### MMV1014559.png

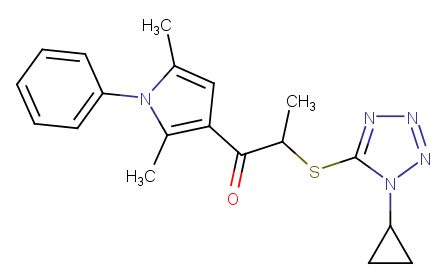

### MMV1014895.png

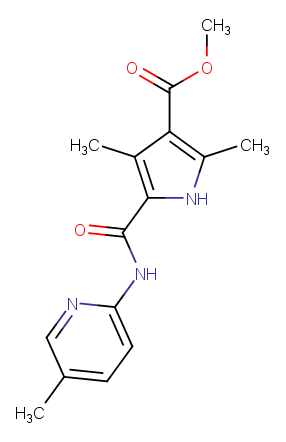

### MMV1015011.png

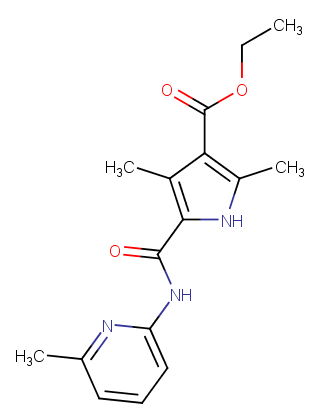

### MMV1021203.png

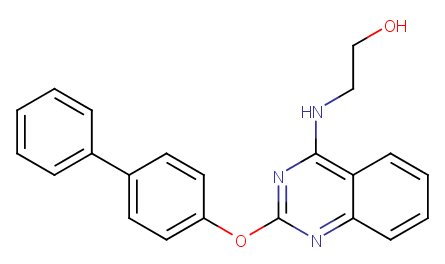

### MMV1025906.png

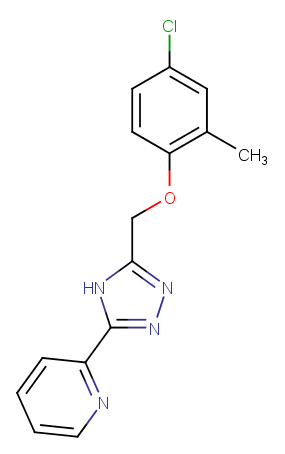

### MMV1026923.png

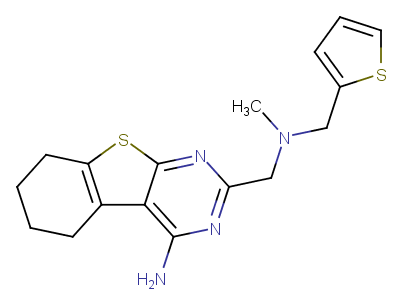

### MMV1028379.png

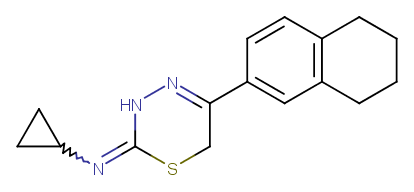

### MMV1028655.png

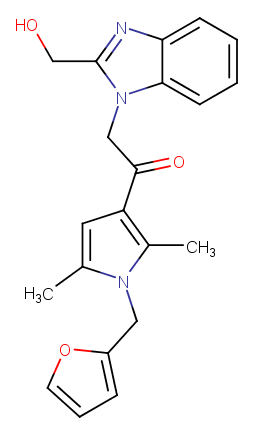

### MMV1028949.png

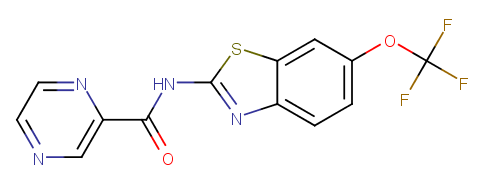

### MMV1029203.png

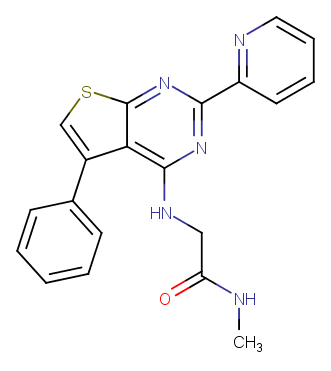

### MMV1029300.png

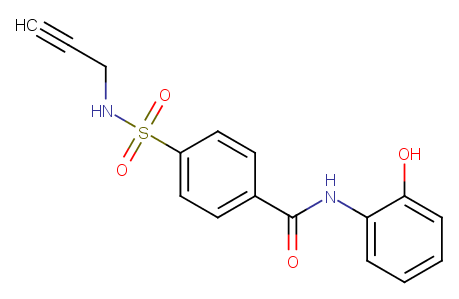

### MMV1029656.png

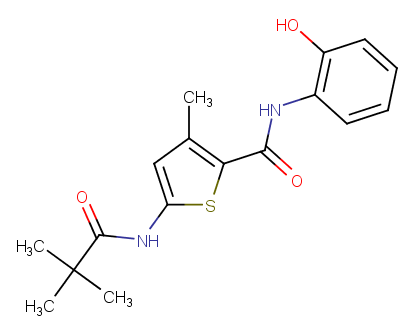

### MMV1030182.png

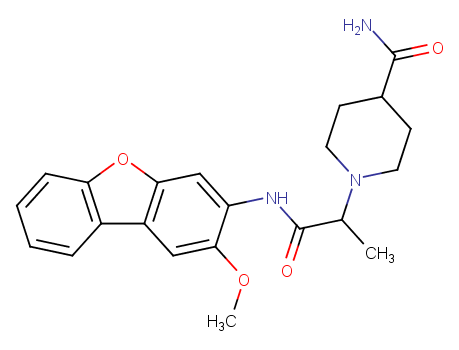

### MMV1030932.png

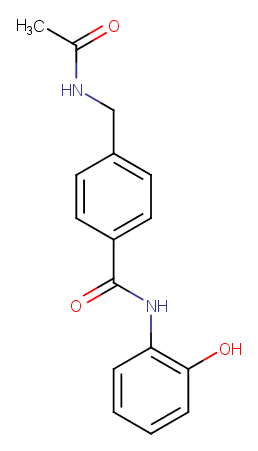

### MMV1031421.png

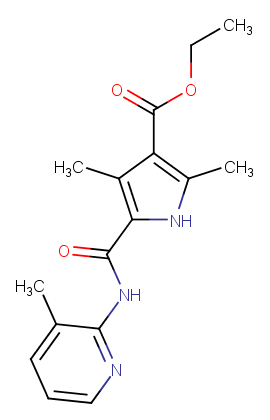

### MMV1031775.png

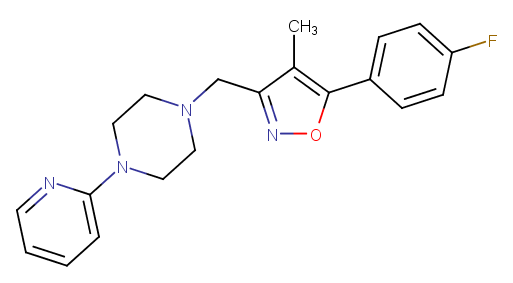

### MMV1034288.png

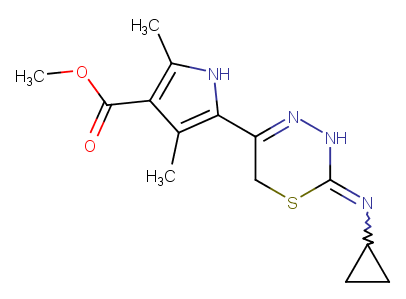

### MMV1034579.png

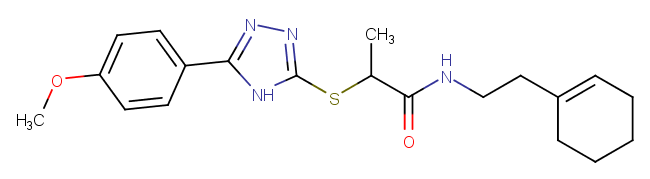

### MMV1034814.png

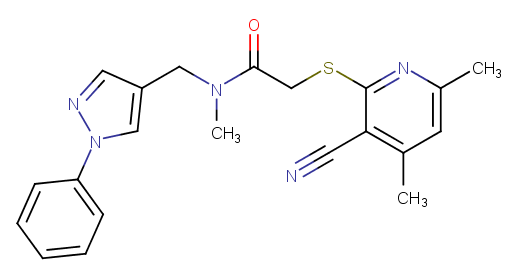

### MMV1035644.png

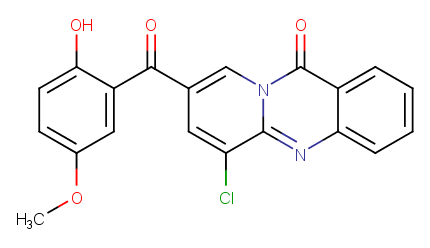

### MMV1036321.png

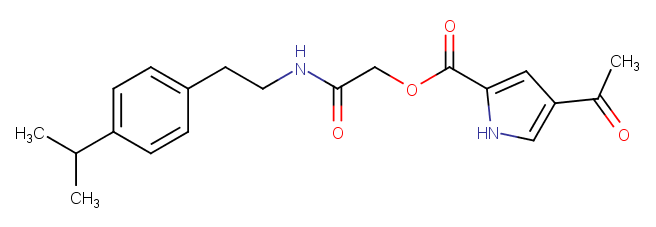

### MMV1038036.png

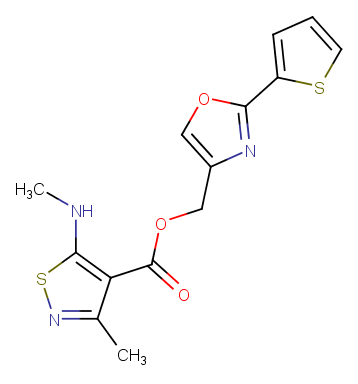

### MMV1038147.png

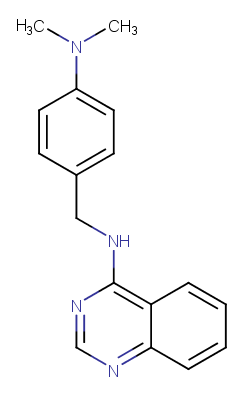

### MMV1042747.png

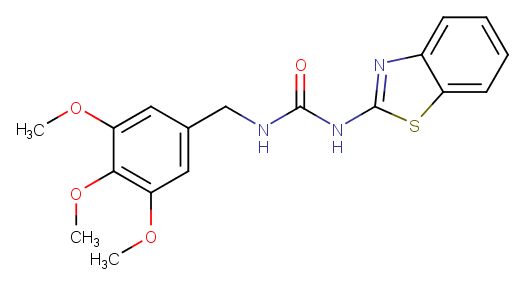

### MMV1042847.png

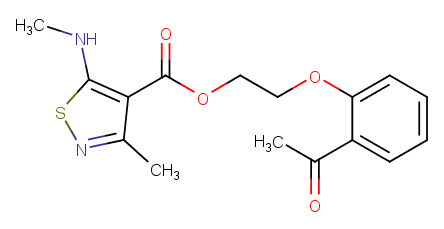

### MMV1042856.png

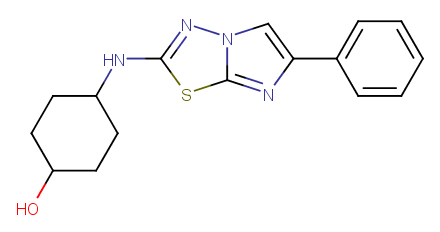

### MMV1042923.png

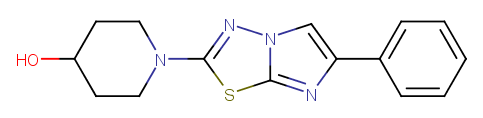

### MMV1042937.png

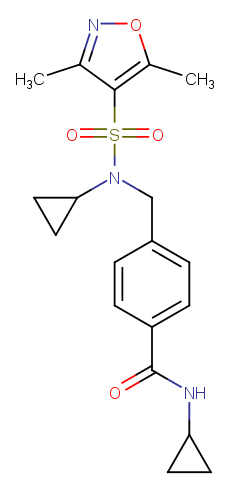

### MMV1043464.png

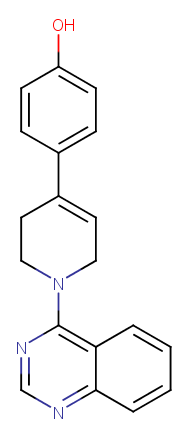

### MMV1043514.png

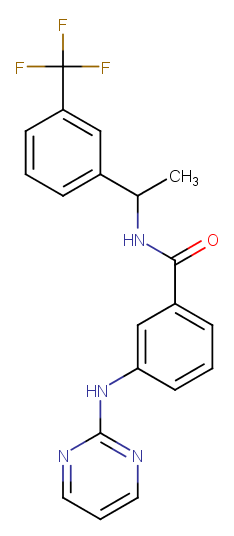

### MMV1043680.png

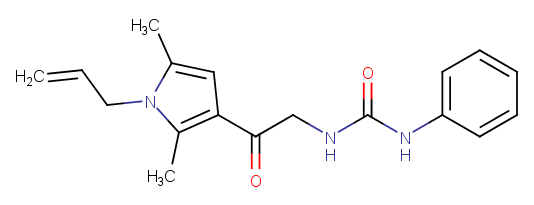

### MMV1043905.png

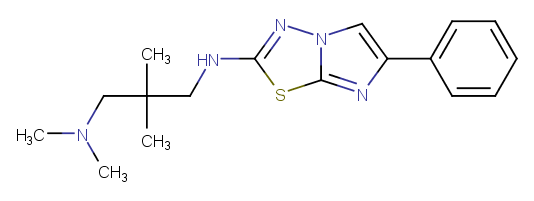

### MMV1044457.png

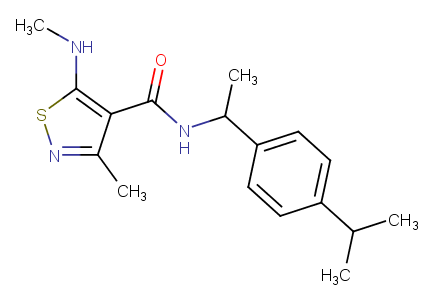

### MMV1045248.png

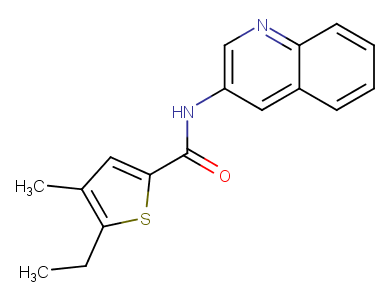

### MMV1045838.png

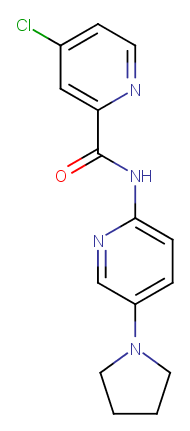

### MMV1046173.png

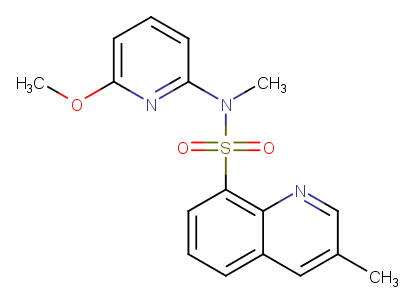

### MMV1046672.png

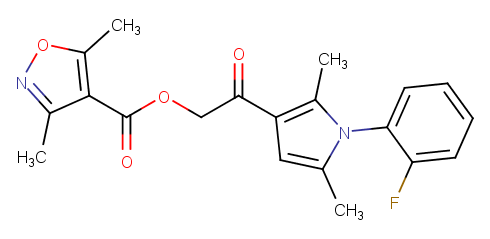

### MMV1046691.png

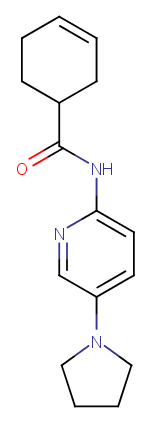

### MMV1047377.png

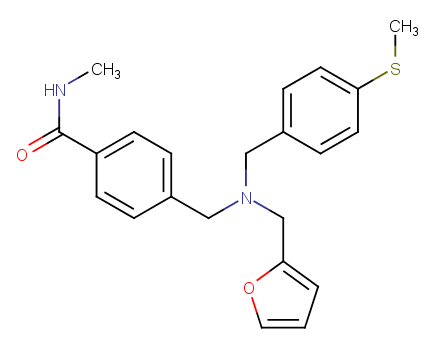

### MMV1047464.png

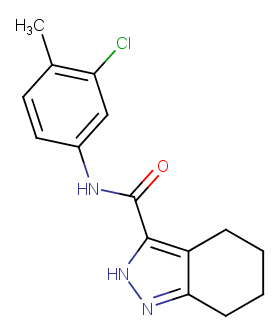

### MMV1048226.png

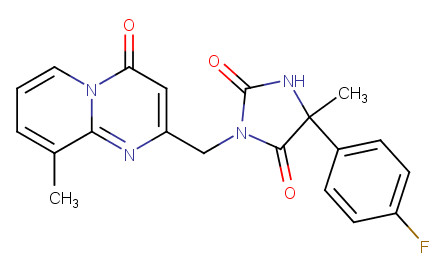

### MMV1048515.png

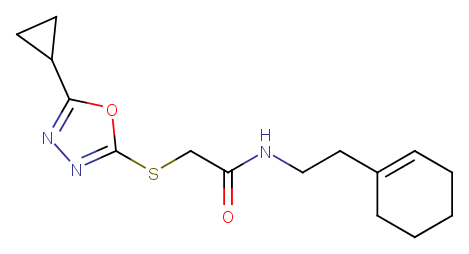

### MMV1048712.png

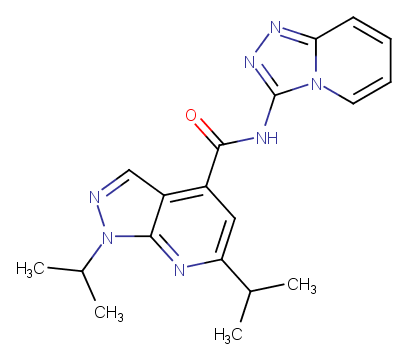

### MMV1048929.png

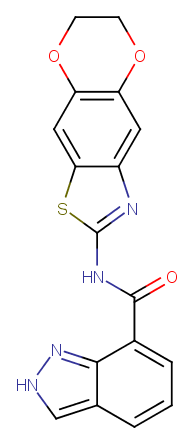

### MMV1049300.png

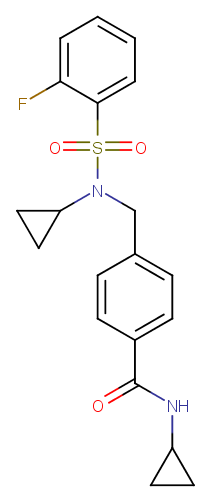

### MMV1049738.png

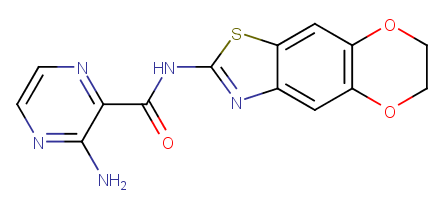

### MMV1050452.png

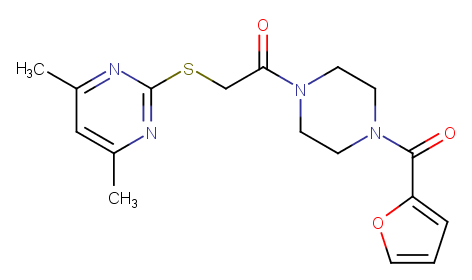

### MMV1050519.png

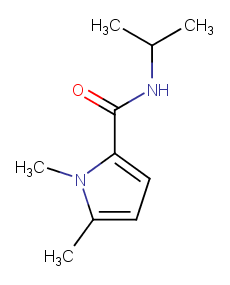
